## Supplementary material for "The marginal brain": Replication package: README.md

Trial-level records for the numerosity-estimation reanalysis are available from the source OSF project, “Endogenous Precision of the Number Sense: Data & Code” (https://osf.io/d6k3m/), associated with Prat-Carrabin and Woodford, “Endogenous precision of the number sense,” eLife 13:RP101277 (2026), https://doi.org/10.7554/eLife.101277.4.

The CSV is not redistributed in this package because file-level reuse terms could not be independently confirmed. Obtain the source file from the OSF project and save it here as published_estimation.csv before running the complete reproduction workflow. The analysis uses anonymized subject IDs and trial-level task responses. It is a secondary reanalysis of the published cohort, not a new or independent human study and not a test of the integrated energy controller.

The analysis used a 787,835-byte source file with SHA-256 2c492fb3a1ba0bfa6e11c8d50e3de759093fbce83e4a81403694221a720e0404 (12,960 trial records). After downloading, verify the file against this fingerprint before reproduction; the campaign-time source manifest records the same checksum.
