## Supplementary material for "The marginal brain": Replication package: analysis_record.md

# Analysis and reproducibility record

## Scope and evidential status

This release combines an analytical resource-allocation model, numerical checks, prospective design simulations, and a secondary reanalysis of a published behavioral dataset. These evidence types are kept separate. The human data describe response-variability associations in the released numerosity-estimation cohort; they do not measure neural energy use, the marginal energy price, latent Fisher precision, or an integrated physiological controller. Simulation rejection rates are design projections under the stated assumptions, not human effects or empirical significance tests.

No preregistration, held-out validation cohort, or cross-validation split is included in the supplied study record. The analysis configuration and seeds make the reported computations reproducible, but do not constitute a preregistration. Cross-validation is not applicable to the descriptive reanalysis and analytical model as implemented here: no individual-level predictive model was trained to estimate held-out human outcomes. This distinction is retained rather than retroactively assigning a confirmatory status to completed analyses.

## Primary computational campaign

The primary campaign was generated from `model/config.json` with master seed 20261006. Its numerical output is `model/results.json`; the recorded run used Python 3.12.14, NumPy 2.5.4 and SciPy 1.18.1. The output stores campaign-time source fingerprint `f8979e7a3eec62e8166af6f09d1c206d8aa347805048fad39d4ef983d8ed1a44` and hashes for 35 source paths. The release also contains a file-by-file SHA-256 manifest. Because the package includes manuscript, figure, build and packaging updates made after the numerical run, these two records serve different purposes: the first identifies the source snapshot associated with the numerical campaign; the second verifies the files distributed now. A targeted Proposition 5 check was subsequently corrected to match the manuscript's estimand, task energy divided by task-plus-competing energy above committed use, excluding committed expenditure. The 16 configured share-price cases were recalculated by independent bisection of the equilibrium residual; the maximum relative errors in the target price and share are both 2.22 × 10⁻¹⁶. This targeted correction does not rerun or alter the other campaign results. The configuration and core numerical modules are retained in the release; the source CSV is an external input and is not included in the curated archive, consistent with `data/README.md`. The delivered numerical estimates and the separately generated matched-support and participant-sign sensitivities are preserved as result files with their analysis scripts and settings.

The primary trial-level family covers 21 shared numerosities (50–70) and three prior-width contrasts, for 63 tests. Participant-cluster bootstrap inference uses 500,000 draws per contrast and numerosity, stratified by the two observed sequence groups. Bonferroni adjustment targets a nominal familywise level of 0.05 conditional on valid individual bootstrap p-values; the adjustment does not establish the finite-sample calibration of the centered bootstrap. The corresponding per-contrast interval level is 99.9206%. The participant-equal sensitivity uses 500,000 draws across its three contrasts (98.3333% per-contrast Bonferroni intervals); the within-block position sensitivity uses 100,000 draws across six contrasts. These are distinct summaries of the same published sample, not independent replications.

| Analysis | Estimate | Reported pattern |
|:--|--:|:--|
| Primary pooled residual variance, width 40 / 20 | Median ratio 2.858 (range 1.978–4.439) | Point estimates above one at 21/21 numerosity levels |
| Primary pooled residual variance, width 60 / 20 | Median ratio 4.251 (range 2.764–6.240) | Point estimates above one at 21/21 levels |
| Primary pooled residual variance, width 60 / 40 | Median ratio 1.447 (range 0.958–2.687) | Point estimates above one at 19/21 levels |
| Participant-equal geometric-mean variance, width 40 / 20 | 2.689 (nominal 98.33% interval 2.299–3.145) | Nominal adjusted p = 0.0000060; sensitivity estimand |
| Participant-equal geometric-mean variance, width 60 / 20 | 3.921 (nominal 98.33% interval 3.209–4.792) | Nominal adjusted p = 0.0000060; sensitivity estimand |
| Participant-equal geometric-mean variance, width 60 / 40 | 1.458 (nominal 98.33% interval 1.179–1.804) | Nominal adjusted p = 0.000066; sensitivity estimand |
| Participant-level aggregate sign, width 40 / 20 | 36/36 positive directions | Exact Bonferroni-adjusted p = 8.73 × 10⁻¹¹ |
| Participant-level aggregate sign, width 60 / 20 | 36/36 positive directions | Exact Bonferroni-adjusted p = 8.73 × 10⁻¹¹ |
| Participant-level aggregate sign, width 60 / 40 | 29/36 positive directions | Exact Bonferroni-adjusted p = 9.38 × 10⁻⁴ |

The participant-equal 60/40 contrast targets a different estimand from the primary fixed-numerosity pooled analysis. The participant sets and available residual degrees of freedom also vary across cells. The main interpretation therefore follows the designated primary family in the analysis code and presents weighting and support checks as sensitivity analyses, not as replacements for the primary result.

The release also contains a reproducible, conditional Gaussian calibration pilot for the centered participant-cluster bootstrap. It generated 100 outer cohorts and used 10,000 inner draws for each of the 63 tests per cohort (seed 20261019; 63 million inner draws). Under this fitted null, 22/100 cohorts had at least one nominal Bonferroni rejection (22.0%; Wilson 95% interval 15.0%–31.1%); the mean unadjusted rejection fraction was 9.06% (cohort-level t interval 8.17%–9.96%). The generator preserves the observed participant-by-cell degrees of freedom, fitted participant–numerosity variance profiles, and sequence strata while imposing equal expected pooled variance across widths. This targeted diagnostic is a model-conditional calibration warning, not a general estimate of the procedure's validity; the finite outer count and Gaussian assumptions are recorded in `model/outputs/conditional_null_calibration/pilot_summary.json`.

## Pair-specific matched-support sensitivity

`model/complete_case_sensitivity.py` performs a separate, post hoc secondary sensitivity analysis. At each numerosity and width pair it retains participants with at least two observations in both conditions, pools within-cell residual sums of squares and degrees of freedom, and resamples participants within the observed sequence strata. The matched set changes by contrast and numerosity. The run uses 500,000 draws for each of 63 tests (31.5 million draws total) and seed 20261013. Its separate output is `model/complete_case_sensitivity.json`; it does not alter `results.json`.

The width 40/20 median ratio is 2.93 (range 2.06–4.74), with point estimates above one at 21/21 levels and median matched support of 28 participants (range 24–32). The width 60/20 median is 4.07 (2.68–6.66), with point estimates above one at 21/21 levels and median support of 20 (13–26). The width 60/40 median is 1.41 (0.80–2.82), with point estimates above one at 17/21 levels and median support of 16 (9–24). This analysis checks sensitivity to cell availability; it is not independent confirmation and does not identify latent precision.

## Prospective simulations and decision rules

The discrimination design simulation uses 200 replications per condition, 20/40/80 participants, 200 trials per state, and specified price ratios of 1, 1.25, 1.5 and 2. The estimation-recovery simulation uses 100 replications per condition, 40 participants and 120 trials per state. The lapse sensitivity uses 200 replications per condition at 40 and 80 participants and raises second-state lapse odds by 50%. Monte Carlo uncertainty is reported with Wilson intervals. For example, under the illustrative doubled-price discrimination condition, the one-sided t-test rejection estimates are 0.805, 0.960 and 0.985 at N=20, 40 and 80; under no price change, the corresponding estimates are 0.040, 0.035 and 0.045. These simulation proportions have finite Monte Carlo precision and are not claims of statistically significant human effects.

## Post hoc participant-level sign-balance sensitivity

`participant_sign_sensitivity.py` adds a deterministic sensitivity analysis to the released 12,960-trial file. For each shared numerosity and width pair, it computes each participant's sample variance in each cell, includes participants with at least two trials in both cells, and compares the variances directly. One-zero pairs are retained as directional signs and equal variance pairs are treated as ties. A two-sided exact binomial sign test is applied to the non-tied positive and negative signs; Bonferroni adjustment is within these 63 numerosity-by-width tests only. The corrected wider-positive counts are 4/21 for 40/20 (numerosities 56, 66, 67 and 70; eligible support 24–32), 1/21 for 60/20 (numerosity 58; support 13–26), and 0/21 for 60/40 (support 9–24). This is a post hoc test of directional balance among eligible participants, not a test of equality of population variances or the primary pooled residual-variance ratio. It uses the same cohort, does not constitute independent replication, and does not calibrate the centered cluster bootstrap. The output records all 63 rows, support, sign and tie counts, exact p-values, adjusted p-values and the source data hash.

When price is unchanged but lapse odds rise by 50%, the shared-lapse fit rejects in 0.775 and 0.950 of simulated studies at N=40 and N=80, compared with 0.050 at both sizes under the state-specific lapse fit. With both price doubling and the lapse change, the state-specific model estimates 0.680 and 0.815 rejection at N=40 and N=80. The results motivate measuring lapses and testing their specification in a future experiment; they do not establish the behavior of every possible lapse process. All lapse fits report optimizer convergence, while boundary contact for at least one fitted parameter is 8.9% for shared-lapse fits and 30.7% for state-specific fits at price ratio 1, and 11.8% and 35.3%, respectively, at price ratio 2 (16,000 fits per model and ratio, from N=80 cohorts used for nested N=40 analyses). The diagnostics aggregate across parameters and do not identify which parameter reached a bound; the nominal state-specific null rejection estimate is therefore specific to this bounded fitting design.

## Data and source integrity

The reanalysis uses 12,960 public trial records from 36 participants and 21 shared numerosity levels. The dataset source, reuse terms and attribution are documented in `data/README.md`. The package manifest records the SHA-256 digest for each distributed file. The CSV digest below is a fingerprint of the local source input used for the recorded analysis, not a digest of a file in the archive; the other two digests identify packaged derived outputs.

- Local source-input fingerprint (CSV not distributed), `data/published_estimation.csv`: `2c492fb3a1ba0bfa6e11c8d50e3de759093fbce83e4a81403694221a720e0404`
- `model/results.json`: `9512971132b1d6b208085d61f42b08ec4d7f00c035abce6b26a69a1da91b1885`
- `model/complete_case_sensitivity.json`: `e07dd7cf00a2bdb52b51f943586088b13c48e082bfd6d09eed3c5476cae21c3e`

The release's automated model suite contains 18 checks. `run_all.py` regenerates the primary model results and model figures, including the participant-clustered reanalysis; the pair-specific complete-case and participant-sign sensitivities are generated by their own scripts. `reproduce.ps1` or `reproduce.sh` orchestrates all three result pipelines, paper figures, TeX/PDF builds and tests. The supplied PDFs are standard LaTeX PDFs and are not tagged for PDF/UA accessibility. Outlet-specific word limits, anonymization, reporting forms and file requirements should be matched when a journal is selected.

## Calibration profile

The encoding model is calibrated at numerosity 60 and predicts the response-variance profile at the other 20 shared numerosity levels by deterministic quadrature. These predictions use no additional fitted parameters, but all 21 levels come from the same published cohort; the comparison is within-cohort and is not an independent-sample validation. The figure-generation script overlays this predicted profile on the matched-numerosity estimates, and the manuscript identifies the calibration anchor in its Results and figure caption.
