## Supplementary material for "The marginal brain": Replication package: complete_case_sensitivity.py

```
"""Secondary, post hoc matched-participant sensitivity for width contrasts.

For each configured numerosity and width pair, this analysis includes only
participants with at least two responses in both widths. It compares pooled
within-participant residual variances on that pair-specific support and uses a
participant-cluster bootstrap stratified by observed block order. The matched
support changes across numerosity and width pair, so this is a secondary
sensitivity analysis rather than a common-population estimand or confirmatory
test.

Run from the model directory with:

    python complete_case_sensitivity.py

The default analysis uses 500,000 draws per contrast-by-numerosity test and
seed 20261013. It writes a standalone JSON alongside this script and never
reads or modifies results.json.
"""
from __future__ import annotations

import argparse
import csv
import hashlib
import itertools
import json
import math
import platform
import sys
import time
from collections import defaultdict
from pathlib import Path

import numpy as np


DEFAULT_REPS = 500_000
DEFAULT_SEED = 20261013
DEFAULT_CHUNK_SIZE = 10_000
ORDER_LABELS = ("Widening", "Narrowing")


def _sha256(path: Path) -> str:
    digest = hashlib.sha256()
    with path.open("rb") as stream:
        for chunk in iter(lambda: stream.read(1024 * 1024), b""):
            digest.update(chunk)
    return digest.hexdigest()


def _load_sufficient_statistics(data_path: Path, widths: tuple[int, ...],
                                numerosities: tuple[int, ...]):
    """Read response cells and return participant order and cell SSE/df."""
    responses = defaultdict(list)
    subject_order = {}
    n_rows = 0
    with data_path.open("r", encoding="utf-8-sig", newline="") as stream:
        reader = csv.DictReader(stream)
        required = {"ID", "Order", "Delta", "Number", "Response"}
        missing = required - set(reader.fieldnames or ())
        if missing:
            raise ValueError(f"Input CSV is missing required columns: {sorted(missing)}")
        for row in reader:
            subject = str(row["ID"])
            order = str(row["Order"]).strip()
            if order not in ORDER_LABELS:
                raise ValueError(f"Unexpected order label for participant {subject}: {order!r}")
            if subject in subject_order and subject_order[subject] != order:
                raise ValueError(f"Order label changes within participant {subject}")
            subject_order[subject] = order
            width = int(float(row["Delta"]))
            numerosity = int(float(row["Number"]))
            if width in widths and numerosity in numerosities:
                responses[(subject, width, numerosity)].append(float(row["Response"]))
            n_rows += 1

    subjects = tuple(sorted(subject_order))
    if not subjects:
        raise ValueError("The input CSV contains no participants")
    stats = {}
    for subject in subjects:
        for width in widths:
            for x in numerosities:
                values = np.asarray(responses.get((subject, width, x), ()), dtype=float)
                df = max(0, int(values.size) - 1)
                sse = float(np.square(values - values.mean()).sum()) if values.size else 0.0
                stats[(subject, width, x)] = (sse, df)
    return subjects, subject_order, stats, n_rows


def _bootstrap_test(sse_a: np.ndarray, df_a: np.ndarray,
                    sse_b: np.ndarray, df_b: np.ndarray,
                    stratum_indices: list[np.ndarray], theta: float,
                    reps: int, rng: np.random.Generator,
                    family_size: int, alpha: float,
                    chunk_size: int):
    """Centered two-sided bootstrap and Bonferroni-dual ratio interval."""
    if reps < 1 or chunk_size < 1:
        raise ValueError("reps and chunk_size must be positive")
    centered = np.empty(reps, dtype=float)
    kept = 0
    attempts = 0
    while kept < reps:
        size = min(chunk_size, reps - kept)
        draw = np.concatenate([
            idx[rng.integers(0, len(idx), size=(size, len(idx)))]
            for idx in stratum_indices
        ], axis=1)
        ss_a = sse_a[draw].sum(axis=1)
        dof_a = df_a[draw].sum(axis=1)
        ss_b = sse_b[draw].sum(axis=1)
        dof_b = df_b[draw].sum(axis=1)
        valid = (ss_a > 0) & (ss_b > 0) & (dof_a > 0) & (dof_b > 0)
        attempts += size
        if np.any(valid):
            values = (np.log(ss_b[valid] / dof_b[valid])
                      - np.log(ss_a[valid] / dof_a[valid]) - theta)
            centered[kept:kept + len(values)] = values
            kept += len(values)

    tail_count = int(np.count_nonzero(np.abs(centered) >= abs(theta)))
    p_value = float((tail_count + 1) / (reps + 1))
    p_adjusted = float(min(1.0, p_value * family_size))

    # Match the finite-replication decision rule used by the primary analysis:
    # invert the plus-one, two-sided Monte Carlo test after Bonferroni correction.
    adjusted_alpha = alpha / family_size
    max_tail_count = int(math.ceil(adjusted_alpha * (reps + 1) - 1.0) - 1)
    if max_tail_count < 0:
        critical = float("inf")
    else:
        order_index = reps - max_tail_count - 1
        critical = float(np.partition(np.abs(centered), order_index)[order_index])
    interval = np.exp([theta - critical, theta + critical])
    significant = bool(p_adjusted <= alpha)
    interval_excludes_one = bool(interval[0] > 1.0 or interval[1] < 1.0)
    return {
        "bootstrap_reps": int(reps),
        "valid_bootstrap_draws": int(reps),
        "attempted_bootstrap_draws": int(attempts),
        "tail_exceedances": tail_count,
        "p_value_centered_two_sided": p_value,
        "p_value_bonferroni": p_adjusted,
        "familywise_95_ci_variance_ratio": [float(interval[0]), float(interval[1])],
        "familywise_ci_log_critical": critical,
        "significant_familywise": significant,
        "familywise_interval_excludes_one": interval_excludes_one,
    }


def analyze(data_path: str | Path, config: dict, reps: int = DEFAULT_REPS,
            seed: int = DEFAULT_SEED, chunk_size: int = DEFAULT_CHUNK_SIZE,
            config_path: str | Path | None = None):
    """Run all fixed-numerosity pair-specific complete-case tests."""
    started = time.perf_counter()
    data_path = Path(data_path).resolve()
    emp_cfg = config["empirical_reanalysis"]
    widths = tuple(int(w) for w in emp_cfg["widths"])
    x_min = int(emp_cfg["shared_number_min"])
    x_max = int(emp_cfg["shared_number_max"])
    numerosities = tuple(range(x_min, x_max + 1))
    alpha = float(emp_cfg["familywise_alpha"])
    pairs = tuple(itertools.combinations(widths, 2))
    family_size = len(numerosities) * len(pairs)
    if not pairs or not numerosities or family_size < 1:
        raise ValueError("Configuration must include at least two widths and one numerosity")

    subjects, subject_order, stats, n_rows = _load_sufficient_statistics(
        data_path, widths, numerosities)
    root_seed = np.random.SeedSequence(int(seed))
    child_seeds = root_seed.spawn(family_size)
    contrast_rows = []
    seed_idx = 0

    for width_a, width_b in pairs:
        for x in numerosities:
            included = [
                subject for subject in subjects
                if stats[(subject, width_a, x)][1] > 0
                and stats[(subject, width_b, x)][1] > 0
            ]
            if not included:
                raise ValueError(f"No pair-complete participants for {width_a}/{width_b} at x={x}")
            local_index = {subject: idx for idx, subject in enumerate(included)}
            order_counts = {
                label: sum(subject_order[s] == label for s in included)
                for label in ORDER_LABELS
            }
            stratum_indices = [
                np.asarray([local_index[s] for s in included if subject_order[s] == label], dtype=int)
                for label in ORDER_LABELS if order_counts[label] > 0
            ]
            sse_a = np.asarray([stats[(s, width_a, x)][0] for s in included], dtype=float)
            df_a = np.asarray([stats[(s, width_a, x)][1] for s in included], dtype=float)
            sse_b = np.asarray([stats[(s, width_b, x)][0] for s in included], dtype=float)
            df_b = np.asarray([stats[(s, width_b, x)][1] for s in included], dtype=float)
            total_sse_a, total_df_a = float(sse_a.sum()), float(df_a.sum())
            total_sse_b, total_df_b = float(sse_b.sum()), float(df_b.sum())
            if min(total_sse_a, total_sse_b, total_df_a, total_df_b) <= 0:
                raise ValueError(f"Non-positive pooled variance support at {width_a}/{width_b}, x={x}")
            variance_a = total_sse_a / total_df_a
            variance_b = total_sse_b / total_df_b
            theta = float(np.log(variance_b / variance_a))
            rng = np.random.default_rng(child_seeds[seed_idx])
            seed_idx += 1
            bootstrap = _bootstrap_test(
                sse_a, df_a, sse_b, df_b, stratum_indices, theta,
                int(reps), rng, family_size, alpha, int(chunk_size))
            if bootstrap["significant_familywise"] != bootstrap["familywise_interval_excludes_one"]:
                raise AssertionError("Bonferroni test and inverted interval disagree")
            contrast_rows.append({
                "width_1": int(width_a),
                "width_2": int(width_b),
                "numerosity": int(x),
                "matched_participants": int(len(included)),
                "matched_participants_by_order": order_counts,
                "residual_df_width_1": int(total_df_a),
                "residual_df_width_2": int(total_df_b),
                "pooled_residual_variance_width_1": float(variance_a),
                "pooled_residual_variance_width_2": float(variance_b),
                "variance_ratio_width_2_over_width_1": float(np.exp(theta)),
                "log_variance_ratio": theta,
                **bootstrap,
            })

    pair_summaries = []
    for width_a, width_b in pairs:
        rows = [r for r in contrast_rows if (r["width_1"], r["width_2"]) == (width_a, width_b)]
        ratios = np.asarray([r["variance_ratio_width_2_over_width_1"] for r in rows])
        participant_counts = np.asarray([r["matched_participants"] for r in rows], dtype=int)
        order_summary = {}
        for label in ORDER_LABELS:
            values = np.asarray([r["matched_participants_by_order"][label] for r in rows], dtype=int)
            order_summary[label] = {
                "median": float(np.median(values)),
                "range": [int(values.min()), int(values.max())],
            }
        significant_x = [r["numerosity"] for r in rows if r["significant_familywise"]]
        pair_summaries.append({
            "width_1": int(width_a),
            "width_2": int(width_b),
            "n_numerosity_tests": len(rows),
            "median_variance_ratio": float(np.median(ratios)),
            "variance_ratio_range": [float(ratios.min()), float(ratios.max())],
            "matched_participants_median": float(np.median(participant_counts)),
            "matched_participants_range": [int(participant_counts.min()), int(participant_counts.max())],
            "matched_participants_by_order": order_summary,
            "significant_numerosities_familywise": int(len(significant_x)),
            "significant_numerosity_values": significant_x,
            "nonsignificant_numerosity_values": [x for x in numerosities if x not in significant_x],
            "median_bonferroni_p_value": float(np.median([r["p_value_bonferroni"] for r in rows])),
            "max_bonferroni_p_value": float(max(r["p_value_bonferroni"] for r in rows)),
            "familywise_ci_excludes_one_count": int(sum(r["familywise_interval_excludes_one"] for r in rows)),
        })

    config_path_resolved = Path(config_path).resolve() if config_path is not None else None
    result = {
        "analysis": "pair_specific_fixed_numerosity_complete_case_sensitivity",
        "status": "secondary_post_hoc_sensitivity_not_confirmatory",
        "metadata": {
            "estimand": (
                "At each fixed numerosity and width pair, log ratio of pooled within-participant "
                "residual variances among participants with at least two observations in both widths; "
                "the participant set may differ by pair and numerosity."
            ),
            "support_rule": "Include participant only when residual df is positive in both widths at that numerosity.",
            "variance_estimator": "sum participant-cell SSE divided by sum participant-cell residual df on shared pair-specific support",
            "resampling": "Participant clusters sampled with replacement jointly across both widths, within observed order strata; stratum sample sizes fixed to matched support.",
            "test": "Centered two-sided cluster-bootstrap tail test on the log variance ratio with plus-one correction.",
            "multiplicity": "Bonferroni correction across all configured width-pair by numerosity tests.",
            "interval": "95% Bonferroni familywise variance-ratio interval inverted from the centered plus-one bootstrap test.",
            "familywise_alpha": alpha,
            "planned_family_size": int(family_size),
            "widths": list(widths),
            "width_pairs": [[int(a), int(b)] for a, b in pairs],
            "numerosities": list(numerosities),
            "n_rows_in_input": int(n_rows),
            "n_participants": int(len(subjects)),
            "observed_order_counts_all_participants": {
                label: int(sum(subject_order[s] == label for s in subjects))
                for label in ORDER_LABELS
            },
            "bootstrap_reps_per_test": int(reps),
            "total_bootstrap_draws_requested": int(reps * family_size),
            "bootstrap_seed": int(seed),
            "seed_stream": "NumPy SeedSequence(seed).spawn(family_size), assigned pair-major then numerosity-major.",
            "bootstrap_chunk_size": int(chunk_size),
            "rng": "numpy.random.default_rng",
            "python_version": platform.python_version(),
            "numpy_version": np.__version__,
            "data_sha256": _sha256(data_path),
            "config_sha256": _sha256(config_path_resolved) if config_path_resolved and config_path_resolved.exists() else None,
            "data_file": data_path.name,
            "config_file": config_path_resolved.name if config_path_resolved else None,
            "caveats": [
                "This is a secondary, post hoc sensitivity analysis; its tests are not confirmatory.",
                "Matched participant support changes across width pairs and numerosities, so the estimand is not a single common-population contrast.",
                "Some order-specific matched supports are small; bootstrap tail inference can be discrete and unstable in those cells.",
                "A positive ratio describes observable within-participant response dispersion and does not by itself identify latent sensory precision or a neural mechanism.",
            ],
        },
        "pairwise_summary": pair_summaries,
        "contrasts": contrast_rows,
    }
    result["metadata"]["runtime_seconds"] = float(time.perf_counter() - started)
    return result


def main(argv: list[str] | None = None):
    model_dir = Path(__file__).resolve().parent
    parser = argparse.ArgumentParser(description=__doc__)
    parser.add_argument("--config", type=Path, default=model_dir / "config.json")
    parser.add_argument("--data", type=Path, default=None,
                        help="CSV path; default comes from empirical_reanalysis.data_path in config.")
    parser.add_argument("--output", type=Path, default=model_dir / "complete_case_sensitivity.json")
    parser.add_argument("--reps", type=int, default=DEFAULT_REPS,
                        help=f"Bootstrap draws per test (default {DEFAULT_REPS}).")
    parser.add_argument("--seed", type=int, default=DEFAULT_SEED,
                        help=f"Fixed bootstrap seed (default {DEFAULT_SEED}).")
    parser.add_argument("--chunk-size", type=int, default=DEFAULT_CHUNK_SIZE)
    args = parser.parse_args(argv)

    config_path = args.config.resolve()
    config = json.loads(config_path.read_text(encoding="utf-8"))
    data_path = args.data.resolve() if args.data else (config_path.parent / config["empirical_reanalysis"]["data_path"]).resolve()
    result = analyze(data_path, config, reps=args.reps, seed=args.seed,
                     chunk_size=args.chunk_size, config_path=config_path)
    output = args.output.resolve()
    output.parent.mkdir(parents=True, exist_ok=True)
    output.write_text(json.dumps(result, indent=2, sort_keys=True) + "\n", encoding="utf-8")
    print(json.dumps({
        "output": str(output),
        "tests": len(result["contrasts"]),
        "reps_per_test": args.reps,
        "total_draws": result["metadata"]["total_bootstrap_draws_requested"],
        "runtime_seconds": result["metadata"]["runtime_seconds"],
        "significant_by_pair": [
            {"pair": f"{r['width_1']}/{r['width_2']}",
             "count": r["significant_numerosities_familywise"],
             "total": r["n_numerosity_tests"]}
            for r in result["pairwise_summary"]
        ],
    }, indent=2))


if __name__ == "__main__":
    main()
```
