## Supplementary material for "The marginal brain": Replication package: conditional_null_calibration.py

```
#!/usr/bin/env python3
"""Conditional null-size pilot for the primary pooled variance-ratio tests.

This file is intentionally isolated from the release source. It imports the
release's centered participant-cluster bootstrap helper so the inner test
matches the implementation under review.

Null model: retain the observed participant-by-numerosity cell degrees of
freedom and participant variance profiles. Within each numerosity, multiply
the participant-specific baseline variance by a width-specific factor chosen
so the expected pooled available-cell variance is the same at all three
widths. Generate independent Gaussian within-cell residual sums of squares,
SSE[i,x,w] = sigma2[i,x,w] * chi-square(df[i,x,w]). The factors correct for
the different participant/df mixes across widths. This is a parametric,
model-conditional calibration, not a general validity claim.

Outer cohorts resample participant design/variance profiles with replacement
within the observed Widening (20) and Narrowing (16) strata. The same sampled
profile is retained across all numerosities and widths in each cohort.
"""

from __future__ import annotations

import argparse
import csv
import hashlib
import json
import math
import platform
import sys
import time
from pathlib import Path

import numpy as np
import scipy


HERE = Path(__file__).resolve().parent
REPLICATION = HERE.parent
DATA_DEFAULT = REPLICATION / "data" / "published_estimation.csv"
RESULTS = HERE / "results.json"
EMPIRICAL_SOURCE = HERE / "spt" / "empirical.py"
WIDTHS = (20, 40, 60)
NUMEROSITIES = tuple(range(50, 71))
ORDER_LABELS = ("Widening", "Narrowing")
FAMILY_SIZE = len(NUMEROSITIES) * 3
ALPHA = 0.05


def sha256(path: Path) -> str:
    h = hashlib.sha256()
    with path.open("rb") as f:
        for block in iter(lambda: f.read(1 << 20), b""):
            h.update(block)
    return h.hexdigest()


def import_release_bootstrap():
    sys.path.insert(0, str(HERE))
    from spt.empirical import _cluster_bootstrap_log_ratio  # type: ignore
    return _cluster_bootstrap_log_ratio


def observed_cells(data_path: Path):
    cells: dict[tuple[str, int, int], list[float]] = {}
    orders: dict[str, str] = {}
    block_trials: dict[tuple[str, int], list[int]] = {}
    first_trial: dict[tuple[str, int], int] = {}
    with data_path.open("r", encoding="utf-8-sig", newline="") as f:
        for row in csv.DictReader(f):
            subject = str(row["ID"])
            width = int(float(row["Delta"]))
            x = int(float(row["Number"]))
            response = float(row["Response"])
            order = str(row["Order"]).strip()
            tloc = int(float(row["tLoc"]))
            tall = int(float(row["tAll"]))
            if subject in orders and orders[subject] != order:
                raise ValueError(f"Order changes within participant {subject}")
            orders[subject] = order
            block_trials.setdefault((subject, width), []).append(tloc)
            first_trial[(subject, width)] = min(tall, first_trial.get((subject, width), tall))
            if width in WIDTHS and x in NUMEROSITIES:
                cells.setdefault((subject, width, x), []).append(response)

    ids = sorted(orders)
    if len(ids) != 36:
        raise ValueError(f"Expected 36 participants, found {len(ids)}")
    for (subject, width), trials in block_trials.items():
        if len(trials) != 120 or sorted(trials) != list(range(1, 121)):
            raise ValueError(f"Invalid 120-trial block for {subject}, width {width}")
        order = orders[subject]
        expected = tuple(sorted(WIDTHS)) if order == "Widening" else tuple(sorted(WIDTHS, reverse=True))
        actual = tuple(w for _, w in sorted((first_trial[(subject, w)], w) for w in WIDTHS))
        if actual != expected:
            raise ValueError(f"Order sequence mismatch for {subject}: {actual} vs {expected}")

    subject_index = {s: i for i, s in enumerate(ids)}
    x_index = {x: i for i, x in enumerate(NUMEROSITIES)}
    width_index = {w: i for i, w in enumerate(WIDTHS)}
    sse = np.zeros((len(ids), len(NUMEROSITIES), len(WIDTHS)), dtype=float)
    df = np.zeros_like(sse, dtype=np.int64)
    trials = np.zeros_like(sse, dtype=np.int64)
    for (subject, width, x), values in cells.items():
        if len(values) < 2:
            continue
        vals = np.asarray(values, dtype=float)
        centered = vals - vals.mean()
        i, j, k = subject_index[subject], x_index[x], width_index[width]
        sse[i, j, k] = float(centered @ centered)
        df[i, j, k] = len(vals) - 1
        trials[i, j, k] = len(vals)

    order_array = np.asarray([orders[s] for s in ids], dtype=object)
    strata = [np.flatnonzero(order_array == label) for label in ORDER_LABELS]
    if tuple(len(z) for z in strata) != (20, 16):
        raise ValueError(f"Unexpected sequence counts: {tuple(len(z) for z in strata)}")
    return ids, order_array, strata, sse, df, trials


def fit_null_profile(sse: np.ndarray, df: np.ndarray, strata: list[np.ndarray]):
    """Fit participant-x base variance and scale widths to a pooled null.

    The baseline participant variance is the within-participant pooled SSE/df
    across widths at that numerosity. The null width scale is then chosen from
    the empirical profile population (20/16 stratum weights) so the ratio of
    expected pooled SSE to expected pooled df equals one across widths.
    """
    n_x, n_w = sse.shape[1], sse.shape[2]
    tau2 = np.zeros((sse.shape[0], n_x), dtype=float)
    np.divide(sse.sum(axis=2), df.sum(axis=2), out=tau2, where=df.sum(axis=2) > 0)
    scales = np.ones((n_x, n_w), dtype=float)
    target = np.zeros(n_x, dtype=float)
    null_means = np.zeros((n_x, n_w), dtype=float)
    for xj in range(n_x):
        numer = np.zeros(n_w, dtype=float)
        denom = np.zeros(n_w, dtype=float)
        for idx in strata:
            n = len(idx)
            numer += n * (df[idx, xj, :] * tau2[idx, xj, None]).mean(axis=0)
            denom += n * df[idx, xj, :].mean(axis=0)
        if np.any(denom <= 0) or np.any(numer <= 0):
            raise ValueError(f"No positive null variance/support at x={NUMEROSITIES[xj]}")
        null_means[xj] = numer / denom
        target[xj] = float(np.exp(np.mean(np.log(null_means[xj]))))
        scales[xj] = target[xj] / null_means[xj]
        if not np.allclose(scales[xj] * null_means[xj], target[xj], rtol=1e-12, atol=1e-12):
            raise AssertionError("Fitted null scales failed equal-variance constraint")
    return tau2, scales, target, null_means


def draw_outer_cohort(rng: np.random.Generator, strata: list[np.ndarray],
                      tau2: np.ndarray, scales: np.ndarray,
                      df_templates: np.ndarray):
    selected = np.concatenate([
        idx[rng.integers(0, len(idx), size=len(idx))] for idx in strata
    ])
    cohort_df = df_templates[selected].copy()
    sigma2 = tau2[selected, :, None] * scales[None, :, :]
    cohort_sse = np.zeros_like(sigma2)
    nonzero = cohort_df > 0
    cohort_sse[nonzero] = sigma2[nonzero] * rng.chisquare(cohort_df[nonzero])
    cohort_strata = [np.arange(0, len(strata[0])), np.arange(len(strata[0]), len(selected))]
    return selected, cohort_sse, cohort_df, cohort_strata


def monte_carlo_interval(successes: int, trials: int, alpha: float = 0.05):
    """Wilson score interval without adding a scipy dependency to output code."""
    from scipy.stats import norm
    z = float(norm.ppf(1 - alpha / 2))
    p = successes / trials
    den = 1 + z * z / trials
    center = (p + z * z / (2 * trials)) / den
    half = z * math.sqrt(p * (1 - p) / trials + z * z / (4 * trials * trials)) / den
    return [max(0.0, center - half), min(1.0, center + half)]


def self_test(bootstrap_fn):
    # Identical participant-level variance contributions imply theta=0 and
    # every stratified bootstrap replicate remains exactly centered at zero.
    toy_sse = np.asarray([[2.0, 2.0], [4.0, 4.0], [6.0, 6.0], [8.0, 8.0]])
    toy_df = np.full((4, 2), 2.0)
    theta = float(np.log(toy_sse[:, 1].sum() / toy_df[:, 1].sum()) -
                  np.log(toy_sse[:, 0].sum() / toy_df[:, 0].sum()))
    assert theta == 0.0
    toy = bootstrap_fn(toy_sse, toy_df, 0, 1, theta, 2_000, 1234, 1,
                       strata_indices=[np.asarray([0, 1]), np.asarray([2, 3])])
    assert toy["pvalue_bootstrap"] == 1.0
    # Confirm the DGP primitive has the specified chi-square mean.
    rng = np.random.default_rng(8675309)
    sample = rng.chisquare(4, size=100_000)
    assert abs(float(sample.mean()) - 4.0) < 0.05
    # A balanced Gaussian null is a small end-to-end size check for the
    # imported release bootstrap. It is deliberately a toy, not an estimate
    # of size under the release's unequal, sparse cell design.
    balanced_rng = np.random.default_rng(314159)
    balanced_df = np.full((36, 2), 8.0)
    balanced_strata = [np.arange(20), np.arange(20, 36)]
    balanced_rejects = 0
    balanced_p = []
    balanced_outer, balanced_inner = 300, 2_000
    balanced_seed_seq = np.random.SeedSequence(271828).spawn(balanced_outer)
    for draw_i, child in enumerate(balanced_seed_seq):
        balanced_sse = balanced_rng.chisquare(8, size=(36, 2))
        balanced_theta = float(
            np.log(balanced_sse[:, 1].sum() / balanced_df[:, 1].sum())
            - np.log(balanced_sse[:, 0].sum() / balanced_df[:, 0].sum())
        )
        child_words = child.generate_state(2, dtype=np.uint32)
        toy_result = bootstrap_fn(
            balanced_sse, balanced_df, 0, 1, balanced_theta, balanced_inner,
            int(child_words[0]) ^ (int(child_words[1]) << 1), 1,
            strata_indices=balanced_strata,
        )
        p = float(toy_result["pvalue_bootstrap"])
        balanced_p.append(p)
        balanced_rejects += int(p < 0.05)
    return {"toy_equal_width_ratio": theta, "toy_bootstrap_pvalue": toy["pvalue_bootstrap"],
            "chi_square_df4_simulated_mean": float(sample.mean()), "chi_square_df4_expected_mean": 4.0,
            "balanced_gaussian_null_size_check": {
                "outer_cohorts": balanced_outer,
                "inner_draws": balanced_inner,
                "strata_counts": [20, 16],
                "raw_alpha": 0.05,
                "rejections": balanced_rejects,
                "rejection_rate": balanced_rejects / balanced_outer,
                "wilson_95": monte_carlo_interval(balanced_rejects, balanced_outer),
                "mean_bootstrap_pvalue": float(np.mean(balanced_p)),
                "note": "balanced two-width toy validation only; not evidence about the empirical unequal-support design",
            }}


def check_against_release(sse: np.ndarray, df: np.ndarray):
    with RESULTS.open("r", encoding="utf-8") as f:
        results = json.load(f)
    primary = results["empirical_reanalysis"]["per_numerosity_cluster_bootstrap"]
    if primary["n_clusters"] != sse.shape[0] or primary["planned_family_size"] != FAMILY_SIZE:
        raise ValueError("Release result metadata does not match reconstructed design")
    max_v_diff, max_df_diff = 0.0, 0
    for xj, row in enumerate(primary["per_numerosity"]):
        pooled_df = df[:, xj, :].sum(axis=0)
        pooled_v = sse[:, xj, :].sum(axis=0) / pooled_df
        for wj, width in enumerate(WIDTHS):
            stored_v = row["pooled_variance_by_width"][str(width)]
            stored_df = row["residual_df_by_width"][str(width)]
            max_v_diff = max(max_v_diff, abs(float(pooled_v[wj]) - float(stored_v)))
            max_df_diff = max(max_df_diff, abs(int(pooled_df[wj]) - int(stored_df)))
    if max_v_diff > 1e-10 or max_df_diff != 0:
        raise AssertionError(f"Reconstructed design differs from release JSON: V={max_v_diff}, df={max_df_diff}")
    return {"n_clusters": int(primary["n_clusters"]),
            "sequence_counts": primary["sequence_counts"],
            "family_size": int(primary["planned_family_size"]),
            "release_inner_reps": int(primary["bootstrap_reps_per_contrast"]),
            "max_abs_pooled_variance_difference": max_v_diff,
            "max_abs_df_difference": max_df_diff}


def run(outer: int, inner: int, seed: int, output_dir: Path, data_path: Path):
    bootstrap_fn = import_release_bootstrap()
    tests = self_test(bootstrap_fn)
    ids, order_array, strata, observed_sse, observed_df, trials = observed_cells(data_path)
    release_check = check_against_release(observed_sse, observed_df)
    tau2, scales, target, null_means = fit_null_profile(observed_sse, observed_df, strata)

    root_rng = np.random.default_rng(seed)
    outer_seeds = root_rng.integers(0, 2**32 - 1, size=outer, dtype=np.uint64)
    test_rows = []
    rep_family = np.zeros(outer, dtype=bool)
    rep_raw_rate = np.zeros(outer, dtype=float)
    rep_bonf_rate = np.zeros(outer, dtype=float)
    rep_mean_p = np.zeros(outer, dtype=float)
    started = time.perf_counter()

    for r, outer_seed in enumerate(outer_seeds):
        rng = np.random.default_rng(int(outer_seed))
        selected, sse_star, df_star, inner_strata = draw_outer_cohort(
            rng, strata, tau2, scales, observed_df
        )
        # Separate child streams mirror the release code's separate bootstrap
        # draw stream for every numerosity-by-width contrast.
        contrast_seeds = np.random.SeedSequence(int(outer_seed) ^ 0x6A09E667).spawn(FAMILY_SIZE)
        seed_i = 0
        raw_rejects = 0
        bonf_rejects = 0
        pvals = []
        for xj, x in enumerate(NUMEROSITIES):
            ss = sse_star[:, xj, :]
            dd = df_star[:, xj, :]
            pooled_df = dd.sum(axis=0)
            pooled_sse = ss.sum(axis=0)
            if np.any(pooled_df <= 0) or np.any(pooled_sse <= 0):
                raise RuntimeError(f"Degenerate simulated cohort at replication {r}, x={x}")
            pooled_v = pooled_sse / pooled_df
            for i, j in ((0, 1), (0, 2), (1, 2)):
                theta = float(np.log(pooled_v[j] / pooled_v[i]))
                seed_words = contrast_seeds[seed_i].generate_state(2, dtype=np.uint32)
                inner_seed = int(seed_words[0]) ^ (int(seed_words[1]) << 1)
                seed_i += 1
                summary = bootstrap_fn(
                    ss, dd, i, j, theta, inner, inner_seed, FAMILY_SIZE,
                    strata_indices=inner_strata, alpha=ALPHA
                )
                p = float(summary["pvalue_bootstrap"])
                p_bonf = min(1.0, p * FAMILY_SIZE)
                is_raw = p < ALPHA
                is_bonf = p_bonf < ALPHA
                raw_rejects += int(is_raw)
                bonf_rejects += int(is_bonf)
                pvals.append(p)
                test_rows.append({
                    "outer_replication": r + 1,
                    "numerosity": x,
                    "width_1": WIDTHS[i],
                    "width_2": WIDTHS[j],
                    "log_variance_ratio": theta,
                    "pvalue_centered_cluster_bootstrap": p,
                    "pvalue_bonferroni_63": p_bonf,
                    "reject_raw_0_05": int(is_raw),
                    "reject_bonferroni_0_05": int(is_bonf),
                })
        rep_family[r] = bonf_rejects > 0
        rep_raw_rate[r] = raw_rejects / FAMILY_SIZE
        rep_bonf_rate[r] = bonf_rejects / FAMILY_SIZE
        rep_mean_p[r] = float(np.mean(pvals))
        if (r + 1) % max(1, outer // 10) == 0:
            elapsed = time.perf_counter() - started
            print(f"completed {r + 1}/{outer} cohorts; elapsed {elapsed:.1f} s", flush=True)

    runtime = time.perf_counter() - started
    output_dir.mkdir(parents=True, exist_ok=True)
    test_csv = output_dir / "pilot_test_results.csv"
    with test_csv.open("w", encoding="utf-8", newline="") as f:
        writer = csv.DictWriter(f, fieldnames=list(test_rows[0]))
        writer.writeheader()
        writer.writerows(test_rows)

    family_hits = int(rep_family.sum())
    # Cohort-level t intervals preserve within-cohort dependence across the 63 tests.
    from scipy.stats import t as student_t
    bonf_mean = float(rep_bonf_rate.mean())
    bonf_se = float(rep_bonf_rate.std(ddof=1) / math.sqrt(outer))
    tcrit = float(student_t.ppf(0.975, outer - 1)) if outer > 1 else float("nan")
    raw_mean = float(rep_raw_rate.mean())
    raw_se = float(rep_raw_rate.std(ddof=1) / math.sqrt(outer))
    p_mean = float(rep_mean_p.mean())
    summary = {
        "title": "Conditional Gaussian null-size pilot for primary pooled available-cell variance ratios",
        "status": "pilot only; no claim of general finite-sample validity",
        "estimand": "theta[x,w2/w1] = log{(sum_i SSE[i,x,w2] / sum_i df[i,x,w2]) / (sum_i SSE[i,x,w1] / sum_i df[i,x,w1])}; cells with fewer than two trials contribute zero SSE and df",
        "null": "for every x and width contrast, the population ratio of expected pooled available-cell residual variance is 1; width scales correct for unequal df/participant composition",
        "null_generator": "participant-x baseline variances fitted by pooled SSE/df across widths; width-specific scale factors make expected pooled SSE / expected pooled df equal at all widths; independent Gaussian within-cell residual SSE via sigma2 * chi-square(df)",
        "outer_sampling": "empirical participant design/variance profiles resampled with replacement within observed sequence strata; each selected profile retained across all 21 numerosities and widths",
        "inner_test": "release _cluster_bootstrap_log_ratio: centered, two-sided, participant-cluster bootstrap stratified by Widening/Narrowing; plus-one p-values; Bonferroni family size 63",
        "pilot": {"outer_cohorts": outer, "inner_bootstrap_draws_per_test": inner,
                  "tests_per_cohort": FAMILY_SIZE, "familywise_alpha": ALPHA,
                  "inner_mc_min_p": 1.0 / (inner + 1), "seed": seed,
                  "runtime_seconds": runtime,
                  "inner_draws_total": outer * FAMILY_SIZE * inner},
        "rejection_summary": {
            "familywise_any_bonferroni_rejection_count": family_hits,
            "familywise_any_bonferroni_rejection_rate": family_hits / outer,
            "familywise_any_bonferroni_rejection_wilson_95": monte_carlo_interval(family_hits, outer),
            "mean_raw_rejection_fraction_across_63_tests": raw_mean,
            "mean_raw_rejection_fraction_cohort_t_95": [max(0.0, raw_mean - tcrit * raw_se), min(1.0, raw_mean + tcrit * raw_se)],
            "mean_bonferroni_rejection_fraction_across_63_tests": bonf_mean,
            "mean_bonferroni_rejection_fraction_cohort_t_95": [max(0.0, bonf_mean - tcrit * bonf_se), min(1.0, bonf_mean + tcrit * bonf_se)],
            "nominal_per_test_bonferroni_threshold": ALPHA / FAMILY_SIZE,
            "mean_bootstrap_pvalue_across_tests": p_mean,
            "pvalue_grid_resolution": 1.0 / (inner + 1),
        },
        "null_scale_diagnostics": {
            "target_expected_pooled_variance_by_numerosity": {str(x): float(target[j]) for j, x in enumerate(NUMEROSITIES)},
            "unadjusted_expected_pooled_variance_by_numerosity_and_width": {str(x): {str(w): float(null_means[j, k]) for k, w in enumerate(WIDTHS)} for j, x in enumerate(NUMEROSITIES)},
            "width_scale_factors_by_numerosity": {str(x): {str(w): float(scales[j, k]) for k, w in enumerate(WIDTHS)} for j, x in enumerate(NUMEROSITIES)},
            "min_width_scale_factor": float(scales.min()),
            "max_width_scale_factor": float(scales.max()),
            "max_expected_width_difference_after_scaling": float(np.max(np.abs(scales * null_means - target[:, None]))),
        },
        "input_validation": release_check,
        "toy_validation": tests,
        "input_sha256": {"published_estimation.csv": sha256(data_path), "empirical.py": sha256(EMPIRICAL_SOURCE)},
        "simulation_script_sha256": sha256(Path(__file__).resolve()),
        "environment": {"python": platform.python_version(), "numpy": np.__version__, "scipy": scipy.__version__},
        "output_csv": test_csv.name,
        "limitations": [
            "Gaussian cell residuals and the fitted common participant-x variance profile are assumptions, not identified by the participant-level SSE/df summaries.",
            "This pilot targets the pooled available-cell estimand and empirical sequence-stratified profile population; it does not calibrate the participant-equal sensitivity or establish general validity.",
            "Only 100 outer cohorts and 10,000 inner draws were run. Familywise-rate uncertainty is wide, and the Bonferroni cutoff is near the inner-bootstrap tail resolution.",
        ],
    }
    (output_dir / "pilot_summary.json").write_text(json.dumps(summary, indent=2) + "\n", encoding="utf-8")
    return summary


def main():
    parser = argparse.ArgumentParser()
    parser.add_argument("--outer", type=int, default=100)
    parser.add_argument("--inner", type=int, default=10_000)
    parser.add_argument("--seed", type=int, default=20261019)
    parser.add_argument("--data", type=Path, default=DATA_DEFAULT,
                        help="path to the published estimation CSV obtained from the source OSF project")
    parser.add_argument("--output-dir", type=Path,
                        default=HERE / "outputs" / "conditional_null_calibration")
    parser.add_argument("--validation-only", action="store_true",
                        help="run code/toy checks without the empirical-design calibration")
    args = parser.parse_args()
    if args.validation_only:
        validation = self_test(import_release_bootstrap())
        args.output_dir.mkdir(parents=True, exist_ok=True)
        (args.output_dir / "toy_validation.json").write_text(
            json.dumps(validation, indent=2) + "\n", encoding="utf-8"
        )
        print(json.dumps(validation, indent=2))
        return
    if not args.data.is_file():
        parser.error(f"trial-level input not found at {args.data}; see ../data/README.md")
    summary = run(args.outer, args.inner, args.seed, args.output_dir, args.data)
    print(json.dumps(summary["rejection_summary"], indent=2))
    print(f"runtime_seconds={summary['pilot']['runtime_seconds']:.3f}")
    print(f"outputs={args.output_dir}")


if __name__ == "__main__":
    main()
```
