## Supplementary material for "The marginal brain": Replication package: fill_numbers.py

```
#!/usr/bin/env python3
"""Fill the numbers of a manuscript from the model's results, so that every number comes from code.

Usage
  python3 fill_numbers.py body.md results.json body_filled.md
  python3 fill_numbers.py body.md results.json complete_case_sensitivity.json body_filled.md
  python3 fill_numbers.py body.md results.json complete_case_sensitivity.json participant_sign_sensitivity.json body_filled.md
  python3 fill_numbers.py --latex numbers.tex results.json
  Any of the above accepts --extra=other_results.json to merge further results files.

Placeholders in the markdown source take the form {{key}} or {{key:spec}}, where spec is a
Python format specification: {{gamma_c:.2f}}, {{n_agents:,}}, {{share:.1%}}, {{p:.2g}}.
Nested keys use dots: {{fit.alpha:.3f}}. Unknown keys are left in place and reported, and the
script exits with status 1 so that a build cannot silently skip them. In LaTeX mode, every
scalar in results.json becomes a macro: fit.alpha becomes \\numFitAlpha, t1 becomes \\numTOne;
floats are written with three significant digits unless the results file stores a string.
"""
import json
import re
import sys

DIGITS = "Zero One Two Three Four Five Six Seven Eight Nine".split()


def flatten(d, prefix=""):
    out = {}
    for k, v in d.items():
        key = f"{prefix}{k}"
        if isinstance(v, dict):
            out.update(flatten(v, key + "."))
        else:
            out[key] = v
    return out


def macro_name(key):
    parts = re.split(r"[^A-Za-z0-9]+", key)
    name = "".join(p[:1].upper() + p[1:] for p in parts if p)
    return "\\num" + "".join(DIGITS[int(c)] if c.isdigit() else c for c in name)


def fmt(value, spec):
    if spec:
        return format(value, spec)
    if isinstance(value, float):
        return format(value, ".3g")
    return str(value)


def main(argv):
    if len(argv) == 3 and argv[0] == "--latex":
        values = flatten(json.load(open(argv[2], encoding="utf-8")))
        lines = ["% Generated from " + argv[2] + " by fill_numbers.py; do not edit by hand."]
        for k, v in sorted(values.items()):
            if isinstance(v, (int, float, str)) and not isinstance(v, bool):
                lines.append(f"\\newcommand{{{macro_name(k)}}}{{{fmt(v, '')}}}  % {k}")
        open(argv[1], "w", encoding="utf-8").write("\n".join(lines) + "\n")
        print(f"Wrote {len(lines) - 1} macros to {argv[1]}")
        return 0
    extras = [a.split("=", 1)[1] for a in argv if a.startswith("--extra=")]
    argv = [a for a in argv if not a.startswith("--extra=")]
    if len(argv) not in (3, 4, 5):
        print(__doc__)
        return 2
    sign_values = None
    if len(argv) == 5:
        source_path, results_path, sensitivity_path, sign_path, output_path = argv
        sensitivity_values = json.load(open(sensitivity_path, encoding="utf-8"))
        sign_values = json.load(open(sign_path, encoding="utf-8"))
    elif len(argv) == 4:
        source_path, results_path, sensitivity_path, output_path = argv
        sensitivity_values = json.load(open(sensitivity_path, encoding="utf-8"))
    else:
        source_path, results_path, output_path = argv
        sensitivity_values = None
    text = open(source_path, encoding="utf-8").read()
    raw_values = json.load(open(results_path, encoding="utf-8"))
    values = flatten(raw_values)
    for extra_path in extras:  # additional results files, e.g. model_predicted_ratios.json
        values.update(flatten(json.load(open(extra_path, encoding="utf-8"))))
    # Express median variance ratios as relative increases for reader-facing summaries.
    for contrast in ("20_40", "20_60", "40_60"):
        key = f"headline.empirical_variance_ratio_{contrast}"
        if key in values:
            values[f"headline.empirical_variance_ratio_{contrast}_increase_pct"] = (
                float(values[key]) - 1.0
            ) * 100.0
    if sign_values is not None:
        for row in sign_values["summary"]:
            contrast = row["contrast"].replace("/", "_")
            values[f"sign_corrected_{contrast}"] = int(row["corrected_numerosities"])
    if "{{complete_case_sensitivity_rows}}" in text and sensitivity_values is not None:
        rows = []
        for row in sorted(sensitivity_values["pairwise_summary"],
                          key=lambda item: (item["width_2"], item["width_1"])):
            n_lo, n_hi = row["matched_participants_range"]
            r_lo, r_hi = row["variance_ratio_range"]
            ratios = [
                item["variance_ratio_width_2_over_width_1"]
                for item in sensitivity_values["contrasts"]
                if item["width_1"] == row["width_1"]
                and item["width_2"] == row["width_2"]
            ]
            above_one = sum(value > 1.0 for value in ratios)
            rows.append(
                f"| {row['width_2']} / {row['width_1']} "
                f"| {row['matched_participants_median']:.0f} ({n_lo}–{n_hi}) "
                f"| {row['median_variance_ratio']:.2f} ({r_lo:.2f}–{r_hi:.2f}) "
                f"| {above_one}/{len(ratios)} |"
            )
        text = text.replace("{{complete_case_sensitivity_rows}}", "\n".join(rows))
    if "{{complete_case_sensitivity_summary}}" in text and sensitivity_values is not None:
        summaries = {}
        for row in sensitivity_values["pairwise_summary"]:
            key = (int(row["width_1"]), int(row["width_2"]))
            n_lo, n_hi = row["matched_participants_range"]
            r_lo, r_hi = row["variance_ratio_range"]
            ratios = [
                item["variance_ratio_width_2_over_width_1"]
                for item in sensitivity_values["contrasts"]
                if item["width_1"] == row["width_1"]
                and item["width_2"] == row["width_2"]
            ]
            above_one = sum(value > 1.0 for value in ratios)
            summaries[key] = (
                f"{row['median_variance_ratio']:.2f} for {key[1]}/{key[0]} "
                f"(range across numerosity {r_lo:.2f}–{r_hi:.2f}; point estimates above one at "
                f"{above_one}/{len(ratios)} levels), "
                f"with median matched support {row['matched_participants_median']:.0f} "
                f"({n_lo}–{n_hi})"
            )
        ordered = [summaries[(20, 40)], summaries[(20, 60)], summaries[(40, 60)]]
        text = text.replace(
            "{{complete_case_sensitivity_summary}}",
            "Its median pooled variance ratios are " + "; ".join(ordered[:2])
            + "; and " + ordered[2] + ".",
        )
    if "{{participant_sign_sensitivity_summary}}" in text and sign_values is not None:
        summaries = {row["contrast"]: row for row in sign_values["summary"]}
        pieces = []
        for contrast in ("40/20", "60/20", "60/40"):
            row = summaries[contrast]
            levels = row["corrected_numerosity_values"]
            level_text = (
                " at numerosity " + ", ".join(str(x) for x in levels)
                if levels else ""
            )
            pieces.append(
                f"{row['corrected_numerosities']}/{row['family_size']} for {contrast}{level_text} "
                f"(eligible support {row['eligible_participant_range'][0]}–"
                f"{row['eligible_participant_range'][1]})"
            )
        text = text.replace(
            "{{participant_sign_sensitivity_summary}}",
            "; ".join(pieces),
        )
    if "{{aggregate_participant_sign_summary}}" in text and sign_values is not None:
        aggregate = sign_values["aggregate_participant_direction"]["contrasts"]
        summaries = {row["contrast"]: row for row in aggregate}
        pieces = []
        for contrast in ("40/20", "60/20", "60/40"):
            row = summaries[contrast]
            pieces.append(
                f"{row['n_positive_wider_variance']}/{row['n_eligible']} participants for "
                f"{contrast}"
            )
        text = text.replace("{{aggregate_participant_sign_summary}}", "; ".join(pieces))
    if "{{aggregate_participant_sign_p_summary}}" in text and sign_values is not None:
        aggregate = sign_values["aggregate_participant_direction"]["contrasts"]
        summaries = {row["contrast"]: row for row in aggregate}
        pieces = [
            f"{summaries[contrast]['bonferroni_adjusted_p_3']:.2g}"
            for contrast in ("40/20", "60/20", "60/40")
        ]
        text = text.replace("{{aggregate_participant_sign_p_summary}}", ", ".join(pieces))
    if "{{aggregate_participant_sign_recap}}" in text and sign_values is not None:
        aggregate = sign_values["aggregate_participant_direction"]["contrasts"]
        summaries = {row["contrast"]: row for row in aggregate}
        pieces = []
        for contrast in ("40/20", "60/20", "60/40"):
            row = summaries[contrast]
            pieces.append(
                f"{row['n_positive_wider_variance']}/{row['n_eligible']} for {contrast}"
            )
        text = text.replace("{{aggregate_participant_sign_recap}}", ", ".join(pieces))
    if "{{participant_sign_sensitivity_rows}}" in text and sign_values is not None:
        rows = []
        for row in sign_values["contrasts"]:
            adjusted = row["bonferroni_adjusted_p_63"]
            rows.append(
                f"| {row['numerosity']} | {row['wider_width']}/{row['narrower_width']} "
                f"| {row['n_eligible']} "
                f"| {row['n_positive_wider_variance']} / {row['n_negative_wider_variance']} "
                f"/ {row['n_tied_variance_differences']} "
                f"| {row['exact_two_sided_sign_p']:.3g} | {adjusted:.3g} |"
            )
        text = text.replace("{{participant_sign_sensitivity_rows}}", "\n".join(rows))
    if "{{empirical_support_summary}}" in text:
        rows = raw_values["empirical_reanalysis"]["per_numerosity_cluster_bootstrap"]["per_numerosity"]
        support = {}
        for pair in ("20/40", "20/60", "40/60"):
            counts = [int(row["participants_with_residual_df_both_widths"][pair]) for row in rows]
            support[pair] = (min(counts), max(counts))
        text = text.replace(
            "{{empirical_support_summary}}",
            ", ".join(f"{lo}–{hi} for {pair}" for pair, (lo, hi) in support.items()),
        )
    if "{{lapse_false_positive_intervals}}" in text or "{{lapse_power_intervals}}" in text:
        unchanged, doubled = raw_values["lapse_robustness"]

        def rate_with_interval(result, key):
            rate = float(result["mc_uncertainty"][key]["estimate"])
            lo, hi = result["mc_uncertainty"][key]["wilson_95"]
            return f"{rate:.3f} (95% Wilson interval {lo:.3f}–{hi:.3f})"

        false_positive = (
            f"estimated null rejection rates of {rate_with_interval(unchanged, 'rejection_shared_N40')} under a shared-lapse fit at N=40 "
            f"and {rate_with_interval(unchanged, 'rejection_shared_N80')} at N=80; "
            f"the corresponding rates with state-specific lapse parameters are "
            f"{rate_with_interval(unchanged, 'rejection_state_N40')} at N=40 and "
            f"{rate_with_interval(unchanged, 'rejection_state_N80')} at N=80"
        )
        price_power = (
            f"{rate_with_interval(doubled, 'rejection_state_N40')} at N=40 and "
            f"{rate_with_interval(doubled, 'rejection_state_N80')} at N=80"
        )
        text = text.replace("{{lapse_false_positive_intervals}}", false_positive)
        text = text.replace("{{lapse_power_intervals}}", price_power)
    if "{{lapse_fit_diagnostics_main}}" in text or "{{lapse_fit_diagnostics_supplement}}" in text:
        unchanged, doubled = raw_values["lapse_robustness"]
        def fit_diag(result, model):
            diag = result["fit_diagnostics"][model]
            return diag["success_rate"], diag["boundary_rate"], diag["fits"]
        shared_1 = fit_diag(unchanged, "shared")
        state_1 = fit_diag(unchanged, "state")
        shared_2 = fit_diag(doubled, "shared")
        state_2 = fit_diag(doubled, "state")
        fits = shared_1[2]
        if not all(item[2] == fits for item in (state_1, shared_2, state_2)):
            raise ValueError("lapse fit diagnostic counts do not match across conditions")
        main_diagnostic = (
            f"Across the N=80 simulated cohorts ({fits:,} participant fits per model and price ratio; "
            "the N=40 analyses use nested participant subsets), optimizer convergence was "
            f"{min(item[0] for item in (shared_1, state_1, shared_2, state_2)):.1%}. "
            "At least one fitted parameter reached a bound in "
            f"{shared_1[1]:.1%} of shared-lapse fits and {state_1[1]:.1%} of state-specific fits "
            "with no price change, and in "
            f"{shared_2[1]:.1%} and {state_2[1]:.1%}, respectively, when price doubled. "
            "These aggregate rates do not identify which parameter reached a bound; the state-specific "
            "null rejection rate near 0.05 is therefore a design-specific projection, not general evidence of calibration."
        )
        supplement_diagnostic = (
            f"Each price-ratio/model cell contains {fits:,} participant fits from the N=80 simulated cohorts "
            "(N=40 analyses use nested subsets), and all fits report optimizer convergence. "
            "Boundary contact, defined as at least one fitted parameter reaching a bound, is "
            f"{shared_1[1]:.1%} for the shared-lapse and {state_1[1]:.1%} for the state-specific fit "
            "at price ratio 1, and "
            f"{shared_2[1]:.1%} and {state_2[1]:.1%}, respectively, at price ratio 2. "
            "These aggregate diagnostics do not identify which parameter reached a bound; frequent "
            "boundary contact qualifies interpretation of the state-specific fit's estimated null rejection rate."
        )
        text = text.replace("{{lapse_fit_diagnostics_main}}", main_diagnostic)
        text = text.replace("{{lapse_fit_diagnostics_supplement}}", supplement_diagnostic)
    if "{{baseline_effective_signal_mass}}" in text:
        gross = raw_values["net_increment"]["baseline"]["gross_E"]
        energy_per_signal = raw_values["config"]["budget"]["baseline"]["e"]
        text = text.replace("{{baseline_effective_signal_mass}}", f"{gross / energy_per_signal:.4f}")
    if "{{estimation_null_calibration}}" in text:
        null = next(row for row in raw_values["estimation_recovery"] if row["price_ratio"] == 1.0)

        def estimation_rate(key):
            rate = float(null[key])
            lo, hi = null["mc_uncertainty"][key]["wilson_95"]
            return f"{rate:.3f} (95% Wilson interval {lo:.3f}–{hi:.3f})"

        text = text.replace(
            "{{estimation_null_calibration}}",
            f"a no-change rejection rate of {estimation_rate('power')} for signed-rank and "
            f"{estimation_rate('power_ttest')} for the t test, each based on "
            f"{null['mc_uncertainty']['power']['replications']} replications",
        )
    if "{{empirical_per_number_rows}}" in text:
        analyses = raw_values["empirical_reanalysis"]["per_numerosity_cluster_bootstrap"]["contrasts"]
        supports = {
            int(row["numerosity"]): row
            for row in raw_values["empirical_reanalysis"]["per_numerosity_cluster_bootstrap"]["per_numerosity"]
        }
        by_number = {}
        for row in analyses:
            by_number.setdefault(int(row["numerosity"]), {})[(int(row["width_1"]), int(row["width_2"]))] = row
        def p_text(value):
            return format(float(value), ".3g")
        rows = []
        for x in sorted(by_number):
            r2040 = by_number[x][(20, 40)]
            r2060 = by_number[x][(20, 60)]
            r4060 = by_number[x][(40, 60)]
            support = supports[x]
            n_by_width = support["participants_with_residual_df_by_width"]
            n_both = support["participants_with_residual_df_both_widths"]
            rows.append(
                f"| {x}; n={n_by_width['20']}/{n_by_width['40']}/{n_by_width['60']}; "
                f"paired n={n_both['20/40']}/{n_both['20/60']}/{n_both['40/60']} "
                f"| {r2040['variance_ratio_width_2_over_width_1']:.2f} ({p_text(r2040['pvalue_bonferroni'])}) "
                f"| {r2060['variance_ratio_width_2_over_width_1']:.2f} ({p_text(r2060['pvalue_bonferroni'])}) "
                f"| {r4060['variance_ratio_width_2_over_width_1']:.2f} ({p_text(r4060['pvalue_bonferroni'])}) |"
            )
        text = text.replace("{{empirical_per_number_rows}}", "\n".join(rows))
    if "{{empirical_equal_weight_rows}}" in text:
        analyses = raw_values["empirical_reanalysis"]["participant_equal_weight_sensitivity"]["contrasts"]
        rows = []
        for row in analyses:
            lo, hi = row["familywise_ci_variance_ratio"]
            rows.append(
                f"| {row['width_2']} / {row['width_1']} | {row['n_participants']} "
                f"| {row['geometric_mean_within_participant_variance_ratio']:.2f} "
                f"| [{lo:.2f}, {hi:.2f}] | {row['pvalue_bonferroni']:.3g} |"
            )
        text = text.replace("{{empirical_equal_weight_rows}}", "\n".join(rows))
    if "{{empirical_order_rows}}" in text:
        analyses = raw_values["empirical_reanalysis"]["order_sequence_sensitivity"]["contrasts"]
        rows = []
        for row in analyses:
            w, n = row["widening"], row["narrowing"]
            wlo, whi = w["familywise_ci_variance_ratio"]
            nlo, nhi = n["familywise_ci_variance_ratio"]
            diff = row["difference_log_ratio_widening_minus_narrowing"]
            dlo = diff["estimate"] - diff["familywise_ci_log_critical"]
            dhi = diff["estimate"] + diff["familywise_ci_log_critical"]
            rows.append(
                f"| {row['width_2']} / {row['width_1']} "
                f"| {w['geometric_mean_within_participant_variance_ratio']:.2f} "
                f"[{wlo:.2f}, {whi:.2f}] "
                f"| {n['geometric_mean_within_participant_variance_ratio']:.2f} "
                f"[{nlo:.2f}, {nhi:.2f}] "
                f"| {diff['estimate']:.2f} [{dlo:.2f}, {dhi:.2f}] |"
            )
        text = text.replace("{{empirical_order_rows}}", "\n".join(rows))
    if "{{empirical_position_rows}}" in text:
        analyses = raw_values["empirical_reanalysis"]["within_block_position_sensitivity"]["contrasts"]
        rows = []
        for row in analyses:
            lo, hi = row["familywise_ci_variance_ratio"]
            mc_lo, mc_hi = row["pvalue_bootstrap_mc_95"]
            rows.append(
                f"| {row['block_half'].replace('_', ' ')} | {row['width_2']} / {row['width_1']} "
                f"| {row['variance_ratio_width_2_over_width_1']:.2f} [{lo:.2f}, {hi:.2f}] "
                f"| {row['pvalue_bonferroni']:.3g} (raw {mc_lo:.3g}–{mc_hi:.3g}) | "
                f"{row['participants_with_df_width_1']}/{row['participants_with_df_width_2']} |"
            )
        text = text.replace("{{empirical_position_rows}}", "\n".join(rows))
    if "{{simulation_summary_rows}}" in text:
        rows = []
        def add_sim_row(label, ratio, n, estimate, uncertainty, reps, test):
            lo, hi = uncertainty["wilson_95"]
            rows.append(
                f"| {label} | {ratio:g} | {n} | {test} | {estimate:.3f} "
                f"| [{lo:.3f}, {hi:.3f}] | {reps} |"
            )
        for result in raw_values["power"]["by_price_ratio"]:
            ratio = result["price_ratio"]
            for n in raw_values["power"]["N_grid"]:
                for suffix, test in (("", "one-sided t"), ("_wilcoxon", "one-sided Wilcoxon")):
                    key = f"power{suffix}_N{n}"
                    add_sim_row("Discrimination", ratio, n, result[key],
                                result["mc_uncertainty"][key],
                                result["mc_uncertainty"][key]["replications"], test)
        for result in raw_values["estimation_recovery"]:
            for key, test in (("power", "one-sided signed-rank"), ("power_ttest", "one-sided t")):
                add_sim_row("Estimation", result["price_ratio"], result["N"], result[key],
                            result["mc_uncertainty"][key],
                            result["mc_uncertainty"][key]["replications"], test)
        for result in raw_values["lapse_robustness"]:
            for model in ("shared", "state"):
                for n in raw_values["config"]["lapse_robustness"]["N_grid"]:
                    key = f"rejection_{model}_N{n}"
                    add_sim_row("Discrimination, lapse sensitivity", result["price_ratio"], n,
                                result[key], result["mc_uncertainty"][key],
                                result["mc_uncertainty"][key]["replications"],
                                f"one-sided t, {model} lapse")
        text = text.replace("{{simulation_summary_rows}}", "\n".join(rows))
    missing, used = [], set()

    def repl(m):
        key, spec = m.group(1).strip(), (m.group(2) or "").strip()
        if key not in values:
            missing.append(key)
            return m.group(0)
        used.add(key)
        try:
            return fmt(values[key], spec)
        except (ValueError, TypeError) as e:
            missing.append(f"{key} (format '{spec}': {e})")
            return m.group(0)

    filled = re.sub(r"\{\{\s*([A-Za-z_][\w.]*)\s*(?::([^}]*))?\}\}", repl, text)
    open(output_path, "w", encoding="utf-8").write(filled)
    print(f"Wrote {output_path} | {len(used)} keys used of {len(values)} in {results_path}")
    unused = sorted(set(values) - used)
    if unused:
        print("Results not quoted in the text: " + ", ".join(unused[:30]) + (" ..." if len(unused) > 30 else ""))
    if missing:
        print("MISSING, left in place: " + ", ".join(missing))
        return 1
    return 0


if __name__ == "__main__":
    sys.exit(main(sys.argv[1:]))
```
