## Supplementary material for "The marginal brain": Replication package: LICENSE

MIT License

Copyright (c) 2026 Pier Luigi Sacco

Permission is hereby granted, free of charge, to any person obtaining a copy of this software and associated
documentation files (the "Software"), to deal in the Software without restriction, including without limitation the
rights to use, copy, modify, merge, publish, distribute, sublicense, and/or sell copies of the Software, and to permit
persons to whom the Software is furnished to do so, subject to the following conditions:

The above copyright notice and this permission notice shall be included in all copies or substantial portions of the
Software.

THE SOFTWARE IS PROVIDED "AS IS", WITHOUT WARRANTY OF ANY KIND, EXPRESS OR IMPLIED, INCLUDING BUT NOT LIMITED TO THE
WARRANTIES OF MERCHANTABILITY, FITNESS FOR A PARTICULAR PURPOSE AND NONINFRINGEMENT. IN NO EVENT SHALL THE AUTHORS OR
COPYRIGHT HOLDERS BE LIABLE FOR ANY CLAIM, DAMAGES OR OTHER LIABILITY, WHETHER IN AN ACTION OF CONTRACT, TORT OR
OTHERWISE, ARISING FROM, OUT OF OR IN CONNECTION WITH THE SOFTWARE OR THE USE OR OTHER DEALINGS IN THE SOFTWARE.
