## Supplementary material for "The marginal brain": Replication package: participant_sign_sensitivity.py

```
"""Exact participant-level sign checks for matched response-variance contrasts.

These deterministic, post hoc analyses target participant-level directions,
not the pooled residual-variance ratio used in the primary analysis. The
numerosity-specific family tests paired sample-variance signs at each level;
the aggregate family first pools each participant's residual sums of squares
and degrees of freedom across the common support and tests the resulting
variance-ratio directions. Both use exact binomial sign tests with separate
Bonferroni families.
"""
from __future__ import annotations

import csv
import hashlib
import json
import math
import statistics
from collections import defaultdict
from itertools import combinations
from pathlib import Path

from scipy.stats import binomtest

ROOT = Path(__file__).resolve().parent
DEFAULT_DATA = ROOT.parent / "data" / "published_estimation.csv"
DEFAULT_OUTPUT = ROOT / "participant_sign_sensitivity.json"


def _sample_variance(values: list[float]) -> float:
    mean = math.fsum(values) / len(values)
    return math.fsum((value - mean) ** 2 for value in values) / (len(values) - 1)


def _data_sha256(path: Path) -> str:
    digest = hashlib.sha256()
    with path.open("rb") as stream:
        for chunk in iter(lambda: stream.read(1024 * 1024), b""):
            digest.update(chunk)
    return digest.hexdigest()


def analyze(data_path: Path = DEFAULT_DATA) -> dict:
    widths = (20, 40, 60)
    numerosities = tuple(range(50, 71))
    pairs = tuple(combinations(widths, 2))
    family_size = len(numerosities) * len(pairs)
    cells: dict[tuple[str, int, int], list[float]] = defaultdict(list)
    orders: dict[str, str] = {}

    with data_path.open("r", encoding="utf-8-sig", newline="") as stream:
        for row in csv.DictReader(stream):
            participant = str(row["ID"])
            width = int(float(row["Delta"]))
            number = int(float(row["Number"]))
            order = str(row["Order"]).strip()
            if participant in orders and orders[participant] != order:
                raise ValueError(f"Order changes within participant {participant}")
            orders[participant] = order
            if width in widths and number in numerosities:
                cells[(participant, width, number)].append(float(row["Response"]))

    if set(orders.values()) != {"Widening", "Narrowing"}:
        raise ValueError("Both recorded block-order sequences must be represented")

    # Aggregate each participant's within-cell residual information over the
    # common numerosity range before computing one variance ratio per contrast.
    # This matches the participant-equal weighting target used in the separate
    # geometric-mean sensitivity while supporting a distribution-free test of
    # its median direction.
    participant_variance: dict[tuple[str, int], float] = {}
    for participant in sorted(orders):
        for width in widths:
            total_sse = total_df = 0.0
            for number in numerosities:
                values = cells.get((participant, width, number), [])
                if len(values) >= 2:
                    mean = math.fsum(values) / len(values)
                    total_sse += math.fsum((value - mean) ** 2 for value in values)
                    total_df += len(values) - 1
            if total_df > 0 and total_sse > 0:
                participant_variance[(participant, width)] = total_sse / total_df

    aggregate_contrasts = []
    for narrower, wider in pairs:
        positive = negative = ties = missing = 0
        ratios: list[float] = []
        for participant in sorted(orders):
            narrow_variance = participant_variance.get((participant, narrower))
            wide_variance = participant_variance.get((participant, wider))
            if narrow_variance is None or wide_variance is None:
                missing += 1
                continue
            ratio = wide_variance / narrow_variance
            ratios.append(ratio)
            log_ratio = math.log(ratio)
            if math.isclose(log_ratio, 0.0, rel_tol=0.0, abs_tol=1e-12):
                ties += 1
            elif log_ratio > 0:
                positive += 1
            else:
                negative += 1

        non_tied = positive + negative
        raw_p = (
            float(binomtest(positive, non_tied, p=0.5, alternative="two-sided").pvalue)
            if non_tied else 1.0
        )
        aggregate_contrasts.append({
            "contrast": f"{wider}/{narrower}",
            "n_eligible": positive + negative + ties,
            "n_missing_incomplete_width_support": missing,
            "n_positive_wider_variance": positive,
            "n_negative_wider_variance": negative,
            "n_tied_variance_ratios": ties,
            "median_participant_variance_ratio": (
                float(statistics.median(ratios)) if ratios else None
            ),
            "geometric_mean_participant_variance_ratio": (
                float(math.exp(math.fsum(math.log(value) for value in ratios) / len(ratios)))
                if ratios else None
            ),
            "exact_two_sided_sign_p": raw_p,
            "bonferroni_adjusted_p_3": min(1.0, raw_p * len(pairs)),
            "significant_familywise_05": bool(raw_p * len(pairs) < 0.05),
        })

    rows = []
    for number in numerosities:
        for narrower, wider in pairs:
            signs_by_order = {
                order: {"positive": 0, "negative": 0, "ties": 0, "eligible": 0}
                for order in ("Widening", "Narrowing")
            }
            positive = negative = ties = missing = one_zero = both_zero = 0
            differences: list[float] = []

            for participant in sorted(orders):
                values_narrow = cells.get((participant, narrower, number), [])
                values_wide = cells.get((participant, wider, number), [])
                if len(values_narrow) < 2 or len(values_wide) < 2:
                    missing += 1
                    continue

                variance_narrow = _sample_variance(values_narrow)
                variance_wide = _sample_variance(values_wide)
                if variance_narrow == 0.0 and variance_wide == 0.0:
                    both_zero += 1
                elif (variance_narrow == 0.0) != (variance_wide == 0.0):
                    one_zero += 1

                difference = variance_wide - variance_narrow
                differences.append(difference)
                if math.isclose(difference, 0.0, rel_tol=0.0, abs_tol=1e-12):
                    sign = "ties"
                    ties += 1
                elif difference > 0:
                    sign = "positive"
                    positive += 1
                else:
                    sign = "negative"
                    negative += 1

                sequence = signs_by_order[orders[participant]]
                sequence[sign] += 1
                sequence["eligible"] += 1

            non_tied = positive + negative
            raw_p = (
                float(binomtest(positive, non_tied, p=0.5, alternative="two-sided").pvalue)
                if non_tied else 1.0
            )
            adjusted_p = min(1.0, raw_p * family_size)
            rows.append({
                "numerosity": number,
                "narrower_width": narrower,
                "wider_width": wider,
                "n_eligible": positive + negative + ties,
                "n_missing_insufficient_cell_trials": missing,
                "n_positive_wider_variance": positive,
                "n_negative_wider_variance": negative,
                "n_tied_variance_differences": ties,
                "n_one_zero_variance_pairs": one_zero,
                "n_both_zero_variance_pairs": both_zero,
                "median_participant_variance_difference": (
                    float(statistics.median(differences)) if differences else None
                ),
                "exact_two_sided_sign_p": raw_p,
                "bonferroni_adjusted_p_63": adjusted_p,
                "significant_familywise_05": bool(adjusted_p < 0.05),
                "signs_by_recorded_sequence": signs_by_order,
            })

    summaries = []
    for narrower, wider in pairs:
        contrast_rows = [
            row for row in rows
            if row["narrower_width"] == narrower and row["wider_width"] == wider
        ]
        supports = [row["n_eligible"] for row in contrast_rows]
        medians = [row["median_participant_variance_difference"] for row in contrast_rows
                   if row["median_participant_variance_difference"] is not None]
        summaries.append({
            "contrast": f"{wider}/{narrower}",
            "corrected_numerosities": sum(row["significant_familywise_05"] for row in contrast_rows),
            "family_size": len(contrast_rows),
            "corrected_numerosity_values": [
                row["numerosity"] for row in contrast_rows if row["significant_familywise_05"]
            ],
            "eligible_participant_range": [min(supports), max(supports)],
            "median_eligible_participants": float(statistics.median(supports)),
            "median_of_numerosity_specific_median_variance_differences": float(statistics.median(medians)),
        })

    return {
        "analysis": "participant-level exact sign sensitivity for matched sample-variance differences",
        "status": "post hoc sensitivity; not preregistered; same published cohort",
        "estimand": "median direction of participant-specific sample-variance differences, conditional on usable paired cells",
        "null_hypothesis": "among eligible participants, positive and negative paired variance differences are equally probable",
        "test": "two-sided exact binomial sign test; ties omitted from the binomial denominator; one-zero pairs retained as directional signs",
        "participant_independence_unit": "participant",
        "minimum_trials_per_cell": 2,
        "widths": list(widths),
        "numerosities": list(numerosities),
        "family_size": family_size,
        "familywise_alpha": 0.05,
        "multiplicity": "Bonferroni within these 63 contrasts only; no correction across distinct sensitivity families",
        "data_sha256": _data_sha256(data_path),
        "n_participants": len(orders),
        "sequence_counts": {
            order: sum(value == order for value in orders.values())
            for order in ("Widening", "Narrowing")
        },
        "summary": summaries,
        "contrasts": rows,
        "aggregate_participant_direction": {
            "estimand": "median direction of participant-specific pooled residual-variance ratios, pooling available within-cell residual sums of squares and degrees of freedom over numerosities 50–70",
            "null_hypothesis": "among eligible participants, positive and negative log variance ratios are equally probable",
            "test": "two-sided exact binomial sign test; ties omitted from the binomial denominator",
            "support": "participants with positive residual degrees of freedom at each width after pooling their available cells over the common numerosity range",
            "family_size": len(pairs),
            "familywise_alpha": 0.05,
            "multiplicity": "Bonferroni across the three width contrasts only; separate from the 63 numerosity-specific tests and not corrected across all sensitivity families",
            "contrasts": aggregate_contrasts,
        },
    }


def main() -> None:
    result = analyze(DEFAULT_DATA)
    DEFAULT_OUTPUT.write_text(json.dumps(result, indent=2) + "\n", encoding="utf-8")
    print(f"Wrote {len(result['contrasts'])} exact sign tests to {DEFAULT_OUTPUT}")
    for row in result["summary"]:
        print(
            f"{row['contrast']}: {row['corrected_numerosities']}/{row['family_size']} corrected; "
            f"eligible participants {row['eligible_participant_range']}"
        )
    for row in result["aggregate_participant_direction"]["contrasts"]:
        print(
            f"aggregate {row['contrast']}: {row['n_positive_wider_variance']}/"
            f"{row['n_eligible']} positive; exact sign p adj = "
            f"{row['bonferroni_adjusted_p_3']:.3g}"
        )


if __name__ == "__main__":
    main()
```
