## Supplementary material for "The marginal brain": Replication package: precompute_estimation.py

```
"""Optional: compute estimation-recovery chunks ahead of run_all.py (each chunk is cached and seeded independently).
Usage: python3 precompute_estimation.py <price_ratio> <first_chunk> <last_chunk>"""
import json, os, sys
sys.path.insert(0, os.path.dirname(os.path.abspath(__file__)))
from spt import behavior, recovery
cfg = json.load(open('config.json'))
cal = behavior.calibrate_estimation(cfg['calibration']['estimation'])
ecfg = cfg['estimation_recovery']
pr, c0, c1 = float(sys.argv[1]), int(sys.argv[2]), int(sys.argv[3])
per_chunk = int(ecfg.get('chunk_size', 10))
n_chunks = (ecfg['reps'] + per_chunk - 1) // per_chunk
os.makedirs('cache', exist_ok=True)
fingerprint = recovery.estimation_cache_fingerprint(cal, ecfg, pr, cfg['seed'], per_chunk)
for ch in range(c0, c1 + 1):
    reps_in_chunk = min(per_chunk, ecfg['reps'] - ch * per_chunk)
    if reps_in_chunk <= 0:
        continue
    path = f'cache/estimation_{fingerprint}_ratio{pr}_chunk{ch}_of{n_chunks}_seed{cfg["seed"]}.json'
    cached = None
    if os.path.exists(path):
        try:
            candidate = json.load(open(path, encoding='utf-8'))
            if (candidate.get('cache_fingerprint') == fingerprint
                    and len(candidate.get('reps', [])) == reps_in_chunk
                    and candidate.get('fit_count') == reps_in_chunk * ecfg['N']):
                cached = candidate
        except (OSError, ValueError, TypeError):
            cached = None
    if cached is None:
        out = recovery.estimation_chunk(cal, ecfg, pr, ch, reps_in_chunk, cfg['seed'])
        out['cache_fingerprint'] = fingerprint
        tmp_path = path + '.tmp'
        with open(tmp_path, 'w', encoding='utf-8') as f:
            json.dump(out, f)
        os.replace(tmp_path, path)
        print('wrote', path, flush=True)
```
