## Supplementary material for "The marginal brain": Replication package: predicted_variance_ratios.py

```
"""Variance ratios that the calibrated estimation model predicts at each shared numerosity (50 to 70), for comparison
with the matched-numerosity reanalysis. The calibration anchor is numerosity 60; the other shared levels are not used for fitting, but are from the same cohort and are not an independent validation sample.
Deterministic (Gauss-Hermite quadrature); writes model_predicted_ratios.json."""
import json, os
import numpy as np
from numpy.polynomial.hermite_e import hermegauss
from scipy.stats import norm, truncnorm

HERE = os.path.dirname(os.path.abspath(__file__))
res = json.load(open(os.path.join(HERE, 'results.json'), encoding='utf-8'))
cal = res['calibration']['estimation']
nu, s0 = cal['nu'], cal['sigma0']
RANGES = {20: (50, 70), 40: (40, 80), 60: (30, 90)}
gx, gw = hermegauss(80)
gw = gw / gw.sum()


def response_variance(x, w):
    lo, hi = RANGES[w]
    s = nu * np.sqrt(w)
    r = x + s * gx
    a, b = (lo - r) / s, (hi - r) / s
    m = truncnorm.mean(a, b, loc=r, scale=s)   # posterior mean under the uniform prior, numerically stable
    k = np.arange(lo, hi + 1)
    P = norm.cdf((k[None, :] + 0.5 - m[:, None]) / s0) - norm.cdf((k[None, :] - 0.5 - m[:, None]) / s0)
    P /= P.sum(1, keepdims=True)
    mean_k = (P * k).sum(1)
    var_k = (P * k**2).sum(1) - mean_k**2
    return float((gw * var_k).sum() + (gw * mean_k**2).sum() - (gw * mean_k).sum()**2)


xs = list(range(50, 71))
V = {w: [response_variance(x, w) for x in xs] for w in RANGES}
out = {'analysis': 'model-predicted response-variance ratios at matched numerosities', 'numerosities': xs,
       'parameters': {'nu': nu, 'sigma0': s0}, 'per_numerosity': {}}
for (num, den) in ((40, 20), (60, 20), (60, 40)):
    ratio = [V[num][i] / V[den][i] for i in range(len(xs))]
    key = f'{den}_{num}'
    out['per_numerosity'][key] = ratio
    out[f'pred_ratio_{key}_median'] = float(np.median(ratio))
    out[f'pred_ratio_{key}_min'] = float(min(ratio))
    out[f'pred_ratio_{key}_max'] = float(max(ratio))
contrasts = res['empirical_reanalysis']['per_numerosity_cluster_bootstrap']['contrasts']
for (den, num) in ((20, 40), (20, 60), (40, 60)):
    key = f'{den}_{num}'
    pred = dict(zip(xs, out['per_numerosity'][key]))
    rows = [r for r in contrasts if r['width_1'] == den and r['width_2'] == num]
    inside = sum(1 for r in rows if r['familywise_ci_variance_ratio'][0] <= pred[r['numerosity']] <= r['familywise_ci_variance_ratio'][1])
    out[f'pred_within_ci_{key}'] = inside
    out[f'pred_n_levels_{key}'] = len(rows)
json.dump(out, open(os.path.join(HERE, 'model_predicted_ratios.json'), 'w', encoding='utf-8'), indent=1)
print({k: round(v, 3) for k, v in out.items() if k.startswith('pred_ratio')})
```
