## Supplementary material for "The marginal brain": Replication package: README.md

# Model and simulation package

This directory contains the analytic and numerical implementation supporting the manuscript. `config.json` defines the primary model and simulation settings; `run_all.py` regenerates the primary results and model figures, including the matched-numerosity participant-clustered reanalysis of the public estimation trials in `../data/`. `complete_case_sensitivity.py` separately regenerates the pair-specific matched-support sensitivity, and `participant_sign_sensitivity.py` regenerates the exact participant-level sign-balance sensitivity. `tests/test_model.py` contains 18 algebraic, numerical, data-integrity and source-to-output checks. When the source trial CSV is absent, its direct runner reports the two data-dependent checks as skipped and runs the remaining checks. `conditional_null_calibration.py` independently reruns the conditional Gaussian-null pilot stored under `outputs/conditional_null_calibration/`; it is a diagnostic for the primary centered bootstrap and is not included in the routine run. The repository-level `reproduce.ps1` and `reproduce.sh` scripts orchestrate all these steps with figure generation, manuscript builds and tests.

Run from this directory:

```powershell
python run_all.py
python tests/test_model.py
```

The trial-level source data are not bundled; obtain them from the OSF project and place published_estimation.csv in ../data/ as described in ../data/README.md.


The results file records the campaign-time source fingerprint. The package manifest covers the current distributed files, including the TeX sources, PDFs and figure assets; its generator is write_package_manifest.py.

In the primary JSON output, fields named `significant_familywise` and `significant_numerosities` indicate whether a bootstrap result crosses the configured nominal Bonferroni threshold. The companion `familywise_ci_*` fields are the corresponding nominal interval outputs. These field names describe the procedure's decision rule; the manuscript reports them as nominal and does not interpret them as evidence that familywise error is empirically controlled.

The pair-specific sensitivity uses 500,000 participant-cluster bootstrap draws per contrast and numerosity with seed 20261013. It writes `complete_case_sensitivity.json` separately and does not alter `results.json`.

The conditional Gaussian-null diagnostic can be rerun separately, using the source CSV obtained from the cited OSF project:

```powershell
python conditional_null_calibration.py --data ..\data\published_estimation.csv --outer 100 --inner 10000 --seed 20261019
```

The recorded run's summary, cohort-level test results, and toy validation are included under `outputs/conditional_null_calibration/`. This diagnostic is separate from the primary analysis and does not establish general bootstrap validity.
