## Supplementary material for "The marginal brain": Replication package: refresh_derived.py

```
"""Compatibility entry point for regenerating the complete analysis package.

The canonical runner reuses only fingerprint-verified empirical-bootstrap and
estimation-recovery caches; all configured simulation campaigns are evaluated.
"""
from run_all import main


if __name__ == "__main__":
    main()
```
