## Supplementary material for "The marginal brain": Replication package: run_all.py

```
"""Regenerates every result and figure of the model from config.json. Usage: python3 run_all.py"""
import json, time, sys, os, platform, hashlib
import numpy as np
import scipy
sys.path.insert(0, os.path.dirname(os.path.abspath(__file__)))
from spt import analytics, behavior, recovery, validation, figures, numerics, empirical


def empirical_cache_fingerprint(data_path, settings):
    """Key the slower trial-level bootstrap cache to source, data, settings, and runtime."""
    digest = hashlib.sha256()
    payload = {'schema': 1, 'settings': settings,
               'python': platform.python_version(), 'numpy': np.__version__,
               'scipy': scipy.__version__}
    digest.update(json.dumps(payload, sort_keys=True, separators=(',', ':')).encode('utf-8'))
    for path in (empirical.__file__, data_path):
        with open(path, 'rb') as stream:
            digest.update(hashlib.sha256(stream.read()).digest())
    return digest.hexdigest()


def main():
    t0 = time.time()
    cfg = json.load(open('config.json'))
    package_root = os.path.dirname(os.path.dirname(os.path.abspath(__file__)))
    source_paths = []
    for subtree in ('model', 'paper'):
        for current, dirs, files in os.walk(os.path.join(package_root, subtree)):
            dirs[:] = [d for d in dirs if d not in {'__pycache__', 'node_modules', 'qa_render',
                                                     'qa_render_delivery', 'qa_render_final', 'figures'}]
            for name in files:
                if name.endswith(('.py', '.js')):
                    source_paths.append(os.path.join(current, name))
    source_paths.extend(os.path.join(package_root, p) for p in (
        'README.md', 'reproduce.ps1', 'reproduce.sh', 'LICENSE',
        'model/README.md', 'model/config.json', 'model/requirements.txt',
        'paper/body.md', 'paper/supplement.md',
        'paper/refs.json', 'paper/build/package.json', 'paper/build/package-lock.json',
        'data/published_estimation.csv'))
    source_manifest = {}
    for path in sorted(set(source_paths)):
        if not os.path.isfile(path):
            continue  # optional build files (e.g. the Word builder's npm manifests) may be absent
        rel = os.path.relpath(path, package_root).replace(os.sep, '/')
        with open(path, 'rb') as stream:
            source_manifest[rel] = hashlib.sha256(stream.read()).hexdigest()
    source_digest = hashlib.sha256(json.dumps(source_manifest, sort_keys=True).encode('utf-8')).hexdigest()
    rng = np.random.default_rng(cfg['seed'])
    res = {'meta': {'seed': cfg['seed'], 'python': platform.python_version(),
                    'numpy': np.__version__, 'scipy': scipy.__version__,
                    'source_sha256': source_digest, 'source_manifest': source_manifest,
                    'source_manifest_scope': 'Campaign-time source fingerprint; subsequent package and manuscript edits are inventoried separately in model/package_manifest.json.',
                    'package_manifest': 'model/package_manifest.json'},
           'config': cfg, 'spec': {'bimodal_prior': numerics.BIMODAL, 'values_averaged_per_color': 5}}
    bm = numerics.BIMODAL
    ce, cd = cfg['calibration']['estimation'], cfg['calibration']['discrimination']
    words = lambda xs: ', '.join(xs[:-1]) + ' and ' + xs[-1] if len(xs) > 1 else xs[0]
    res['spec'].update({
        'bimodal_mean_1': bm['means'][0], 'bimodal_mean_2': bm['means'][1],
        'bimodal_sd_1': bm['sds'][0], 'bimodal_sd_2': bm['sds'][1],
        'bimodal_weight_1': bm['weights'][0], 'bimodal_weight_2': bm['weights'][1],
        'est_calibration_ranges': words([f'{lo} to {hi}' for lo, hi in zip(ce['lows'], ce['highs'])]),
        'disc_calibration_widths': words(sorted({f'{int(p[0])}' for p in cd['points']}, key=float)),
        'chunk_size': cfg['estimation_recovery'].get('chunk_size', 10)})
    print('Stage 2: symbolic derivations')
    res['analytics'] = analytics.derive()
    print('Calibration to reported values')
    res['calibration'] = {'estimation': behavior.calibrate_estimation(cfg['calibration']['estimation']),
                          'discrimination': behavior.calibrate_discrimination(cfg['calibration']['discrimination']),
                          'source': cfg['calibration']['source']}
    ercfg = cfg['empirical_reanalysis']
    data_path = os.path.normpath(os.path.join(os.path.dirname(__file__), ercfg['data_path']))
    empirical_key = empirical_cache_fingerprint(data_path, ercfg)
    empirical_cache_dir = os.path.join(os.path.dirname(__file__), 'cache')
    os.makedirs(empirical_cache_dir, exist_ok=True)
    empirical_cache_path = os.path.join(empirical_cache_dir, f'empirical_{empirical_key}.json')
    cached_empirical = None
    if os.path.isfile(empirical_cache_path):
        try:
            with open(empirical_cache_path, encoding='utf-8') as stream:
                cache_payload = json.load(stream)
            if cache_payload.get('fingerprint') == empirical_key:
                cached_empirical = cache_payload.get('result')
        except (OSError, json.JSONDecodeError):
            cached_empirical = None
    if cached_empirical is None:
        print('Stage 3: participant-clustered empirical analysis (cache miss)')
        res['empirical_reanalysis'] = empirical.reanalyze_estimation(
            data_path, widths=ercfg['widths'], anchor_number=ercfg['anchor_number'],
            shared_numbers=range(ercfg['shared_number_min'], ercfg['shared_number_max'] + 1),
            published_sd=ercfg['published_sd'], familywise_alpha=ercfg['familywise_alpha'],
            bootstrap_reps=ercfg['bootstrap_reps'], bootstrap_seed=ercfg['bootstrap_seed'],
            equal_weight_bootstrap_reps=ercfg['participant_equal_weight_bootstrap_reps'],
            position_bootstrap_reps=ercfg['position_sensitivity_bootstrap_reps'])
        temporary_cache_path = empirical_cache_path + '.tmp'
        with open(temporary_cache_path, 'w', encoding='utf-8') as stream:
            json.dump({'fingerprint': empirical_key, 'result': res['empirical_reanalysis']},
                      stream, indent=1, default=float)
        os.replace(temporary_cache_path, empirical_cache_path)
    else:
        print('Stage 3: participant-clustered empirical analysis (verified cache hit)')
        res['empirical_reanalysis'] = cached_empirical
    print('Stage 4: budget, robustness, signatures, ablations, Proposition 5')
    res['budget'] = validation.budget_analysis(cfg['budget'])
    res['P5_construction'] = validation.constructive_prop5()
    res['robustness'] = validation.robustness_numeric(cfg['numerics'])
    res['signatures'] = validation.signatures()
    res['ablations'] = validation.ablations(cfg['budget'])
    res['net_increment'] = validation.net_increment_analysis(cfg['budget'])
    res['integer_sweep'] = validation.integer_sweep()
    res['displacement_remark'] = validation.displacement_remark()
    res['calibration']['estimation_integer_prior_refit'] = validation.calibration_integer_prior_refit(cfg['calibration']['estimation'])
    res['calibration']['estimation_discrete_prior_sd'] = validation.calibration_discrete_prior(
        res['calibration']['estimation'], cfg['calibration']['estimation'])
    print('Stage 4: power (discrimination) and recovery (estimation)')
    pw = []
    for pr in cfg['price_ratios']:
        pw.append(recovery.discrimination_power(rng, res['calibration']['discrimination'], cfg['power'], pr))
        print('   price ratio', pr, {k: v for k, v in pw[-1].items() if k.startswith('power')})
    res['power'] = {'N_grid': cfg['power']['N_grid'], 'by_price_ratio': pw, 'design': cfg['power']}
    er = []
    ecfg = cfg['estimation_recovery']
    per_chunk = int(ecfg.get('chunk_size', 10))
    n_chunks = (ecfg['reps'] + per_chunk - 1) // per_chunk
    os.makedirs('cache', exist_ok=True)
    for pr in ecfg['price_ratios']:
        chunks = []
        fingerprint = recovery.estimation_cache_fingerprint(
            res['calibration']['estimation'], ecfg, pr, cfg['seed'], per_chunk)
        for ch in range(n_chunks):
            reps_in_chunk = min(per_chunk, ecfg['reps'] - ch * per_chunk)
            path = f'cache/estimation_{fingerprint}_ratio{pr}_chunk{ch}_of{n_chunks}_seed{cfg["seed"]}.json'
            cached = None
            if os.path.exists(path):
                try:
                    candidate = json.load(open(path, encoding='utf-8'))
                    if (candidate.get('cache_fingerprint') == fingerprint
                            and len(candidate.get('reps', [])) == reps_in_chunk
                            and candidate.get('fit_count') == reps_in_chunk * ecfg['N']):
                        cached = candidate
                except (OSError, ValueError, TypeError):
                    cached = None
            if cached is None:
                out = recovery.estimation_chunk(res['calibration']['estimation'], ecfg, pr,
                                                ch, reps_in_chunk, cfg['seed'])
                out['cache_fingerprint'] = fingerprint
                tmp_path = path + '.tmp'
                with open(tmp_path, 'w', encoding='utf-8') as f:
                    json.dump(out, f)
                os.replace(tmp_path, path)
                chunks.append(out)
            else:
                chunks.append(cached)
        er.append(recovery.estimation_aggregate(chunks, pr, ecfg))
        print('   estimation, price ratio', pr, er[-1])
    res['estimation_recovery'] = er
    # headline numbers for the text (flat keys)
    b = res['budget']
    res['headline'] = {
        'el_sigma_price': 0.25, 'el_energy_price': -0.5,
        'common_factor_price_x2': 2**0.25,
        'slope_price': -0.5, 'slope_technology': 0.5,
        'calib_est_nu': res['calibration']['estimation']['nu'],
        'calib_est_sigma0': res['calibration']['estimation']['sigma0'],
        'calib_disc_nu': res['calibration']['discrimination']['nu'],
        'calib_disc_eta': res['calibration']['discrimination']['eta'],
        'baseline_task_share': b['baseline_task_share_of_margin_use'],
        'el_mu_taskscale': b['el_mu_taskscale_numeric'],
        'el_sigma_M_baseline': b['el_sigma_M_baseline'],
        'tighten30_mu_ratio': b['tighten30_mu_ratio'],
        'tighten30_sigma_ratio': b['tighten30_sigma_ratio'],
        'tighten30_energy_ratio': b['tighten30_task_energy_ratio'],
        'P5_max_err': max(res['P5_construction']['max_rel_err_mu'], res['P5_construction']['max_rel_err_share']),
        'numeric_worst_closed_form_err': res['robustness']['worst_closed_form_err'],
        'numeric_worst_common_factor_err': res['robustness']['worst_common_factor_err'],
        'empirical_n_rows': res['empirical_reanalysis']['n_rows'],
        'empirical_n_subjects': res['empirical_reanalysis']['n_subjects'],
        'empirical_paired_n': res['empirical_reanalysis']['per_numerosity_cluster_bootstrap']['n_clusters'],
        'empirical_shared_number_count': len(res['empirical_reanalysis']['per_numerosity_cluster_bootstrap']['shared_numerosities']),
        'empirical_family_size': res['empirical_reanalysis']['per_numerosity_cluster_bootstrap']['planned_family_size'],
        'empirical_familywise_interval_level_percent': res['empirical_reanalysis']['per_numerosity_cluster_bootstrap']['familywise_interval_level_percent'],
    }
    for anchor in res['empirical_reanalysis']['anchors']:
        res['headline'][f"empirical_sd_width_{anchor['prior_width']}"] = anchor['observed_sd']
        res['headline'][f"empirical_sd_difference_width_{anchor['prior_width']}"] = anchor['difference']
        res['headline'][f"empirical_n_width_{anchor['prior_width']}"] = anchor['n_trials']
    res['headline']['empirical_max_abs_sd_difference'] = max(
        abs(anchor['difference']) for anchor in res['empirical_reanalysis']['anchors'])
    for row in res['empirical_reanalysis']['per_numerosity_cluster_bootstrap']['pairwise_summary']:
        key = f"empirical_variance_ratio_{row['width_1']}_{row['width_2']}"
        res['headline'][key] = row['median_variance_ratio']
        res['headline'][f"{key}_min"] = row['min_variance_ratio']
        res['headline'][f"{key}_max"] = row['max_variance_ratio']
        res['headline'][f"{key}_significant_numerosities"] = row['significant_numerosities']
        res['headline'][f"{key}_max_adjusted_pvalue"] = row['max_adjusted_pvalue']
        res['headline'][f"{key}_max_adjusted_pvalue_mcse"] = row['max_adjusted_pvalue_raw_tail_mcse']
    for row in res['empirical_reanalysis']['participant_equal_weight_sensitivity']['contrasts']:
        key = f"empirical_equal_weight_ratio_{row['width_1']}_{row['width_2']}"
        res['headline'][key] = row['geometric_mean_within_participant_variance_ratio']
        res['headline'][f"{key}_lo"] = row['familywise_ci_variance_ratio'][0]
        res['headline'][f"{key}_hi"] = row['familywise_ci_variance_ratio'][1]
        res['headline'][f"{key}_p"] = row['pvalue_bonferroni']
    for row in res['empirical_reanalysis']['order_sequence_sensitivity']['contrasts']:
        pair_key = f"{row['width_1']}_{row['width_2']}"
        for order, key in (("Widening", "widening"), ("Narrowing", "narrowing")):
            estimate = row[key]
            short = order.lower()
            lo, hi = estimate['familywise_ci_variance_ratio']
            res['headline'][f"empirical_order_{short}_ratio_{pair_key}"] = estimate[
                'geometric_mean_within_participant_variance_ratio']
            res['headline'][f"empirical_order_{short}_ci_{pair_key}"] = f"{lo:.2f}–{hi:.2f}"
    for row in res['empirical_reanalysis']['within_block_position_sensitivity']['contrasts']:
        pair_key = f"{row['width_1']}_{row['width_2']}"
        half = row['block_half']
        res['headline'][f"empirical_position_ratio_{half}_{pair_key}"] = row[
            'variance_ratio_width_2_over_width_1']
        res['headline'][f"empirical_position_p_{half}_{pair_key}"] = row['pvalue_bonferroni']
        res['headline'][f"empirical_position_significant_{half}_{pair_key}"] = int(
            row['significant_familywise'])
    position_rows = res['empirical_reanalysis']['within_block_position_sensitivity']['contrasts']
    res['headline']['empirical_position_all_ratios_above_one'] = all(
        row['variance_ratio_width_2_over_width_1'] > 1 for row in position_rows
    )
    res['headline']['empirical_position_all_contrasts_supported'] = sum(
        row['significant_familywise'] for row in position_rows)
    res['headline']['empirical_position_min_adjusted_pvalue'] = min(
        row['pvalue_bonferroni'] for row in position_rows)
    res['headline']['empirical_position_max_adjusted_pvalue'] = max(
        row['pvalue_bonferroni'] for row in position_rows)
    res['headline']['empirical_position_primary_contrasts_supported'] = sum(
        row['significant_familywise'] for row in position_rows
        if (row['width_1'], row['width_2']) in {(20, 40), (20, 60)})
    ints = res['robustness']['integer_n']
    res['headline']['integer_ratio_n3'] = ints[2]['sigma_ratio_integer']
    res['headline']['integer_ratio_n10'] = ints[3]['sigma_ratio_integer']
    res['headline']['integer_ratio_n100'] = ints[5]['sigma_ratio_integer']
    res['headline']['el_mu_M_baseline'] = b['el_mu_M']['rho1.0_c100.0']
    res['headline']['el_mu_M_hard_rho1'] = b['el_mu_M']['rho1.0_chard']
    res['headline']['el_mu_M_hard_rho2'] = b['el_mu_M']['rho2.0_chard']
    res['headline']['el_mu_M_rho005'] = res['ablations']['el_mu_M_by_rho']['rho0.05']
    sc = res['signatures']['cases']['scarcity_with_technology_change (mu x2, e x0.71, K x0.8)']
    res['headline']['padamsey_like_slope'] = sc['slope']
    for row in pw:
        for N in cfg['power']['N_grid']:
            res['headline'][f"power_disc_ratio{row['price_ratio']}_N{N}"] = row[f'power_N{N}']
            lo, hi = row['mc_uncertainty'][f'power_N{N}']['wilson_95']
            res['headline'][f"power_disc_ratio{row['price_ratio']}_N{N}_mc95_lo"] = lo
            res['headline'][f"power_disc_ratio{row['price_ratio']}_N{N}_mc95_hi"] = hi
    for row in er:
        res['headline'][f"power_est_ratio{row['price_ratio']}_N{row['N']}"] = row['power']
        res['headline'][f"bias_est_median_ratio{row['price_ratio']}"] = row['bias_log_f_median']
        lo, hi = row['mc_uncertainty']['power']['wilson_95']
        res['headline'][f"power_est_ratio{row['price_ratio']}_N{row['N']}_mc95_lo"] = lo
        res['headline'][f"power_est_ratio{row['price_ratio']}_N{row['N']}_mc95_hi"] = hi
        res['headline'][f"nonconverged_fits_est_ratio{row['price_ratio']}"] = 1 - row['fit_diagnostics']['success_rate']
        res['headline'][f"boundary_fits_est_ratio{row['price_ratio']}"] = row['fit_diagnostics']['boundary_rate']
        res['headline'][f"estimation_fit_success_ratio{row['price_ratio']:g}"] = row['fit_diagnostics']['success_rate']
        res['headline'][f"estimation_boundary_rate_ratio{row['price_ratio']:g}"] = row['fit_diagnostics']['boundary_rate']
    fs = res['calibration']['estimation']['fitted_sd']
    res['headline']['calib_est_fitted_sd_narrow'], res['headline']['calib_est_fitted_sd_medium'], res['headline']['calib_est_fitted_sd_wide'] = fs
    fp = [row for row in pw if row['price_ratio'] == 1.0][0]
    res['headline']['false_pos_max'] = max(fp[f'power_N{N}'] for N in cfg['power']['N_grid'])
    rl = cfg['lapse_robustness']
    rng_l = np.random.default_rng(cfg['seed'] + 7)
    res['lapse_robustness'] = [recovery.discrimination_lapse_robustness(rng_l, res['calibration']['discrimination'], cfg['power'],
                                pr, rl['lapse_odds_mult'], rl['N_grid'], rl['reps']) for pr in (1.0, 2.0)]
    lr0, lr2 = res['lapse_robustness']
    res['headline'].update({'lapse_fp_shared': lr0['rejection_shared_N40'], 'lapse_fp_state': lr0['rejection_state_N40'],
                            'lapse_fp_state_N80': lr0['rejection_state_N80'],
                            'lapse_bias_shared': lr0['bias_log_f_shared'], 'lapse_bias_state': lr0['bias_log_f_state'],
                            'lapse_power_shared': lr2['rejection_shared_N40'], 'lapse_power_state': lr2['rejection_state_N40'],
                            'lapse_power_state_N80': lr2['rejection_state_N80']})
    res['headline']['false_pos_min'] = min(fp[f'power_N{N}'] for N in cfg['power']['N_grid'])
    ir = res['calibration']['estimation_integer_prior_refit']
    res['headline']['calib_int_nu'], res['headline']['calib_int_sigma0'] = ir['nu'], ir['sigma0']
    res['headline']['calib_int_nu_change'] = ir['nu'] / res['calibration']['estimation']['nu'] - 1
    pts = cfg['calibration']['discrimination']['points']
    by_w = {}
    for p in pts:
        by_w.setdefault(int(p[0]), []).append(p)
    parts, first = [], True
    for wv in sorted(by_w):
        items = []
        for p in by_w[wv]:
            items.append(f'{p[2]:.2f} at a difference of {p[1]:g}' if first else f'{p[2]:.2f} at {p[1]:g}')
            first = False
        parts.append(' and '.join(items) + f' for width {wv}')
    res['spec']['disc_points_text'] = ', and '.join(parts)
    est1 = [row for row in er if row['price_ratio'] == 1.0][0]
    res['headline']['false_pos_est'] = est1['power']
    res['headline']['nonconverged_fits_est_max'] = max(1 - row['fit_diagnostics']['success_rate'] for row in er)
    res['headline']['boundary_fits_est_max'] = max(row['fit_diagnostics']['boundary_rate'] for row in er)
    na = res['net_increment']
    res['headline']['net_over_gross_baseline'] = na['baseline']['net_over_gross']
    res['headline']['reallocated_share_baseline'] = 1 - na['baseline']['net_over_gross']
    res['headline']['lemma3_approx_rel_err'] = na['baseline_approx_rel_err']
    res['headline']['net_over_gross_M008'] = na['margin_sweep'][0]['net_over_gross']
    res['headline']['net_over_gross_M002'] = na['margin_sweep'][-1]['net_over_gross']
    res['headline']['same_price_net_over_gross_c1'] = na['same_price_varying_stiffness'][0]['net_over_gross']
    res['headline']['same_price_net_over_gross_c1e5'] = na['same_price_varying_stiffness'][-1]['net_over_gross']
    ins = res['integer_sweep']
    res['headline']['integer_no_effect_max'] = ins['max_nc_without_effect']
    res['headline']['integer_range_3_10_lo'], res['headline']['integer_range_3_10_hi'] = ins['range_3_10']
    res['headline']['integer_range_10_30_lo'], res['headline']['integer_range_10_30_hi'] = ins['range_10_30']
    dr = res['displacement_remark']
    res['headline']['displacement_fixed_slope'] = dr['fixed (D = 0.6 E0 in both states)']['slope']
    res['headline']['displacement_half_slope'] = dr['displacement falls by half the energy decline']['slope']
    res['headline']['displacement_more_slope'] = dr['displacement falls by 1.5 times the energy decline']['slope']
    res['headline']['displacement_baseline_D'] = dr['fixed (D = 0.6 E0 in both states)']['D0']
    res['headline']['displacement_large_change_D1'] = dr['displacement falls by 1.5 times the energy decline']['D1']
    ds = res['calibration']['estimation_discrete_prior_sd']
    res['headline']['calib_discrete_sd_narrow'], res['headline']['calib_discrete_sd_medium'], res['headline']['calib_discrete_sd_wide'] = ds
    ci_widths = [u['wilson_95'][1] - u['wilson_95'][0]
                 for group in (pw, er, res['lapse_robustness'])
                 for row in group
                 for u in row['mc_uncertainty'].values()]
    res['headline']['mc_ci95_width_min'], res['headline']['mc_ci95_width_max'] = min(ci_widths), max(ci_widths)
    res['headline']['disc_sd_subject_log_f'] = float(np.mean([row['sd_subject_log_f_hat'] for row in pw]))
    res['headline']['disc_max_abs_bias_log_f'] = float(max(abs(row['bias_log_f']) for row in pw))
    res['headline']['disc_fit_success_min'] = float(min(row['fit_diagnostics']['success_rate'] for row in pw))
    res['headline']['disc_fit_success_max'] = float(max(row['fit_diagnostics']['success_rate'] for row in pw))
    res['headline']['disc_fit_boundary_min'] = float(min(row['fit_diagnostics']['boundary_rate'] for row in pw))
    res['headline']['disc_fit_boundary_max'] = float(max(row['fit_diagnostics']['boundary_rate'] for row in pw))
    lapse_diags = [row['fit_diagnostics'][model]
                   for row in res['lapse_robustness'] for model in ('shared', 'state')]
    res['headline']['lapse_fit_success_min'] = float(min(row['success_rate'] for row in lapse_diags))
    res['headline']['lapse_fit_success_max'] = float(max(row['success_rate'] for row in lapse_diags))
    res['headline']['lapse_fit_boundary_min'] = float(min(row['boundary_rate'] for row in lapse_diags))
    res['headline']['lapse_fit_boundary_max'] = float(max(row['boundary_rate'] for row in lapse_diags))
    os.makedirs('figures', exist_ok=True)
    figures.fig_signatures(res, 'figures/fig1_signatures')
    figures.fig_power(res, 'figures/fig2_power')
    res['meta']['runtime_seconds'] = round(time.time() - t0, 1)
    json.dump(res, open('results.json', 'w'), indent=1, default=float)
    print('Done in %.1f s; results.json and figures written.' % (time.time() - t0))


if __name__ == '__main__':
    main()
```
