## Supplementary material for "The marginal brain": Replication package: analytics.py

```
"""Stage 2: symbolic derivations (sympy). Every claim in the propositions is checked here."""
import sympy as sp


def _el(expr, var):
    """Elasticity d log(expr) / d log(var)."""
    return sp.simplify(sp.diff(sp.log(expr), var) * var)


def derive():
    R = {}
    # ---------------- Encoding layer (Prat-Carrabin & Woodford) ----------------
    # Pointwise problem behind the optimal per-signal allocation:
    # min_s p^a / s^2 + nu * s, with s = sqrt(I1(x)) and nu the multiplier on  int s dx <= K
    p, nu, s, a = sp.symbols('p nu s a', positive=True)
    f = p**a / s**2 + nu * s
    s_star = sp.solve(sp.Eq(sp.diff(f, s), 0), s)[0]
    R['pointwise_s_star'] = str(s_star)
    R['s_star_propto_pi_pow_a3'] = bool(sp.simplify(sp.diff(sp.simplify(s_star / p**(a / 3)), p)) == 0)
    R['pointwise_soc_positive'] = bool(sp.simplify(sp.diff(f, s, 2).subs(s, s_star)).is_positive)

    # With sqrt(I1) = K pi^(a/3) / Z_a, Z_a = int pi^(a/3) dx, the loss for n signals is Z^3/(n K^2)
    n, K, lam, Z, pix = sp.symbols('n K lambda Z pi_x', positive=True)
    L = Z**3 / (n * K**2)
    nstar = sp.solve(sp.Eq(sp.diff(L + lam * n, n), 0), n)[0]
    Ix = nstar * (K * pix**(a / 3) / Z)**2
    sigma = sp.simplify(1 / sp.sqrt(Ix))
    Lstar = sp.simplify(L.subs(n, nstar))
    R['n_star'] = str(nstar)
    R['sigma_x'] = str(sigma)
    R['loss_star'] = str(Lstar)
    R['loss_equals_lambda_n'] = bool(sp.simplify(Lstar - lam * nstar) == 0)
    R['el_sigma_lambda'] = str(_el(sigma, lam))
    R['el_n_lambda'] = str(_el(nstar, lam))
    R['el_sigma_K'] = str(_el(sigma, K))
    R['el_n_K'] = str(_el(nstar, K))
    R['el_sigma_lambda_free_of_prior_task_capacity'] = bool(
        all(sp.diff(_el(sigma, lam), v) == 0 for v in (Z, pix, a, K)))

    # Uniform prior of width w: baseline exponents (Proposition 1)
    w, xx = sp.symbols('w x', positive=True)
    for aval, name in [(1, 'estimation'), (2, 'discrimination')]:
        Zu = sp.integrate((1 / w)**sp.Rational(aval, 3), (xx, 0, w))
        sig_u = sp.simplify(sigma.subs({a: aval, Z: Zu, pix: 1 / w}))
        R[f'uniform_{name}_sigma'] = str(sig_u)
        R[f'uniform_{name}_exp_w'] = str(_el(sig_u, w))
        R[f'uniform_{name}_exp_lambda'] = str(_el(sig_u, lam))
        R[f'uniform_{name}_n_exp_w'] = str(_el(nstar.subs(Z, Zu), w))

    # ---------------- Price of energy ----------------
    e, mu = sp.symbols('e mu', positive=True)
    E = sp.simplify((e * nstar).subs(lam, e * mu))           # task energy
    sig_mu = sigma.subs(lam, e * mu)
    R['task_energy'] = str(E)
    R['el_E_mu'] = str(_el(E, mu))
    R['loss_equals_mu_times_E'] = bool(sp.simplify(Lstar.subs(lam, e * mu) - mu * E) == 0)

    # Sign test (Proposition 6): d log sigma and d log E along price (x), energy-per-signal (y), capacity (z)
    x, y, z = sp.symbols('x y z', real=True)
    dls = _el(sig_mu, mu) * x + _el(sig_mu, e) * y + _el(sig_mu, K) * z
    dlE = _el(E, mu) * x + _el(E, e) * y + _el(E, K) * z
    R['dlog_sigma'] = str(sp.simplify(dls))
    R['dlog_E'] = str(sp.simplify(dlE))
    R['slope_price'] = str(sp.simplify((dls / dlE).subs({y: 0, z: 0})))
    R['slope_energy_per_signal'] = str(sp.simplify((dls / dlE).subs({x: 0, z: 0})))
    R['slope_capacity'] = str(sp.simplify((dls / dlE).subs({x: 0, y: 0})))
    R['sigma2E_invariant_to_mu'] = bool(sp.simplify(sp.diff(sp.simplify(sig_mu**2 * E), mu)) == 0)
    R['sigma2E_varies_with_e_or_K'] = bool(sp.simplify(sp.diff(sp.simplify(sig_mu**2 * E), K)) != 0)

    # Stakes enter only through lambda / beta (identification remark)
    beta = sp.symbols('beta', positive=True)
    nb = sp.solve(sp.Eq(sp.diff(beta * L + lam * n, n), 0), n)[0]
    R['stakes_enter_as_lambda_over_beta'] = bool(sp.simplify(nb - nstar.subs(lam, lam / beta)) == 0)

    # Robustness: cost lambda * n^kappa
    kap = sp.symbols('kappa', positive=True)
    nk = (Z**3 / (kap * lam * K**2))**(1 / (kap + 1))
    foc_k = sp.diff(L + lam * n**kap, n).subs(n, nk)
    vals = [{Z: 1.3, K: 0.7, lam: 0.01, kap: kv} for kv in (0.5, 1.0, 2.0, 3.0)]
    R['kappa_foc_check'] = bool(all(abs(float(foc_k.subs(vv))) < 1e-9 for vv in vals))
    sk = sp.simplify(1 / sp.sqrt(nk * (K * pix**(a / 3) / Z)**2))
    R['kappa_el_sigma_lambda'] = str(sp.simplify(_el(sk, lam)))
    Ek = e * nk.subs(lam, e * mu)
    skm = sk.subs(lam, e * mu)
    R['kappa_slope_price'] = str(sp.simplify(_el(skm, mu) / _el(Ek, mu)))
    R['kappa_slope_capacity'] = str(sp.simplify(_el(skm, K) / _el(Ek, K)))

    # ---------------- Regulated set point (Proposition 4) ----------------
    # Equilibrium: mu = c * (C + E_task + E_other - Bbar), E_task = sqrt(e) A / sqrt(mu), E_other = (v/mu)^(1/rho)
    # Written as F(mu) = mu/c + M - E_task - E_other = 0, with M = Bbar - C the reallocation margin
    M, c, v, rho, A = sp.symbols('M c v rho A', positive=True)
    Et = sp.sqrt(e) * A / sp.sqrt(mu)
    Eo = (v / mu)**(1 / rho)
    F = mu / c + M - Et - Eo
    Fmu = sp.simplify(sp.diff(F, mu))
    R['F_mu'] = str(Fmu)
    R['F_mu_positive'] = bool(Fmu.is_positive)
    R['dmu_dM_negative'] = bool(sp.simplify(-sp.diff(F, M) / Fmu).is_negative)
    R['dmu_dv_positive'] = bool(sp.simplify(-sp.diff(F, v) / Fmu).is_positive)
    R['dmu_dc_positive'] = bool(sp.simplify(-sp.diff(F, c) / Fmu).is_positive)
    elA = sp.simplify((A / mu) * (-sp.diff(F, A) / Fmu))
    target = (Et / mu) / Fmu
    R['el_mu_task_scale_equals_share_formula'] = bool(sp.simplify(elA - target) == 0)
    lim = sp.limit(F, c, sp.oo)
    R['hard_cap_limit'] = str(sp.simplify(lim))
    R['hard_cap_limit_is_M_minus_demand'] = bool(sp.simplify(lim - (M - Et - Eo)) == 0)

    # Closed-form identities and invariant checks.
    # Proposition 4 for every k > 0: F_mu = d/(k mu) + E/(2 mu) + E_o/(rho mu) with d = (mu/c)^(1/k)
    k = sp.symbols('k', positive=True)
    Fk = (mu / c)**(1 / k) + M - Et - Eo
    Fk_mu = sp.simplify(sp.diff(Fk, mu))
    R['F_mu_general_k_positive'] = bool(Fk_mu.is_positive)
    d = (mu / c)**(1 / k)
    R['F_mu_general_k_decomposition_ok'] = bool(sp.simplify(Fk_mu - (d / (k * mu) + Et / (2 * mu) + Eo / (rho * mu))) == 0)

    # Lemma 3 (net increment): perturb the task's scale from 0; first-order net increase in total use per unit of task energy
    eps = sp.symbols('epsilon', nonnegative=True)
    Fe = (mu / c)**(1 / k) + M - eps * Et - Eo
    dmu_deps = sp.simplify(-sp.diff(Fe, eps) / sp.diff(Fe, mu)).subs(eps, 0)
    dB_deps = sp.simplify(sp.diff(d, mu) * dmu_deps)
    phi = sp.simplify(dB_deps / Et)
    phi_target = (d / k) / (d / k + Eo / rho)
    R['lemma3_phi'] = str(phi)
    R['lemma3_phi_formula_ok'] = bool(sp.simplify(phi - phi_target) == 0)
    R['lemma3_phi_k1_ok'] = bool(sp.simplify(phi.subs(k, 1) - 1 / (1 + c * Eo / (rho * mu))) == 0)
    # elasticity of phi with respect to the price: (1 - phi)(1/k + 1/rho) > 0
    el_phi = sp.simplify(sp.diff(sp.log(phi_target), mu) * mu)
    R['lemma3_el_phi_mu_ok'] = bool(sp.simplify(el_phi - (1 - phi_target) * (1 / k + 1 / rho)) == 0)
    R['lemma3_hard_cap_limit_phi'] = str(sp.limit(phi_target.subs(k, 1), c, sp.oo))

    # Remark (measurement): measured increment E - D; slope scales by (E - D)/(E (1 - delta)), delta = dD/dE
    Ev, Dv, dE, delta = sp.symbols('E_v D_v dE delta', real=True)
    dlogEm = (dE - delta * dE) / (Ev - Dv)
    factor = sp.simplify((dE / Ev) / dlogEm)
    R['remark_slope_factor'] = str(factor)
    R['remark_proportional_displacement_preserves_slope'] = bool(sp.simplify(factor.subs(Dv, delta * Ev)) == 1)
    return R


if __name__ == '__main__':
    import json
    print(json.dumps(derive(), indent=1))
```
