## Supplementary material for "The marginal brain": Replication package: behavior.py

```
"""Behavioral layer: the Gaussian-representation model of Prat-Carrabin & Woodford with a state factor f.
Representation sd = nu * w^alpha * f, with f = (mu_state / mu_baseline)^(1/4) (Proposition 2)."""
import numpy as np
from numpy.polynomial.hermite_e import hermegauss
from scipy.stats import norm
from scipy.optimize import minimize, least_squares
from scipy.special import expit, logit

_GH_X, _GH_W = hermegauss(24)
_GH_W = _GH_W / _GH_W.sum()
_TINY = 1e-300


def posterior_mean_uniform(r, s, lo, hi):
    """Posterior mean of x given r ~ N(x, s^2) and a uniform prior on [lo, hi] (truncated normal mean)."""
    al = (lo - r) / s
    be = (hi - r) / s
    Zs = np.maximum(norm.cdf(be) - norm.cdf(al), 1e-12)
    return r + s * (norm.pdf(al) - norm.pdf(be)) / Zs


def response_probs(m, sigma0, lo, hi, k):
    """P(response = k) for motor noise N(m, sigma0^2) truncated to [lo-0.5, hi+0.5] and rounded."""
    num = norm.cdf((k + 0.5 - m) / sigma0) - norm.cdf((k - 0.5 - m) / sigma0)
    den = norm.cdf((hi + 0.5 - m) / sigma0) - norm.cdf((lo - 0.5 - m) / sigma0)
    return num / np.maximum(den, 1e-12)


def estimation_moments(x, s, sigma0, lo, hi):
    """Mean and sd of the response to number x."""
    ks = np.arange(lo, hi + 1)
    r = x + s * _GH_X
    m = posterior_mean_uniform(r, s, lo, hi)
    P = response_probs(m[:, None], sigma0, lo, hi, ks[None, :])
    P = (_GH_W[:, None] * P).sum(axis=0)
    P /= P.sum()
    mean = (P * ks).sum()
    return mean, np.sqrt((P * (ks - mean)**2).sum())


def calibrate_estimation(cfg):
    lows, highs, x0, target = cfg['lows'], cfg['highs'], cfg['x'], np.array(cfg['sd'])
    alpha = cfg['alpha']

    def resid(p):
        nu, s0 = np.exp(p)
        out = []
        for lo, hi in zip(lows, highs):
            w = hi - lo
            out.append(estimation_moments(x0, nu * w**alpha, s0, lo, hi)[1])
        return np.array(out) - target
    sol = least_squares(resid, np.log([1.0, 2.0]))
    nu, s0 = np.exp(sol.x)
    return dict(nu=float(nu), sigma0=float(s0), fitted_sd=list(map(float, resid(sol.x) + target)), target_sd=list(map(float, target)))


def calibrate_discrimination(cfg):
    pts = np.array(cfg['points'])
    alpha = cfg['alpha']

    def resid(p):
        nu = np.exp(p[0]); eta = expit(p[1])
        w, d, pc = pts[:, 0], pts[:, 1], pts[:, 2]
        return eta / 2 + (1 - eta) * norm.cdf(d / (np.sqrt(2) * nu * w**alpha)) - pc
    sol = least_squares(resid, [np.log(0.5), logit(0.05)])
    return dict(nu=float(np.exp(sol.x[0])), eta=float(expit(sol.x[1])), residuals=list(map(float, resid(sol.x))))


# ---------------- simulation ----------------
def simulate_estimation(rng, xs, s, sigma0, lo, hi):
    r = xs + s * rng.standard_normal(xs.shape)
    m = posterior_mean_uniform(r, s, lo, hi)
    a = norm.cdf((lo - 0.5 - m) / sigma0)
    b = norm.cdf((hi + 0.5 - m) / sigma0)
    u = a + (b - a) * rng.random(xs.shape)
    resp = m + sigma0 * norm.ppf(np.clip(u, 1e-12, 1 - 1e-12))
    return np.clip(np.round(resp), lo, hi)


def simulate_discrimination(rng, T, s, eta, lo, hi):
    xr = rng.integers(lo, hi + 1, size=(T, 5)).mean(axis=1)
    xb = rng.integers(lo, hi + 1, size=(T, 5)).mean(axis=1)
    p = eta / 2 + (1 - eta) * norm.cdf((xr - xb) / (np.sqrt(2) * s))
    choice_r = rng.random(T) < p
    return xr - xb, choice_r


# ---------------- likelihoods ----------------
def nll_estimation(par, data, lo, hi, alpha):
    """par = (log nu, log sigma0, log f); data = list of (xs, resp, state) with state 0/1."""
    nu, s0, f = np.exp(np.clip(par, -12.0, 12.0))
    w = hi - lo
    tot = 0.0
    for xs, resp, state in data:
        s = nu * w**alpha * (f if state == 1 else 1.0)
        r = xs[:, None] + s * _GH_X[None, :]
        m = posterior_mean_uniform(r, s, lo, hi)
        P = response_probs(m, s0, lo, hi, resp[:, None])
        lik = (P * _GH_W[None, :]).sum(axis=1)
        tot -= np.log(np.maximum(lik, _TINY)).sum()
    return tot


def nll_discrimination(par, data, w, alpha):
    """par = (log nu, logit eta, log f); data = list of (diff, choice_r, state). Returns (nll, gradient)."""
    par = np.clip(par, [-12.0, -15.0, -5.0], [12.0, 15.0, 5.0])
    nu, eta, f = np.exp(par[0]), expit(par[1]), np.exp(par[2])
    tot = 0.0
    g = np.zeros(3)
    for d, ch, state in data:
        s = nu * w**alpha * (f if state == 1 else 1.0)
        z = d / (np.sqrt(2) * s)
        Phi, phi = norm.cdf(z), norm.pdf(z)
        p = np.clip(eta / 2 + (1 - eta) * Phi, 1e-12, 1 - 1e-12)
        tot -= np.where(ch, np.log(p), np.log1p(-p)).sum()
        dl_dp = -np.where(ch, 1.0 / p, -1.0 / (1 - p))
        dp_dlognu = (1 - eta) * phi * (-z)
        dp_dlogiteta = eta * (1 - eta) * (0.5 - Phi)
        g[0] += (dl_dp * dp_dlognu).sum()
        g[1] += (dl_dp * dp_dlogiteta).sum()
        if state == 1:
            g[2] += (dl_dp * dp_dlognu).sum()
    return tot, g


def fit(nll, x0, args, jac=False):
    """Fit a likelihood on the same parameter bounds used by its objective.

    Returning convergence and boundary diagnostics prevents downstream recovery
    summaries from silently treating failed or constrained fits as ordinary fits.
    """
    if nll not in _FIT_BOUNDS:
        raise ValueError(f"No parameter bounds registered for {nll.__name__}")
    bounds = _FIT_BOUNDS[nll]
    if len(x0) != len(bounds):
        raise ValueError(f"{nll.__name__} expects {len(bounds)} parameters, received {len(x0)}")
    res = minimize(nll, x0, args=args, method='L-BFGS-B', jac=jac, bounds=bounds,
                   options={'maxiter': 2000, 'ftol': 1e-12, 'gtol': 1e-7, 'maxls': 50})
    at_bound = [bool(abs(float(x) - lo) <= 1e-5 or abs(float(x) - hi) <= 1e-5)
                for x, (lo, hi) in zip(res.x, bounds)]
    diagnostics = {
        'success': bool(res.success),
        'status': int(res.status),
        'message': str(res.message),
        'iterations': int(res.nit),
        'at_bound': at_bound,
    }
    return res.x, bool(res.success), diagnostics


def nll_discrimination_state_lapse(par, data, w, alpha):
    """par = (log nu, logit eta in state 0, log f, logit eta in state 1). Returns (nll, gradient)."""
    par = np.clip(par, [-12.0, -15.0, -5.0, -15.0], [12.0, 15.0, 5.0, 15.0])
    nu, f = np.exp(par[0]), np.exp(par[2])
    tot = 0.0
    g = np.zeros(4)
    for d, ch, state in data:
        eta = expit(par[3] if state == 1 else par[1])
        s = nu * w**alpha * (f if state == 1 else 1.0)
        z = d / (np.sqrt(2) * s)
        Phi, phi = norm.cdf(z), norm.pdf(z)
        p = np.clip(eta / 2 + (1 - eta) * Phi, 1e-12, 1 - 1e-12)
        tot -= np.where(ch, np.log(p), np.log1p(-p)).sum()
        dl_dp = -np.where(ch, 1.0 / p, -1.0 / (1 - p))
        dp_dlognu = (1 - eta) * phi * (-z)
        dp_dlogiteta = eta * (1 - eta) * (0.5 - Phi)
        g[0] += (dl_dp * dp_dlognu).sum()
        g[3 if state == 1 else 1] += (dl_dp * dp_dlogiteta).sum()
        if state == 1:
            g[2] += (dl_dp * dp_dlognu).sum()
    return tot, g


_FIT_BOUNDS = {
    nll_estimation: [(-12.0, 12.0), (-12.0, 12.0), (-12.0, 12.0)],
    nll_discrimination: [(-12.0, 12.0), (-15.0, 15.0), (-5.0, 5.0)],
    nll_discrimination_state_lapse: [(-12.0, 12.0), (-15.0, 15.0), (-5.0, 5.0), (-15.0, 15.0)],
}
```
