## Supplementary material for "The marginal brain": Replication package: empirical.py

```
"""Participant-clustered reanalysis of published estimation trials.

The width comparisons are made separately at each shared numerosity.  Within
each participant × width × numerosity cell, responses are centered on that
cell's mean; sums of squares and residual degrees of freedom are then pooled
over participant clusters.  This avoids treating repeated trials as
independent observations and avoids comparing widths with different mixes of
stimulus numerosity.
"""
import csv
from collections import defaultdict
from itertools import combinations
from pathlib import Path

import numpy as np


def _wilson_interval(successes, trials, alpha=0.05):
    """Wilson interval for a binomial Monte Carlo tail probability."""
    from scipy.stats import norm
    z = float(norm.ppf(1 - alpha / 2))
    p = successes / trials
    den = 1 + z * z / trials
    center = (p + z * z / (2 * trials)) / den
    half = z * np.sqrt(p * (1 - p) / trials + z * z / (4 * trials * trials)) / den
    return [float(max(0.0, center - half)), float(min(1.0, center + half))]


def _bootstrap_summary(centered, theta, family_size, alpha=0.05):
    """Summarize centered draws with intervals inverted from the absolute-tail test."""
    reps = len(centered)
    tail_count = int(np.count_nonzero(np.abs(centered) >= abs(theta)))
    pvalue = (1 + tail_count) / (reps + 1)
    # Use the plus-one estimate in the SE so a zero observed tail count does
    # not misleadingly imply zero Monte Carlo uncertainty. Also retain a
    # Wilson interval for the underlying simulated exceedance probability.
    pvalue_mcse = float(np.sqrt(pvalue * (1.0 - pvalue) / reps))
    tail_fraction_mc_95 = _wilson_interval(tail_count, reps)

    # Invert the exact plus-one Monte Carlo decision rule used above. A value
    # is significant after Bonferroni correction iff its observed absolute
    # log ratio exceeds this critical absolute centered-bootstrap deviation.
    # This keeps the plotted intervals and corrected p-values decision-dual,
    # including the finite-replication plus-one adjustment.
    adjusted_alpha = alpha / family_size
    max_tail_count = int(np.ceil(adjusted_alpha * (reps + 1) - 1.0) - 1)
    if max_tail_count < 0:
        critical = np.inf
    else:
        ordered = np.sort(np.abs(centered))
        critical = float(ordered[reps - max_tail_count - 1])
    interval = np.exp([theta - critical, theta + critical])
    return {
        "pvalue_bootstrap": float(pvalue),
        "pvalue_bootstrap_mcse": pvalue_mcse,
        "pvalue_bootstrap_mc_95": tail_fraction_mc_95,
        "familywise_ci_variance_ratio": [float(interval[0]), float(interval[1])],
        "familywise_ci_log_critical": float(critical),
    }


def _cluster_bootstrap_log_ratio(sse, df, i, j, theta, reps, seed,
                                 family_size, strata_indices=None, alpha=0.05,
                                 chunk_size=10_000):
    """Centered paired participant bootstrap, optionally stratified by sequence."""
    n_subjects = sse.shape[0]
    rng = np.random.default_rng(seed)
    if strata_indices is None:
        strata_indices = [np.arange(n_subjects)]
    if sum(len(idx) for idx in strata_indices) != n_subjects or any(len(idx) == 0 for idx in strata_indices):
        raise ValueError("bootstrap strata must partition all participants and be nonempty")
    centered = np.empty(reps, dtype=float)
    kept = 0
    while kept < reps:
        size = min(chunk_size, reps - kept)
        draw = np.concatenate([
            idx[rng.integers(0, len(idx), size=(size, len(idx)))]
            for idx in strata_indices
        ], axis=1)
        ss = sse[draw].sum(axis=1)
        dof = df[draw].sum(axis=1)
        valid = (ss[:, i] > 0) & (ss[:, j] > 0) & (dof[:, i] > 0) & (dof[:, j] > 0)
        if np.any(valid):
            boot_theta = np.log(ss[valid, j] / dof[valid, j]) - np.log(ss[valid, i] / dof[valid, i])
            values = boot_theta - theta
            centered[kept:kept + len(values)] = values
            kept += len(values)
    return _bootstrap_summary(centered, theta, family_size, alpha)


def _centered_mean_bootstrap(values, reps, rng, family_size, alpha=0.05,
                             chunk_size=10_000):
    """Bootstrap the mean log ratio, keeping the observed participant count."""
    values = np.asarray(values, dtype=float)
    theta = float(values.mean())
    centered = np.empty(int(reps), dtype=float)
    kept = 0
    while kept < len(centered):
        size = min(chunk_size, len(centered) - kept)
        draw = rng.integers(0, len(values), size=(size, len(values)))
        centered[kept:kept + size] = values[draw].mean(axis=1) - theta
        kept += size
    return theta, _bootstrap_summary(centered, theta, family_size, alpha)


def _centered_stratified_mean_bootstrap(values, strata_indices, reps, rng,
                                        family_size, alpha=0.05,
                                        chunk_size=10_000):
    """Bootstrap the mean while preserving the participant count in each stratum."""
    values = np.asarray(values, dtype=float)
    if sum(len(idx) for idx in strata_indices) != len(values) or any(len(idx) == 0 for idx in strata_indices):
        raise ValueError("bootstrap strata must partition the included participants and be nonempty")
    theta = float(values.mean())
    centered = np.empty(int(reps), dtype=float)
    kept = 0
    while kept < len(centered):
        size = min(chunk_size, len(centered) - kept)
        draw = np.concatenate([
            idx[rng.integers(0, len(idx), size=(size, len(idx)))]
            for idx in strata_indices
        ], axis=1)
        centered[kept:kept + size] = values[draw].mean(axis=1) - theta
        kept += size
    return theta, _bootstrap_summary(centered, theta, family_size, alpha)


def _centered_difference_bootstrap(values_a, values_b, reps, rng,
                                   family_size, alpha=0.05,
                                   chunk_size=10_000):
    """Bootstrap a difference between independent sequence-specific means."""
    values_a = np.asarray(values_a, dtype=float)
    values_b = np.asarray(values_b, dtype=float)
    theta = float(values_a.mean() - values_b.mean())
    centered = np.empty(int(reps), dtype=float)
    kept = 0
    while kept < len(centered):
        size = min(chunk_size, len(centered) - kept)
        draw_a = rng.integers(0, len(values_a), size=(size, len(values_a)))
        draw_b = rng.integers(0, len(values_b), size=(size, len(values_b)))
        boot_difference = values_a[draw_a].mean(axis=1) - values_b[draw_b].mean(axis=1)
        centered[kept:kept + size] = boot_difference - theta
        kept += size
    return theta, _bootstrap_summary(centered, theta, family_size, alpha)


def reanalyze_estimation(path, widths=(20, 40, 60), anchor_number=60,
                         shared_numbers=range(50, 71), published_sd=(4.2, 6.8, 8.4),
                         familywise_alpha=0.05, bootstrap_reps=500_000,
                         bootstrap_seed=20261006,
                         equal_weight_bootstrap_reps=500_000,
                         position_bootstrap_reps=100_000):
    """Recreate published anchor SDs and compare widths at fixed numerosity.

    Bootstrap resampling treats participants, rather than trials, as clusters.
    For each fixed numerosity x, the estimate is

        V[x,w] = sum_i SSE[i,x,w] / sum_i df[i,x,w].

    A cell with fewer than two trials contributes zero residual degrees of
    freedom. The p-value is the centered two-sided cluster-bootstrap tail
    probability for log(V[x,w2] / V[x,w1]); p-values are Bonferroni-adjusted
    over every planned shared numerosity and pairwise width contrast. The
    participant-equal-weighted sensitivity aggregates each participant's
    within-cell residual variance over the common numerosity support, computes
    paired log variance ratios, and resamples participants as clusters.
    """
    widths = tuple(int(w) for w in widths)
    if len(widths) < 2 or len(set(widths)) != len(widths):
        raise ValueError("widths must contain at least two distinct conditions")
    shared_numbers = tuple(int(x) for x in shared_numbers)
    if not shared_numbers or len(set(shared_numbers)) != len(shared_numbers):
        raise ValueError("shared_numbers must be a nonempty set of distinct numerosities")
    pairs = list(combinations(range(len(widths)), 2))
    planned_family_size = len(shared_numbers) * len(pairs)

    groups = defaultdict(list)
    cells = defaultdict(list)
    position_cells = defaultdict(list)
    subject_orders = {}
    first_trial_by_subject_width = {}
    trial_indices_by_subject_width = defaultdict(list)
    subjects = set()
    n_rows = 0
    with Path(path).open("r", encoding="utf-8-sig", newline="") as stream:
        for row in csv.DictReader(stream):
            width = int(float(row["Delta"]))
            x = int(float(row["Number"]))
            response = float(row["Response"])
            subject = str(row["ID"])
            order = str(row["Order"]).strip()
            t_all = int(float(row["tAll"]))
            t_loc = int(float(row["tLoc"]))
            if not 1 <= t_loc <= 120:
                raise ValueError(f"Within-block trial index is outside 1–120 for participant {subject}")
            if order not in {"Widening", "Narrowing"}:
                raise ValueError(f"Unexpected block-order label for participant {subject}: {order!r}")
            if subject in subject_orders and subject_orders[subject] != order:
                raise ValueError(f"Block-order label changes within participant {subject}")
            subject_orders[subject] = order
            block_key = (subject, width)
            trial_indices_by_subject_width[block_key].append(t_loc)
            first_trial_by_subject_width[block_key] = min(
                t_all, first_trial_by_subject_width.get(block_key, t_all))
            groups[(width, x)].append(response)
            if x in shared_numbers and width in widths:
                cells[(subject, width, x)].append(response)
                position_cells[(subject, width, x)].append((t_loc, response))
            subjects.add(subject)
            n_rows += 1

    expected_trial_indices = set(range(1, 121))
    for block_key, trial_indices in trial_indices_by_subject_width.items():
        if len(trial_indices) != 120 or set(trial_indices) != expected_trial_indices:
            subject, width = block_key
            raise ValueError(
                f"Participant {subject}, width {width} must have exactly one row for each tLoc 1–120"
            )

    observed_by_width = {
        width: {x for w, x in groups if w == width}
        for width in widths
    }
    common_observed = set.intersection(*(observed_by_width[w] for w in widths))
    absent = [x for x in shared_numbers if x not in common_observed]
    if absent:
        raise ValueError(f"Configured shared numerosities are not observed in every width: {absent}")

    anchors = []
    for width, target in zip(widths, published_sd):
        values = np.asarray(groups[(width, int(anchor_number))], dtype=float)
        if values.size < 2:
            raise ValueError(f"No repeated observations at anchor x={anchor_number}, width={width}")
        observed_sd = float(np.std(values, ddof=1))
        anchors.append({
            "prior_width": width,
            "numerosity": int(anchor_number),
            "n_trials": int(values.size),
            "observed_sd": observed_sd,
            "published_sd": float(target),
            "difference": observed_sd - float(target),
        })

    subject_ids = sorted(subjects)
    subject_index = {subject: i for i, subject in enumerate(subject_ids)}
    order_labels = ("Widening", "Narrowing")
    expected_width_order = {
        "Widening": tuple(sorted(widths)),
        "Narrowing": tuple(sorted(widths, reverse=True)),
    }
    observed_width_order = {}
    for subject in subject_ids:
        by_period = sorted(
            (first_trial_by_subject_width[(subject, width)], width)
            for width in widths
        )
        observed = tuple(width for _, width in by_period)
        order = subject_orders[subject]
        if observed != expected_width_order[order]:
            raise ValueError(
                f"Observed block order for participant {subject} conflicts with {order}: {observed}"
            )
        observed_width_order.setdefault(order, observed)
    order_array = np.asarray([subject_orders[s] for s in subject_ids], dtype=object)
    order_indices = [np.flatnonzero(order_array == order) for order in order_labels]
    if any(len(idx) == 0 for idx in order_indices):
        raise ValueError("Both Widening and Narrowing sequences must be represented")
    order_counts = {label: int(len(idx)) for label, idx in zip(order_labels, order_indices)}
    width_index = {width: i for i, width in enumerate(widths)}
    contrasts = []
    per_numerosity = []
    equal_sse = np.zeros((len(subject_ids), len(widths)), dtype=float)
    equal_df = np.zeros_like(equal_sse)
    rng_seeds = np.random.SeedSequence(int(bootstrap_seed)).spawn(len(shared_numbers) * len(pairs))
    seed_index = 0
    for x in shared_numbers:
        sse = np.zeros((len(subject_ids), len(widths)), dtype=float)
        df = np.zeros_like(sse)
        trials = np.zeros_like(sse, dtype=int)
        for subject in subject_ids:
            for width in widths:
                values = cells.get((subject, width, x), ())
                if len(values) < 2:
                    continue
                values = np.asarray(values, dtype=float)
                i, j = subject_index[subject], width_index[width]
                centered = values - values.mean()
                sse[i, j] = float(centered @ centered)
                df[i, j] = len(values) - 1
                trials[i, j] = len(values)
        pooled_df = df.sum(axis=0)
        pooled_sse = sse.sum(axis=0)
        equal_sse += sse
        equal_df += df
        if np.any(pooled_df <= 0) or np.any(pooled_sse <= 0):
            raise ValueError(f"A width has no within-cell residual information at numerosity {x}")
        pooled_variance = pooled_sse / pooled_df
        paired_support = {
            f"{widths[i]}/{widths[j]}": int(np.count_nonzero((df[:, i] > 0) & (df[:, j] > 0)))
            for i, j in pairs
        }
        x_contrasts = []
        for i, j in pairs:
            width_1, width_2 = widths[i], widths[j]
            theta = float(np.log(pooled_variance[j] / pooled_variance[i]))
            requested_reps = int(bootstrap_reps)
            seed_words = rng_seeds[seed_index].generate_state(2, dtype=np.uint32)
            seed_index += 1
            boot = _cluster_bootstrap_log_ratio(
                sse, df, i, j, theta, requested_reps,
                seed=int(seed_words[0]) ^ (int(seed_words[1]) << 1),
                family_size=planned_family_size,
                strata_indices=order_indices,
                alpha=familywise_alpha,
            )
            row = {
                "numerosity": int(x),
                "width_1": width_1,
                "width_2": width_2,
                "variance_ratio_width_2_over_width_1": float(np.exp(theta)),
                "log_variance_ratio": theta,
                **boot,
                "pvalue_bonferroni": float(min(1.0, boot["pvalue_bootstrap"] * planned_family_size)),
                "bootstrap_reps": requested_reps,
                "participants_with_df_width_1": int(np.count_nonzero(df[:, i] > 0)),
                "participants_with_df_width_2": int(np.count_nonzero(df[:, j] > 0)),
                "participants_with_df_both_widths": paired_support[f"{width_1}/{width_2}"],
                "pooled_residual_df_width_1": int(pooled_df[i]),
                "pooled_residual_df_width_2": int(pooled_df[j]),
                "significant_familywise": bool(boot["pvalue_bootstrap"] * planned_family_size < familywise_alpha),
            }
            contrasts.append(row)
            x_contrasts.append(row)
        per_numerosity.append({
            "numerosity": int(x),
            "pooled_variance_by_width": {
                str(width): float(pooled_variance[j]) for j, width in enumerate(widths)
            },
            "residual_df_by_width": {
                str(width): int(pooled_df[j]) for j, width in enumerate(widths)
            },
            "participants_with_residual_df_by_width": {
                str(width): int(np.count_nonzero(df[:, j] > 0)) for j, width in enumerate(widths)
            },
            "participants_with_residual_df_both_widths": paired_support,
            "contrasts": x_contrasts,
        })

    summaries = []
    for i, j in pairs:
        width_1, width_2 = widths[i], widths[j]
        rows = [r for r in contrasts if r["width_1"] == width_1 and r["width_2"] == width_2]
        ratios = np.asarray([r["variance_ratio_width_2_over_width_1"] for r in rows])
        adjusted = np.asarray([r["pvalue_bonferroni"] for r in rows])
        largest_adjusted = max(rows, key=lambda r: r["pvalue_bonferroni"])
        summaries.append({
            "width_1": width_1,
            "width_2": width_2,
            "n_numerosities": len(rows),
            "median_variance_ratio": float(np.median(ratios)),
            "min_variance_ratio": float(np.min(ratios)),
            "max_variance_ratio": float(np.max(ratios)),
            "significant_numerosities": int(sum(r["significant_familywise"] for r in rows)),
            "max_adjusted_pvalue": float(np.max(adjusted)),
            "max_adjusted_pvalue_raw_tail_mcse": float(largest_adjusted["pvalue_bootstrap_mcse"]),
            "min_adjusted_pvalue": float(np.min(adjusted)),
        })

    # Equal-weight sensitivity: each participant contributes one within-person
    # variance ratio over the configured common numerosity support. This is a
    # distinct estimand from pooled SSE / pooled df.
    participant_variances = np.full_like(equal_sse, np.nan)
    np.divide(equal_sse, equal_df, out=participant_variances, where=equal_df > 0)
    complete = np.all(np.isfinite(participant_variances) & (participant_variances > 0), axis=1)
    complete_order = order_array[complete]
    complete_strata = [np.flatnonzero(complete_order == order) for order in order_labels]
    if any(len(idx) == 0 for idx in complete_strata):
        raise ValueError("Each block-order sequence needs complete participants for the equal-weight sensitivity")
    equal_weight_rows = []
    equal_weight_seed_seq = np.random.SeedSequence(int(bootstrap_seed) + 9173).spawn(len(pairs))
    for pair_no, (i, j) in enumerate(pairs):
        log_ratios = np.log(participant_variances[complete, j] / participant_variances[complete, i])
        rng = np.random.default_rng(equal_weight_seed_seq[pair_no])
        theta, boot = _centered_stratified_mean_bootstrap(
            log_ratios, complete_strata, equal_weight_bootstrap_reps, rng,
            len(pairs), familywise_alpha)
        equal_weight_rows.append({
            "width_1": widths[i],
            "width_2": widths[j],
            "n_participants": int(np.count_nonzero(complete)),
            "geometric_mean_within_participant_variance_ratio": float(np.exp(theta)),
            "median_within_participant_variance_ratio": float(np.median(np.exp(log_ratios))),
            **boot,
            "pvalue_bonferroni": float(min(1.0, boot["pvalue_bootstrap"] * len(pairs))),
            "significant_familywise": bool(boot["pvalue_bootstrap"] * len(pairs) < familywise_alpha),
            "bootstrap_reps": int(equal_weight_bootstrap_reps),
        })

    # Sequence sensitivity: equal-participant mean log variance ratios within
    # each observed sequence, with familywise intervals over six group-specific
    # contrasts. The sequence difference is descriptive because assignment to
    # order is not documented as randomized and block period is confounded with
    # width within each sequence.
    sequence_contrasts = []
    sequence_seed_seq = np.random.SeedSequence(int(bootstrap_seed) + 20_261_013).spawn(
        len(pairs) * 3)
    seed_index = 0
    for i, j in pairs:
        group_rows = {}
        group_logs = {}
        for order in order_labels:
            mask = (order_array == order) & complete
            log_ratios = np.log(
                participant_variances[mask, j] / participant_variances[mask, i]
            )
            if len(log_ratios) < 2:
                raise ValueError(f"At least two complete participants are required in {order}")
            rng = np.random.default_rng(sequence_seed_seq[seed_index])
            seed_index += 1
            theta, boot = _centered_mean_bootstrap(
                log_ratios, equal_weight_bootstrap_reps, rng,
                family_size=len(order_labels) * len(pairs), alpha=familywise_alpha)
            row = {
                "order": order,
                "n_participants": int(len(log_ratios)),
                "geometric_mean_within_participant_variance_ratio": float(np.exp(theta)),
                **boot,
                "pvalue_bonferroni": float(min(
                    1.0, boot["pvalue_bootstrap"] * len(order_labels) * len(pairs))),
                "bootstrap_reps": int(equal_weight_bootstrap_reps),
            }
            group_rows[order] = row
            group_logs[order] = log_ratios
        rng = np.random.default_rng(sequence_seed_seq[seed_index])
        seed_index += 1
        interaction_theta, interaction_boot = _centered_difference_bootstrap(
            group_logs["Widening"], group_logs["Narrowing"],
            equal_weight_bootstrap_reps, rng, family_size=len(pairs),
            alpha=familywise_alpha)
        sequence_contrasts.append({
            "width_1": widths[i],
            "width_2": widths[j],
            "widening": group_rows["Widening"],
            "narrowing": group_rows["Narrowing"],
            "difference_log_ratio_widening_minus_narrowing": {
                "estimate": interaction_theta,
                **interaction_boot,
                "pvalue_bonferroni": float(min(
                    1.0, interaction_boot["pvalue_bootstrap"] * len(pairs))),
                "bootstrap_reps": int(equal_weight_bootstrap_reps),
            },
        })

    # Within-block position sensitivity: split each 120-trial block into its
    # first and second halves, re-center within participant × width × numerosity
    # × half, and pool cell residual SSE/df. This checks whether the matched-
    # numerosity pattern is present in both halves; it does not separate width
    # from block period or identify arbitrary carry-over effects.
    half_labels = ("first_half", "second_half")
    position_sse = np.zeros((len(subject_ids), len(widths), len(half_labels)), dtype=float)
    position_df = np.zeros_like(position_sse)
    position_trials = np.zeros_like(position_sse, dtype=int)
    for (subject, width, _x), observations in position_cells.items():
        si, wi = subject_index[subject], width_index[width]
        obs = np.asarray(observations, dtype=float)
        halves = (obs[:, 0] <= 60, obs[:, 0] > 60)
        for hi, mask in enumerate(halves):
            values = obs[mask, 1]
            if len(values) < 2:
                continue
            centered = values - values.mean()
            position_sse[si, wi, hi] += float(centered @ centered)
            position_df[si, wi, hi] += len(values) - 1
            position_trials[si, wi, hi] += len(values)

    position_contrasts = []
    position_seed_seq = np.random.SeedSequence(int(bootstrap_seed) + 20_261_014).spawn(
        len(half_labels) * len(pairs))
    seed_index = 0
    for hi, half in enumerate(half_labels):
        for i, j in pairs:
            sse, df = position_sse[:, :, hi], position_df[:, :, hi]
            pooled_df, pooled_sse = df.sum(axis=0), sse.sum(axis=0)
            if np.any(pooled_df <= 0) or np.any(pooled_sse <= 0):
                raise ValueError(f"A width has no within-cell residual information in {half}")
            pooled_variance = pooled_sse / pooled_df
            theta = float(np.log(pooled_variance[j] / pooled_variance[i]))
            seed_words = position_seed_seq[seed_index].generate_state(2, dtype=np.uint32)
            seed_index += 1
            boot = _cluster_bootstrap_log_ratio(
                sse, df, i, j, theta, int(position_bootstrap_reps),
                seed=int(seed_words[0]) ^ (int(seed_words[1]) << 1),
                family_size=len(half_labels) * len(pairs),
                strata_indices=order_indices,
                alpha=familywise_alpha,
            )
            position_contrasts.append({
                "block_half": half,
                "width_1": widths[i],
                "width_2": widths[j],
                "variance_ratio_width_2_over_width_1": float(np.exp(theta)),
                "log_variance_ratio": theta,
                **boot,
                "pvalue_bonferroni": float(min(
                    1.0, boot["pvalue_bootstrap"] * len(half_labels) * len(pairs))),
                "significant_familywise": bool(
                    boot["pvalue_bootstrap"] * len(half_labels) * len(pairs)
                    < familywise_alpha),
                "bootstrap_reps": int(position_bootstrap_reps),
                "participants_with_df_width_1": int(np.count_nonzero(df[:, i] > 0)),
                "participants_with_df_width_2": int(np.count_nonzero(df[:, j] > 0)),
                "pooled_residual_df_width_1": int(pooled_df[i]),
                "pooled_residual_df_width_2": int(pooled_df[j]),
                "trials_width_1": int(position_trials[:, i, hi].sum()),
                "trials_width_2": int(position_trials[:, j, hi].sum()),
            })

    return {
        "source_file": Path(path).name,
        "n_rows": int(n_rows),
        "n_subjects": int(len(subject_ids)),
        "anchor_number": int(anchor_number),
        "anchors": anchors,
        "per_numerosity_cluster_bootstrap": {
            "method": "within-participant × width × numerosity residual sums of squares divided by pooled residual degrees of freedom; paired participant-cluster bootstrap stratified by recorded block order at fixed numerosity",
            "shared_numerosities": list(shared_numbers),
            "widths": list(widths),
            "n_clusters": int(len(subject_ids)),
            "sequence_counts": order_counts,
            "observed_block_orders": {
                order: list(observed_width_order[order]) for order in order_labels
            },
            "familywise_alpha": float(familywise_alpha),
            "planned_family_size": int(planned_family_size),
            "adjustment": "Bonferroni over every configured shared numerosity × pairwise width contrast",
            "bootstrap_test": "centered, two-sided paired participant-cluster bootstrap of log variance ratios, resampling within observed block-order groups",
            "familywise_interval_method": "Bonferroni interval obtained by inversion of the plus-one absolute centered-bootstrap test on the log-ratio scale",
            "bootstrap_seed": int(bootstrap_seed),
            "bootstrap_reps_per_contrast": int(bootstrap_reps),
            "familywise_interval_level_percent": float(100 * (1 - familywise_alpha / planned_family_size)),
            "per_numerosity": per_numerosity,
            "contrasts": contrasts,
            "pairwise_summary": summaries,
        },
        "participant_equal_weight_sensitivity": {
            "estimand": "geometric mean of within-participant residual-variance ratios, with each participant equally weighted after pooling residual SSE and df across the configured common numerosities",
            "common_numerosities": list(shared_numbers),
            "complete_participants": int(np.count_nonzero(complete)),
            "sequence_counts": {order: int(len(idx)) for order, idx in zip(order_labels, complete_strata)},
            "familywise_alpha": float(familywise_alpha),
            "family_size": int(len(pairs)),
            "bootstrap_reps": int(equal_weight_bootstrap_reps),
            "contrasts": equal_weight_rows,
        },
        "order_sequence_sensitivity": {
            "estimand": "sequence-specific geometric mean of within-participant variance ratios after pooling residual sums of squares and degrees of freedom over the configured common numerosity support",
            "common_numerosities": list(shared_numbers),
            "sequence_counts": order_counts,
            "sequence_group_intervals_family_size": int(len(order_labels) * len(pairs)),
            "between_sequence_intervals_family_size": int(len(pairs)),
            "bootstrap_reps": int(equal_weight_bootstrap_reps),
            "interval_note": "Group-specific intervals are Bonferroni-adjusted across sequence × width-pair estimates; difference intervals are adjusted across width pairs. They do not identify causal order effects.",
            "contrasts": sequence_contrasts,
        },
        "within_block_position_sensitivity": {
            "estimand": "pooled within-participant × width × numerosity × block-half residual-variance ratios over the common numerosity support",
            "block_halves": {"first_half": "tLoc 1–60", "second_half": "tLoc 61–120"},
            "common_numerosities": list(shared_numbers),
            "sequence_counts": order_counts,
            "family_size": int(len(half_labels) * len(pairs)),
            "familywise_alpha": float(familywise_alpha),
            "bootstrap_reps": int(position_bootstrap_reps),
            "interval_note": "Participant-cluster bootstrap stratified by recorded block order; Bonferroni correction spans three width pairs × two block halves. This is a descriptive within-block sensitivity and does not identify block-period or carry-over effects.",
            "contrasts": position_contrasts,
        },
    }
```
