## Supplementary material for "The marginal brain": Replication package: figures.py

```
"""Figures: the two signatures, the price-margin relation, and power."""
import numpy as np
import matplotlib
matplotlib.use('Agg')
import matplotlib.pyplot as plt
from . import model as Mo

plt.rcParams.update({'font.size': 8, 'axes.spines.top': False, 'axes.spines.right': False,
                     'savefig.dpi': 300, 'font.family': 'DejaVu Serif',
                     'mathtext.fontset': 'stix', 'axes.labelsize': 8,
                     'xtick.labelsize': 7, 'ytick.labelsize': 7})


def fig_signatures(res, path):
    fig, ax = plt.subplots(1, 3, figsize=(10.0, 5.1))
    # (a) common factor across tasks and prior widths
    ws = np.linspace(20, 80, 50)
    for a, name, col in [(1, 'estimation', '#17365D'), (2, 'discrimination', '#C65A46')]:
        base = Mo.uniform_sigma(ws, 1.0, 1.0, a)
        hi = Mo.uniform_sigma(ws, 1.0, 2.0, a)
        ref = Mo.uniform_sigma(20.0, 1.0, 1.0, a)
        ax[0].loglog(ws, base / ref, color=col, lw=1.5, label=f'{name}, baseline')
        ax[0].loglog(ws, hi / ref, color=col, lw=1.5, ls='--', label=f'{name}, λ ×2')
    from matplotlib.ticker import FixedLocator, NullLocator, FormatStrFormatter
    ax[0].xaxis.set_major_locator(FixedLocator([20, 30, 40, 60, 80]))
    ax[0].xaxis.set_minor_locator(NullLocator())
    ax[0].xaxis.set_major_formatter(FormatStrFormatter('%g'))
    ax[0].yaxis.set_major_locator(FixedLocator([1.0, 1.5, 2.0, 3.0]))
    ax[0].yaxis.set_minor_locator(NullLocator())
    ax[0].yaxis.set_major_formatter(FormatStrFormatter('%g'))
    ax[0].set_xlabel('prior width w')
    ax[0].set_ylabel('imprecision (relative to w = 20)')
    ax[0].set_title('A  Common-factor scaling', loc='left', fontsize=8.5, weight='bold', pad=6)
    ax[0].legend(fontsize=6.1, frameon=False, loc='upper center', bbox_to_anchor=(0.5, -0.30),
                 ncol=2, handlelength=1.7, columnspacing=0.8)
    # (b) sign test in (dlog E, dlog sigma)
    t = np.linspace(-0.45, 0.45, 2)
    ax[1].plot(t, -0.5 * t, color='#17365D', lw=1.5, label='budget-price path (−½)')
    ax[1].plot(t, 0.5 * t, color='#C65A46', lw=1.5, label='technology path (+½)')
    cases = res['signatures']['cases']
    marks = {'price_rise_supply_tightening (mu x2)': ('o', 'p ×2'),
             'capacity_loss (K x0.8)': ('s', 'capacity ×0.8'),
             'cheaper_signals (e x0.71)': ('^', 'energy/signal ×0.71'),
             'scarcity_with_technology_change (mu x2, e x0.71, K x0.8)': ('D', 'p ×2 + tech'),
             'stakes_up (beta x1.5, price-like)': ('v', 'stakes ×1.5')}
    for k, (m, lab) in marks.items():
        c = cases[k]
        ax[1].plot(c['dlog_E'], c['dlog_sigma'], m, color='#2A7F7F', mec='#17365D', mew=0.4,
                   ms=4.2, label=lab)
    ax[1].axhline(0, color='0.7', lw=0.5)
    ax[1].axvline(0, color='0.7', lw=0.5)
    ax[1].set_xlabel('change in log task energy')
    ax[1].set_ylabel('change in log imprecision')
    ax[1].set_title('B  Joint-measurement sign test', loc='left', fontsize=8.5, weight='bold', pad=6)
    ax[1].legend(fontsize=5.8, frameon=False, loc='upper center', bbox_to_anchor=(0.5, -0.30),
                 ncol=2, handlelength=1.5, columnspacing=0.7)
    # (c) price against the reallocation margin
    M = np.array(res['budget']['M_grid'])
    for rho, col in [(0.5, '#2A7F7F'), (1.0, '#17365D'), (2.0, '#C65A46')]:
        ax[2].loglog(M, res['budget']['curves'][f'rho{rho}_c100.0'], color=col, lw=1.5, label=f'regulated, ρ={rho:g}')
    ax[2].loglog(M, res['budget']['curves']['rho1.0_chard'], color='#17365D', lw=1, ls=':', label='hard cap, ρ=1')
    ax[2].set_xlabel('reallocation margin M (set point = 1)')
    ax[2].set_ylabel('shadow price of energy')
    ax[2].set_title('C  Shadow price and margin', loc='left', fontsize=8.5, weight='bold', pad=6)
    ax[2].legend(fontsize=5.9, frameon=False, loc='upper center', bbox_to_anchor=(0.5, -0.30),
                 ncol=2, handlelength=1.5, columnspacing=0.7)
    for axis in ax:
        axis.tick_params(axis='both', labelsize=6.5, length=2.5)
    fig.subplots_adjust(left=0.07, right=0.99, top=0.88, bottom=0.38, wspace=0.46)
    fig.savefig(path + '.png', bbox_inches='tight')
    fig.savefig(path + '.pdf', bbox_inches='tight')
    plt.close(fig)


def fig_power(res, path):
    # The journal-width figure is typeset near single-column width. Use a
    # compact canvas and larger labels so text remains legible after inclusion.
    fig, ax = plt.subplots(1, 1, figsize=(5.8, 3.8))
    pw = res['power']
    Ns = pw['N_grid']
    colors = {1.0: '#7B8790', 1.25: '#17365D', 1.5: '#2A7F7F', 2.0: '#C65A46'}
    for row in pw['by_price_ratio']:
        pr = row['price_ratio']
        y = [row[f'power_N{N}'] for N in Ns]
        ci = [row['mc_uncertainty'][f'power_N{N}']['wilson_95'] for N in Ns]
        yerr = np.array([[v - lo for v, (lo, hi) in zip(y, ci)],
                         [hi - v for v, (lo, hi) in zip(y, ci)]])
        factor = pr ** 0.25
        lab = r'$p_1/p_0=1$ (no change)' if pr == 1.0 else rf'$p_1/p_0={pr:g}$ ($f={factor:.3f}$)'
        ax.errorbar(Ns, y, yerr=yerr, fmt='o-' if pr != 1.0 else 'o--', ms=3.2, lw=1.25,
                    capsize=2, label=lab, color=colors[pr])
    for row in res['estimation_recovery']:
        if row['price_ratio'] != 1.0:
            lo, hi = row['mc_uncertainty']['power']['wilson_95']
            ax.errorbar([row['N']], [row['power']],
                        yerr=[[row['power'] - lo], [hi - row['power']]],
                        fmt='*', color='#B08D57', mec='#775f34', mew=0.5, ms=8, capsize=2,
                        label=rf"estimation, $p_1/p_0={row['price_ratio']:g}$")
    ax.axhline(0.05, color='0.55', lw=0.65, ls='--')
    ax.axhline(0.8, color='0.55', lw=0.65, ls=':')
    ax.set_xlabel('participants (two-state within-subject design)')
    ax.set_ylabel('rejection rate (one-sided, alpha 0.05)')
    ax.xaxis.label.set_size(9.2)
    ax.yaxis.label.set_size(9.2)
    ax.tick_params(axis='both', labelsize=8.2, length=3)
    ax.set_ylim(0, 1.08)
    ax.set_yticks([0, 0.25, 0.50, 0.75, 1.00])
    ax.set_xticks(Ns)
    ax.legend(fontsize=8.2, frameon=False, loc='upper center', bbox_to_anchor=(0.5, -0.20),
              ncol=3, handlelength=1.5, columnspacing=0.8, labelspacing=0.5)
    fig.subplots_adjust(left=0.14, right=0.98, top=0.95, bottom=0.30)
    fig.savefig(path + '.png', bbox_inches='tight')
    fig.savefig(path + '.pdf', bbox_inches='tight')
    plt.close(fig)
```
