## Supplementary material for "The marginal brain": Replication package: model.py

```
"""Numerical closed forms: endogenous precision (Prat-Carrabin & Woodford) under a regulated energy budget."""
import numpy as np
from scipy.optimize import brentq
import math


def competing_use_value(Eo, v, rho):
    """Isoelastic value of competing energy use, including the log limit rho=1."""
    if Eo <= 0 or v <= 0 or rho <= 0:
        raise ValueError("Eo, v and rho must be positive")
    if rho == 1:
        return v * math.log(Eo)
    return v * math.expm1((1.0 - rho) * math.log(Eo)) / (1.0 - rho)


def marginal_competing_use_value(Eo, v, rho):
    """Marginal value V'(Eo)=v Eo^(-rho), including rho=1."""
    if Eo <= 0 or v <= 0 or rho <= 0:
        raise ValueError("Eo, v and rho must be positive")
    return v * Eo ** (-rho)


def Z_a(pdf, dx, a):
    """Z_a = int pi(x)^(a/3) dx on a grid."""
    return float(np.sum(pdf**(a / 3.0)) * dx)


def n_star(Z, K, lam, kappa=1.0):
    """Optimal number of signals for cost lam * n^kappa (kappa = 1 is the baseline)."""
    return (Z**3 / (kappa * lam * K**2))**(1.0 / (kappa + 1.0))


def sigma_x(Z, K, lam, pdf_x, a, kappa=1.0):
    """Imprecision I(x)^(-1/2) at the optimum."""
    n = n_star(Z, K, lam, kappa)
    I = n * (K * pdf_x**(a / 3.0) / Z)**2
    return 1.0 / np.sqrt(I)


def uniform_sigma(w, K, lam, a):
    pdf = 1.0 / w
    Z = w * pdf**(a / 3.0)
    return sigma_x(Z, K, lam, pdf, a)


def task_energy(Z, K, e, mu, kappa=1.0):
    return e * n_star(Z, K, e * mu, kappa)


def budget_F(mu, M, c, v, rho, e, A, k=1.0, hard_cap=False):
    """Equilibrium condition of the regulated set point.
    Regulatory marginal cost R'(d) = c * d^k above the set point, so mu = c d^k and d = (mu/c)^(1/k).
    F(mu) = d(mu) + M - E_task(mu) - E_other(mu); E_task = sqrt(e) A / sqrt(mu); E_other = (v/mu)^(1/rho)."""
    Et = np.sqrt(e) * A / np.sqrt(mu)
    Eo = (v / mu)**(1.0 / rho)
    d = 0.0 if hard_cap else (mu / c)**(1.0 / k)
    return d + M - Et - Eo


def budget_mu(M, c, v, rho, e, A, k=1.0, hard_cap=False):
    """Shadow price of energy at the regulated set point (or at the hard cap)."""
    if hard_cap and M <= 0:
        raise ValueError('hard cap needs a positive margin')
    lo, hi = 1e-12, 1.0
    while budget_F(hi, M, c, v, rho, e, A, k, hard_cap) < 0:
        hi *= 2.0
        if hi > 1e12:
            raise RuntimeError('no bracket')
    while budget_F(lo, M, c, v, rho, e, A, k, hard_cap) > 0:
        lo /= 10.0
    return brentq(lambda m: budget_F(m, M, c, v, rho, e, A, k, hard_cap), lo, hi, xtol=1e-15, rtol=1e-13, maxiter=500)


def budget_state(M, c, v, rho, e, A, k=1.0, hard_cap=False):
    mu = budget_mu(M, c, v, rho, e, A, k, hard_cap)
    Et = np.sqrt(e) * A / np.sqrt(mu)
    Eo = (v / mu)**(1.0 / rho)
    d = 0.0 if hard_cap else (mu / c)**(1.0 / k)
    total_above_committed = Et + Eo           # = M + d
    return dict(mu=mu, E_task=Et, E_other=Eo, excess=d, task_share_of_margin_use=Et / total_above_committed)
```
