## Supplementary material for "The marginal brain": Replication package: numerics.py

```
"""Model-independent numerical checks: direct optimization of the encoding for arbitrary priors and constraints."""
import numpy as np
from scipy.optimize import minimize


BIMODAL = dict(means=(30.0, 70.0), sds=(6.0, 8.0), weights=(1.0, 0.6))


def prior_on_grid(kind, x):
    if kind == 'uniform':
        pdf = np.ones_like(x)
    elif kind == 'loguniform':
        pdf = 1.0 / x
    elif kind == 'bimodal':
        (m1, m2), (s1, s2), (w1, w2) = BIMODAL['means'], BIMODAL['sds'], BIMODAL['weights']
        pdf = w1 * np.exp(-0.5 * ((x - m1) / s1)**2) + w2 * np.exp(-0.5 * ((x - m2) / s2)**2)
    else:
        raise ValueError(kind)
    dx = x[1] - x[0]
    return pdf / (pdf.sum() * dx)


def optimize_encoding(pdf, x, a, K, lam, gamma=1.0, kappa=1.0):
    """Minimize int pi^a/(n I1) dx + lam n^kappa subject to int I1^(gamma/2) dx = K (gamma = 1: Prat-Carrabin & Woodford).
    Parametrization: I1^(gamma/2) = K softmax(theta)/dx, n = exp(eta). Analytic gradient."""
    dx = x[1] - x[0]
    m = len(x)
    pa = pdf**a

    def unpack(par):
        th, eta = par[:m], par[m]
        wts = np.exp(th - th.max())
        wts /= wts.sum()
        u = K * wts / dx
        t = u**(2.0 / gamma)
        return wts, t, np.exp(eta)

    def obj(par):
        wts, t, n = unpack(par)
        Lterm = np.sum(pa / (n * t)) * dx
        val = Lterm + lam * n**kappa
        dobj_dt = -pa * dx / (n * t**2)
        q = dobj_dt * (2.0 / gamma) * t          # d obj / d log w_i
        g_th = q - wts * q.sum()
        g_eta = -Lterm + kappa * lam * n**kappa
        return val, np.concatenate([g_th, [g_eta]])

    par0 = np.concatenate([np.zeros(m), [np.log(10.0)]])
    res = minimize(obj, par0, jac=True, method='L-BFGS-B', options=dict(maxiter=20000, ftol=1e-15, gtol=1e-12))
    wts, t, n = unpack(res.x)
    sigma = 1.0 / np.sqrt(n * t)
    return dict(sigma=sigma, n=n, t=t, success=bool(res.success), fun=float(res.fun))


def integer_n_deviation(G, lam_ratio=2.0, scales=(0.3, 1, 3, 10, 30, 100)):
    """With loss G/n and cost lam n, compare the integer optimum with the continuous one.
    Returns, for each scale of n*, the realized sigma ratio between prices lam and lam*lam_ratio
    against the continuous prediction lam_ratio^(1/4)."""
    out = []
    for sc in scales:
        lam0 = G / sc**2               # continuous n* = sqrt(G/lam0) = sc
        ratios = []
        for lam in (lam0, lam0 * lam_ratio):
            nc = np.sqrt(G / lam)
            cands = np.unique(np.maximum(1, [np.floor(nc), np.ceil(nc)]))
            vals = G / cands + lam * cands
            nb = cands[np.argmin(vals)]
            ratios.append(1.0 / np.sqrt(nb))
        out.append(dict(n_star_continuous=float(sc), sigma_ratio_integer=float(ratios[1] / ratios[0]),
                        sigma_ratio_continuous=float(lam_ratio**0.25)))
    return out
```
