## Supplementary material for "The marginal brain": Replication package: recovery.py

```
"""Stage 4 (empirical anchoring): parameter recovery and power for the state factor f = (mu1/mu0)^(1/4)."""
import hashlib
import json
from pathlib import Path

import numpy as np
import scipy
from scipy.stats import ttest_1samp, wilcoxon
from scipy.special import expit, logit
from . import behavior as B


def _rate_summary(events):
    """Monte Carlo standard error and two-sided 95% Wilson interval for a rate."""
    events = np.asarray(events, dtype=bool)
    n = int(events.size)
    if n == 0:
        raise ValueError('Cannot summarize an empty set of Monte Carlo events')
    k = int(events.sum())
    p = k / n
    z = 1.959963984540054
    den = 1 + z*z/n
    center = (p + z*z/(2*n)) / den
    half = z * np.sqrt(p*(1-p)/n + z*z/(4*n*n)) / den
    return {'replications': n, 'events': k, 'estimate': float(p),
            'mc_se': float(np.sqrt(p*(1-p)/n)),
            'wilson_95': [float(max(0.0, center-half)), float(min(1.0, center+half))]}


def estimation_cache_fingerprint(cal, cfg, price_ratio, seed, per_chunk):
    """Fingerprint inputs and model source so stale chunks cannot be silently reused."""
    payload = {'cache_schema': 2, 'calibration': cal, 'config': cfg,
               'price_ratio': float(price_ratio), 'seed': int(seed),
               'numpy': np.__version__, 'scipy': scipy.__version__,
               'per_chunk': int(per_chunk)}
    digest = hashlib.sha256(json.dumps(payload, sort_keys=True, separators=(',', ':')).encode())
    for source in (Path(__file__), Path(B.__file__)):
        digest.update(source.read_bytes())
    return digest.hexdigest()[:20]


def _subject_params(rng, base_nu, sd_log_nu, f, sd_log_f):
    return base_nu * np.exp(sd_log_nu * rng.standard_normal()), f * np.exp(sd_log_f * rng.standard_normal())


def discrimination_power(rng, cal, cfg, price_ratio):
    """Within-subject design: baseline state and manipulated state, discrimination task.
    Returns per-rep arrays of fitted log f for the largest N; smaller N use nested subsets."""
    lo, hi, T = cfg['lo'], cfg['hi'], cfg['trials_per_state']
    w = hi - lo
    alpha = 0.75
    f_true = price_ratio**0.25
    Nmax = max(cfg['N_grid'])
    est = np.zeros((cfg['reps'], Nmax))
    fit_success = np.zeros((cfg['reps'], Nmax), dtype=bool)
    fit_boundary = np.zeros((cfg['reps'], Nmax), dtype=bool)
    for r in range(cfg['reps']):
        for i in range(Nmax):
            nu_i, f_i = _subject_params(rng, cal['nu'], cfg['sd_log_nu'], f_true, cfg['sd_log_f'])
            eta_i = expit(logit(cal['eta']) + cfg['sd_logit_eta'] * rng.standard_normal())
            data = []
            for state in (0, 1):
                s = nu_i * w**alpha * (f_i if state == 1 else 1.0)
                d, ch = B.simulate_discrimination(rng, T, s, eta_i, lo, hi)
                data.append((d, ch, state))
            x0 = np.array([np.log(cal['nu']), logit(cal['eta']), 0.0])
            par, ok, diag = B.fit(B.nll_discrimination, x0, (data, w, alpha), jac=True)
            est[r, i] = par[2]
            fit_success[r, i] = ok
            fit_boundary[r, i] = any(diag['at_bound'])
    out = dict(price_ratio=price_ratio, f_true=f_true, log_f_true=float(np.log(f_true)),
               fit_diagnostics={'fits': int(fit_success.size),
                                'success_rate': float(fit_success.mean()),
                                'boundary_rate': float(fit_boundary.mean())})
    out['mc_uncertainty'] = {}
    for N in cfg['N_grid']:
        sub = est[:, :N]
        pvals = np.array([ttest_1samp(sub[r], 0.0, alternative='greater').pvalue for r in range(sub.shape[0])])
        pw = np.array([wilcoxon(sub[r], alternative='greater').pvalue for r in range(sub.shape[0])])
        out[f'power_N{N}'] = float(np.mean(pvals < cfg['alpha']))
        out[f'power_wilcoxon_N{N}'] = float(np.mean(pw < cfg['alpha']))
        out[f'mean_log_f_hat_N{N}'] = float(sub.mean())
        out['mc_uncertainty'][f'power_N{N}'] = _rate_summary(pvals < cfg['alpha'])
        out['mc_uncertainty'][f'power_wilcoxon_N{N}'] = _rate_summary(pw < cfg['alpha'])
    out['bias_log_f'] = float(est.mean() - np.log(f_true))
    out['sd_subject_log_f_hat'] = float(est.std())
    return out


def estimation_recovery(rng, cal, cfg, price_ratio):
    lo, hi, T, N = cfg['lo'], cfg['hi'], cfg['trials_per_state'], cfg['N']
    alpha = 0.5
    f_true = price_ratio**0.25
    all_logf, powers, powers_t = [], [], []
    fit_success, fit_boundary = [], []
    for r in range(cfg['reps']):
        logf = np.zeros(N)
        for i in range(N):
            nu_i, f_i = _subject_params(rng, cal['nu'], cfg['sd_log_nu'], f_true, cfg['sd_log_f'])
            s0_i = cal['sigma0'] * np.exp(cfg['sd_log_sigma0'] * rng.standard_normal())
            data = []
            for state in (0, 1):
                xs = rng.integers(lo, hi + 1, size=T).astype(float)
                s = nu_i * (hi - lo)**alpha * (f_i if state == 1 else 1.0)
                resp = B.simulate_estimation(rng, xs, s, s0_i, lo, hi)
                data.append((xs, resp, state))
            x0 = np.log([cal['nu'], cal['sigma0'], 1.0])
            par, ok, diag = B.fit(B.nll_estimation, x0, (data, lo, hi, alpha))
            logf[i] = par[2]
            fit_success.append(ok)
            fit_boundary.append(any(diag['at_bound']))
        all_logf.append(logf)
        powers.append(wilcoxon(logf, alternative='greater').pvalue < cfg['alpha'])
        powers_t.append(ttest_1samp(logf, 0.0, alternative='greater').pvalue < cfg['alpha'])
    all_logf = np.concatenate(all_logf)
    out = dict(price_ratio=price_ratio, f_true=f_true, N=N, reps=cfg['reps'],
                power=float(np.mean(powers)), power_ttest=float(np.mean(powers_t)),
                bias_log_f_mean=float(all_logf.mean() - np.log(f_true)),
                bias_log_f_median=float(np.median(all_logf) - np.log(f_true)),
                sd_subject_log_f_hat=float(all_logf.std()),
                fit_diagnostics={'fits': len(fit_success),
                                 'success_rate': float(np.mean(fit_success)),
                                 'boundary_rate': float(np.mean(fit_boundary))},
                mean_log_f_hat=float(all_logf.mean()))
    out['mc_uncertainty'] = {'power': _rate_summary(powers),
                             'power_ttest': _rate_summary(powers_t)}
    return out


# ---------------- chunked estimation recovery (independent seeded streams per chunk) ----------------
def estimation_chunk(cal, cfg, price_ratio, chunk_id, reps_in_chunk, base_seed):
    """One chunk of replications with its own stream, so that results do not depend on how chunks are scheduled."""
    rng = np.random.default_rng([base_seed, int(round(price_ratio * 1000)), chunk_id])
    lo, hi, T, N = cfg['lo'], cfg['hi'], cfg['trials_per_state'], cfg['N']
    f_true = price_ratio**0.25
    reps, successes, boundaries = [], [], []
    for r in range(reps_in_chunk):
        logf = np.zeros(N)
        for i in range(N):
            nu_i, f_i = _subject_params(rng, cal['nu'], cfg['sd_log_nu'], f_true, cfg['sd_log_f'])
            s0_i = cal['sigma0'] * np.exp(cfg['sd_log_sigma0'] * rng.standard_normal())
            data = []
            for state in (0, 1):
                xs = rng.integers(lo, hi + 1, size=T).astype(float)
                s = nu_i * (hi - lo)**0.5 * (f_i if state == 1 else 1.0)
                resp = B.simulate_estimation(rng, xs, s, s0_i, lo, hi)
                data.append((xs, resp, state))
            par, ok, diag = B.fit(B.nll_estimation, np.log([cal['nu'], cal['sigma0'], 1.0]), (data, lo, hi, 0.5))
            logf[i] = par[2]
            successes.append(ok)
            boundaries.append(any(diag['at_bound']))
        reps.append(logf.tolist())
    return dict(reps=reps, fit_successes=int(sum(successes)),
                fit_count=len(successes), boundary_fits=int(sum(boundaries)))


def estimation_aggregate(chunks, price_ratio, cfg):
    reps = [np.array(r) for ch in chunks for r in ch['reps']]
    allf = np.concatenate(reps)
    f_true = price_ratio**0.25
    pw = np.mean([wilcoxon(r, alternative='greater').pvalue < cfg['alpha'] for r in reps])
    pt = np.mean([ttest_1samp(r, 0.0, alternative='greater').pvalue < cfg['alpha'] for r in reps])
    n_fits = sum(ch['fit_count'] for ch in chunks)
    out = dict(price_ratio=price_ratio, f_true=f_true, N=cfg['N'], reps=len(reps),
                power=float(pw), power_ttest=float(pt),
                bias_log_f_mean=float(allf.mean() - np.log(f_true)),
                bias_log_f_median=float(np.median(allf) - np.log(f_true)),
                sd_subject_log_f_hat=float(allf.std()),
                fit_diagnostics={'fits': int(n_fits),
                                 'success_rate': float(sum(ch['fit_successes'] for ch in chunks) / n_fits),
                                 'boundary_rate': float(sum(ch['boundary_fits'] for ch in chunks) / n_fits)},
                mean_log_f_hat=float(allf.mean()))
    rejections_w = [wilcoxon(r, alternative='greater').pvalue < cfg['alpha'] for r in reps]
    rejections_t = [ttest_1samp(r, 0.0, alternative='greater').pvalue < cfg['alpha'] for r in reps]
    out['mc_uncertainty'] = {'power': _rate_summary(rejections_w),
                             'power_ttest': _rate_summary(rejections_t)}
    return out


def discrimination_lapse_robustness(rng, cal, cfg, price_ratio, lapse_odds_mult, N_grid, reps):
    """State-dependent lapses (odds multiplied in the second state), fitted with a shared lapse rate and with
    state-specific lapse rates. Returns the rejection rates of the one-sided t-test on log f and the mean estimates."""
    lo, hi, T = cfg['lo'], cfg['hi'], cfg['trials_per_state']
    w, alpha, f_true = hi - lo, 0.75, price_ratio**0.25
    N = max(N_grid)
    est = {'shared': np.zeros((reps, N)), 'state': np.zeros((reps, N))}
    fit_ok = {'shared': [], 'state': []}
    fit_boundary = {'shared': [], 'state': []}
    for r in range(reps):
        for i in range(N):
            nu_i, f_i = _subject_params(rng, cal['nu'], cfg['sd_log_nu'], f_true, cfg['sd_log_f'])
            le = logit(cal['eta']) + cfg['sd_logit_eta'] * rng.standard_normal()
            data = []
            for state in (0, 1):
                s = nu_i * w**alpha * (f_i if state == 1 else 1.0)
                eta_s = expit(le + (np.log(lapse_odds_mult) if state == 1 else 0.0))
                d, ch = B.simulate_discrimination(rng, T, s, eta_s, lo, hi)
                data.append((d, ch, state))
            x0 = np.array([np.log(cal['nu']), logit(cal['eta']), 0.0])
            par, ok, diag = B.fit(B.nll_discrimination, x0, (data, w, alpha), jac=True)
            est['shared'][r, i] = par[2]
            fit_ok['shared'].append(ok)
            fit_boundary['shared'].append(any(diag['at_bound']))
            x1 = np.array([np.log(cal['nu']), logit(cal['eta']), 0.0, logit(cal['eta'])])
            par, ok, diag = B.fit(B.nll_discrimination_state_lapse, x1, (data, w, alpha), jac=True)
            est['state'][r, i] = par[2]
            fit_ok['state'].append(ok)
            fit_boundary['state'].append(any(diag['at_bound']))
    out = dict(price_ratio=price_ratio, lapse_odds_mult=lapse_odds_mult, N_grid=list(N_grid), reps=reps)
    out['fit_diagnostics'] = {
        model: {'fits': len(fit_ok[model]), 'success_rate': float(np.mean(fit_ok[model])),
                'boundary_rate': float(np.mean(fit_boundary[model]))}
        for model in ('shared', 'state')}
    out['mc_uncertainty'] = {}
    for k, e in est.items():
        for n_ in N_grid:
            p = np.array([ttest_1samp(e[r, :n_], 0.0, alternative='greater').pvalue for r in range(reps)])
            out[f'rejection_{k}_N{n_}'] = float(np.mean(p < cfg['alpha']))
            out['mc_uncertainty'][f'rejection_{k}_N{n_}'] = _rate_summary(p < cfg['alpha'])
        out[f'bias_log_f_{k}'] = float(e.mean() - np.log(f_true))
    return out
```
