## Supplementary material for "The marginal brain": Replication package: validation.py

```
"""Stage 4: sensitivity, robustness, ablations, rivals, and the constructive check for Proposition 5."""
import hashlib
from pathlib import Path
import numpy as np
from . import model as Mo, numerics as Nu


def _el_mu_M(M, c, v, rho, e, A, k=1.0, hard=False, h=1e-5):
    m1 = Mo.budget_mu(M * (1 + h), c, v, rho, e, A, k, hard)
    m0 = Mo.budget_mu(M * (1 - h), c, v, rho, e, A, k, hard)
    return (np.log(m1) - np.log(m0)) / (np.log(1 + h) - np.log(1 - h))


def budget_analysis(bcfg):
    b = bcfg['baseline']
    Ms = np.linspace(bcfg['M_min'], bcfg['M_max'], bcfg['M_points'])
    curves, elasticities = {}, {}
    for rho in bcfg['rho_values']:
        for c in bcfg['c_values'] + ['hard']:
            hard = c == 'hard'
            cc = 1.0 if hard else c
            curves[f'rho{rho}_c{c}'] = [Mo.budget_mu(M, cc, b['v'], rho, b['e'], b['A'], b['k'], hard) for M in Ms]
            elasticities[f'rho{rho}_c{c}'] = float(_el_mu_M(b['M'], cc, b['v'], rho, b['e'], b['A'], b['k'], hard))
    monotone = bool(all(np.all(np.diff(v) < 0) for v in curves.values()))
    # regulatory-cost exponent k
    mono_k = {}
    for k in bcfg['k_values']:
        mus = [Mo.budget_mu(M, b['c'], b['v'], b['rho'], b['e'], b['A'], k) for M in Ms]
        mono_k[f'k{k}'] = bool(np.all(np.diff(mus) < 0))
    st = Mo.budget_state(b['M'], b['c'], b['v'], b['rho'], b['e'], b['A'], b['k'])
    # price-taker: elasticity of mu with respect to the task's scale A, numerical and formula
    h = 1e-5
    m1 = Mo.budget_mu(b['M'], b['c'], b['v'], b['rho'], b['e'], b['A'] * (1 + h), b['k'])
    m0 = Mo.budget_mu(b['M'], b['c'], b['v'], b['rho'], b['e'], b['A'] * (1 - h), b['k'])
    el_A = (np.log(m1) - np.log(m0)) / (np.log(1 + h) - np.log(1 - h))
    mu = st['mu']
    Fmu = 1.0 / b['c'] + st['E_task'] / (2 * mu) + st['E_other'] / (b['rho'] * mu)
    el_A_formula = (st['E_task'] / mu) / Fmu
    # a 30 percent tightening of the margin at baseline
    mu_t = Mo.budget_mu(0.7 * b['M'], b['c'], b['v'], b['rho'], b['e'], b['A'], b['k'])
    return dict(M_grid=Ms.tolist(), curves=curves, monotone_all=monotone, monotone_by_k=mono_k,
                el_mu_M=elasticities, el_sigma_M_baseline=float(elasticities[f"rho{b['rho']}_c{b['c']}"] / 4.0),
                baseline_mu=float(mu), baseline_task_energy=float(st['E_task']), baseline_other_energy=float(st['E_other']),
                baseline_task_share_of_margin_use=float(st['task_share_of_margin_use']),
                el_mu_taskscale_numeric=float(el_A), el_mu_taskscale_formula=float(el_A_formula),
                tighten30_mu_ratio=float(mu_t / mu), tighten30_sigma_ratio=float((mu_t / mu)**0.25),
                tighten30_task_energy_ratio=float((mu_t / mu)**-0.5))


def constructive_prop5(s_values=(0.001, 0.01, 0.05, 0.2), mu_values=(0.1, 1.0, 10.0, 100.0), rho=1.0):
    """For any target task share s of use above committed expenditure and any price mu, construct an equilibrium."""
    max_err_mu, max_err_s = 0.0, 0.0
    for s in s_values:
        for mu in mu_values:
            e = 1.0
            A = s * np.sqrt(mu)                 # E_task = A / sqrt(mu) = s
            Eo = 1 - s                           # normalize use above commitment: E_task + E_other = 1
            v = mu * Eo**rho                    # V'(Eo) = v Eo^(-rho) = mu
            d = 0.01                              # positive regulatory excess at the target equilibrium
            c = mu / d                           # k = 1, so d = mu / c
            Bbar = 1.0
            C = d                                 # ensures M = Bbar - C = 1 - d
            M = Bbar - C
            # Use an independent monotone bisection on the equilibrium residual
            # rather than validating the construction through the same solver
            # used to generate the model's other numerical results.
            lo, hi = 1e-12, max(1.0, mu)
            while Mo.budget_F(hi, M, c, v, rho, e, A) < 0:
                hi *= 2.0
            while Mo.budget_F(lo, M, c, v, rho, e, A) > 0:
                lo *= 0.1
            for _ in range(160):
                mid = (lo + hi) / 2.0
                if Mo.budget_F(mid, M, c, v, rho, e, A) < 0:
                    lo = mid
                else:
                    hi = mid
            mu_hat = (lo + hi) / 2.0
            E_hat = np.sqrt(e) * A / np.sqrt(mu_hat)
            Eo_hat = (v / mu_hat)**(1.0 / rho)
            s_hat = E_hat / (E_hat + Eo_hat)
            max_err_mu = max(max_err_mu, abs(mu_hat / mu - 1))
            max_err_s = max(max_err_s, abs(s_hat / s - 1))
    return dict(max_rel_err_mu=float(max_err_mu), max_rel_err_share=float(max_err_s),
                n_pairs=len(s_values) * len(mu_values),
                validation_method="Independent monotone bisection of the equilibrium residual; the share uses task energy divided by task plus competing energy, excluding committed use.",
                validation_source_sha256=hashlib.sha256(Path(__file__).read_bytes()).hexdigest())


def robustness_numeric(ncfg):
    """Model-independent checks with the direct optimizer: priors, constraint exponent gamma, cost exponent kappa."""
    x = np.linspace(ncfg['x_min'], ncfg['x_max'], ncfg['grid_points'])
    dx = x[1] - x[0]
    lam0, lam1, K = ncfg['lam0'], ncfg['lam1'], ncfg['K']
    out = {'priors': {}, 'gammas': {}, 'kappas': {}}
    worst_cf, worst_cfac = 0.0, 0.0
    for kind in ncfg['priors']:
        pdf = Nu.prior_on_grid(kind, x)
        for a in (1, 2):
            r0 = Nu.optimize_encoding(pdf, x, a, K, lam0)
            r1 = Nu.optimize_encoding(pdf, x, a, K, lam1)
            cf = Mo.sigma_x(Mo.Z_a(pdf, dx, a), K, lam0, pdf, a)
            e_cf = float(np.max(np.abs(r0['sigma'] / cf - 1)))
            ratio = r1['sigma'] / r0['sigma']
            e_fac = float(np.max(np.abs(ratio / (lam1 / lam0)**0.25 - 1)))
            out['priors'][f'{kind}_a{a}'] = dict(max_rel_err_closed_form=e_cf, max_rel_err_common_factor=e_fac)
            worst_cf, worst_cfac = max(worst_cf, e_cf), max(worst_cfac, e_fac)
    out['worst_closed_form_err'] = worst_cf
    out['worst_common_factor_err'] = worst_cfac
    # constraint exponent gamma: price elasticity of sigma stays 1/4; width exponent changes
    xu = lambda w: np.linspace(50 - w / 2, 50 + w / 2, ncfg['grid_points'])
    for g in ncfg['gammas']:
        res = {}
        for a in (1, 2):
            sig = {}
            for w in (20.0, 60.0):
                xx = xu(w)
                pdf = Nu.prior_on_grid('uniform', xx)
                for lam in (lam0, lam1):
                    sig[(w, lam)] = np.median(Nu.optimize_encoding(pdf, xx, a, K, lam, gamma=g)['sigma'])
            el_lam = np.log(sig[(20.0, lam1)] / sig[(20.0, lam0)]) / np.log(lam1 / lam0)
            el_w = np.log(sig[(60.0, lam0)] / sig[(20.0, lam0)]) / np.log(3.0)
            res[f'a{a}'] = dict(el_sigma_lambda=float(el_lam), el_sigma_w=float(el_w))
        out['gammas'][f'gamma{g}'] = res
    # cost exponent kappa: elasticity 1/(2(kappa+1)); slopes -1/2 (price) and kappa/2 (capacity)
    xx = xu(40.0)
    pdf = Nu.prior_on_grid('uniform', xx)
    for kap in ncfg['kappas']:
        r0 = Nu.optimize_encoding(pdf, xx, 1, K, lam0, kappa=kap)
        r1 = Nu.optimize_encoding(pdf, xx, 1, K, lam1, kappa=kap)
        rK = Nu.optimize_encoding(pdf, xx, 1, 0.8 * K, lam0, kappa=kap)
        s0, s1, sK = (np.median(r['sigma']) for r in (r0, r1, rK))
        el = np.log(s1 / s0) / np.log(lam1 / lam0)
        slope_price = np.log(s1 / s0) / np.log(r1['n'] / r0['n'])      # energy proportional to n at fixed e
        slope_cap = np.log(sK / s0) / np.log(rK['n'] / r0['n'])
        out['kappas'][f'kappa{kap}'] = dict(el_sigma_lambda=float(el), el_target=1 / (2 * (kap + 1)),
                                            slope_price=float(slope_price), slope_capacity=float(slope_cap),
                                            slope_capacity_target=kap / 2)
    out['integer_n'] = Nu.integer_n_deviation(1.0)
    return out


def signatures():
    """Predicted (dlog sigma, dlog E, slope) for manipulations and rival accounts (Proposition 6, Stage 4 rivals)."""
    def sig(x=0.0, y=0.0, z=0.0):
        ds = x / 4 + y / 4 - z / 2
        dE = -x / 2 + y / 2 - z
        return dict(dlog_sigma=ds, dlog_E=dE, slope=(ds / dE) if dE != 0 else None)
    cases = {
        'price_rise_supply_tightening (mu x2)': sig(x=np.log(2.0)),
        'price_rise_opportunity_cost (mu x2, via value of other uses)': sig(x=np.log(2.0)),
        'stakes_up (beta x1.5, price-like)': sig(x=-np.log(1.5)),
        'capacity_loss (K x0.8)': sig(z=np.log(0.8)),
        'cheaper_signals (e x0.71)': sig(y=np.log(0.71)),
        'scarcity_with_technology_change (mu x2, e x0.71, K x0.8)': sig(x=np.log(2.0), y=np.log(0.71), z=np.log(0.8)),
        'cheap_cognition (no metabolic effect)': dict(dlog_sigma=0.0, dlog_E=0.0, slope=None),
    }
    # General condition (Proposition 6): with technology index T = dlog e - 2 dlog K and price change x = dlog mu,
    # slope = (x + T) / (2 (T - x)), negative exactly when |T| < |x|. Checked on a random sample.
    rng = np.random.default_rng(7)
    xs, ys, zs = rng.normal(size=(3, 20000)) * 0.5
    ok = True
    for x, y, z in zip(xs, ys, zs):
        r = sig(x=x, y=y, z=z)
        if r['slope'] is None:
            continue
        T = y - 2 * z
        ok &= abs(r['slope'] - (x + T) / (2 * (T - x))) < 1e-9
        if abs(abs(T) - abs(x)) > 1e-9:
            ok &= (r['slope'] < 0) == (abs(T) < abs(x))
    return dict(cases={k: {kk: (float(vv) if vv is not None else None) for kk, vv in v.items()} for k, v in cases.items()},
                slope_formula_and_sign_condition_verified=bool(ok),
                padamsey_like_technology_index=float(np.log(0.71) - 2 * np.log(0.8)), price_index_mu_x2=float(np.log(2.0)))


def ablations(bcfg):
    b = bcfg['baseline']
    # (i) fixed price: no budget layer, so the margin cannot move precision
    fixed_price_ratio = 1.0
    # (ii) fixed number of signals (capacity-constrained models): sigma does not depend on the price; width exponent 1
    w1, w2 = 20.0, 60.0
    sig_fixed_n = lambda w, a: 1.0 / np.sqrt(10.0 * (1.0 / w)**2)   # n fixed at 10, I1 = (K/w)^2 with K = 1
    el_w_fixed_n = np.log(sig_fixed_n(w2, 1) / sig_fixed_n(w1, 1)) / np.log(w2 / w1)
    # (iii) curvature of the value of competing uses: elasticity of the price to the margin shrinks with rho
    rhos = [0.05, 0.25, 0.5, 1.0, 2.0]
    els = {f'rho{r}': float(_el_mu_M(b['M'], b['c'], b['v'], r, b['e'], b['A'], b['k'])) for r in rhos}
    return dict(fixed_price_sigma_ratio_under_margin_change=fixed_price_ratio,
                fixed_n_el_sigma_w=float(el_w_fixed_n), fixed_n_el_sigma_lambda=0.0,
                el_mu_M_by_rho=els)


# Budget comparative statics, measurement checks and numerical validation.
def net_increment_analysis(bcfg):
    """Lemma 3: gross task energy E against the net increase in total use when the task is switched on."""
    b = bcfg['baseline']

    def pair(M, c, v, rho=b['rho'], k=b['k']):
        m1 = Mo.budget_mu(M, c, v, rho, b['e'], b['A'], k)
        m0 = Mo.budget_mu(M, c, v, rho, b['e'], 0.0, k)
        d1, d0 = (m1 / c)**(1 / k), (m0 / c)**(1 / k)
        E = np.sqrt(b['e']) * b['A'] / np.sqrt(m1)
        E0 = np.sqrt(b['e']) * b['A'] / np.sqrt(m0)
        # The first-order expansion is around the no-task equilibrium (epsilon=0).
        # Its comparative-static coefficient must therefore use baseline d0 and Eo0,
        # not their post-task values at m1.
        Eo0 = (v / m0)**(1 / rho)
        phi_approx = (d0 / k) / (d0 / k + Eo0 / rho)
        net_dB = d1 - d0
        first_order_net_dB = phi_approx * E0
        return dict(mu=m1, mu_without_task=m0, gross_E=E, gross_E_no_task=E0,
                    net_dB=net_dB, net_over_gross=net_dB / E,
                    net_over_unit_gross=net_dB / E0, phi_approx=phi_approx,
                    first_order_net_dB=first_order_net_dB,
                    first_order_approx_rel_err=abs(first_order_net_dB / net_dB - 1))
    base = pair(b['M'], b['c'], b['v'])
    sweep = [dict(M=M, **pair(M, b['c'], b['v'])) for M in (0.08, 0.05, 0.035, 0.02)]
    # same price, different regulatory stiffness: adjust v to hold the price at its baseline value
    target = base['mu']
    same_price = []
    for c in (1.0, 10.0, 100.0, 1000.0, 1e5):
        lo, hi = -25.0, 8.0
        for _ in range(200):
            mid = 0.5 * (lo + hi)
            if Mo.budget_mu(b['M'], c, np.exp(mid), b['rho'], b['e'], b['A'], b['k']) > target:
                hi = mid
            else:
                lo = mid
        v = np.exp(0.5 * (lo + hi))
        r = pair(b['M'], c, v)
        same_price.append(dict(c=c, mu=r['mu'], net_over_gross=r['net_over_gross']))
    return dict(baseline=base,
                baseline_approx_rel_err=base['first_order_approx_rel_err'],
                margin_sweep=sweep,
                gross_falls_with_price=bool(all(np.diff([s['gross_E'] for s in sweep]) < 0)),
                net_rises_with_price=bool(all(np.diff([s['net_dB'] for s in sweep]) > 0)),
                same_price_varying_stiffness=same_price)


def integer_sweep(G=1.0, lam_ratio=2.0, lo=0.3, hi=30.0, n=3000):
    """Response of an integer number of signals to a doubled price, across the continuous optimum."""
    def best(lam):
        nc = np.sqrt(G / lam)
        cand = np.arange(max(1, int(np.floor(nc)) - 1), int(np.ceil(nc)) + 2)
        return cand[np.argmin(G / cand + lam * cand)]
    ncs = np.linspace(lo, hi, n)
    ratios = np.array([np.sqrt(best(G / nc**2) / best(lam_ratio * G / nc**2)) for nc in ncs])
    out = dict(max_nc_without_effect=float(ncs[ratios == 1.0].max()),
               switch_1_2=float(np.sqrt(2.0)), switch_2_3=float(np.sqrt(6.0)),
               continuous_ratio=float(lam_ratio**0.25))
    for a, b in ((1.5, 3.0), (3.0, 10.0), (10.0, 30.0)):
        sel = (ncs >= a) & (ncs < b)
        out[f'range_{a:g}_{b:g}'] = [float(ratios[sel].min()), float(ratios[sel].max())]
    return out


def displacement_remark(price_ratio=2.0):
    """Sign test when the measured increment is net of activity displaced within the region (E_meas = E - D)."""
    x = np.log(price_ratio)
    E0 = 1.0
    E1 = E0 * price_ratio**-0.5
    s_ratio = price_ratio**0.25
    rows = {}
    scenarios = (
        ('proportional (D = 0.6 E)', 0.6 * E0, 0.6 * E1),
        ('fixed (D = 0.6 E0 in both states)', 0.6 * E0, 0.6 * E0),
        ('displacement falls by half the energy decline', 0.6 * E0,
         0.6 * E0 + 0.5 * (E1 - E0)),
        # Choose a sufficiently large but feasible D0; the earlier D0=0.3
        # parameterization implied D1<0, which has no physical interpretation.
        ('displacement falls by 1.5 times the energy decline', 0.6 * E0,
         0.6 * E0 + 1.5 * (E1 - E0)),
    )
    for label, D0, D1 in scenarios:
        net0, net1 = E0 - D0, E1 - D1
        if min(D0, D1, net0, net1) < 0:
            raise ValueError(f'Infeasible displacement scenario: {label}')
        slope = np.log(s_ratio) / np.log(net1 / net0)
        rows[label] = {'slope': float(slope), 'D0': float(D0), 'D1': float(D1),
                       'net_energy0': float(net0), 'net_energy1': float(net1)}
    return rows


def calibration_discrete_prior(cal, cfg, nsim=200000, seed=11):
    """Monte Carlo SDs of estimates at numerosity 60 with the experiment's integer prior (the model uses a continuous one)."""
    from scipy.stats import norm
    rg = np.random.default_rng(seed)
    out = []
    for lo, hi in zip(cfg['lows'], cfg['highs']):
        s = cal['nu'] * (hi - lo)**cfg['alpha']
        r = cfg['x'] + s * rg.standard_normal(nsim)
        sup = np.arange(lo, hi + 1)
        post = np.empty(nsim)
        for i in range(0, nsim, 20000):
            lw = -0.5 * ((r[i:i + 20000, None] - sup[None, :]) / s)**2
            lw -= lw.max(axis=1, keepdims=True)
            wts = np.exp(lw)
            post[i:i + 20000] = (wts * sup).sum(1) / wts.sum(1)
        a = norm.cdf((lo - 0.5 - post) / cal['sigma0'])
        bb = norm.cdf((hi + 0.5 - post) / cal['sigma0'])
        u = a + (bb - a) * rg.random(nsim)
        resp = np.clip(np.round(post + cal['sigma0'] * norm.ppf(np.clip(u, 1e-12, 1 - 1e-12))), lo, hi)
        out.append(float(resp.std()))
    return out


def calibration_integer_prior_refit(cfg):
    """Re-fit the estimation calibration under the experiment's integer prior, by deterministic Gauss-Hermite quadrature."""
    from numpy.polynomial.hermite_e import hermegauss
    from scipy.stats import norm
    from scipy.optimize import least_squares
    gx, gw = hermegauss(60)
    gw = gw / gw.sum()

    def sd(nu, s0, lo, hi):
        s = nu * (hi - lo)**cfg['alpha']
        sup = np.arange(lo, hi + 1)
        r = cfg['x'] + s * gx
        lw = -0.5 * ((r[:, None] - sup[None, :]) / s)**2
        lw -= lw.max(1, keepdims=True)
        w = np.exp(lw)
        m = (w * sup).sum(1) / w.sum(1)
        P = norm.cdf((sup[None, :] + 0.5 - m[:, None]) / s0) - norm.cdf((sup[None, :] - 0.5 - m[:, None]) / s0)
        P /= (norm.cdf((hi + 0.5 - m) / s0) - norm.cdf((lo - 0.5 - m) / s0))[:, None]
        P = (gw[:, None] * P).sum(0)
        P /= P.sum()
        mu = (P * sup).sum()
        return float(np.sqrt((P * (sup - mu)**2).sum()))
    target = np.array(cfg['sd'])
    rngs = list(zip(cfg['lows'], cfg['highs']))
    sol = least_squares(lambda p: np.array([sd(*np.exp(p), lo, hi) for lo, hi in rngs]) - target, np.log([1.0, 3.0]))
    nu, s0 = np.exp(sol.x)
    return dict(nu=float(nu), sigma0=float(s0), fitted_sd=[sd(nu, s0, lo, hi) for lo, hi in rngs])
```
