## Supplementary material for "The marginal brain": Replication package: test_model.py

```
"""Invariant tests. Run with the package's documented Python environment."""
import os, sys, json, csv, subprocess, tempfile, re
import numpy as np
from scipy.optimize import check_grad
ROOT = os.path.dirname(os.path.dirname(os.path.abspath(__file__)))
sys.path.insert(0, ROOT)
from spt import analytics, model as Mo, numerics as Nu, validation as V, behavior as B, recovery as R, empirical as Emp
import complete_case_sensitivity as CCS
import participant_sign_sensitivity as PSS

CFG = json.load(open(os.path.join(ROOT, 'config.json')))


def test_symbolic_results():
    r = analytics.derive()
    flags = [k for k, v in r.items() if isinstance(v, bool)]
    assert all(r[k] for k in flags), [k for k in flags if not r[k]]
    assert r['el_sigma_lambda'] == '1/4' and r['el_n_lambda'] == '-1/2' and r['el_E_mu'] == '-1/2'
    assert r['uniform_estimation_exp_w'] == '1/2' and r['uniform_discrimination_exp_w'] == '3/4'
    assert r['slope_price'] == '-1/2' and r['slope_energy_per_signal'] == '1/2' and r['slope_capacity'] == '1/2'


def test_numeric_optimizer_matches_closed_form_and_common_factor():
    x = np.linspace(10, 90, 120)
    dx = x[1] - x[0]
    for kind in ('uniform', 'loguniform', 'bimodal'):
        pdf = Nu.prior_on_grid(kind, x)
        for a in (1, 2):
            r0 = Nu.optimize_encoding(pdf, x, a, 1.0, 1e-3)
            r1 = Nu.optimize_encoding(pdf, x, a, 1.0, 3e-3)
            cf = Mo.sigma_x(Mo.Z_a(pdf, dx, a), 1.0, 1e-3, pdf, a)
            assert np.max(np.abs(r0['sigma'] / cf - 1)) < 1e-5
            assert np.max(np.abs(r1['sigma'] / r0['sigma'] / 3**0.25 - 1)) < 1e-5


def test_budget_comparative_statics():
    b = CFG['budget']['baseline']
    mu = lambda **kw: Mo.budget_mu(kw.get('M', b['M']), kw.get('c', b['c']), kw.get('v', b['v']), b['rho'], b['e'], b['A'])
    assert mu(M=0.04) > mu(M=0.05) > mu(M=0.06)
    assert mu(v=0.06) > mu(v=0.05)
    assert mu(c=200.0) > mu(c=100.0)
    hard = Mo.budget_mu(b['M'], 1.0, b['v'], b['rho'], b['e'], b['A'], hard_cap=True)
    assert abs(Mo.budget_F(hard, b['M'], 1.0, b['v'], b['rho'], b['e'], b['A'], hard_cap=True)) < 1e-10


def test_isoelastic_value_log_limit():
    Eo, v = 0.7, 0.05
    assert np.isclose(Mo.competing_use_value(Eo, v, 1.0), v * np.log(Eo))
    assert np.isclose(Mo.marginal_competing_use_value(Eo, v, 1.0), v / Eo)
    for rho in (0.5, 2.0):
        h = 1e-6
        fd = (Mo.competing_use_value(Eo + h, v, rho)
              - Mo.competing_use_value(Eo - h, v, rho)) / (2 * h)
        assert np.isclose(fd, Mo.marginal_competing_use_value(Eo, v, rho), rtol=1e-8)
    for rho in (1 - 1e-7, 1 + 1e-7):
        assert np.isclose(Mo.competing_use_value(Eo, v, rho), v * np.log(Eo), atol=1e-8)


def test_proposition5_construction():
    r = V.constructive_prop5()
    assert r['max_rel_err_mu'] < 1e-10 and r['max_rel_err_share'] < 1e-10
    assert r['validation_method'].startswith('Independent monotone bisection')
    import hashlib
    with open(V.__file__, 'rb') as stream:
        assert r['validation_source_sha256'] == hashlib.sha256(stream.read()).hexdigest()


def test_sign_condition():
    assert V.signatures()['slope_formula_and_sign_condition_verified']


def test_calibration_and_gradient():
    ce = B.calibrate_estimation(CFG['calibration']['estimation'])
    assert max(abs(f - t) for f, t in zip(ce['fitted_sd'], ce['target_sd'])) < 0.2
    cd = B.calibrate_discrimination(CFG['calibration']['discrimination'])
    assert max(abs(r) for r in cd['residuals']) < 0.02
    rng = np.random.default_rng(0)
    data = [(*B.simulate_discrimination(rng, 200, 0.2 * 30**0.75 * (1.2 if st else 1.0), 0.14, 35, 65), st) for st in (0, 1)]
    f = lambda p: B.nll_discrimination(p, data, 30, 0.75)[0]
    g = lambda p: B.nll_discrimination(p, data, 30, 0.75)[1]
    assert check_grad(f, g, np.array([np.log(0.2), -1.8, 0.1])) < 1e-3
    par, ok, diag = B.fit(B.nll_discrimination,
                          np.array([np.log(0.2), -1.8, 0.0]), (data, 30, 0.75), jac=True)
    assert ok and diag['success'] and len(diag['at_bound']) == 3
    assert all(lo <= x <= hi for x, (lo, hi) in zip(par, B._FIT_BOUNDS[B.nll_discrimination]))


def test_monte_carlo_uncertainty_includes_boundary_rates():
    zero = R._rate_summary([False] * 40)
    one = R._rate_summary([True] * 40)
    assert zero['mc_se'] == 0 and zero['wilson_95'][1] > 0
    assert one['mc_se'] == 0 and one['wilson_95'][0] < 1
    assert zero['replications'] == one['replications'] == 40


def test_estimation_chunk_partial_count_and_cache_fingerprint():
    cal = B.calibrate_estimation(CFG['calibration']['estimation'])
    cfg = {'lo': 40, 'hi': 80, 'trials_per_state': 8, 'N': 2,
           'sd_log_nu': 0.2, 'sd_log_sigma0': 0.2, 'sd_log_f': 0.03}
    chunk = R.estimation_chunk(cal, cfg, 1.25, 0, 1, 17)
    assert len(chunk['reps']) == 1 and chunk['fit_count'] == 2
    assert len(chunk['reps'][0]) == 2
    fp0 = R.estimation_cache_fingerprint(cal, cfg, 1.25, 17, 10)
    cfg_changed = dict(cfg, trials_per_state=9)
    fp1 = R.estimation_cache_fingerprint(cal, cfg_changed, 1.25, 17, 10)
    assert fp0 != fp1


def test_pair_specific_complete_case_sensitivity_structure_direction_and_duality():
    # Synthetic data give larger within-person dispersion at wider priors,
    # while keeping six participants balanced across observed order strata.
    with tempfile.TemporaryDirectory(dir=ROOT) as temp_dir:
        data_path = os.path.join(temp_dir, 'synthetic_estimation.csv')
        with open(data_path, 'w', newline='', encoding='utf-8') as stream:
            writer = csv.writer(stream)
            writer.writerow(['ID', 'Order', 'Delta', 'Number', 'Response'])
            for participant in range(6):
                order = 'Widening' if participant < 3 else 'Narrowing'
                for width, scale in ((20, 1.0), (40, 2.0), (60, 4.0)):
                    for x in (50, 51):
                        center = 100.0 * participant + x
                        for value in (-scale, 0.0, scale):
                            writer.writerow([f'p{participant}', order, width, x, center + value])

        config = {'empirical_reanalysis': {
            'widths': [20, 40, 60], 'shared_number_min': 50,
            'shared_number_max': 51, 'familywise_alpha': 0.05,
        }}
        result = CCS.analyze(data_path, config, reps=200, seed=321, chunk_size=32)

    assert result['status'] == 'secondary_post_hoc_sensitivity_not_confirmatory'
    assert result['metadata']['planned_family_size'] == 6
    assert result['metadata']['total_bootstrap_draws_requested'] == 1200
    assert len(result['contrasts']) == 6 and len(result['pairwise_summary']) == 3
    expected_ratios = {(20, 40): 4.0, (20, 60): 16.0, (40, 60): 4.0}
    for row in result['contrasts']:
        pair = (row['width_1'], row['width_2'])
        assert np.isclose(row['variance_ratio_width_2_over_width_1'], expected_ratios[pair])
        assert row['matched_participants'] == 6
        assert row['matched_participants_by_order'] == {'Widening': 3, 'Narrowing': 3}
        assert row['significant_familywise'] == row['familywise_interval_excludes_one']
        lo, hi = row['familywise_95_ci_variance_ratio']
        assert row['significant_familywise'] == (lo > 1.0 or hi < 1.0)
    assert all(row['significant_numerosities_familywise'] == 2
               for row in result['pairwise_summary'])


def test_results_file_consistency():
    path = os.path.join(ROOT, 'results.json')
    if not os.path.exists(path):
        return
    r = json.load(open(path))
    assert r['config'] == CFG, 'results.json was not generated from the active config.json'
    assert r['empirical_reanalysis']['n_rows'] == 12960
    paired = r['empirical_reanalysis']['per_numerosity_cluster_bootstrap']
    assert paired['n_clusters'] == 36
    assert paired['shared_numerosities'] == list(range(50, 71))
    assert paired['planned_family_size'] == 63 and len(paired['contrasts']) == 63
    assert len(paired['pairwise_summary']) == 3
    assert len(paired['per_numerosity']) == 21
    for level in paired['per_numerosity']:
        by_width = level['participants_with_residual_df_by_width']
        by_pair = level['participants_with_residual_df_both_widths']
        assert set(by_pair) == {'20/40', '20/60', '40/60'}
        assert by_pair['20/40'] <= min(by_width['20'], by_width['40'])
        assert by_pair['20/60'] <= min(by_width['20'], by_width['60'])
        assert by_pair['40/60'] <= min(by_width['40'], by_width['60'])
    assert all(row['participants_with_df_both_widths'] > 0 for row in paired['contrasts'])
    assert [x['significant_numerosities'] for x in paired['pairwise_summary']] == [
        sum(row['significant_familywise'] for row in paired['contrasts']
            if (row['width_1'], row['width_2']) == (a, b))
        for a, b in ((20, 40), (20, 60), (40, 60))]
    expected_net = V.net_increment_analysis(CFG['budget'])
    assert np.isclose(r['net_increment']['baseline']['phi_approx'], expected_net['baseline']['phi_approx'], rtol=1e-12)
    assert np.isclose(r['headline']['lemma3_approx_rel_err'], expected_net['baseline_approx_rel_err'], rtol=1e-12)
    h = r['headline']
    assert abs(h['common_factor_price_x2'] - 2**0.25) < 1e-12
    for row in r['power']['by_price_ratio']:
        assert all(u['replications'] == CFG['power']['reps'] for u in row['mc_uncertainty'].values())
        if row['price_ratio'] >= 1.5:
            p = [row[f'power_N{N}'] for N in r['power']['N_grid']]
            assert p[-1] >= p[0]
    for row in r['estimation_recovery']:
        assert row['reps'] == CFG['estimation_recovery']['reps']
        assert all(u['replications'] == CFG['estimation_recovery']['reps']
                   for u in row['mc_uncertainty'].values())
    for row in r['lapse_robustness']:
        assert row['reps'] == CFG['lapse_robustness']['reps']
        assert all(u['replications'] == CFG['lapse_robustness']['reps']
                   for u in row['mc_uncertainty'].values())
    false_pos = [row for row in r['power']['by_price_ratio'] if row['price_ratio'] == 1.0][0]
    assert all(false_pos[f'power_N{N}'] <= 0.1 for N in r['power']['N_grid'])
    position = r['empirical_reanalysis']['within_block_position_sensitivity']['contrasts']
    assert len(position) == 6
    assert h['empirical_position_all_contrasts_supported'] == sum(
        bool(row['significant_familywise']) for row in position)
    assert np.isclose(h['empirical_position_min_adjusted_pvalue'],
                      min(row['pvalue_bonferroni'] for row in position))
    assert np.isclose(h['empirical_position_max_adjusted_pvalue'],
                      max(row['pvalue_bonferroni'] for row in position))
    assert h['empirical_position_all_ratios_above_one'] == all(
        row['variance_ratio_width_2_over_width_1'] > 1 for row in position)


def test_net_increment_and_displacement_invariants():
    cfg = CFG['budget']
    na = V.net_increment_analysis(cfg)
    assert na['gross_falls_with_price'] and na['net_rises_with_price']
    assert na['baseline_approx_rel_err'] < 0.05
    sp_ = [r['net_over_gross'] for r in na['same_price_varying_stiffness']]
    assert all(abs(r['mu'] / na['baseline']['mu'] - 1) < 1e-6 for r in na['same_price_varying_stiffness'])
    assert sp_[0] > 0.4 and sp_[-1] < 1e-3
    ins = V.integer_sweep()
    assert abs(ins['max_nc_without_effect'] - 6**0.5) < 0.01
    dr = V.displacement_remark()
    assert abs(dr['proportional (D = 0.6 E)']['slope'] + 0.5) < 1e-9
    high_displacement = dr['displacement falls by 1.5 times the energy decline']
    assert high_displacement['slope'] > 0
    assert min(high_displacement[k] for k in ('D0', 'D1', 'net_energy0', 'net_energy1')) >= 0
    r = analytics.derive()
    for key in ('F_mu_general_k_positive', 'lemma3_phi_formula_ok', 'lemma3_phi_k1_ok', 'lemma3_el_phi_mu_ok',
                'remark_proportional_displacement_preserves_slope'):
        assert r[key], key


def test_lemma3_first_order_coefficient_uses_no_task_equilibrium():
    b = CFG['budget']['baseline']
    mu0 = Mo.budget_mu(b['M'], b['c'], b['v'], b['rho'], b['e'], 0.0, b['k'])
    d0 = (mu0 / b['c']) ** (1 / b['k'])
    Eo0 = (b['v'] / mu0) ** (1 / b['rho'])
    phi0 = (d0 / b['k']) / (d0 / b['k'] + Eo0 / b['rho'])

    eps = 1e-3
    mu_eps = Mo.budget_mu(b['M'], b['c'], b['v'], b['rho'], b['e'], eps * b['A'], b['k'])
    d_eps = (mu_eps / b['c']) ** (1 / b['k'])
    E_eps = np.sqrt(b['e']) * eps * b['A'] / np.sqrt(mu_eps)
    finite_difference = (d_eps - d0) / E_eps
    assert abs(finite_difference / phi0 - 1) < 5e-5

    result = V.net_increment_analysis(CFG['budget'])
    assert np.isclose(result['baseline']['phi_approx'], phi0, rtol=1e-12)
    expected_first_order = phi0 * result['baseline']['gross_E_no_task']
    assert np.isclose(result['baseline']['first_order_net_dB'], expected_first_order, rtol=1e-12)
    exact_increment = result['baseline']['net_dB']
    assert np.isclose(result['baseline_approx_rel_err'],
                      abs(expected_first_order / exact_increment - 1), rtol=1e-12)
    assert result['baseline_approx_rel_err'] < 0.02


def test_published_trial_data_reanalysis():
    c = CFG['empirical_reanalysis']
    path = os.path.normpath(os.path.join(ROOT, c['data_path']))
    result = Emp.reanalyze_estimation(
        path, widths=c['widths'], anchor_number=c['anchor_number'],
        shared_numbers=range(c['shared_number_min'], c['shared_number_max'] + 1),
        published_sd=c['published_sd'], familywise_alpha=c['familywise_alpha'],
        bootstrap_reps=4000, bootstrap_seed=c['bootstrap_seed'],
        equal_weight_bootstrap_reps=8000, position_bootstrap_reps=4000)
    assert result['n_rows'] == 12960 and result['n_subjects'] == 36
    assert len(result['anchors']) == 3 and max(abs(x['difference']) for x in result['anchors']) < 0.15
    paired = result['per_numerosity_cluster_bootstrap']
    assert paired['shared_numerosities'] == list(range(c['shared_number_min'], c['shared_number_max'] + 1))
    assert paired['n_clusters'] == 36 and paired['planned_family_size'] == 63
    assert paired['sequence_counts'] == {'Widening': 20, 'Narrowing': 16}
    assert paired['observed_block_orders'] == {
        'Widening': [20, 40, 60], 'Narrowing': [60, 40, 20]}
    assert len(paired['contrasts']) == 63 and len(paired['pairwise_summary']) == 3
    assert all(row['participants_with_df_width_1'] > 0 and row['participants_with_df_width_2'] > 0
               for row in paired['contrasts'])
    assert all(row['pvalue_bonferroni'] >= row['pvalue_bootstrap'] for row in paired['contrasts'])
    assert all(np.isfinite(row['pvalue_bootstrap_mcse']) and row['pvalue_bootstrap_mcse'] > 0
               for row in paired['contrasts'])
    assert all(len(row['pvalue_bootstrap_mc_95']) == 2 and row['pvalue_bootstrap_mc_95'][1] > 0
               for row in paired['contrasts'])
    for row in paired['contrasts']:
        lo, hi = row['familywise_ci_variance_ratio']
        assert row['significant_familywise'] == (lo > 1 or hi < 1)
    equal = result['participant_equal_weight_sensitivity']
    assert equal['complete_participants'] == 36
    assert len(equal['contrasts']) == 3
    position = result['within_block_position_sensitivity']
    assert len(position['contrasts']) == 6
    assert {row['block_half'] for row in position['contrasts']} == {'first_half', 'second_half'}
    assert all(row['bootstrap_reps'] == 4000 for row in position['contrasts'])
    assert all(row['participants_with_df_width_1'] > 0 and row['participants_with_df_width_2'] > 0
               for row in position['contrasts'])


def test_sequence_table_labels_match_ratio_direction():
    paper = os.path.normpath(os.path.join(ROOT, '..', 'paper'))
    template = os.path.join(paper, 'supplement.md')
    main_template = os.path.join(paper, 'body.md')
    results = os.path.join(ROOT, 'results.json')
    sensitivity = os.path.join(ROOT, 'complete_case_sensitivity.json')
    sign_sensitivity = os.path.join(ROOT, 'participant_sign_sensitivity.json')
    with tempfile.TemporaryDirectory(dir=ROOT) as temp_dir:
        output = os.path.join(temp_dir, 'supplement_filled.md')
        subprocess.run([sys.executable, os.path.join(ROOT, 'fill_numbers.py'),
                        template, results, sensitivity, sign_sensitivity,
                        '--extra=' + os.path.join(ROOT, 'model_predicted_ratios.json'), output],
                       check=True, capture_output=True, text=True)
        text = open(output, encoding='utf-8').read()
        main_output = os.path.join(temp_dir, 'main_filled.md')
        subprocess.run([sys.executable, os.path.join(ROOT, 'fill_numbers.py'),
                        main_template, results, sensitivity, sign_sensitivity,
                        '--extra=' + os.path.join(ROOT, 'model_predicted_ratios.json'), main_output],
                       check=True, capture_output=True, text=True)
        main_text = open(main_output, encoding='utf-8').read()
    table = text.split('| Prior-width contrast | Widening sequence', 1)[1].split('**Table S4.', 1)[0]
    assert '| 40 / 20 |' in table
    assert '| 60 / 20 |' in table
    assert '| 60 / 40 |' in table
    assert '| 20 / 40 |' not in table and '| 20 / 60 |' not in table
    combined = main_text + text  # the full reanalysis and its diagnostics are reported in the supplement
    assert '{{complete_case_sensitivity_summary}}' not in combined
    assert '2.93 for 40/20' in combined and '4.07 for 60/20' in combined and '1.41 for 60/40' in combined
    assert '{{lapse_fit_diagnostics_main}}' not in combined
    assert '{{lapse_fit_diagnostics_supplement}}' not in text
    assert '30.7%' in text and '35.3%' in text
    assert '0.0021 normalized units' in combined  # the numerical results are reported in Supplementary Section S9
    assert '0.0021 normalized units' in text
    discussion = main_text.split('## Discussion', 1)[1].split('## Conclusion', 1)[0]
    assert 'without preregistration or an independent validation sample' in discussion
    assert 'not as prospectively preregistered confirmatory testing' not in combined
    assert 'The fixed-numerosity contrasts are assembled in Table S2' in text
    sequence_results = json.load(open(results, encoding='utf-8'))['empirical_reanalysis']['order_sequence_sensitivity']['contrasts']
    for row in sequence_results:
        label = f"| {row['width_2']} / {row['width_1']} |"
        widening = row['widening']['geometric_mean_within_participant_variance_ratio']
        narrowing = row['narrowing']['geometric_mean_within_participant_variance_ratio']
        table_row = next((line for line in table.splitlines() if line.startswith(label)), '')
        assert table_row
        assert f"| {widening:.2f} [" in table_row and f"| {narrowing:.2f} [" in table_row
        assert widening > 1 and narrowing > 1
    assert text.count('**Table S3.') == 1
    assert text.count('| Prior-width contrast | Paired participants | Equal-weight geometric mean ratio |') == 1
    assert 'Table S10' not in text
    assert '**Table S6.** Simulated rejection rates' in text
    assert 'Fit diagnostics are summarized separately' in text
    assert '4/21 for 40/20' in combined
    assert '1/21 for 60/20' in combined
    assert '0/21 for 60/40' in combined
    assert '**Table S8.** Exact participant-level sign-balance results' in text
    s7 = text.split('## S7.', 1)[1].split('**Table S8.', 1)[0]
    s8_rows = [line for line in s7.splitlines()
               if line.startswith('| ') and line.split('|')[1].strip().isdigit()]
    assert len(s8_rows) == 63
    assert sum(line.split('|')[1].strip() == '50' for line in s8_rows) == 3
    assert 'one-zero pair remains directional' in text
    assert '{{' not in main_text and '{{' not in text


def test_participant_sign_sensitivity_is_reproducible_and_targets_all_63_contrasts():
    result = PSS.analyze()
    saved = json.load(open(os.path.join(ROOT, 'participant_sign_sensitivity.json'), encoding='utf-8'))
    assert result == saved
    assert result['family_size'] == 63
    assert len(result['contrasts']) == 63
    assert result['data_sha256'] == PSS._data_sha256(PSS.DEFAULT_DATA)
    summary = {row['contrast']: row['corrected_numerosities'] for row in result['summary']}
    assert summary == {'40/20': 4, '60/20': 1, '60/40': 0}
    assert all(row['n_eligible'] == row['n_positive_wider_variance'] +
               row['n_negative_wider_variance'] + row['n_tied_variance_differences']
               for row in result['contrasts'])
    aggregate = result['aggregate_participant_direction']
    assert aggregate['family_size'] == 3
    aggregate_rows = {row['contrast']: row for row in aggregate['contrasts']}
    assert {contrast: (row['n_positive_wider_variance'], row['n_negative_wider_variance'],
                       row['n_tied_variance_ratios'])
            for contrast, row in aggregate_rows.items()} == {
                '40/20': (36, 0, 0), '60/20': (36, 0, 0), '60/40': (29, 7, 0)
            }
    assert all(row['bonferroni_adjusted_p_3'] < 0.001
               for row in aggregate_rows.values())
    assert all(row['n_eligible'] == result['n_participants']
               for row in aggregate_rows.values())


def test_tex_manuscripts_and_pdfs_are_complete():
    paper = os.path.normpath(os.path.join(ROOT, '..', 'paper'))
    main_tex = open(os.path.join(paper, 'Manuscript.tex'), encoding='utf-8').read()
    supp_tex = open(os.path.join(paper, 'Supplementary_Materials.tex'), encoding='utf-8').read()
    captions = re.findall(r'\\caption\{(.*?)\}\\label\{fig:figure\d+\}', main_tex, flags=re.S)
    assert len(captions) == 6
    assert all(len(caption) >= 40 for caption in captions)
    figure_alt = re.findall(r'\\includegraphics\[alt=\{([\s\S]*?)\},width=', main_tex)
    assert len(figure_alt) == 6 and all(len(alt) >= 40 for alt in figure_alt)
    assert main_tex.count('\\includegraphics') == 6
    # The supplied PDFs are not tagged; do not claim tagged-PDF support.
    assert '\\DocumentMetadata{lang=en-US,pdfversion=2.0,tagging=on}' not in main_tex
    assert '\\tagpdfsetup{table/header-rows={1}}' not in main_tex
    assert main_tex.count('\\begin{figure}[H]\\centering') == 6
    assert 'textbackslash{}(' not in supp_tex
    assert ('Under no price change, estimated rejection rates at N = 20, 40 and 80 are '
            '0.04 (0.02--0.08), 0.04 (0.02--0.07) and 0.045 (0.02--0.08)'
            in supp_tex)
    for name in ('Manuscript.pdf', 'Supplementary_Materials.pdf'):
        with open(os.path.join(paper, name), 'rb') as stream:
            assert stream.read(5) == b'%PDF-'


def test_every_bibliography_entry_is_cited_in_the_manuscript_or_supplement():
    paper = os.path.normpath(os.path.join(ROOT, '..', 'paper'))
    refs = json.load(open(os.path.join(paper, 'refs.json'), encoding='utf-8'))
    source = '\n'.join(open(os.path.join(paper, name), encoding='utf-8').read()
                        for name in ('body.md', 'supplement.md'))
    groups = re.findall(r'\[\[([^\[\]]+)\]\]', source)
    cited = {key.strip().lstrip('@') for group in groups for key in group.split(';')}
    assert cited == set(refs), {
        'uncited': sorted(set(refs) - cited),
        'undefined': sorted(cited - set(refs)),
    }


if __name__ == '__main__':
    tests = [v for k, v in dict(globals()).items() if k.startswith('test_')]
    source_data = os.path.normpath(os.path.join(ROOT, CFG['empirical_reanalysis']['data_path']))
    source_data_tests = {
        'test_published_trial_data_reanalysis',
        'test_participant_sign_sensitivity_is_reproducible_and_targets_all_63_contrasts',
    }
    passed = 0
    skipped = 0
    for t in tests:
        if t.__name__ in source_data_tests and not os.path.isfile(source_data):
            print(f'skipped {t.__name__}: source trial CSV is not present ({source_data})')
            skipped += 1
            continue
        t()
        print('passed', t.__name__)
        passed += 1
    print(f'{passed} tests passed; {skipped} skipped; {len(tests)} total')
```
