## Supplementary material for "The marginal brain": Replication package: write_package_manifest.py

```
"""Write a SHA-256 inventory for the curated replication-package contents."""
from __future__ import annotations

import hashlib
import json
from pathlib import Path

ROOT = Path(__file__).resolve().parents[1]
OUTPUT = ROOT / "model" / "package_manifest.json"

# This inventory mirrors the files shipped in the replication ZIP. The manifest
# file is intentionally excluded from its own file list to avoid a hash cycle.
PACKAGE_FILES = (
    "README.md", "LICENSE", "reproduce.ps1", "reproduce.sh", "requirements.txt", "verify.sh",
    "data/README.md",
    "model/README.md", "model/LICENSE", "model/config.json",
    "model/analysis_record.md", "model/complete_case_sensitivity.py",
    "model/complete_case_sensitivity.json", "model/model_predicted_ratios.json", "model/predicted_variance_ratios.py", "model/participant_sign_sensitivity.py",
    "model/participant_sign_sensitivity.json", "model/conditional_null_calibration.py",
    "model/outputs/conditional_null_calibration/pilot_summary.json",
    "model/outputs/conditional_null_calibration/pilot_test_results.csv",
    "model/outputs/conditional_null_calibration/toy_validation.json",
    "model/fill_numbers.py", "model/precompute_estimation.py", "model/refresh_derived.py",
    "model/requirements.txt", "model/results.json", "model/run_all.py",
    "model/tests/test_model.py", "model/spt/__init__.py", "model/spt/analytics.py",
    "model/spt/behavior.py", "model/spt/empirical.py", "model/spt/figures.py",
    "model/spt/model.py", "model/spt/numerics.py", "model/spt/recovery.py",
    "model/spt/validation.py", "model/write_package_manifest.py",
    "model/figures/fig1_signatures.pdf", "model/figures/fig1_signatures.png",
    "model/figures/fig2_power.pdf", "model/figures/fig2_power.png",
    "paper/body.md", "paper/fig6_empirical.py", "paper/fig_common_factor.py", "paper/fig_rest.py",
    "paper/fig_schematic.py", "paper/refs.json", "paper/supplement.md", "paper/typeset.py",
    "paper/Manuscript.pdf", "paper/Manuscript.tex",
    "paper/Supplementary_Materials.pdf", "paper/Supplementary_Materials.tex",
    "paper/build/markdown_to_tex.py", "paper/build/compile_tex.ps1", "paper/build/compile_tex.sh",
    "paper/build/mathfix.py",
    "paper/figures/fig1_schematic.pdf", "paper/figures/fig1_schematic.png",
    "paper/figures/fig2_common_factor.pdf", "paper/figures/fig2_common_factor.png",
    "paper/figures/fig3_budget.pdf", "paper/figures/fig3_budget.png",
    "paper/figures/fig4_sign_test.pdf", "paper/figures/fig4_sign_test.png",
    "paper/figures/fig5_power.pdf", "paper/figures/fig5_power.png",
    "paper/figures/fig6_empirical.pdf", "paper/figures/fig6_empirical.png",
)


def sha256(path: Path) -> str:
    digest = hashlib.sha256()
    with path.open("rb") as stream:
        for chunk in iter(lambda: stream.read(1024 * 1024), b""):
            digest.update(chunk)
    return digest.hexdigest()


def main() -> None:
    missing = [name for name in PACKAGE_FILES if not (ROOT / name).is_file()]
    if missing:
        raise FileNotFoundError("Replication-package files missing: " + ", ".join(missing))
    files = {name: sha256(ROOT / name) for name in PACKAGE_FILES}
    canonical = json.dumps(files, sort_keys=True, separators=(",", ":")).encode("utf-8")
    payload = {
        "manifest_format": 1,
        "scope": "SHA-256 inventory of the curated replication ZIP contents at package-build time.",
        "self_exclusion": "model/package_manifest.json is excluded from its own file list.",
        "campaign_source_manifest": "model/results.json meta.source_manifest records the separate campaign-time source snapshot.",
        "files": files,
        "file_inventory_sha256": hashlib.sha256(canonical).hexdigest(),
    }
    OUTPUT.write_text(json.dumps(payload, indent=2, ensure_ascii=False) + "\n", encoding="utf-8")
    print(f"Wrote {len(files)} file hashes to {OUTPUT}")


if __name__ == "__main__":
    main()
```
