## Supplementary material for "The marginal brain": Replication package: compile_tex.ps1

```
$ErrorActionPreference = 'Stop'
$paper = Split-Path -Parent (Split-Path -Parent $MyInvocation.MyCommand.Path)
$python = $env:REPRO_PYTHON
$pythonPrefix = @()
if (-not $python) {
    $launcher = Get-Command py -ErrorAction SilentlyContinue
    if ($launcher) {
        & $launcher.Source -3 -c 'import sys' *> $null
        if ($LASTEXITCODE -eq 0) { $python = $launcher.Source; $pythonPrefix = @('-3') }
    }
}
if (-not $python) {
    $candidate = Get-Command python -ErrorAction SilentlyContinue
    if ($candidate -and $candidate.Source -notmatch '\\WindowsApps\\') { $python = $candidate.Source }
}
if (-not $python) { throw 'Python 3 is required. Set REPRO_PYTHON or add Python to PATH.' }

$latexPaths = [System.Collections.Generic.List[string]]::new()
foreach ($engine in @('lualatex.exe', 'xelatex.exe')) {
    $command = Get-Command $engine -ErrorAction SilentlyContinue
    if ($command) { $latexPaths.Add($command.Source); continue }
    $miKTeX = Join-Path 'C:\Program Files\MiKTeX\miktex\bin\x64' $engine
    if (Test-Path -LiteralPath $miKTeX) { $latexPaths.Add($miKTeX) }
}
if ($latexPaths.Count -eq 0) {
    throw 'LuaLaTeX or XeLaTeX was not found. Install MiKTeX or TeX Live and add one engine to PATH.'
}

Push-Location $paper
try {
    foreach ($name in @('Manuscript', 'Supplementary_Materials')) {
        $converterArgs = @()
        if ($name -eq 'Manuscript') { $sourceName = '_main_manuscript.md' }
        else { $sourceName = '_supplement_manuscript.md'; $converterArgs = @('--supplement') }
        & $python @pythonPrefix 'build\markdown_to_tex.py' $sourceName 'refs.json' "$name.tex" @converterArgs
        if ($LASTEXITCODE -ne 0) { throw "LaTeX source generation failed for $name." }
    }
    $output = 'build\tex_output'
    New-Item -ItemType Directory -Force -Path $output | Out-Null
    $outputOption = "-output-directory=$output"
    foreach ($name in @('Manuscript', 'Supplementary_Materials')) {
        $compiled = $false
        foreach ($latexPath in $latexPaths) {
            $engineSucceeded = $true
            for ($pass = 1; $pass -le 2; $pass++) {
                & $latexPath '-interaction=nonstopmode' '-halt-on-error' $outputOption "$name.tex"
                if ($LASTEXITCODE -ne 0) { $engineSucceeded = $false; break }
            }
            if ($engineSucceeded) { $compiled = $true; break }
        }
        if (-not $compiled) { throw "LuaLaTeX and XeLaTeX could not compile $name." }
        Copy-Item -LiteralPath (Join-Path $output "$name.pdf") -Destination "$name.pdf" -Force
    }
} finally { Pop-Location }
Write-Output 'Both TeX sources and PDFs were generated.'
```
