## Supplementary material for "The marginal brain": Replication package: compile_tex.sh

```
#!/bin/sh
set -eu
PAPER=$(CDPATH= cd -- "$(dirname -- "$0")/.." && pwd)
PYTHON=${REPRO_PYTHON:-${PYTHON:-python3}}
ENGINE=${TEX_ENGINE:-lualatex}
command -v "$ENGINE" >/dev/null 2>&1 || ENGINE=xelatex
command -v "$ENGINE" >/dev/null 2>&1 || { echo 'LuaLaTeX or XeLaTeX is required to build the manuscript PDFs.' >&2; exit 1; }
cd "$PAPER"
"$PYTHON" build/markdown_to_tex.py _main_manuscript.md refs.json Manuscript.tex
"$PYTHON" build/markdown_to_tex.py _supplement_manuscript.md refs.json Supplementary_Materials.tex --supplement
mkdir -p build/tex_output
for name in Manuscript Supplementary_Materials; do
  # LuaLaTeX first; XeLaTeX when the LuaTeX font database cannot load the OpenType fonts (minimal TeX installations)
  if ! "$ENGINE" -interaction=nonstopmode -halt-on-error -output-directory=build/tex_output "$name.tex" >/dev/null; then
    ENGINE=xelatex
    xelatex -interaction=nonstopmode -halt-on-error -output-directory=build/tex_output "$name.tex" >/dev/null
  fi
  "$ENGINE" -interaction=nonstopmode -halt-on-error -output-directory=build/tex_output "$name.tex" >/dev/null
  cp "build/tex_output/$name.pdf" "$name.pdf"
done
printf '%s\n' "Both TeX sources and PDFs were generated with $ENGINE."
```
