## Supplementary material for "The marginal brain": Replication package: markdown_to_tex.py

```
#!/usr/bin/env python3
"""Convert the filled manuscript Markdown into a portable, self-contained TeX source."""

from __future__ import annotations

import argparse
import json
import re
import unicodedata
from collections import defaultdict
from pathlib import Path


TEX_ESCAPES = {
    "\\": r"\textbackslash{}",
    "#": r"\#",
    "$": r"\$",
    "%": r"\%",
    "&": r"\&",
    "_": r"\_",
    "{": r"\{",
    "}": r"\}",
    "~": r"\textasciitilde{}",
    "^": r"\textasciicircum{}",
}

UNICODE_TEX_ESCAPES = {
    "–": "--", "—": "---", "…": r"\ldots{}", "′": r"\ensuremath{^\prime}",
    "’": "'", "“": "``", "”": "''", "ç": r"\c{c}", "ć": r"\'{c}",
    "ě": r"\v{e}", "í": r"\'{i}", "\u00a0": "~",
}


def tex_escape(text: str) -> str:
    return "".join(UNICODE_TEX_ESCAPES.get(char, TEX_ESCAPES.get(char, char)) for char in text)


def markdown_inline(text: str, references: dict) -> str:
    protected: list[str] = []

    def hold(value: str) -> str:
        protected.append(value)
        return f"ZZTEXTOKEN{len(protected) - 1}ZZ"

    # Web addresses are typeset verbatim with \url, so underscores and hyphens in them are neither
    # escaped nor read as subscripts, and long addresses can break across lines.
    def protect_url(m):
        url = m.group(0)
        trail = ""
        while url and url[-1] in ".,;:":
            trail = url[-1] + trail
            url = url[:-1]
        return hold(r"\url{" + url + "}") + trail
    text = re.sub(r"https?://[^\s<>()\]]+", protect_url, text)

    # Preserve TeX math before escaping ordinary prose.  The source manuscript
    # uses Markdown-style \(...\) and \[...\] delimiters; escaping their
    # backslashes would print the delimiters literally in the compiled PDF.
    def math_content(value: str) -> str:
        replacements = {
            "∝": r"\propto", "∗": r"\ast", "∈": r"\in",
            "≤": r"\leq", "≥": r"\geq", "≠": r"\neq",
            "≈": r"\approx", "×": r"\times", "Σ": r"\sum",
            "α": r"\alpha", "β": r"\beta", "γ": r"\gamma",
            "δ": r"\delta", "κ": r"\kappa", "λ": r"\lambda",
            "μ": r"\mu", "ν": r"\nu", "π": r"\pi",
            "ρ": r"\rho", "σ": r"\sigma", "θ": r"\theta",
            "−": "-", "–": "-", "′": r"\prime",
        }
        for symbol, command in replacements.items():
            value = value.replace(symbol, command)
        superscripts = str.maketrans("⁰¹²³⁴⁵⁶⁷⁸⁹⁺⁻⁼", "0123456789+-=")
        value = re.sub(r"[⁰¹²³⁴⁵⁶⁷⁸⁹⁺⁻⁼]+", lambda m: "^{" + m.group(0).translate(superscripts) + "}", value)
        return value

    def hold_math(match: re.Match[str]) -> str:
        token = match.group(0)
        if token.startswith(r"\["):
            return hold(r"\[" + math_content(token[2:-2].strip()) + r"\]")
        if token.startswith(r"\("):
            return hold(r"\(" + math_content(token[2:-2].strip()) + r"\)")
        if token.startswith("$$"):
            return hold(r"\[" + math_content(token[2:-2].strip()) + r"\]")
        return hold(r"\(" + math_content(token[1:-1].strip()) + r"\)")

    text = re.sub(r"\\\[[\s\S]*?\\\]|\\\([\s\S]*?\\\)|\$\$[\s\S]*?\$\$|(?<!\\)\$[^$\n]+?\$", hold_math, text)

    # Map mathematical Unicode in prose to the same Latin Modern math fonts
    # used by the companion manuscripts.  Keeping these symbols in math mode
    # avoids font-encoding substitutions and missing Greek/superscript glyphs.
    text = re.sub(
        r"([0-9])([⁰¹²³⁴⁵⁶⁷⁸⁹⁺⁻]+)",
        lambda m: m.group(1) + hold(r"\textsuperscript{" + m.group(2).translate(str.maketrans("⁰¹²³⁴⁵⁶⁷⁸⁹⁺⁻", "0123456789+-")) + "}"),
        text,
    )
    text = text.replace("√K", hold(r"\ensuremath{\sqrt{K}}"))
    text_symbols = {
        "∝": r"\ensuremath{\propto}", "∗": r"\ensuremath{\ast}",
        "∈": r"\ensuremath{\in}", "≤": r"\ensuremath{\leq}",
        "≥": r"\ensuremath{\geq}", "≠": r"\ensuremath{\neq}",
        "≈": r"\ensuremath{\approx}", "×": r"\ensuremath{\times}",
        "−": r"\ensuremath{-}", "∫": r"\ensuremath{\int}",
        "∂": r"\ensuremath{\partial}",
        "→": r"\ensuremath{\rightarrow}", "Σ": r"\ensuremath{\Sigma}",
        "Δ": r"\ensuremath{\Delta}", "α": r"\ensuremath{\alpha}",
        "β": r"\ensuremath{\beta}", "γ": r"\ensuremath{\gamma}",
        "δ": r"\ensuremath{\delta}", "ε": r"\ensuremath{\varepsilon}",
        "θ": r"\ensuremath{\theta}", "κ": r"\ensuremath{\kappa}",
        "λ": r"\ensuremath{\lambda}", "μ": r"\ensuremath{\mu}",
        "ν": r"\ensuremath{\nu}", "π": r"\ensuremath{\pi}",
        "ρ": r"\ensuremath{\rho}", "σ": r"\ensuremath{\sigma}",
        "φ": r"\ensuremath{\phi}", "ω": r"\ensuremath{\omega}",
        "½": r"\ensuremath{\frac{1}{2}}", "¼": r"\ensuremath{\frac{1}{4}}",
        "¾": r"\ensuremath{\frac{3}{4}}", "¹": r"\textsuperscript{1}",
        "²": r"\textsuperscript{2}", "³": r"\textsuperscript{3}",
        "⁴": r"\textsuperscript{4}", "⁵": r"\textsuperscript{5}",
        "⁷": r"\textsuperscript{7}", "⁻": r"\textsuperscript{-}",
        "₀": r"\textsubscript{0}", "₁": r"\textsubscript{1}",
        "₂": r"\textsubscript{2}", "ₐ": r"\textsubscript{a}",
        "ₒ": r"\textsubscript{o}", "ᵢ": r"\textsubscript{i}",
        "ᵘ": r"\textsuperscript{u}", "ᵏ": r"\textsuperscript{k}",
        # Use TeX-native punctuation so the T1 Latin Modern output has no
        # missing Unicode glyphs under either pdfTeX or XeTeX.
        "–": "--", "—": "---", "…": r"\ldots{}", "′": r"\ensuremath{^\prime}",
        "’": "'", "“": "``", "”": "''",
    }
    text_accents = {
        "ç": r"\c{c}", "ć": r"\'{c}", "ě": r"\v{e}", "í": r"\'{i}",
    }
    for letter, command in text_accents.items():
        text = text.replace(letter, hold(command))
    for letter, command in {
        "B": r"\ensuremath{\bar{B}}",
    }.items():
        text = text.replace(letter + "\u0304", hold(command))
    for symbol, value in text_symbols.items():
        text = text.replace(symbol, hold(value))

    def cite(match: re.Match[str]) -> str:
        parts = [part.strip() for part in match.group(1).split(";") if part.strip()]
        keys: list[str] = []
        locator = ""
        narrative = False
        for part in parts:
            if part.startswith("@"):
                narrative = True
                part = part[1:]
            if "|" in part:
                key, loc = part.split("|", 1)
                part = key.strip()
                locator = loc.strip()
            part = part.strip()
            if part in references:
                keys.append(part)
        if not keys:
            return match.group(0)
        command = "citet" if narrative and len(keys) == 1 else "citep"
        optional = f"[{tex_escape(locator)}]" if locator else ""
        return hold(rf"\{command}{optional}{{{','.join(keys)}}}")

    text = re.sub(r"\[\[([^\]]+)\]\]", cite, text)

    def code(match: re.Match[str]) -> str:
        return hold(r"\texttt{" + tex_escape(match.group(1)) + "}")

    text = re.sub(r"`([^`]+)`", code, text)

    def sub(match: re.Match[str]) -> str:
        base = tex_escape(match.group(1))
        val = match.group(2).strip("{}")
        return hold(base + r"\textsubscript{" + tex_escape(val) + "}")

    def sup(match: re.Match[str]) -> str:
        base = tex_escape(match.group(1))
        val = match.group(2).strip("{}")
        return hold(base + r"\textsuperscript{" + tex_escape(val) + "}")

    text = re.sub(r"([A-Za-z0-9Α-Ωα-ω]+)_(\{[^{}]*\}|[A-Za-z0-9]+)", sub, text)
    text = re.sub(r"([A-Za-z0-9Α-Ωα-ω)\]]+)\^(\{[^{}]*\}|[A-Za-z0-9]+)", sup, text)

    def style(pattern: str, command: str, value: str) -> str:
        return re.sub(pattern, lambda m: hold(rf"\{command}{{{markdown_inline(m.group(1), references)}}}"), value)

    text = style(r"\*\*(.+?)\*\*", "textbf", text)
    text = style(r"(?<!\*)\*([^*]+?)\*(?!\*)", "emph", text)
    text = tex_escape(text)
    for i in range(len(protected) - 1, -1, -1):
        text = text.replace(f"ZZTEXTOKEN{i}ZZ", protected[i])
    return text


def parse_front_matter(source: str) -> tuple[dict[str, str], str]:
    if not source.startswith("---\n"):
        return {}, source
    _, front, rest = source.split("---", 2)
    values: dict[str, str] = {}
    for line in front.strip().splitlines():
        if ":" in line:
            key, value = line.split(":", 1)
            values[key.strip().lower()] = value.strip().strip("\"'")
    return values, rest.lstrip("\r\n")


def author_tex(author_field: str, affiliation: str) -> str:
    """Authors and affiliations. A single shared affiliation is printed without markers; several affiliations
    keep their superscript markers (^{a}, ^{a,b}) and are printed one per line, followed by the correspondence line."""
    names = [name.strip() for name in re.split(r"\s+and\s+", author_field) if name.strip()]
    markers = re.findall(r"\^\{([a-z])\}", affiliation)
    multiple = len(set(markers)) > 1
    def name_tex(name):
        m = re.search(r"\s*\^\{([a-z,\s]+)\}\s*$", name)
        bare = name[:m.start()] if m else name
        sup = (r"\textsuperscript{" + m.group(1).replace(" ", "") + "}") if (m and multiple) else ""
        return tex_escape(bare.strip()) + sup
    author_block = r"\author{" + " and ".join(name_tex(n) for n in names) + "}"
    corr = ""
    aff = affiliation
    if "Correspondence:" in aff:
        aff, corr = aff.split("Correspondence:", 1)
        aff = aff.strip().rstrip(".").strip()
    if multiple:
        parts = [p for p in re.split(r"\^\{([a-z])\}\s*", aff) if p is not None]
        lines, i = [], 1
        while i < len(parts):
            label, text = parts[i], parts[i + 1].strip().rstrip(";").strip().rstrip(".")
            lines.append(r"\textsuperscript{" + label + "}" + tex_escape(text))
            i += 2
    else:
        lines = [tex_escape(re.sub(r"\^\{[a-z]\}\s*", "", aff).strip())]
    if corr:
        lines.append("Correspondence: " + tex_escape(corr.strip()))
    return author_block + "\n" + r"\address{" + r" \\ ".join(lines) + "}"


def author_list(entry: dict) -> str:
    return ", ".join(entry.get("authors", [])) or entry.get("org", "")


def citation_label(entry: dict, year: str) -> str:
    authors = entry.get("authors", [])
    surnames = [author.split(",", 1)[0].strip() for author in authors]
    if len(surnames) == 0:
        surname_label = entry.get("org", "Anonymous")
    elif len(surnames) == 1:
        surname_label = surnames[0]
    elif len(surnames) == 2:
        surname_label = f"{surnames[0]} and {surnames[1]}"
    else:
        surname_label = f"{surnames[0]} et al."
    return f"{surname_label}({year})"


def year_suffixes(refs: dict) -> dict[str, str]:
    groups: dict[tuple[tuple[str, ...], str], list[str]] = defaultdict(list)
    for key, item in refs.items():
        identity = (tuple(item.get("authors", [])), str(item.get("year", "")))
        groups[identity].append(key)
    suffixes: dict[str, str] = {}
    for keys in groups.values():
        if len(keys) > 1:
            # Letter suffixes follow the same title order as an alphabetized
            # author-date reference list.
            ordered_keys = sorted(keys, key=lambda key: (refs[key].get("title", "").casefold(), key))
            for index, key in enumerate(ordered_keys):
                suffixes[key] = chr(ord("a") + index)
    return suffixes


def bibliography_tex(keys: list[str], refs: dict) -> str:
    # Keep the bibliography compact enough to avoid a nearly empty final page.
    # DOI identifiers use compact linked labels to reduce wrapping in the reference list.
    out = [
        r"\begingroup",
        r"\footnotesize",
        r"\setlength{\parskip}{0pt}",
        r"\begin{thebibliography}{99}",
        r"\setlength{\itemsep}{0pt}",
        r"\setlength{\parsep}{0pt}",
    ]
    suffixes = year_suffixes(refs)
    def sort_key(key: str) -> tuple[str, str, str, str]:
        item = refs.get(key, {})
        authors = item.get("authors", [])
        first = authors[0].split(",", 1)[0].strip() if authors else item.get("org", "")
        normalized_author = "".join(
            char for char in unicodedata.normalize("NFKD", first).casefold()
            if not unicodedata.combining(char)
        )
        title = "".join(
            char for char in unicodedata.normalize("NFKD", item.get("title", "")).casefold()
            if not unicodedata.combining(char)
        )
        return (normalized_author, str(item.get("year", "")), title, key)

    for key in sorted(keys, key=sort_key):
        item = refs.get(key)
        if not item:
            continue
        authors = tex_escape(author_list(item))
        raw_year = str(item.get("year", ""))
        year = tex_escape(raw_year + suffixes.get(key, ""))
        title = tex_escape(item.get("title", ""))
        journal = tex_escape(item.get("journal", item.get("publisher", "")))
        volume = tex_escape(item.get("volume", ""))
        issue = tex_escape(item.get("issue", ""))
        pages = tex_escape(item.get("pages", ""))
        doi = item.get("doi", "")
        detail = journal
        if volume:
            detail += f", {volume}"
        if issue:
            detail += f"({issue})"
        if pages:
            detail += f", {pages}"
        label = tex_escape(citation_label(item, year))
        line = rf"\bibitem[{label}]{{{key}}} {authors} ({year}). \emph{{{title}}}. {detail}."
        if doi:
            line += rf" DOI: \href{{https://doi.org/{doi}}}{{{tex_escape(doi)}}}."
        elif item.get("url"):
            line += rf" \url{{{item['url']}}}."
        out.append(line)
    out.append(r"\end{thebibliography}")
    out.append(r"\endgroup")
    return "\n".join(out)


def table_cells(line: str) -> list[str]:
    return [cell.strip() for cell in line.strip().strip("|").split("|")]


FIGURE_ALT_TEXT = {
    "1": "Three-stage schematic. Panel A shows the shared-budget identities for committed use, the reallocation margin, task energy and competing uses. Panel B equates marginal regulatory cost with the marginal value of competing uses to define the shadow price. Panel C maps that price to effective cost per signal and the resulting imprecision response.",
    "2": "Line chart of imprecision versus prior width for estimation and discrimination at baseline and doubled effective signal price. Estimation and discrimination retain distinct width slopes of one-half and three-quarters; doubling the price shifts both curves upward by the common fourth-root factor.",
    "3": "Two side-by-side panels. Panel A plots the shadow price against the reallocation margin for three competing-use curvatures and a hard-cap benchmark. Panel B contrasts gross task energy with the net change in total use as the margin tightens; both energy measures are reported in units of 10⁻³ times the normalized set point.",
    "4": "Sign-test plot of changes in log imprecision against changes in log task energy. A pure price change follows slope minus one-half, whereas a pure technology change follows slope plus one-half. Model markers include a mixed scarcity-and-technology change on the price-dominant side.",
    "5": "Simulation chart of rejection rates across effective price ratios, sample sizes, and task designs. Rejection rises with larger price changes and larger samples under the stated design assumptions. Error bars are 95 percent Wilson intervals; horizontal references mark the nominal 0.05 test size and 0.80 rejection rate.",
    "6": "Three-panel plot of pooled residual-variance ratios for prior-width contrasts 40 over 20, 60 over 20, and 60 over 40 across 21 shared numerosities. The equality line marks a ratio of one. Navy points show the observed estimates at each numerosity; rust curves show the model-implied width-scaling profile calibrated at numerosity 60 and displayed across the shared range. Panel subtitles summarize each contrast's median estimate and number of numerosity-specific estimates above one.",
}


def figure_alt_text(caption: str, number: str) -> str:
    """Return a concise plain-language description of the figure's main content."""
    text = FIGURE_ALT_TEXT.get(number, re.sub(r"\*\*|__|\*|_", "", caption))
    return tex_escape(re.sub(r"\s+", " ", text).strip())


def table_widths(rows: list[list[str]], columns: int) -> list[float]:
    lengths = []
    for col in range(columns):
        longest = max((len(re.sub(r"[*`]", "", row[col])) for row in rows if col < len(row)), default=8)
        column_values = [re.sub(r"[*`]", "", row[col]) for row in rows if col < len(row)]
        unbreakable = max((len(word) for value in column_values for word in re.findall(r"[A-Za-z0-9./%]+", value)), default=6)
        lengths.append(max(max(6.0, min(float(longest), 72.0)) ** 0.78, float(unbreakable + 1) ** 1.15))
    total = sum(lengths)
    return [0.94 * value / total for value in lengths]


def render_table(rows: list[list[str]], caption: str | None, refs: dict, number: int) -> str:
    if not rows:
        return ""
    columns = max(len(row) for row in rows)
    rows = [row + [""] * (columns - len(row)) for row in rows]
    widths = table_widths(rows, columns)
    spec = "@{}" + "".join(rf">{{\raggedright\arraybackslash}}p{{{w:.4f}\textwidth}}" for w in widths) + "@{}"
    label = f"tab:table{number}"
    result = [r"\begingroup\small\setlength{\tabcolsep}{1pt}\renewcommand{\arraystretch}{1.12}", rf"\begin{{longtable}}{{{spec}}}"]
    if caption:
        # The Markdown source includes its manuscript-facing table number;
        # LaTeX supplies the number, so keep only the descriptive caption.
        cap = re.sub(r"^\s*\*\*Table\s+S?\d+\.\*\*\s*", "", caption)
        cap = re.sub(r"^\s*Table\s+S?\d+\.\s*", "", cap)
        result.append(rf"\caption{{{markdown_inline(cap, refs)}}}\label{{{label}}}\\")
    headers = [markdown_inline(cell, refs) for cell in rows[0]]
    result.append(r"\toprule")
    result.append(" & ".join(r"\textbf{" + cell + "}" for cell in headers) + r" \\")
    result.extend([r"\midrule", r"\endfirsthead", r"\toprule", " & ".join(r"\textbf{" + cell + "}" for cell in headers) + r" \\", r"\midrule", r"\endhead"])
    for row in rows[1:]:
        result.append(" & ".join(markdown_inline(cell, refs) for cell in row) + r" \\")
    result.extend([r"\bottomrule", r"\end{longtable}", r"\endgroup"])
    return "\n".join(result)


def table_caption_tex(caption: str, refs: dict, number: int) -> str:
    """Format a Markdown table caption using the generated LaTeX table number."""
    cap = re.sub(r"^\s*\*\*Table\s+S?\d+\.\*\*\s*", "", caption)
    cap = re.sub(r"^\s*Table\s+S?\d+\.\s*", "", cap)
    return rf"\caption{{{markdown_inline(cap, refs)}}}\label{{tab:table{number}}}\\"


def attach_caption_to_table(table_tex: str, caption: str, refs: dict, number: int) -> str:
    """Insert a post-table Markdown caption into its preceding longtable."""
    begin = table_tex.find(r"\begin{longtable}")
    if begin < 0:
        raise ValueError("A table caption followed a block that is not a longtable.")
    line_end = table_tex.find("\n", begin)
    if line_end < 0:
        raise ValueError("Malformed longtable block: missing begin-line terminator.")
    caption_tex = table_caption_tex(caption, refs, number)
    return table_tex[:line_end + 1] + caption_tex + "\n" + table_tex[line_end + 1:]


def convert_body(body: str, refs: dict, supplement: bool) -> tuple[str, list[str]]:
    lines = body.splitlines()
    out: list[str] = []
    cited: list[str] = []
    table_number = 0
    figure_number = 0
    paragraph: list[str] = []
    pending_caption: str | None = None
    last_table_index: int | None = None
    last_table_number: int | None = None
    in_abstract = False

    for match in re.finditer(r"\[\[([^\]]+)\]\]", body):
        for part in match.group(1).split(";"):
            key = part.strip().lstrip("@").split("|")[0].strip()
            if key in refs and key not in cited:
                cited.append(key)

    def flush_paragraph() -> None:
        nonlocal paragraph
        text = " ".join(part.strip() for part in paragraph).strip()
        paragraph = []
        if text:
            out.append(markdown_inline(text, refs) + "\n\n")

    i = 0
    while i < len(lines):
        line = lines[i].strip()
        if not line:
            flush_paragraph()
            i += 1
            continue

        if line.startswith("**Table"):
            flush_paragraph()
            if last_table_index is not None and last_table_number is not None:
                out[last_table_index] = attach_caption_to_table(
                    out[last_table_index], line, refs, last_table_number
                )
                last_table_index = None
                last_table_number = None
            else:
                pending_caption = line
            i += 1
            continue

        # A caption printed after a table belongs to that immediately preceding
        # table only. Any intervening content ends that association.
        last_table_index = None
        last_table_number = None

        figure = re.match(r"^@@FIGURE\s*(\d+)\s*\|\s*([^|]+?)\s*\|\s*(.*?)@@$", line)
        if figure:
            flush_paragraph()
            pending_caption = None
            figure_number += 1
            num, image, caption = figure.groups()
            image = image.replace("\\", "/")
            vector_image = re.sub(r"\.png$", ".pdf", image, flags=re.IGNORECASE)
            if vector_image != image and Path(vector_image).is_file():
                image = vector_image
            out.append(r"\begin{figure}[H]\centering" + "\n")
            alt = figure_alt_text(caption, num)
            out.append(rf"\includegraphics[alt={{{alt}}},width=0.96\linewidth]{{{image}}}" + "\n")
            out.append(rf"\caption{{{markdown_inline(caption, refs)}}}\label{{fig:figure{num}}}" + "\n")
            out.append(r"\end{figure}" + "\n\n")
            i += 1
            continue

        if line.startswith("|") and i + 1 < len(lines) and lines[i + 1].strip().startswith("|"):
            flush_paragraph()
            raw_rows: list[list[str]] = []
            while i < len(lines) and lines[i].strip().startswith("|"):
                cells = table_cells(lines[i])
                if not all(re.fullmatch(r":?-{2,}:?", cell or "-") for cell in cells):
                    raw_rows.append(cells)
                i += 1
            table_number += 1
            out.append(render_table(raw_rows, pending_caption, refs, table_number) + "\n\n")
            last_table_index = len(out) - 1
            last_table_number = table_number
            pending_caption = None
            continue

        heading = re.match(r"^(#{1,4})\s+(.+)$", line)
        if heading:
            flush_paragraph()
            if pending_caption:
                out.append(markdown_inline(pending_caption, refs) + "\n\n")
                pending_caption = None
            level, title = len(heading.group(1)), heading.group(2)
            if level == 2 and title.lower() == "abstract":
                out.append(r"\begin{abstract}" + "\n")
                in_abstract = True
            else:
                if in_abstract:
                    out.append(r"\end{abstract}" + "\n")
                    in_abstract = False
                command = {1: "section", 2: "section", 3: "subsection", 4: "subsubsection"}[level]
                unnumbered_availability = level == 2 and title.strip().lower() in {"data and code availability", "competing interests", "funding", "author contributions"}
                unnumbered_supplement_heading = supplement and (
                    (level == 2 and re.match(r"S\d+\.", title))
                    or (level == 3 and re.match(r"S9\.\d+\.", title))
                )
                star = "*" if unnumbered_availability or unnumbered_supplement_heading else ""
                out.append(rf"\{command}{star}{{{markdown_inline(title, refs)}}}" + "\n\n")
            i += 1
            continue

        if line.startswith("- ") or re.match(r"^\d+\.\s+", line):
            flush_paragraph()
            if pending_caption:
                out.append(markdown_inline(pending_caption, refs) + "\n\n")
                pending_caption = None
            ordered = bool(re.match(r"^\d+\.\s+", line))
            env = "enumerate" if ordered else "itemize"
            out.append(rf"\begin{{{env}}}" + "\n")
            while i < len(lines):
                item = lines[i].strip()
                match = re.match(r"^\d+\.\s+(.*)$", item) if ordered else re.match(r"^-\s+(.*)$", item)
                if not match:
                    break
                out.append(r"\item " + markdown_inline(match.group(1), refs) + "\n")
                i += 1
            out.append(rf"\end{{{env}}}" + "\n\n")
            continue

        if line.startswith("> "):
            flush_paragraph()
            out.append(r"\begin{quote}" + "\n" + markdown_inline(line[2:], refs) + "\n" + r"\end{quote}" + "\n\n")
            i += 1
            continue

        paragraph.append(line)
        i += 1

    flush_paragraph()
    if in_abstract:
        out.append(r"\end{abstract}" + "\n")
    if pending_caption:
        out.append(markdown_inline(pending_caption, refs) + "\n\n")
    return "".join(out), cited


def document_preamble(meta: dict[str, str], supplement: bool) -> str:
    title = markdown_inline(meta.get("title", "Manuscript"), {})
    authors = author_tex(meta.get("authors", ""), meta.get("affiliations", ""))
    pdf_title = re.sub(r"\s+", " ", meta.get("title", "Manuscript")).strip()
    pdf_authors = re.sub(r"\^\{[^}]*\}", "", meta.get("authors", ""))
    pdf_authors = re.sub(r"\s+and\s+", "; ", pdf_authors).strip()
    preamble = r"""\ExplSyntaxOn\cs_undefine:N \qed\ExplSyntaxOff
\documentclass[10pt]{article}
\usepackage[margin=1in]{geometry}
\usepackage[round,authoryear]{natbib}
\makeatletter
\newcommand{\biboptions}[1]{}
\setcounter{secnumdepth}{0}
\gdef\fm@address{}
\newcommand{\address}[1]{\gdef\fm@address{#1}}
\newcommand{\fm@printtitle}{\begin{center}{\Large\bfseries\@title\par}\vspace{0.8em}{\@author\par}\vspace{0.4em}{\small\fm@address\par}\end{center}\vspace{0.6em}\global\let\fm@printtitle\relax}
\newenvironment{frontmatter}{}{\fm@printtitle}
\let\fm@origabstract\abstract \let\fm@origendabstract\endabstract
\renewenvironment{abstract}{\fm@printtitle\fm@origabstract}{\fm@origendabstract}
\newenvironment{keyword}{\par\noindent\textbf{Keywords:} }{\par\medskip}
\makeatother
\usepackage{fontspec}
\usepackage{microtype}
\usepackage{amsmath,amssymb}
\usepackage{graphicx}
\usepackage{float}
\usepackage{longtable}
\usepackage{array,ragged2e,booktabs}
\usepackage{caption}
\usepackage{xurl}
\usepackage[hidelinks]{hyperref}
HYPERREF_METADATA
\usepackage{lineno}
\biboptions{round,authoryear}
\providecommand{\textsubscript}[1]{\ensuremath{_{\mathrm{#1}}}}
\providecommand{\textsuperscript}[1]{\ensuremath{^{\mathrm{#1}}}}
\captionsetup{font=small,labelfont=bf,justification=justified,singlelinecheck=false}
\renewcommand{\arraystretch}{1.12}
\setlength{\tabcolsep}{5pt}
\setlength{\parindent}{1.25em}
\setlength{\parskip}{0.35em}
\setlength{\emergencystretch}{2em}
\clubpenalty=10000
\widowpenalty=10000
\pagestyle{plain}
\begin{document}
\linenumbers
"""
    metadata = rf"\hypersetup{{pdftitle={{{tex_escape(pdf_title)}}},pdfauthor={{{tex_escape(pdf_authors)}}}}}"
    preamble = preamble.replace("HYPERREF_METADATA", metadata)
    top = r"\begin{frontmatter}" + "\n" + rf"\title{{{title}}}" + "\n" + authors + "\n"
    return preamble + top


def main() -> None:
    parser = argparse.ArgumentParser()
    parser.add_argument("markdown", type=Path)
    parser.add_argument("references", type=Path)
    parser.add_argument("output", type=Path)
    parser.add_argument("--supplement", action="store_true")
    args = parser.parse_args()

    source = args.markdown.read_text(encoding="utf-8")
    meta, body = parse_front_matter(source)
    refs = json.loads(args.references.read_text(encoding="utf-8"))
    converted, cited = convert_body(body, refs, args.supplement)
    if "\\begin{abstract}" in converted and "\\end{abstract}" not in converted:
        converted += "\\end{abstract}\n"
    if args.supplement:
        converted = "\\end{frontmatter}\n\\renewcommand{\\thetable}{S\\arabic{table}}\n" + converted
    else:
        abstract_end = "\\end{abstract}"
        if abstract_end not in converted:
            raise ValueError("The main manuscript must contain an abstract within front matter.")
        keyword_text = meta.get("keywords", "").replace(";", ",").strip()
        keyword_block = ""
        if keyword_text:
            keyword_block = "\\begin{keyword}\n" + markdown_inline(keyword_text, refs) + "\n\\end{keyword}\n"
        converted = converted.replace(
            abstract_end,
            abstract_end + "\n" + keyword_block + "\\end{frontmatter}",
            1,
        )
    bibliography = bibliography_tex(cited, refs) if cited else ""
    args.output.parent.mkdir(parents=True, exist_ok=True)
    args.output.write_text(document_preamble(meta, args.supplement) + converted + bibliography + "\n\\end{document}\n", encoding="utf-8")
    figures = converted.count("\\begin{figure}")
    tables = converted.count("\\begin{longtable}")
    print(f"Wrote {args.output.name}: {len(cited)} cited references, {figures} figures, {tables} tables")


if __name__ == "__main__":
    main()
```
