## Supplementary material for "The marginal brain": Replication package: mathfix.py

```
"""Convert the plain-text mathematics of the filled manuscript into ^{...} and _{...} markup, which the extended
Word builder renders as true superscripts and subscripts. Citation keys, code spans and figure paths are protected."""
import re, sys

SUB = {'₀': '0', '₁': '1', '₂': '2', 'ₐ': 'a', 'ₒ': 'o'}


def convert(seg):
    for u, c in SUB.items():
        seg = seg.replace(u, '_{' + c + '}')
    seg = re.sub(r'(?<=\b[nEL])\*(?!\*)', '^{∗}', seg)            # optimum marks n*, E*, L*
    out, i = [], 0
    while i < len(seg):                                            # ^( ... ) with balanced parentheses
        if seg.startswith('^(', i):
            depth, j = 0, i + 1
            while j < len(seg):
                depth += {'(': 1, ')': -1}.get(seg[j], 0)
                if depth == 0:
                    break
                j += 1
            out.append('^{' + seg[i + 2:j] + '}')
            i = j + 1
        else:
            out.append(seg[i]); i += 1
    seg = ''.join(out)
    seg = re.sub(r'\^(?!\{)([½¼¾]|[A-Za-zκργ0-9]+)', r'^{\1}', seg)   # single-token superscripts
    seg = re.sub(r'(?<=\b[A-Z])_(?!\{)([A-Za-z0-9])(?![A-Za-z0-9_])', r'_{\1}', seg)  # Z_a and similar subscripts
    return seg


def process_line(line):
    if line.startswith('@@FIGURE'):
        head, path, cap = line.split(' | ', 2)
        return ' | '.join([head, path, process_line_text(cap)])
    if line.startswith(('title:', 'authors:', 'affiliations:', 'correspondence:', 'keywords:', 'style:', 'budget:')):
        return line
    return process_line_text(line)


def process_line_text(text):
    parts = re.split(r'(\[\[[^\]]+\]\]|`[^`]*`|\[PLS:[^\]]*\])', text)
    return ''.join(p if (p.startswith(('`', '[[', '[PLS:'))) else convert(p) for p in parts)


src = open(sys.argv[1], encoding='utf-8').read().split('\n')
open(sys.argv[2], 'w', encoding='utf-8').write('\n'.join(process_line(l) for l in src))
```
