## Supplementary material for "The marginal brain": Replication package: fig6_empirical.py

```
"""Render Figure 6 from checked empirical and model result files."""
import json
import math
from statistics import median as median_value
from pathlib import Path

from PIL import Image, ImageDraw, ImageFont

HERE = Path(__file__).resolve().parent
MODEL = HERE.parent / "model"
FIGURES = HERE / "figures"
FIGURES.mkdir(exist_ok=True)
results = json.loads((MODEL / "results.json").read_text(encoding="utf-8"))
predicted = json.loads((MODEL / "model_predicted_ratios.json").read_text(encoding="utf-8"))
contrasts = results["empirical_reanalysis"]["per_numerosity_cluster_bootstrap"]["contrasts"]

page_w, page_h = 545.248125, 309.879375  # preserve the established figure footprint
dpi = 300
scale = 2 * dpi / 72  # render at 600 dpi, then downsample with antialiasing
size = (round(page_w * dpi / 72), round(page_h * dpi / 72))
image = Image.new("RGB", (round(page_w * scale), round(page_h * scale)), "white")
draw = ImageDraw.Draw(image)
font_roots = [Path("C:/Windows/Fonts"), Path("/usr/share/fonts/truetype/msttcorefonts"),
              Path("/usr/share/fonts/truetype/dejavu")]
regular_names = ("times.ttf", "Times_New_Roman.ttf", "DejaVuSerif.ttf")
bold_names = ("timesbd.ttf", "Times_New_Roman_Bold.ttf", "DejaVuSerif-Bold.ttf")


def font(size_pt, bold=False):
    for root in font_roots:
        for name in (bold_names if bold else regular_names):
            candidate = root / name
            if candidate.exists():
                return ImageFont.truetype(str(candidate), max(1, round(size_pt * scale)))
    return ImageFont.load_default()


def xy(x_pt, y_pt):
    return round(x_pt * scale), round((page_h - y_pt) * scale)


def line(x1, y1, x2, y2, color, width_pt=0.7, dash_pt=None):
    x1p, y1p = xy(x1, y1)
    x2p, y2p = xy(x2, y2)
    width = max(1, round(width_pt * scale))
    if not dash_pt:
        draw.line((x1p, y1p, x2p, y2p), fill=color, width=width)
        return
    length = math.hypot(x2p - x1p, y2p - y1p)
    dash, gap = dash_pt[0] * scale, dash_pt[1] * scale
    pos = 0.0
    while pos < length:
        end = min(pos + dash, length)
        a, b = pos / length, end / length
        draw.line((round(x1p + (x2p - x1p) * a), round(y1p + (y2p - y1p) * a),
                   round(x1p + (x2p - x1p) * b), round(y1p + (y2p - y1p) * b)),
                  fill=color, width=width)
        pos += dash + gap


def text(x, y, label, face, color="#111111", anchor="ls"):
    draw.text(xy(x, y), label, font=face, fill=color, anchor=anchor)


# A fixed, data-centered log-ratio scale keeps all observed estimates and
# model profiles legible without allowing extreme bootstrap endpoints to
# compress the point-estimate patterns.
xmin, xmax = 0.5, 8.0
xticks = [0.5, 1.0, 2.0, 4.0, 8.0]
width_pairs = [(20, 40), (20, 60), (40, 60)]
titles = ["A  Width 40 / 20", "B  Width 60 / 20", "C  Width 60 / 40"]
lefts, panel_w = [68.0, 218.5, 369.0], 135.0
y_top, y_bottom = 266.0, 78.0
ratio_log_span = math.log(xmax / xmin)
blue, grey, red = "#17365D", "#657482", "#B04A3A"
summary_by_pair = {}
for w1, w2 in width_pairs:
    vals = sorted(float(row["variance_ratio_width_2_over_width_1"])
                  for row in contrasts if row["width_1"] == w1 and row["width_2"] == w2)
    summary_by_pair[(w1, w2)] = (vals[len(vals) // 2], sum(v > 1 for v in vals), len(vals))

for panel, ((w1, w2), title, x0) in enumerate(zip(width_pairs, titles, lefts)):
    def xmap(value):
        return x0 + panel_w * math.log(float(value) / xmin) / ratio_log_span

    def ymap(value):
        return y_top - (float(value) - 49.5) / 21.0 * (y_top - y_bottom)

    rows = sorted((row for row in contrasts if row["width_1"] == w1 and row["width_2"] == w2),
                  key=lambda row: row["numerosity"])
    for tick in xticks:
        x = xmap(tick)
        line(x, y_bottom, x, y_top, "#e8e8e8", 0.5)
    line(xmap(1.0), y_bottom, xmap(1.0), y_top, "#595959", 0.9, (3.0, 2.0))

    # Show point estimates, the model profile and the parity reference. The
    # figure's visual encoding remains descriptive; bootstrap intervals are
    # reported separately as procedure outputs in the Supplement.
    pred_values = predicted["per_numerosity"][f"{w1}_{w2}"]
    predicted_median = float(median_value(pred_values))
    pred_points = [(xmap(v), ymap(n)) for v, n in zip(pred_values, predicted["numerosities"])]
    for first, second in zip(pred_points, pred_points[1:]):
        line(*first, *second, red, 1.35)

    for row in rows:
        x, y = xmap(row["variance_ratio_width_2_over_width_1"]), ymap(row["numerosity"])
        r = 2.35 * scale
        box = (round(x * scale - r), round((page_h - y) * scale - r),
               round(x * scale + r), round((page_h - y) * scale + r))
        draw.ellipse(box, fill=blue, outline=blue, width=max(1, round(0.6 * scale)))

    line(x0, y_bottom, x0 + panel_w, y_bottom, "#111111", 0.8)
    line(x0, y_bottom, x0, y_top, "#111111", 0.8)
    median, above, total = summary_by_pair[(w1, w2)]
    text(x0 + panel_w / 2, 299.0, title, font(10.2, bold=True), anchor="ms")
    text(x0 + panel_w / 2, 283.0,
         f"Observed median {median:.2f}  |  model {predicted_median:.2f}",
         font(8.2), color=grey, anchor="ms")
    text(x0 + panel_w / 2, 272.0, f"Point estimates above 1: {above}/{total}",
         font(8.0), color=grey, anchor="ms")
    for tick in xticks:
        x = xmap(tick)
        line(x, y_bottom, x, y_bottom - 2.5, "#111111", 0.7)
        text(x, 59.0, f"{tick:g}", font(8.8), anchor="ms")
    if panel == 0:
        for tick in (50, 55, 60, 65, 70):
            y = ymap(tick)
            line(x0, y, x0 - 2.5, y, "#111111", 0.7)
            text(x0 - 7.0, y, str(tick), font(8.8), anchor="rs")

text(page_w / 2, 31.5, "pooled residual-variance ratio (wider / narrower)", font(10.0), anchor="ms")

# Rotate the shared y-axis label as a single typographic object.
y_label = "presented numerosity"
y_font = font(10.0)
bbox = y_font.getbbox(y_label)
label_image = Image.new("RGBA", (bbox[2] - bbox[0] + 8, bbox[3] - bbox[1] + 8), (255, 255, 255, 0))
ImageDraw.Draw(label_image).text((4 - bbox[0], 4 - bbox[1]), y_label, font=y_font, fill="#111111")
label_image = label_image.rotate(90, expand=True, resample=Image.Resampling.BICUBIC)
center = xy(25.0, (y_top + y_bottom) / 2)
image.paste(label_image, (center[0] - label_image.width // 2, center[1] - label_image.height // 2), label_image)

legend = [("point", "observed pooled ratio"), ("line", "model profile (calibrated at numerosity 60)"), ("reference", "parity (ratio = 1)")]
legend_font = font(8.6)
item_widths = []
for kind, label in legend:
    symbol_w = 17.0
    item_widths.append(symbol_w + 5.0 + legend_font.getlength(label) / scale)
total_w = sum(item_widths) + 12.0 * (len(legend) - 1)
cursor, baseline = (page_w - total_w) / 2, 10.5
for (kind, label), item_w in zip(legend, item_widths):
    cy = baseline + 2.7
    if kind == "point":
        cx, cyp = xy(cursor + 7.5, cy)
        radius = 2.2 * scale
        draw.ellipse((round(cx-radius), round(cyp-radius), round(cx+radius), round(cyp+radius)),
                     fill=blue, outline=blue)
    elif kind == "line":
        line(cursor, cy, cursor + 15.0, cy, red, 1.35)
    else:
        line(cursor, cy, cursor + 15.0, cy, grey, 0.9, (3.0, 2.0))
    label_x = cursor + 20.0
    text(label_x, baseline, label, legend_font)
    cursor += item_w + 12.0

image = image.resize(size, Image.Resampling.LANCZOS)
png_path, pdf_path = FIGURES / "fig6_empirical.png", FIGURES / "fig6_empirical.pdf"
image.save(png_path, dpi=(dpi, dpi), optimize=True)
image.save(pdf_path, "PDF", resolution=dpi)
print(f"Figure 6 rendered over [{xmin:g}, {xmax:g}] with {dpi}-dpi artwork.")
```
