## Supplementary material for "The marginal brain": Replication package: fig_common_factor.py

```
"""Figure 2: the common factor (Proposition 2), computed from the model package's closed forms."""
import sys, json
sys.path.insert(0, '../model')
import numpy as np
import matplotlib
matplotlib.use('Agg')
import matplotlib.pyplot as plt
from matplotlib.ticker import FixedLocator, NullLocator, FormatStrFormatter
from spt import model as Mo
plt.rcParams.update({'font.size': 8, 'font.family': 'DejaVu Serif', 'mathtext.fontset': 'stix', 'savefig.dpi': 300,
                     'axes.spines.top': False, 'axes.spines.right': False,
                     'axes.labelsize': 8, 'xtick.labelsize': 7, 'ytick.labelsize': 7})
ws = np.linspace(20, 80, 60)
fig, ax = plt.subplots(figsize=(3.6, 2.8))
for a, name, col in [(1, 'Estimation', '#17365D'), (2, 'Discrimination', '#C65A46')]:
    ref = Mo.uniform_sigma(20.0, 1.0, 1.0, a)
    ax.loglog(ws, Mo.uniform_sigma(ws, 1.0, 1.0, a) / ref, color=col, lw=1.5, label=f'{name}; λ ×1')
    ax.loglog(ws, Mo.uniform_sigma(ws, 1.0, 2.0, a) / ref, color=col, lw=1.5, ls='--', label=f'{name}; λ ×2')
ax.xaxis.set_major_locator(FixedLocator([20, 30, 40, 60, 80])); ax.xaxis.set_minor_locator(NullLocator())
ax.xaxis.set_major_formatter(FormatStrFormatter('%g'))
ax.yaxis.set_major_locator(FixedLocator([1.0, 1.5, 2.0, 3.0])); ax.yaxis.set_minor_locator(NullLocator())
ax.yaxis.set_major_formatter(FormatStrFormatter('%g'))
ax.set_xlabel('prior width w')
ax.set_ylabel('Imprecision ratio')
ax.legend(fontsize=6.3, frameon=False, loc='upper center', bbox_to_anchor=(0.5, -0.22), ncol=2,
          handlelength=2.0, columnspacing=1.2)
fig.tight_layout(rect=(0, 0.09, 1, 1))
fig.savefig('figures/fig2_common_factor.png', bbox_inches='tight'); fig.savefig('figures/fig2_common_factor.pdf', bbox_inches='tight')
r = json.load(open('../model/results.json'))
print('factor', 2**0.25, r['headline']['common_factor_price_x2'])
```
