## Supplementary material for "The marginal brain": Replication package: fig_rest.py

```
"""Figures 3 to 5 for the technical paper, from the model package and its results file."""
import sys, json, shutil
sys.path.insert(0, '../model')
import numpy as np
import matplotlib
matplotlib.use('Agg')
import matplotlib.pyplot as plt
from spt import model as Mo
plt.rcParams.update({'font.size': 9, 'font.family': 'DejaVu Serif', 'mathtext.fontset': 'stix', 'savefig.dpi': 300,
                     'axes.spines.top': False, 'axes.spines.right': False,
                     'axes.labelsize': 9, 'xtick.labelsize': 8.3, 'ytick.labelsize': 8.3})
res = json.load(open('../model/results.json'))
b = res['config']['budget']['baseline']
# Figure 3: two compact, logically linked panels.  The artwork is generated
# at final single-column width, with short in-panel labels to preserve
# legibility without a long vertical axis title crossing the panel boundary.
fig, ax = plt.subplots(1, 2, figsize=(4.42, 2.72), sharey=False)
M = np.array(res['budget']['M_grid'])
for rho, col in [(0.5, '#2A7F7F'), (1.0, '#17365D'), (2.0, '#C65A46')]:
    ax[0].loglog(M, res['budget']['curves'][f'rho{rho}_c100.0'], color=col, lw=1.35, label=rf'$\rho={rho:g}$')
ax[0].loglog(M, res['budget']['curves']['rho1.0_chard'], color='#17365D', lw=1.2, ls=':', label='cap')
ax[0].set_ylabel(r'$\mu$', fontsize=7.2, labelpad=1)
ax[0].set_xlabel(r'margin $M$', fontsize=7.2, labelpad=2)
ax[0].legend(fontsize=7.4, frameon=False, loc='upper center', bbox_to_anchor=(0.5, -0.34),
             ncol=2, handlelength=0.9, columnspacing=0.45, labelspacing=0.1,
             borderaxespad=0.0, handletextpad=0.35)
ax[0].set_title('A   Price vs. margin', loc='left', fontsize=8.5, weight='bold', pad=5)
ax[0].grid(which='major', color='0.91', lw=0.45)
ax[0].tick_params(axis='both', labelsize=6.6, length=2.0, pad=1.8)
Ms = np.linspace(0.015, 0.15, 60)
gross, net = [], []
for m in Ms:
    m1 = Mo.budget_mu(m, b['c'], b['v'], b['rho'], b['e'], b['A'], b['k'])
    m0 = Mo.budget_mu(m, b['c'], b['v'], b['rho'], b['e'], 0.0, b['k'])
    gross.append(np.sqrt(b['e']) * b['A'] / np.sqrt(m1)); net.append((m1 - m0) / b['c'])
ax[1].plot(Ms, np.array(gross) * 1e3, color='#17365D', lw=1.4, label='$E$')
ax[1].plot(Ms, np.array(net) * 1e3, color='#C65A46', lw=1.4, label='$\u0394 B$')
ax[1].set_xlabel(r'margin $M$', fontsize=7.2, labelpad=2)
ax[1].set_title(r'B   Energy ($10^{-3}\bar{B}$)', loc='left', fontsize=8.2, weight='bold', pad=5)
ax[1].legend(fontsize=7.4, frameon=False, loc='upper center', bbox_to_anchor=(0.5, -0.34),
             ncol=2, handlelength=0.9, columnspacing=0.7, labelspacing=0.1,
             borderaxespad=0.0, handletextpad=0.35)
ax[1].set_ylim(0, None)
ax[1].set_yticks([0, 1, 2, 3])
ax[1].grid(axis='y', which='major', color='0.91', lw=0.45)
ax[1].tick_params(axis='both', labelsize=6.6, length=2.0, pad=1.8)
ax[1].set_xticks([0.05, 0.10, 0.15])
ax[1].set_xlim(0.012, 0.155)
fig.subplots_adjust(left=0.13, right=0.99, top=0.83, bottom=0.29, wspace=0.58)
fig.savefig('figures/fig3_budget.png', bbox_inches='tight', pad_inches=0.03)
fig.savefig('figures/fig3_budget.pdf', bbox_inches='tight', pad_inches=0.03)
plt.close(fig)
# Figure 4: the sign test
fig, ax = plt.subplots(figsize=(6.3, 3.1))
t = np.linspace(-0.45, 0.45, 2)
ax.plot(t, -0.5 * t, color='#17365D', lw=1.6, label='budget price change (slope −½)')
ax.plot(t, 0.5 * t, color='#C65A46', lw=1.6, label='technology change (slope +½)')
cases = res['signatures']['cases']
marks = {'price_rise_supply_tightening (mu x2)': ('o', 'p ×2'), 'capacity_loss (K x0.8)': ('s', 'capacity ×0.8'),
         'cheaper_signals (e x0.71)': ('^', 'energy per signal ×0.71'),
         'scarcity_with_technology_change (mu x2, e x0.71, K x0.8)': ('D', 'p ×2 with both technology changes'),
         'stakes_up (beta x1.5, price-like)': ('v', 'stakes ×1.5')}
for k, (m, lab) in marks.items():
    c = cases[k]; ax.plot(c['dlog_E'], c['dlog_sigma'], m, color='#2A7F7F', ms=4.5, mec='#17365D', mew=0.45, label=lab)
ax.axhline(0, color='0.75', lw=0.5); ax.axvline(0, color='0.75', lw=0.5)
ax.set_xlabel('change in log task energy'); ax.set_ylabel('change in log imprecision')
ax.set_xticks([-0.4, -0.2, 0.0, 0.2, 0.4]); ax.set_xlim(-0.5, 0.5)
ax.legend(fontsize=8.5, frameon=False, loc='upper center', bbox_to_anchor=(0.5, -0.25), ncol=2,
          handlelength=1.8, columnspacing=1.0, labelspacing=0.4)
fig.subplots_adjust(left=0.14, right=0.97, bottom=0.35, top=0.93)
fig.savefig('figures/fig4_sign_test.png', bbox_inches='tight'); fig.savefig('figures/fig4_sign_test.pdf', bbox_inches='tight'); plt.close(fig)
# Figure 5: power, from the model package
shutil.copy('../model/figures/fig2_power.png', 'figures/fig5_power.png')
shutil.copy('../model/figures/fig2_power.pdf', 'figures/fig5_power.pdf')
print('net/gross at M=0.05:', round(net[np.argmin(abs(Ms-0.05))]/gross[np.argmin(abs(Ms-0.05))], 3))
```
