## Supplementary material for "The marginal brain": Replication package: fig_schematic.py

```
"""Figure 1: compact, three-stage map of the shared-budget model."""
import matplotlib
matplotlib.use('Agg')
import matplotlib.pyplot as plt
from matplotlib.patches import FancyBboxPatch

plt.rcParams.update({
    'font.family': 'DejaVu Serif',
    'font.size': 9.2,
    'mathtext.fontset': 'stix',
    'savefig.dpi': 300,
    'pdf.fonttype': 42,
})

# The source canvas is close to the manuscript's single-column width.  This
# keeps labels legible after LaTeX includes the PDF at 0.96\linewidth.
fig, ax = plt.subplots(figsize=(4.42, 3.05))
ax.set_xlim(0, 4.42)
ax.set_ylim(0, 3.05)
ax.axis('off')

navy = '#17365D'
rows = [
    (2.14, '#E8F0F4', 'A', 'Shared energy budget',
     r'$M=\bar{B}-C,\qquad E=en,\qquad d+M=E+E_o$'),
    (1.18, '#F5F0E3', 'B', 'Equilibrium shadow price',
     r'$\mu=R\prime(d)=V\prime(E_o)$'),
    (0.22, '#E8EEF5', 'C', 'Precision response',
     r'$\lambda=e\mu/\beta,\qquad \sigma\propto\lambda^{1/4}$'),
]
box_x, box_w, box_h = 0.28, 3.86, 0.72
for y, fill, letter, title, equation in rows:
    ax.add_patch(FancyBboxPatch(
        (box_x, y), box_w, box_h,
        boxstyle='round,pad=0.025,rounding_size=0.06',
        facecolor=fill, edgecolor=navy, linewidth=0.85,
    ))
    ax.text(box_x + 0.18, y + box_h - 0.14, letter,
            ha='left', va='top', color=navy, fontsize=9.4, weight='bold')
    ax.text(box_x + 0.47, y + box_h - 0.14, title,
            ha='left', va='top', color=navy, fontsize=9.0, weight='bold')
    ax.text(box_x + box_w / 2, y + 0.20, equation,
            ha='center', va='center', color='#202833', fontsize=10.1)

# A single, vertical path makes the dependency legible and keeps the
# transmission pathway uncluttered.
for y0, y1 in [(2.14, 1.90), (1.18, 0.94)]:
    ax.annotate('', xy=(2.21, y1), xytext=(2.21, y0),
                arrowprops=dict(arrowstyle='-|>', color=navy,
                                linewidth=1.0, mutation_scale=10))

fig.subplots_adjust(left=0.015, right=0.985, top=0.985, bottom=0.015)
fig.savefig('figures/fig1_schematic.png', bbox_inches='tight', pad_inches=0.02)
fig.savefig('figures/fig1_schematic.pdf', bbox_inches='tight', pad_inches=0.02)
plt.close(fig)
print('Figure 1 rendered: budget to shadow price to precision response')
```
