## Supplementary material for "The marginal brain": Replication package: fig1_schematic.pdf

**A** **Shared energy budget**

$$M = \bar{B} - C, \qquad E = en, \qquad d + M = E + E_o$$

**B** **Equilibrium shadow price**

$$\mu = R'(d) = V'(E_o)$$

**C** **Precision response**

$$\lambda = e\mu/\beta, \qquad \sigma \propto \lambda^{1/4}$$
