## Supplementary material for "The marginal brain": Replication package: Manuscript.pdf

### Model

#### Endogenous precision

We build on the endogenous-precision model developed for the perception of numerical magnitudes (Prat-Carrabin and Woodford, 2026). A magnitude $\mathrm{x}$, drawn from a prior density $\pi$, is encoded with effective signal mass $\mathrm{n}$, a continuous activity measure in the optimization rather than a literal count of individual spikes. Each independent signal carries Fisher information $\mathrm{I}_1(\mathrm{x})$, and the allocation of precision across magnitudes is bounded by $\int \mathrm{I}_1(\mathrm{x})^{\frac{1}{2}}\, \mathrm{dx} \leq \mathrm{K}$, a constraint used in efficient-coding models of unidimensional channels (Prat-Carrabin and Woodford write its right-hand side as $\sqrt{K}$, so K here corresponds to their

$\sqrt{K}$). Because Fisher information adds across independent signals, total precision is $I(x) = n\, I_1(x)$; in the high-precision regime, an efficient decoder attains imprecision $\sigma(x) = I(x)^{-\frac{1}{2}}$. The task loss is approximated by $L_1[I] = \int \pi(x)/I(x)\, dx$ for estimation and $L_2[I] = \int \pi(x)^2/I(x)\, dx$ for discrimination. The observer chooses $I_1$ and the continuous effective activity n to minimize $L_a + \lambda n$, with a = 1 for estimation and a = 2 for discrimination. Under a uniform prior of width w, imprecision scales as $w^{\frac{1}{2}}$ in estimation and $w^{\frac{3}{4}}$ in discrimination, as supported by the cited numerosity data (Prat-Carrabin and Woodford, 2026). In this model, $\lambda$ is a behavioral price held fixed across the observer's states; the next section endogenizes its state dependence through the energy budget.

#### The brain's energy budget

We embed this choice in a shared energy budget. Total use B comprises committed use C, gross task energy $E = e\, n$, and energy $E_o$ allocated to competing uses. For $v > 0$ and $\rho > 0$, the latter's value is $V(E_o) = v({E_o}^{1-\rho} - 1)/(1-\rho)$ for $\rho \neq 1$ and $V(E_o) = v \log E_o$ for $\rho = 1$; both give $V'(E_o) = v{E_o}^{-\rho}$, including the configured baseline $\rho = 1$, and make V increasing and concave. Committed use includes maintenance and intrinsic activity, which together make up most resting consumption (Jamadar et al., 2025). Physiological theory and evidence motivate possible supply constraints through vascular architecture and oxygen-delivery homeostasis (Herculano-Houzel and Rothman, 2022), conserved energy use per neuron across brain sizes (Herculano-Houzel, 2011), and observed redistribution of metabolic activity across cortical regions during attention (Bruckmaier et al., 2020). Together these observations motivate the common-budget structure formalized here. We represent it with a set point $\bar{B}$ and a regulatory cost whose marginal value is $R'(d) = c\, d^k$, where $d = B - \bar{B}$ is use above the set point, $c > 0$ measures regulatory stiffness and $k > 0$ its convexity; a hard cap is the limit as c grows without bound. Local neurovascular coupling can increase supply during activation even when total use remains near rest (Jamadar et al., 2025). Accordingly, the set point may refer to the whole brain or to a region with bounded supply. The reallocation margin $M = \bar{B} - C$ is the energy available to the task and competing uses before regulation binds; changes in supply enter through this margin.

#### Equilibrium and the price of precision

The brain minimizes task loss $L_a$ plus regulatory cost R, less the value V of competing uses. At an interior optimum, the marginal regulatory cost equals the marginal value in competing uses: $\mu = R'(B - \bar{B}) = V'(E_o)$. This defines $\mu$, the shadow price of energy. When task stakes scale loss by $\beta$, define $p = \mu/\beta$ as the budget price per unit loss and $\lambda = ep = e\mu/\beta$ as the effective price per signal; under $\beta = 1$, these reduce to $p = \mu$ and $\lambda = e\mu$. The ratio of $\lambda$ equals the ratio of p only when e is fixed, and the ratio of p equals that of $\mu$ only when $\beta$ is fixed. The model therefore separates energy per signal from the budget's marginal value and makes clear why cross-state price comparisons require stable stakes. Neurophysiological work estimates energy costs of signaling (Laughlin et al., 1998), and neural coding may adapt under scarcity at some cost to precision (Padamsey et al., 2022); our model isolates the additional state-dependent price transmitted through $\mu$. Gross encoding energy E differs from net increment $\Delta B$, the change in total use after competing uses reallocate. Whole-brain imaging reports aggregate task-evoked changes of the latter kind (Jamadar et al., 2025).

Figure 1 gives a visual map of the model's transmission chain from the shared-budget constraint through the shadow price to precision. The three stages make the relationships and equilibrium feedback visible at a glance.

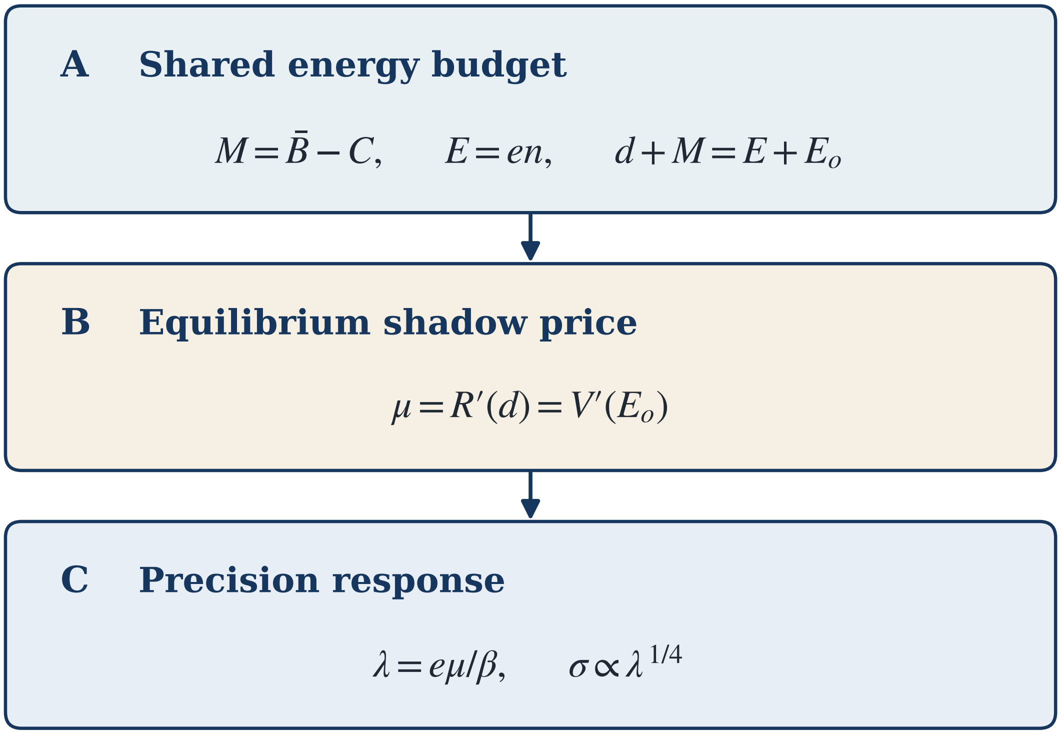


**Figure 1:** The model's three linked stages. Panel A states the shared-budget identities $\mathrm{M} = \bar{B} - \mathrm{C}$ and $\mathrm{d} + \mathrm{M} = \mathrm{E} + \mathrm{E_o}$, with gross task energy $\mathrm{E} = \mathrm{en}$. Panel B shows the equilibrium condition $\mu = \mathrm{R'(d)} = \mathrm{V'(E_o)}$, equating the marginal regulatory cost and value of competing uses. Panel C maps the shadow price into the per-signal price $\lambda = \mathrm{e}\mu/\beta$ and the imprecision response $\sigma \propto \lambda^{1/4}$, at fixed task and coding parameters. The arrows trace price transmission from the budget equilibrium to encoding; the budget is solved jointly, so task demand also feeds back into the equilibrium price.

**Lemma 1 (separation).** *Let $\pi$ be a normalized density on support $\mathcal{X}$, with $0 < Z_a = \int_{\mathcal{X}} \pi(x)^{a/3}\,dx < \infty$, and let $K > 0$ and $n > 0$. Under admissible information allocations satisfying $\int_{\mathcal{X}} I_1(x)^{1/2}\,dx \leq K$, the optimum satisfies $I_1(x)^{1/2} \propto \pi(x)^{a/3}$, and the expected loss after allocation is $Z_a^3/(nK^2)$.*

**Proposition 1 (baseline).** *For $\lambda > 0$ and the interior continuous optimum over $n > 0$, the optimal number of signals is $n^* = Z_a^{3/2}/(K\lambda^{1/2})$ and imprecision is $\sigma(x) = Z_a^{1/4}\lambda^{1/4}K^{-1/2}\pi(x)^{-a/3}$; under a uniform prior of width $w$, holding capacity $K$ and effective signal price $\lambda$ fixed, $\sigma \propto w^{1/2}$ in estimation ($a = 1$) and $\sigma \propto w^{3/4}$ in discrimination ($a = 2$).*

**Proposition 2 (common factor).** *For a fixed task specification (prior, loss, and decoder), the elasticity of imprecision with respect to the effective per-signal price $\lambda = e\mu/\beta$ equals $1/4$ at every magnitude and capacity. If the energy per signal $e$, the capacity $K$, the task specification, and the stakes $\beta$ are unchanged across states, a change that moves the shadow price from $\mu_0$ to $\mu_1$ multiplies imprecision by $(\mu_1/\mu_0)^{1/4}$ in every task that draws on the same budget and leaves the exponents in $w$ unchanged.*

The fourth-root law is the source of the model's main behavioral prediction. Because every task that draws on the same budget faces the same shadow price, a doubling of the price raises imprecision by a factor of 1.19 in every task and at every magnitude, while the prior and the task continue to determine how imprecision is distributed. In the coordinates of Figure 2, the price shifts the curves of estimation and discrimination upward by the same distance and leaves their slopes with prior width, one half and three quarters, unchanged. The prediction is therefore a restriction across tasks: within a change of state that leaves coding technology and stakes unchanged, estimation and discrimination must show the same state factor, which a model with a price fitted separately to each task does not imply. The law rests only on the additivity of Fisher information and on a cost that is linear in the number of signals, so that a different form of the efficient-coding bound alters the slopes with prior width without altering the law, a convex cost of signals preserves its sign, and integer counts make the response lumpy only when few signals are used (Supplementary Section S9.1).

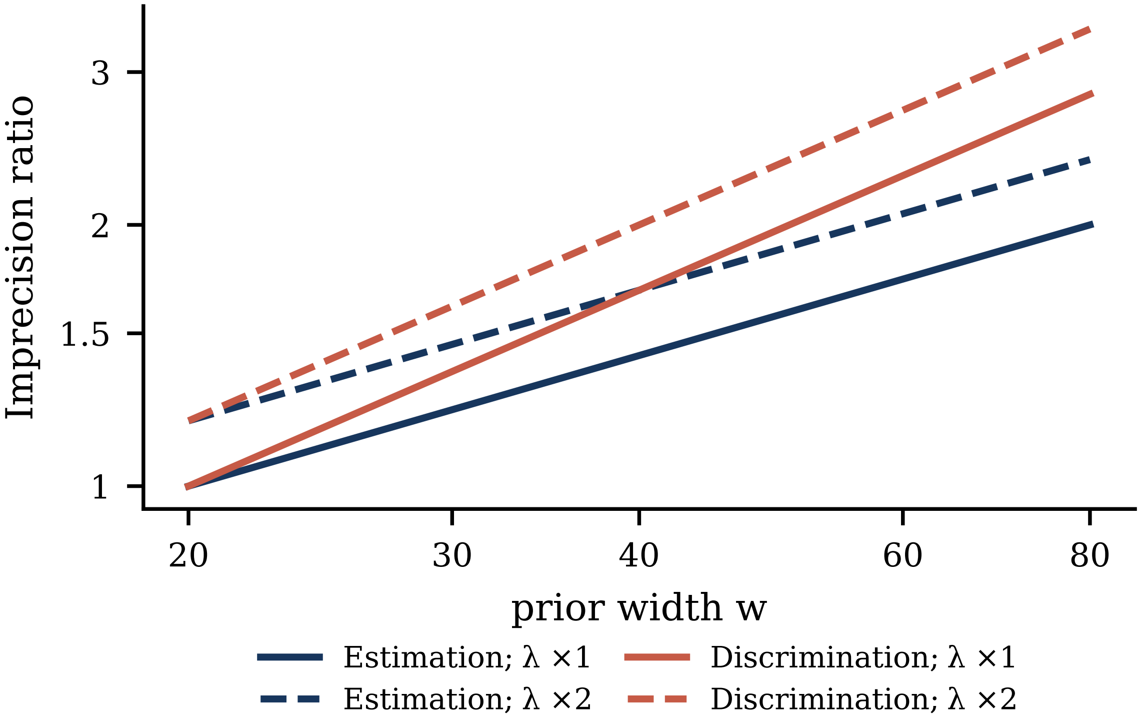


**Figure 2:** Imprecision as a function of prior width in estimation and in discrimination, at baseline and doubled effective per-signal prices $\lambda$, relative to its value at $\mathrm{w} = 20$ under the baseline price. In logarithmic coordinates the slopes are $\frac{1}{2}$ and $\frac{3}{4}$, as in the model of Prat-Carrabin and Woodford; doubling the effective price shifts both curves upward by the common factor $2^{\frac{1}{4}}$ while leaving the slopes unchanged.

**Proposition 3 (the price of joules).** *At the optimum and at fixed stakes, the task's energy is* $E^* = e\, n^* \propto e^{1/2}\mu^{-1/2}K^{-1}$*, and the expected loss satisfies* $\beta L^* = \mu E^*$*.*

The same optimum relates the behavioral cost of imprecision to its energetic cost. At the optimum, the expected loss of the task, valued at its stakes, equals the energy that the encoding population spends valued at the shadow price, and that energy falls with the square root of the price (Proposition 3). This is the familiar relation between expenditure and marginal value, and it has a direct consequence for the debate on the cost of cognition: a high price produces a small expenditure, so that a computation can draw few joules and still carry a large cost in the currency of the decision, and the number of joules spent does not reveal how much a computation matters.

#### The budget and the price

The shadow price is set by the budget as a whole. In equilibrium, it equates the marginal regulatory cost of raising total use with the marginal value of energy in competing uses. Through the fourth-root law, whatever raises the price raises imprecision in every task that draws on the budget (Figure 3A). A task whose demand is small relative to the rest of the budget takes this price as given, as a small buyer takes the price of a large market (Lemma 2), so that the price of a computation is imported from the state of the whole budget and can be high even when the computation itself draws little energy.

**Proposition 4 (the margin).** *For every convexity* $k > 0$ *of regulation, the budget has a unique equilibrium shadow price* $\mu$*, which falls with the reallocation margin* $M$ *and rises with the value* $v$ *of competing uses and with the stiffness* $c$ *of regulation; under the hard cap, an equilibrium exists if and only if* $M > 0$*.*

**Lemma 2 (price taking).** *The elasticity of* $\mu$ *with respect to the scale of the task's demand equals* $(E/\mu)/F'(\mu)$, *where* $F(\mu) = 0$ *is the equilibrium condition of the budget (Methods), and it vanishes as the task's share of the energy above committed use goes to zero.*

Switching a task on raises the demand for energy, and the budget accommodates it in two ways, since regulation lets total use rise while competing uses give up part of their energy. The share that appears as an increase in total use follows the logic of incidence in a market: it equals the responsiveness of regulated supply relative to the combined responsiveness of supply and of competing demand, so that the stiffer the regulation and the more elastic the competing uses, the larger the part of the task's energy that is withdrawn from other uses instead of added to the total (Lemma 3). The whole-brain increment therefore understates the energy that the task spends, and the two can move in opposite directions: as the margin tightens, the higher price lowers the task's own energy while the share passed through to total use rises, and the net increment grows (Figure 3B). The second effect dominates when most of the task's energy is reallocated and regulation is quadratic, and it can fail when regulation is strongly convex and competing uses are strongly curved (Methods).

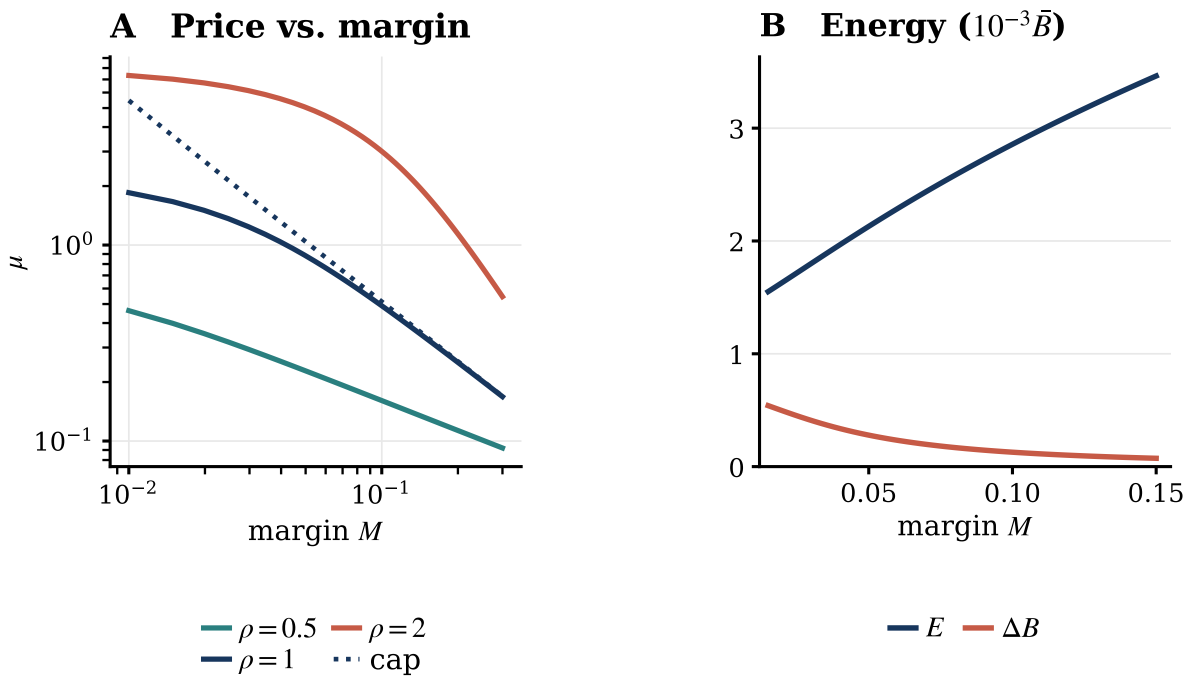


**Figure 3:** Price transmission and energy reallocation across the shared budget. (A) Shadow price $\mu$ over the normalized margin M under regulated curves with k = 1 and c = 100 for competing-use curvatures $\rho = 0.5$, 1, and 2; the dotted line is the hard-cap benchmark at $\rho = 1$. A narrower margin raises $\mu$ across the displayed specifications. (B) Gross task energy E and the net change in total use $\Delta$B under the illustrative baseline, with the set point normalized to one; both energy measures are plotted in units of $10^{-3}$ times this set point. As M tightens, gross task expenditure falls while $\Delta$B rises because less demand can be offset by reallocating competing use, distinguishing the task population's direct draw from the aggregate energy response.

**Lemma 3 (the net increment).** *Introduce task demand as* $\varepsilon E^u(\mu_0)$ *around the no-task equilibrium* $\mu_0$. *As* $\varepsilon \to 0$, *total use changes by* $\Delta B = \varepsilon\phi_0 E^u(\mu_0) + o(\varepsilon)$ *and competing use falls by* $\varepsilon(1-\phi_0)E^u(\mu_0) + o(\varepsilon)$, *where* $\phi_0 = (d/k)/(d/k + E_o/\rho)$ *is evaluated at that no-task equilibrium. For a local comparison of no-task equilibria that differ through* $M$ *alone, holding* $c, v, k, \rho, e, K, \beta$ *and task scale fixed, the first-order coefficient* $\phi_0 E^u(\mu_0)$ *rises with* $\mu_0$ *if and only if* $(1-\phi_0)(1/k + 1/\rho) > 1/2$. *This is a local result for an infinitesimal task; it does not establish monotonicity for finite task demand. A sufficient condition is* $\phi_0 < 1/2$ *and* $1/k + 1/\rho \geq 1$.

**Proposition 6 (sign test).** *For the same task, prior, loss, decoder, and task scale across states, at the interior continuous optimum described above, let $x = \Delta \log p$, $y = \Delta \log e$ and $z = \Delta \log K$, where $p = \mu/\beta$ is the budget price per unit loss, and define $T = y - 2z$. Then $\Delta \log \sigma = (x + T)/4$ and $\Delta \log E = (T - x)/2$, while $\Delta \log(\sigma^2 E) = T$ for any change. For a nonzero pure price change ($x \neq 0, T = 0$), the log–log slope of imprecision against task energy is $-1/2$; for a nonzero pure technology change ($x = 0, T \neq 0$), it is $+1/2$. The finite slope is defined only when $T \neq x$; when $T = x \neq 0$, task energy is unchanged while imprecision changes, and when $x = T = 0$, both are unchanged.*

Two conditions qualify the test. Changes in the stakes of the task act exactly as changes in the price, since precision responds to the price per unit of loss, and a state that changes the value of rewards, as hunger does for food, must therefore be held apart by incentives whose value does not depend on the state. The energy of the encoding population is also measured together with displaced activity: a regional task-evoked change in glucose uptake records the task's energy minus the activity that it displaces in the same region, which preserves the slopes when displacement is proportional to the task's energy and can bend or reverse them when it is not (Supplementary Section S9.3). When both conditions are met, a pure price change also satisfies an overidentifying restriction that links the change in imprecision to the change in energy, and either measure then estimates the price ratio itself.

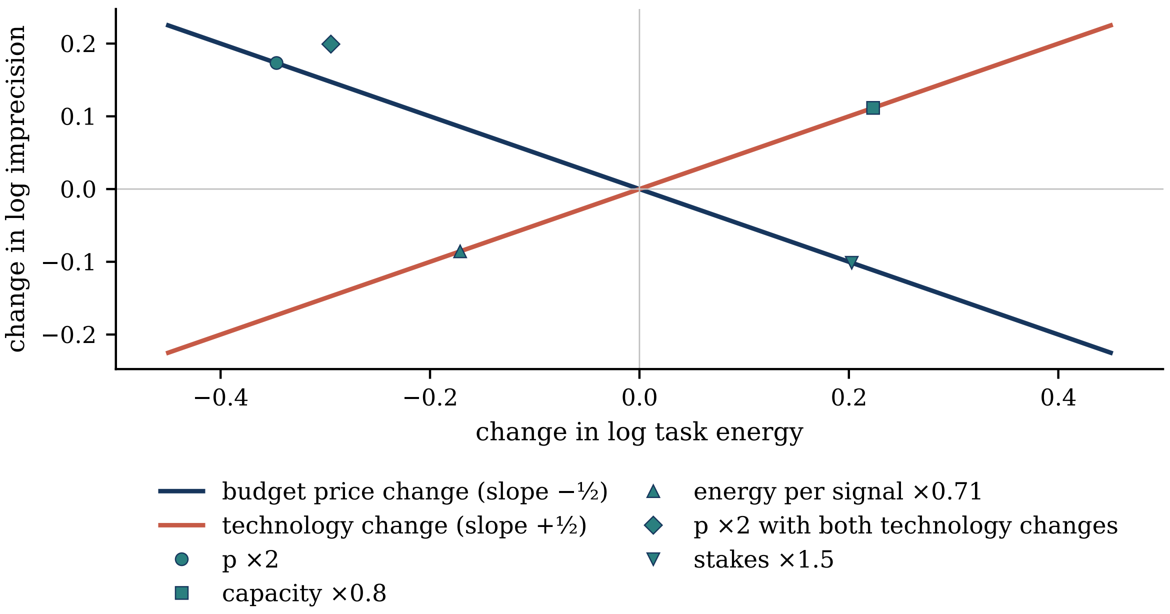


**Figure 4:** Conditional sign restriction for the budget price and technology. Changes in log imprecision are plotted against changes in log task energy. A pure change in p at fixed e and K moves the measures in opposite directions with slope $-\frac{1}{2}$; a pure technology change moves them together with slope $+\frac{1}{2}$. The markers show model comparative statics, including a mixed case in which a doubling of p outweighs lower energy per signal and capacity. The mapping identifies the combined technology index T; additional measurements of e and K resolve its components.

#### Simulation-based design assessment

The behavioral half of the test can be assessed by simulation, with the behavioral layer calibrated to the published numerosity tasks and a within-subject design with two states (Methods). Rejection rates rise with the size of the price change and with the number of participants (Figure 5): in the discrimination task, a doubling of the price, which raises imprecision by about a fifth, is detected in nearly all simulated studies with 40 participants, whereas a rise of a quarter requires substantially larger samples. The simulations also expose the main threat to the inference. When the rate of lapses changes between states, a fit that assumes a common lapse rate attributes the change to imprecision and reports a price change that did not occur, whereas a fit with state-specific lapse rates removes this bias at a cost in power, so that lapses must be measured state by state. Rejection rates, Monte Carlo intervals and fit diagnostics are reported in Supplementary Section S9.4.

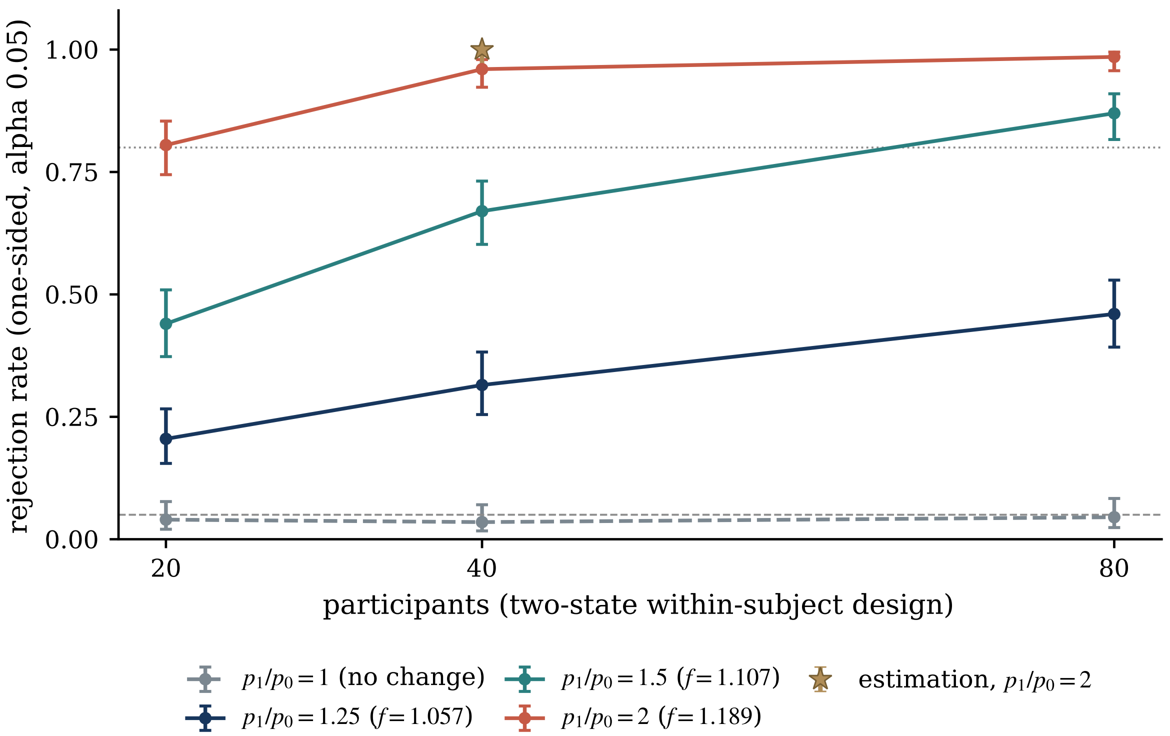


**Figure 5:** Simulated rejection rates for detecting a state-dependent change in the effective per-signal price in within-subject designs. Discrimination curves report the one-sided t-test on fitted log f with 200 trials per state; effective per-signal price ratios of 1, 1.25, 1.5 and 2 correspond to the labeled common factors in imprecision. The star marks the distinct estimation design using a one-sided signed-rank test, 120 trials per state and a doubled effective per-signal price. Error bars are 95% Wilson intervals from 200 replications for discrimination and 100 for estimation. Horizontal lines mark the nominal 0.05 test size and 0.80 rejection rate; lapse rates are held constant across these main design simulations.

#### Matched-numerosity response variability

The encoding layer can be confronted with the published estimation data of Prat-Carrabin and Woodford (2026), released as 12,960 trials from 36 participants (Prat-Carrabin and Woodford, n.d.) (Methods), whose reconstructed standard deviations reproduce the published summaries and thereby anchor the calibration of the behavioral layer to the raw trials. Holding numerosity fixed at the 21 levels that all three priors include, residual response variance rises with prior width, with median ratios of 2.86 for widths 40 and 20 and 4.25 for widths 60 and 20; the pooled ratio exceeds one at all 21 numerosities in both comparisons with width 20 and at 19 of the 21 for widths 60 and 40 (Figure 6). Participant-cluster bootstrap intervals and multiplicity-adjusted p-values are reported in the Supplementary Materials as nominal diagnostics, since a calibration pilot under a fitted null indicates that the procedure is liberal (Supplementary Section S10). The calibrated encoding layer, whose two parameters were fitted only at numerosity 60, predicts these ratios out of sample (2.73, 4.57 and 1.68), a descriptive comparison, since the calibration targets condition-level variability while the ratios use within-participant residuals. The comparison checks the encoding component on which the price operates and leaves the price, neural energy and the budget untested, and, because prior width is linked to block position in the published design, it does not identify a causal effect of width; the inferential apparatus and its sensitivity analyses are reported in Supplementary Sections S6 to S8.

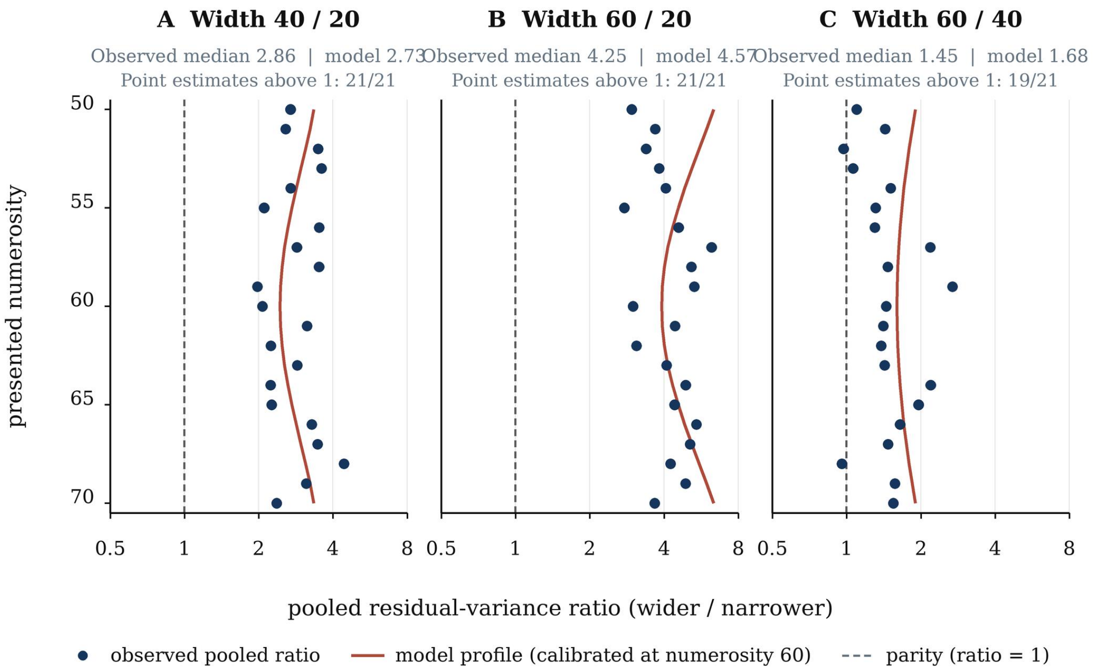


**Figure 6:** Matched-numerosity response variability in the published estimation cohort. Panels A–C show pooled residual-variance ratios for prior-width pairs 40/20, 60/20, and 60/40 at the 21 shared numerosities. Navy points are observed ratios after centering within participant × width × numerosity cells; the dashed vertical reference marks equal residual variance. Rust curves show the model-implied width-scaling profile, calibrated to published condition-level standard-deviation targets at numerosity 60 and displayed across the shared range. Each panel reports the observed median and corresponding model-profile median; a second subtitle gives the number of numerosity-specific observed point estimates above one: 21/21 for 40/20 and 60/20, and 19/21 for 60/40.

#### Derivations of the encoding results

For Lemma 1, set $\mathrm{s(x)} = \mathrm{I_1(x)}^{\frac{1}{2}}$. At fixed n, the objective becomes $(1/\mathrm{n})\int \pi(\mathrm{x})^{\mathrm{a}}/\mathrm{s(x)}^2\mathrm{dx}$ under $\int \mathrm{s(x)dx} \leq \mathrm{K}$. Pointwise minimization of $\pi(\mathrm{x})^{\mathrm{a}}/\mathrm{s(x)}^2 + \nu \mathrm{s(x)}$, where $\nu$ is the multiplier on the information-allocation bound, gives $\mathrm{s(x)} \propto \pi(\mathrm{x})^{\mathrm{a}/3}$; the positive second derivative confirms a minimum. Normalizing to exhaust K yields $\mathrm{s(x)} = \mathrm{K}\pi(\mathrm{x})^{\mathrm{a}/3}/\mathrm{Z_a}$ and loss $\mathrm{Z_a}^3/(\mathrm{nK}^2)$. Thus the optimal shape depends on the prior, while n scales total precision without changing that shape.

For Proposition 1, minimize $\mathrm{Z_a}^3/(\mathrm{nK}^2) + \lambda \mathrm{n}$ over $\mathrm{n} > 0$. The first-order condition gives $\mathrm{n}^* = \mathrm{Z_a}^{3/2}/(\mathrm{K}\lambda^{\frac{1}{2}})$, at which the allocation loss and signal cost are equal. Substitution yields $\mathrm{I(x)} = \mathrm{n}^*\mathrm{s(x)}^2 = \mathrm{KZ_a}^{-\frac{1}{2}}\lambda^{-\frac{1}{2}}\pi(\mathrm{x})^{2\mathrm{a}/3}$ and $\sigma(\mathrm{x}) = \mathrm{Z_a}^{\frac{1}{4}}\lambda^{\frac{1}{4}}\mathrm{K}^{-\frac{1}{2}}\pi(\mathrm{x})^{-\mathrm{a}/3}$. Under a uniform prior of width w, $\mathrm{Z_a} = \mathrm{w}^{1-\mathrm{a}/3}$, so $\sigma \propto \mathrm{w}^{(1+\mathrm{a})/4}$ and $\mathrm{n}^* \propto \mathrm{w}^{(3-\mathrm{a})/2}$. These expressions make the task-specific width scaling and common price elasticity explicit. The derivation therefore isolates a common price response while retaining the distinct way each task uses prior information.

The closed-form optimum separates the task-specific prior shape from the common response to price:

$$n^* = \frac{Z_a^{3/2}}{K\lambda^{1/2}}, \qquad \sigma(x) = Z_a^{1/4}\lambda^{1/4}K^{-1/2}\pi(x)^{-a/3}. \tag{1}$$

The paired expressions in (1) show why the same fourth-root price response applies across prior distributions while the prior continues to determine the shape of imprecision across stimuli.

Proposition 2 follows because $\log \sigma$ is affine in $\log \lambda$ with coefficient $\frac{1}{4}$ at each x, and $\lambda =$ e p, where p $= \mu/\beta$ is the budget price per unit loss. The cross-state fourth-root comparison maps to a $\mu$ ratio only when e, K and $\beta$ are held fixed; with stable e, the $\lambda$ ratio equals the p ratio, and with stable $\beta$, the p ratio equals the $\mu$ ratio. Under the generalized bound $\int \mathrm{I}_1{}^{\gamma/2}\mathrm{dx} \leq \mathrm{K}$, the optimal allocation still separates from n and the optimized loss remains proportional to 1/n; hence $\mathrm{n}^* \propto \lambda^{-\frac{1}{2}}$ and $\sigma \propto \lambda^{\frac{1}{4}}$, although the exponents in w change. A convex signal cost $\lambda \mathrm{n}^\kappa$ instead gives $\mathrm{n}^* \propto \lambda^{-1/(\kappa+1)}$ and $\sigma \propto \lambda^{1/(2(\kappa+1))}$. The discrete-count calculation uses the baseline specification $\gamma = \kappa = 1$: for integer n, it minimizes $\mathrm{G/n} + \lambda\mathrm{n}$ over $\mathrm{n} \in \{1,2,\ldots\}$, with $\mathrm{G} = \mathrm{Z_a}^3/\mathrm{K}^2$. Comparing adjacent integer costs yields intervals in which the optimal count changes; the reported 3–10 range is computed by evaluating these exact discrete optima on a fine grid of continuous optima.

For Proposition 3, multiplying $n^*$ by $e$ gives $E^* \propto e^{1/2}\mu^{-1/2}K^{-1}$ when $\beta$ is fixed. In loss units, $L^* = \lambda n^*$ with $\lambda = e\mu/\beta$; equivalently, $\beta L^* = \mu E^*$. Under the normalization $\beta = 1$, this reduces to $L^* = \lambda n^* = \mu E^*$. Together, these expressions bridge prior-weighted precision, task energy, and valued loss, allowing the budget mechanism to determine a behavioral price rather than treating that price as an external input.

### Derivations of the budget results

At each state, the finite-demand allocation solves the joint problem $\max_{n>0,\, E_o>0,\, d>0}\{V(E_o) - \beta L_a(n) - R(d) : en + E_o = M + d\}$, with $e, K, \beta, c, v, k, \rho > 0$ and $R'(d) = cd^k$. At an interior solution, the first-order conditions are $\beta[-L_a'(n)]/e = V'(E_o) = R'(d) = \mu$, so the per-signal price is $\lambda = e\mu/\beta$. The equilibrium price is therefore determined jointly with task demand; the price-taking result in Lemma 2 is the limiting case as this task's demand becomes negligible relative to the rest of the budget.

For Proposition 4, write $\mathrm{d}(\mu) = (\mu/\mathrm{c})^{1/\mathrm{k}}$, $\mathrm{E}(\mu) \propto \mu^{-\frac{1}{2}}$, and $\mathrm{E_o}(\mu) = (\mathrm{v}/\mu)^{1/\rho}$. The equilibrium condition and its derivative are:

$$F(\mu) = d(\mu) + M - E(\mu) - E_o(\mu) = 0, \qquad F'(\mu) = \frac{d(\mu)}{k\mu} + \frac{E(\mu)}{2\mu} + \frac{E_o(\mu)}{\rho\mu} > 0. \tag{2}$$

Strict monotonicity in (2) gives at most one equilibrium price; for finite c it tends to opposite signs at the positive-price boundaries, establishing existence and uniqueness. Implicit differentiation gives $\partial\mu/\partial\mathrm{M} = -1/\mathrm{F}' < 0$ and positive comparative statics with respect to v and c. Under the hard cap d = 0, the residual tends to M at large $\mu$, so a finite equilibrium requires M > 0. This monotonicity is the analytical basis for the budget curves in Figure 3.

For Lemma 2, scale the task demand by s so the equilibrium equation contains $\mathrm{sE}(\mu)$. Implicit differentiation at s = 1 gives $\partial\mu/\partial\mathrm{s} = \mathrm{E/F'}$, and the corresponding elasticity is $(\mathrm{E}/\mu)/\mathrm{F}'$. As task demand becomes negligible, this elasticity tends to zero: the budget sets the price, while a small task takes it as given.

For Lemma 3, introduce task demand gradually as $\varepsilon\mathrm{E^u}(\mu)$, where $\mathrm{E^u}$ is task energy at unit scale, and denote the no-task equilibrium by $\mu_0$. At that baseline, $\mathrm{d}\mu/\mathrm{d}\varepsilon = \mathrm{E^u}(\mu_0)/F_0'$, where $F_0' = \mathrm{d}(\mu_0)/(\mathrm{k}\mu_0) + \mathrm{E_o}(\mu_0)/(\rho\mu_0)$. Since $\mathrm{d}\Delta\mathrm{B}/\mathrm{d}\varepsilon = \mathrm{d}'(\mu_0)\mathrm{d}\mu/\mathrm{d}\varepsilon$, the first-order expansion and its allocation share are:

$$\Delta B = \phi_0 \varepsilon E^u(\mu_0) + o(\varepsilon), \qquad \phi_0 = \frac{d(\mu_0)/k}{d(\mu_0)/k + E_o(\mu_0)/\rho}. \tag{3}$$

The coefficient in (3), $\phi_0$, measures the local share of an infinitesimal task-energy increase that remains in total use after competing demand adjusts. For a local comparison of no-task states differing through M alone, holding c, v, k, $\rho$, e, K, task stakes and task scale fixed, $\phi_0$ has elasticity $(1-\phi_0)(1/\mathrm{k} + 1/\rho)$ with respect to $\mu_0$. Because $\mathrm{E}^{\mathrm{u}}(\mu_0) \propto {\mu_0}^{-\frac{1}{2}}$, the first-order net increment has elasticity $(1-\phi_0)(1/\mathrm{k} + 1/\rho) - \frac{1}{2}$ and rises with $\mu_0$ under this comparison when $(1-\phi_0)(1/\mathrm{k} + 1/\rho) > \frac{1}{2}$. In particular, $\phi_0 < \frac{1}{2}$ and $1/\mathrm{k} + 1/\rho \geq 1$ provide a transparent sufficient parameter condition. The numerical baseline separately compares this no-task first-order expression with the exact finite-task equilibrium change.

For Proposition 5, normalize task-plus-competing use to one. For any target share $s \in (0, 1)$ and price $\mu > 0$, set $e = 1$, choose the task-scale primitive so that $E(\mu) = s$, and set $E_o = 1 - s$. Choose any $d \in (0, 1)$ and let $M = 1 - d$; then $d + M = E + E_o = 1$. Choosing $\bar{B} > M$ and $C = \bar{B} - M$ keeps committed use positive, while $c = \mu/d^k$ and $v = \mu E_o^\rho$ give $R'(d) = V'(E_o) = \mu$. The resulting share of energy above committed use is $E/(E + E_o) = s$. This establishes the construction for the full interior domain; the grid of sixteen pairs checks its numerical implementation, with maximum relative error $2.22 \times 10^{-16}$. Varying $c$ while adjusting $v$ to hold $\mu$ fixed shows that the net-to-gross increment ratio is not uniquely determined by price. The construction makes precise why the task share and net increment are complementary observations rather than substitutes for the joint price-and-precision restrictions.

For the joint non-identification claim, take $k = \rho = 1$ and hold the task-demand coefficient $A$ fixed, so $E(\mu) = A\mu^{-1/2}$. Choose a common $\Delta > 0$ and two distinct prices $\mu_j$ for which $E_j = A\mu_j^{-1/2} > 2\Delta$. Set $r_j = E_j/(2E_j - \Delta)$, $\mu_{0j} = r_j\mu_j$, $v_j = \mu_j E_j$, $c_j = \mu_j(1 - r_j)/\Delta$, and $M_j = E_j/r_j - r_j\Delta/(1 - r_j)$. These primitives are positive. They imply $E_o(\mu_j) = E_j$ and $E_o(\mu_{0j}) = E_j/r_j = 2E_j - \Delta$, so each task share is 1/2; they also imply $d(\mu_j) - d(\mu_{0j}) = \Delta$. The two distinct prices therefore generate the same task share and net increment with the same task technology $A$, while the budget primitives differ. This construction clarifies why aggregate shares and net increments are complementary observations, while the joint behavioral-energy restrictions carry cross-state information about price.

#### The sign test, measurement, and numerical verification

For each fixed task specification across states, define the budget price per unit loss $p = \mu/\beta$, so that $\lambda = ep$ is the effective price per signal, and let $E = en^*$ denote gross encoding energy. Taking log changes in $\sigma \propto (ep)^{1/4}K^{-1/2}$ and $E \propto e^{1/2}p^{-1/2}K^{-1}$ gives the joint log-change restriction:

$$\begin{aligned} \Delta \log \sigma &= \frac{x+y}{4} - \frac{z}{2}, \qquad \Delta \log E = \frac{y-x}{2} - z, \\ \frac{\Delta \log \sigma}{\Delta \log E} &= \frac{x+T}{2(T-x)}, \quad \Delta \log(\sigma^2 E) = T. \end{aligned} \tag{4}$$

Here T = y−2z is the combined technology index. The joint restriction in (4) defines the finite slope only when $\Delta\log E \neq 0$ ($T \neq x$); when $T = x \neq 0$, gross energy is unchanged while imprecision changes, and when x = T = 0 both measures are unchanged. A pure budget-price change has T = 0 and slope $-\frac{1}{2}$; a pure technology change has x = 0 and slope $+\frac{1}{2}$. The pair of observables therefore identifies the change in budget price p and the combined index T under the model, but not $\mu$ separately when $\beta$ changes, nor e and K separately. At fixed p, offsetting technology changes with $T = \Delta\log(e/K^2) = 0$ leave both $\sigma$ and E unchanged; these outcomes alone therefore cannot distinguish them from unchanged technology. Price claims about $\mu$ consequently require stable or measured stakes.

### Competing interests

The authors declare no competing interests.

### Funding

This work received no specific funding.

### Author contributions

Fabio Lokwani-Di Matteo: formal analysis, software, validation, writing – original draft, writing – review and editing. Pier Luigi Sacco: conceptualization, methodology, formal analysis, writing – original draft, writing – review and editing.
