## Supplementary material for "The marginal brain": Replication package: Manuscript.tex

\ExplSyntaxOn\cs_undefine:N \qed\ExplSyntaxOff
\documentclass[10pt]{article}
\usepackage[margin=1in]{geometry}
\usepackage[round,authoryear]{natbib}
\makeatletter
\newcommand{\biboptions}[1]{}
\setcounter{secnumdepth}{0}
\gdef\fm@address{}
\newcommand{\address}[1]{\gdef\fm@address{#1}}
\newcommand{\fm@printtitle}{\begin{center}{\Large\bfseries\@title\par}\vspace{0.8em}{\@author\par}\vspace{0.4em}{\small\fm@address\par}\end{center}\vspace{0.6em}\global\let\fm@printtitle\relax}
\newenvironment{frontmatter}{}{\fm@printtitle}
\let\fm@origabstract\abstract \let\fm@origendabstract\endabstract
\renewenvironment{abstract}{\fm@printtitle\fm@origabstract}{\fm@origendabstract}
\newenvironment{keyword}{\par\noindent\textbf{Keywords:} }{\par\medskip}
\makeatother
\usepackage{fontspec}
\usepackage{microtype}
\usepackage{amsmath,amssymb}
\usepackage{graphicx}
\usepackage{float}
\usepackage{longtable}
\usepackage{array,ragged2e,booktabs}
\usepackage{caption}
\usepackage{xurl}
\usepackage[hidelinks]{hyperref}
\hypersetup{pdftitle={The marginal brain},pdfauthor={Fabio Lokwani-Di Matteo; Pier Luigi Sacco}}
\usepackage{lineno}
\biboptions{round,authoryear}
\providecommand{\textsubscript}[1]{\ensuremath{_{\mathrm{#1}}}}
\providecommand{\textsuperscript}[1]{\ensuremath{^{\mathrm{#1}}}}
\captionsetup{font=small,labelfont=bf,justification=justified,singlelinecheck=false}
\renewcommand{\arraystretch}{1.12}
\setlength{\tabcolsep}{5pt}
\setlength{\parindent}{1.25em}
\setlength{\parskip}{0.35em}
\setlength{\emergencystretch}{2em}
\clubpenalty=10000
\widowpenalty=10000
\pagestyle{plain}
\begin{document}
\linenumbers
\begin{frontmatter}
\title{The marginal brain}
\author{Fabio Lokwani-Di Matteo\textsuperscript{a} and Pier Luigi Sacco\textsuperscript{a,b}}
\address{\textsuperscript{a}Department of Neurosciences, Imaging and Clinical Sciences, University of Chieti-Pescara, Italy \\ \textsuperscript{b}metaLAB (at) Harvard, Harvard University, Cambridge, MA, USA \\ Correspondence: Pier Luigi Sacco,, }
\begin{abstract}
Goal-directed cognition raises the brain's energy use only a few percent above resting use, a figure often read as evidence that cognition is metabolically cheap. Models of endogenous precision derive the precision of neural representations from a cost on encoding activity whose unit price is a free parameter. Embedding endogenous precision in a regulated, shared energy budget makes this cost the shadow price of energy at the margin of the budget. Neither the task's share of the budget nor the net whole-brain increment identifies this price; when tasks face a common price ratio and coding technology is stable, imprecision changes by the exact fourth root of that ratio, so that the cost of information rescales while its shape is preserved; and the co-movement of imprecision and stimulus-linked task energy separates price changes from changes in coding technology. In 12,960 published estimation trials, response variance at matched numerosities rises with prior width in line with the ratios that the calibrated model predicts out of sample, and simulations indicate that a doubled price is detectable with existing paradigms when state-dependent lapses are modeled.

\end{abstract}
\begin{keyword}
endogenous precision, shadow price, brain energy budget, efficient coding, cognitive effort, numerosity
\end{keyword}
\end{frontmatter}
\section{Introduction}

Functional imaging finds that task-evoked changes in brain energy are modest relative to resting use, which dominates the brain's measured budget \citep{jamadar2025,raichle_mintun2006,gusnard_raichle2001}, and foundational PET and fMRI studies document systematic regional decreases from resting baselines during goal-directed tasks, which calls for a distinction between local responses and aggregate changes from baseline \citep{raichle2001,gusnard_raichle2001}. The small aggregate change has been read as evidence that cognition is metabolically inexpensive, since withdrawing from a task would save little total energy \citep{kondrakiewicz2025}. The aggregate change is, however, a net quantity, which combines the energy that the task recruits with the activity that it displaces elsewhere in a shared system, and if task demand is partly met by reallocating energy from competing uses, a small change in total use need not imply a small marginal value of energy. The distinction matters for theories of effort, which have emphasized opportunity costs over metabolic expenditure \citep{kurzban2013} and which recent reviews leave without a specified source of cost \citep{otto2026}, and for current efforts to make cognitive models both biologically grounded and predictively informative \citep{haueis2026a,haueis2026b}. The quantity that matters for behavior is the marginal cost of the joules that a task draws, and the question is how that cost shapes the precision of the representations that guide behavior.

Two lines of work bear on this question. In neuroscience, vascular supply and the homeostasis of oxygen delivery constrain regional metabolism \citep{herculanohouzel2022}, attention redistributes metabolic activity across cortical regions \citep{bruckmaier2020}, the energy cost of spikes bounds sustainable activity in neural populations \citep{lennie2003}, and cellular energy budgets quantify the costs of action potentials and synaptic transmission \citep{attwell2001,harris2012}; these observations do not specify how the marginal value of a shared budget enters the computations that a behavioral model represents. In economics and in the cognitive science of decision, noisy-representation and rational-inattention models formalize how precision and the use of information respond to goals and to the costs of information \citep{woodford2020,mackowiak2023,dean2023}, related work links adaptive sensory allocation in mice to reward and to limited processing resources \citep{grujic2022}, and resource-rational models assume constraints that vary from model to model and resist unification \citep{ma2020}. Energy-efficient coding has long priced information in energy \citep{laughlin1998,balasubramanian2001}, and it has been argued that cognitive systems optimize energy \citep{markman2011}, yet to our knowledge no model derives the precision of the representations that guide decisions from the brain's energy budget through a price.

The closest formal point of departure is the endogenous-precision model of numerosity judgments of \citet{pratcarrabin2026}, in which an encoding-and-decoding model with task-specific losses, an efficient-coding constraint and a per-signal cost generates prior-width exponents of one half for estimation and three quarters for discrimination, which their human data support. The fourth-root response of imprecision to a change in the per-signal price follows from the same Fisher-information and linear-cost structure, and we use it as a conditional implication of the retained encoding layer. The extension developed here closes that layer with a shared energy budget, in which the unit price of a signal is set endogenously by the energy per signal and the marginal value of the budget; this closure yields comparative statics across competing tasks and restrictions that link task allocation, gross and net energy changes, and joint measurements of imprecision and stimulus-linked energy.

We embed the endogenous-precision encoding problem in a common energy budget with committed use, competing demands and regulation around a set point. At equilibrium, the marginal regulatory cost determines a shadow price \ensuremath{\mu}, which, combined with the energy per signal e and the task stakes \ensuremath{\beta}, sets the effective per-signal price \ensuremath{\lambda} = e\ensuremath{\mu}/\ensuremath{\beta}, so that behavioral precision responds to the state of the whole budget instead of a price fitted separately to each task, and three results follow from this construction. First, neither a task's share of the energy above committed use nor the net whole-brain increment identifies the shadow price, because both depend on how the rest of the budget is allocated and on how strongly total use is regulated. Second, when technology and stakes are stable and tasks face a common price ratio, imprecision changes by the exact fourth root of that ratio while retaining its task-specific scaling with prior width, which imposes a restriction across tasks and implies that the cost of information rescales with the price while its shape is preserved. Third, joint changes in imprecision and stimulus-linked task energy distinguish price-dominant from technology-dominant changes under explicit conditions on displacement.

Two empirical components complement the analysis. The first reanalyzes the numerosity-estimation experiment of \citet{pratcarrabin2026}, in which 36 adults (20 women, 15 men and one non-binary participant, with a mean age of 21.4 years) reported their best estimate of the number of dots in arrays presented for 500 ms, with numbers drawn from uniform priors over 50 to 70, 40 to 80 and 30 to 90 in three successive blocks, whose order of widths was increasing for 20 participants and decreasing for 16; the authors have released the resulting 12,960 trials, 360 per participant, as open data \citep{pratcarrabin_data}. Holding numerosity fixed and resampling participants, residual response variance rises with prior width, with median ratios of 2.86 for widths 40 and 20 and 4.25 for widths 60 and 20, close to the ratios of 2.73 and 4.57 that the calibrated encoding layer predicts out of sample. The second component consists of simulations calibrated to the published numerosity tasks, which show that a doubling of the price can be detected with existing paradigms, provided that lapses that change across states are modeled. We present the model, derive its results on precision, on the budget and on identification, report the reanalysis and the design simulations, and discuss rival accounts and limitations; derivations and analysis details are given in Methods and in the Supplementary Materials.

\section{Model}

\subsection{Endogenous precision}

We build on the endogenous-precision model developed for the perception of numerical magnitudes \citep{pratcarrabin2026}. A magnitude x, drawn from a prior density \ensuremath{\pi}, is encoded with effective signal mass n, a continuous activity measure in the optimization rather than a literal count of individual spikes. Each independent signal carries Fisher information I\textsubscript{1}(x), and the allocation of precision across magnitudes is bounded by \ensuremath{\int} I\textsubscript{1}(x)\textsuperscript{\ensuremath{\frac{1}{2}}} dx \ensuremath{\leq} K, a constraint used in efficient-coding models of unidimensional channels (Prat-Carrabin and Woodford write its right-hand side as \ensuremath{\sqrt{K}}, so K here corresponds to their \ensuremath{\sqrt{K}}). Because Fisher information adds across independent signals, total precision is I(x) = n I\textsubscript{1}(x); in the high-precision regime, an efficient decoder attains imprecision \ensuremath{\sigma}(x) = I(x)\textsuperscript{\ensuremath{-}\ensuremath{\frac{1}{2}}}. The task loss is approximated by L\textsubscript{1}[I] = \ensuremath{\int} \ensuremath{\pi}(x)/I(x) dx for estimation and L\textsubscript{2}[I] = \ensuremath{\int} \ensuremath{\pi}(x)\textsuperscript{2}/I(x) dx for discrimination. The observer chooses I\textsubscript{1} and the continuous effective activity n to minimize L\textsubscript{a} + \ensuremath{\lambda}n, with a = 1 for estimation and a = 2 for discrimination. Under a uniform prior of width w, imprecision scales as w\textsuperscript{\ensuremath{\frac{1}{2}}} in estimation and w\textsuperscript{\ensuremath{\frac{3}{4}}} in discrimination, as supported by the cited numerosity data \citep{pratcarrabin2026}. In this model, \ensuremath{\lambda} is a behavioral price held fixed across the observer's states; the next section endogenizes its state dependence through the energy budget.

\subsection{The brain's energy budget}

We embed this choice in a shared energy budget. Total use B comprises committed use C, gross task energy E = e n, and energy E\textsubscript{o} allocated to competing uses. For v > 0 and \ensuremath{\rho} > 0, the latter's value is V(E\textsubscript{o}) = v(E\textsubscript{o}\textsuperscript{1\ensuremath{-}\ensuremath{\rho}} \ensuremath{-} 1)/(1\ensuremath{-}\ensuremath{\rho}) for \ensuremath{\rho} \ensuremath{\neq} 1 and V(E\textsubscript{o}) = v log E\textsubscript{o} for \ensuremath{\rho} = 1; both give V\ensuremath{^\prime}(E\textsubscript{o}) = vE\textsubscript{o}\textsuperscript{\ensuremath{-}\ensuremath{\rho}}, including the configured baseline \ensuremath{\rho} = 1, and make V increasing and concave. Committed use includes maintenance and intrinsic activity, which together make up most resting consumption \citep{jamadar2025}. Physiological theory and evidence motivate possible supply constraints through vascular architecture and oxygen-delivery homeostasis \citep{herculanohouzel2022}, conserved energy use per neuron across brain sizes \citep{herculanohouzel2011}, and observed redistribution of metabolic activity across cortical regions during attention \citep{bruckmaier2020}. Together these observations motivate the common-budget structure formalized here. We represent it with a set point \ensuremath{\bar{B}} and a regulatory cost whose marginal value is R\ensuremath{^\prime}(d) = c d\textsuperscript{k}, where d = B \ensuremath{-} \ensuremath{\bar{B}} is use above the set point, c > 0 measures regulatory stiffness and k > 0 its convexity; a hard cap is the limit as c grows without bound. Local neurovascular coupling can increase supply during activation even when total use remains near rest \citep{jamadar2025}. Accordingly, the set point may refer to the whole brain or to a region with bounded supply. The reallocation margin M = \ensuremath{\bar{B}} \ensuremath{-} C is the energy available to the task and competing uses before regulation binds; changes in supply enter through this margin.

\subsection{Equilibrium and the price of precision}

The brain minimizes task loss L\textsubscript{a} plus regulatory cost R, less the value V of competing uses. At an interior optimum, the marginal regulatory cost equals the marginal value in competing uses: \ensuremath{\mu} = R\ensuremath{^\prime}(B \ensuremath{-} \ensuremath{\bar{B}}) = V\ensuremath{^\prime}(E\textsubscript{o}). This defines \ensuremath{\mu}, the shadow price of energy. When task stakes scale loss by \ensuremath{\beta}, define p = \ensuremath{\mu}/\ensuremath{\beta} as the budget price per unit loss and \ensuremath{\lambda} = ep = e\ensuremath{\mu}/\ensuremath{\beta} as the effective price per signal; under \ensuremath{\beta} = 1, these reduce to p = \ensuremath{\mu} and \ensuremath{\lambda} = e\ensuremath{\mu}. The ratio of \ensuremath{\lambda} equals the ratio of p only when e is fixed, and the ratio of p equals that of \ensuremath{\mu} only when \ensuremath{\beta} is fixed. The model therefore separates energy per signal from the budget's marginal value and makes clear why cross-state price comparisons require stable stakes. Neurophysiological work estimates energy costs of signaling \citep{laughlin1998}, and neural coding may adapt under scarcity at some cost to precision \citep{padamsey2022}; our model isolates the additional state-dependent price transmitted through \ensuremath{\mu}. Gross encoding energy E differs from net increment \ensuremath{\Delta}B, the change in total use after competing uses reallocate. Whole-brain imaging reports aggregate task-evoked changes of the latter kind \citep{jamadar2025}.

Figure 1 gives a visual map of the model's transmission chain from the shared-budget constraint through the shadow price to precision. The three stages make the relationships and equilibrium feedback visible at a glance.

\begin{figure}[H]\centering
\includegraphics[alt={Three-stage schematic. Panel A shows the shared-budget identities for committed use, the reallocation margin, task energy and competing uses. Panel B equates marginal regulatory cost with the marginal value of competing uses to define the shadow price. Panel C maps that price to effective cost per signal and the resulting imprecision response.},width=0.96\linewidth]{figures/fig1_schematic.pdf}
\caption{The model's three linked stages. Panel A states the shared-budget identities M = \ensuremath{\bar{B}} \ensuremath{-} C and d + M = E + E\textsubscript{o}, with gross task energy E = en. Panel B shows the equilibrium condition \ensuremath{\mu} = R\ensuremath{^\prime}(d) = V\ensuremath{^\prime}(E\textsubscript{o}), equating the marginal regulatory cost and value of competing uses. Panel C maps the shadow price into the per-signal price \ensuremath{\lambda} = e\ensuremath{\mu}/\ensuremath{\beta} and the imprecision response \ensuremath{\sigma} \ensuremath{\propto} \ensuremath{\lambda}\textsuperscript{1/4}, at fixed task and coding parameters. The arrows trace price transmission from the budget equilibrium to encoding; the budget is solved jointly, so task demand also feeds back into the equilibrium price.}\label{fig:figure1}
\end{figure}

Read together, the panels clarify two quantities that can otherwise be conflated: gross task expenditure E and the net whole-brain increment \ensuremath{\Delta}B after competing use adjustments. An observed energy change alone therefore does not identify the shadow price without information about the reallocation margin. By connecting the budget equilibrium to an encoding response, the diagram turns the proposed mechanism into a testable link between resource conditions and behavioral precision.

\subsection{Model timing and scope}

We represent each physiological state as a static allocation equilibrium. The model assumes a separation between the budget state and within-trial encoding, treating the former as fixed over the latter; it does not specify or estimate a price-adjustment speed. In this model, supply reductions lower M, competing demands raise v, and tighter regulation raises c; coding technology enters through e and K. The allocation rule equates marginal values across energy uses; an adaptive signal could convey \ensuremath{\mu} to encoding populations, with neuromodulatory processes as one candidate implementation. When prices are region-specific, \ensuremath{\mu} is the price of the budget used by the encoding population. Precision denotes Fisher information, and costs are expressed in metabolic terms.

\section{Results}

\subsection{Precision and its price}

The encoding layer can be read as a buyer of signals: because Fisher information adds across independent signals, the expected loss of a task falls with the inverse of the number of signals, while their cost rises linearly with that number, and the optimal number equates the marginal reduction in expected loss with the price of a signal. Two consequences organize the rest of the analysis. The allocation of precision across magnitudes depends only on the prior and on the task, since it is fixed before the number of signals is chosen, and it concentrates precision where magnitudes are frequent, in proportion to a power of the prior density that differs between estimation and discrimination (Lemma 1). The number of signals, in turn, falls with the square root of their price, so that the demand for signals has a constant price elasticity of \ensuremath{-}\ensuremath{\frac{1}{2}}, and imprecision, which falls with the square root of the number of signals, rises with the fourth root of the price (Propositions 1 and 2).

\textbf{Lemma 1 (separation).} \emph{Let} \(\pi\) \emph{be a normalized density on support} \(\mathcal X\)\emph{, with} \(0 < Z_{a} = \int_{\mathcal X}\pi(x)^{a/3}\,dx < \infty\)\emph{, and let} \(K>0\) \emph{and} \(n>0\)\emph{. Under admissible information allocations satisfying} \(\int_{\mathcal X} I_{1}(x)^{1/2}\,dx \leq K\)\emph{, the optimum satisfies} \(I_{1}(x)^{1/2} \propto \pi(x)^{a/3}\)\emph{, and the expected loss after allocation is} \(Z_{a}^{3}/(nK^{2})\)\emph{.}

\textbf{Proposition 1 (baseline).} \emph{For} \(\lambda>0\) \emph{and the interior continuous optimum over} \(n>0\)\emph{, the optimal number of signals is} \(n^{\ast} = Z_{a}^{3/2}/(K\lambda^{1/2})\) \emph{and imprecision is} \(\sigma(x) = Z_{a}^{1/4}\lambda^{1/4}K^{-1/2}\pi(x)^{-a/3}\)\emph{; under a uniform prior of width} \(w\)\emph{, holding capacity} \(K\) \emph{and effective signal price} \(\lambda\) \emph{fixed,} \(\sigma \propto w^{1/2}\) \emph{in estimation (}\(a = 1\)\emph{) and} \(\sigma \propto w^{3/4}\) \emph{in discrimination (}\(a = 2\)\emph{).}

\textbf{Proposition 2 (common factor).} \emph{For a fixed task specification (prior, loss, and decoder), the elasticity of imprecision with respect to the effective per-signal price} \(\lambda = e\mu/\beta\) \emph{equals} \(1/4\) \emph{at every magnitude and capacity. If the energy per signal} \(e\)\emph{, the capacity} \(K\)\emph{, the task specification, and the stakes} \(\beta\) \emph{are unchanged across states, a change that moves the shadow price from} \(\mu_0\) \emph{to} \(\mu_1\) \emph{multiplies imprecision by} \((\mu_1/\mu_0)^{1/4}\) \emph{in every task that draws on the same budget and leaves the exponents in} \(w\) \emph{unchanged.}

The fourth-root law is the source of the model's main behavioral prediction. Because every task that draws on the same budget faces the same shadow price, a doubling of the price raises imprecision by a factor of 1.19 in every task and at every magnitude, while the prior and the task continue to determine how imprecision is distributed. In the coordinates of Figure 2, the price shifts the curves of estimation and discrimination upward by the same distance and leaves their slopes with prior width, one half and three quarters, unchanged. The prediction is therefore a restriction across tasks: within a change of state that leaves coding technology and stakes unchanged, estimation and discrimination must show the same state factor, which a model with a price fitted separately to each task does not imply. The law rests only on the additivity of Fisher information and on a cost that is linear in the number of signals, so that a different form of the efficient-coding bound alters the slopes with prior width without altering the law, a convex cost of signals preserves its sign, and integer counts make the response lumpy only when few signals are used (Supplementary Section S9.1).

\begin{figure}[H]\centering
\includegraphics[alt={Line chart of imprecision versus prior width for estimation and discrimination at baseline and doubled effective signal price. Estimation and discrimination retain distinct width slopes of one-half and three-quarters; doubling the price shifts both curves upward by the common fourth-root factor.},width=0.96\linewidth]{figures/fig2_common_factor.pdf}
\caption{Imprecision as a function of prior width in estimation and in discrimination, at baseline and doubled effective per-signal prices \ensuremath{\lambda}, relative to its value at w = 20 under the baseline price. In logarithmic coordinates the slopes are \ensuremath{\frac{1}{2}} and \ensuremath{\frac{3}{4}}, as in the model of Prat-Carrabin and Woodford; doubling the effective price shifts both curves upward by the common factor 2\textsuperscript{\ensuremath{\frac{1}{4}}} while leaving the slopes unchanged.}\label{fig:figure2}
\end{figure}

\textbf{Proposition 3 (the price of joules).} \emph{At the optimum and at fixed stakes, the task's energy is} \(E^{\ast} = e\,n^{\ast} \propto e^{1/2}\mu^{-1/2}K^{-1}\)\emph{, and the expected loss satisfies} \(\beta L^{\ast} = \mu E^{\ast}\)\emph{.}

The same optimum relates the behavioral cost of imprecision to its energetic cost. At the optimum, the expected loss of the task, valued at its stakes, equals the energy that the encoding population spends valued at the shadow price, and that energy falls with the square root of the price (Proposition 3). This is the familiar relation between expenditure and marginal value, and it has a direct consequence for the debate on the cost of cognition: a high price produces a small expenditure, so that a computation can draw few joules and still carry a large cost in the currency of the decision, and the number of joules spent does not reveal how much a computation matters.

\subsection{The budget and the price}

The shadow price is set by the budget as a whole. In equilibrium, it equates the marginal regulatory cost of raising total use with the marginal value of energy in competing uses. Through the fourth-root law, whatever raises the price raises imprecision in every task that draws on the budget (Figure 3A). A task whose demand is small relative to the rest of the budget takes this price as given, as a small buyer takes the price of a large market (Lemma 2), so that the price of a computation is imported from the state of the whole budget and can be high even when the computation itself draws little energy.

\textbf{Proposition 4 (the margin).} \emph{For every convexity} \(k > 0\) \emph{of regulation, the budget has a unique equilibrium shadow price} \(\mu\)\emph{, which falls with the reallocation margin} \(M\) \emph{and rises with the value} \(v\) \emph{of competing uses and with the stiffness} \(c\) \emph{of regulation; under the hard cap, an equilibrium exists if and only if} \(M > 0\)\emph{.}

\textbf{Lemma 2 (price taking).} \emph{The elasticity of} \(\mu\) \emph{with respect to the scale of the task's demand equals} \((E/\mu)/F'(\mu)\)\emph{, where} \(F(\mu) = 0\) \emph{is the equilibrium condition of the budget (Methods), and it vanishes as the task's share of the energy above committed use goes to zero.}

Switching a task on raises the demand for energy, and the budget accommodates it in two ways, since regulation lets total use rise while competing uses give up part of their energy. The share that appears as an increase in total use follows the logic of incidence in a market: it equals the responsiveness of regulated supply relative to the combined responsiveness of supply and of competing demand, so that the stiffer the regulation and the more elastic the competing uses, the larger the part of the task's energy that is withdrawn from other uses instead of added to the total (Lemma 3). The whole-brain increment therefore understates the energy that the task spends, and the two can move in opposite directions: as the margin tightens, the higher price lowers the task's own energy while the share passed through to total use rises, and the net increment grows (Figure 3B). The second effect dominates when most of the task's energy is reallocated and regulation is quadratic, and it can fail when regulation is strongly convex and competing uses are strongly curved (Methods).

\begin{figure}[H]\centering
\includegraphics[alt={Two side-by-side panels. Panel A plots the shadow price against the reallocation margin for three competing-use curvatures and a hard-cap benchmark. Panel B contrasts gross task energy with the net change in total use as the margin tightens; both energy measures are reported in units of 10⁻³ times the normalized set point.},width=0.96\linewidth]{figures/fig3_budget.pdf}
\caption{Price transmission and energy reallocation across the shared budget. (A) Shadow price \ensuremath{\mu} over the normalized margin M under regulated curves with k = 1 and c = 100 for competing-use curvatures \ensuremath{\rho} = 0.5, 1, and 2; the dotted line is the hard-cap benchmark at \ensuremath{\rho} = 1. A narrower margin raises \ensuremath{\mu} across the displayed specifications. (B) Gross task energy E and the net change in total use \ensuremath{\Delta}B under the illustrative baseline, with the set point normalized to one; both energy measures are plotted in units of 10\textsuperscript{-3} times this set point. As M tightens, gross task expenditure falls while \ensuremath{\Delta}B rises because less demand can be offset by reallocating competing use, distinguishing the task population's direct draw from the aggregate energy response.}\label{fig:figure3}
\end{figure}

\textbf{Lemma 3 (the net increment).} \emph{Introduce task demand as} \(\varepsilon E^{u}(\mu_0)\) \emph{around the no-task equilibrium} \(\mu_0\)\emph{. As} \(\varepsilon\to0\)\emph{, total use changes by} \(\Delta B = \varepsilon\phi_0 E^{u}(\mu_0)+o(\varepsilon)\) \emph{and competing use falls by} \(\varepsilon(1-\phi_0)E^{u}(\mu_0)+o(\varepsilon)\)\emph{, where} \(\phi_0 = (d/k)/(d/k + E_{o}/\rho)\) \emph{is evaluated at that no-task equilibrium. For a local comparison of no-task equilibria that differ through} \(M\) \emph{alone, holding} \(c,v,k,\rho,e,K,\beta\) \emph{and task scale fixed, the first-order coefficient} \(\phi_0 E^{u}(\mu_0)\) \emph{rises with} \(\mu_0\) \emph{if and only if} \((1-\phi_0)(1/k + 1/\rho)>1/2\)\emph{. This is a local result for an infinitesimal task; it does not establish monotonicity for finite task demand. A sufficient condition is} \(\phi_0<1/2\) \emph{and} \(1/k+1/\rho\geq1\)\emph{.}

\textbf{Proposition 5 (non-identification).} \emph{For any task share} \(s \in (0,1)\) \emph{of energy allocated above committed use,} \(E/(E+E_{o})\)\emph{, and any price} \(\mu > 0\)\emph{, there exist positive primitives of the budget whose interior equilibrium reproduces both. At a fixed price, the ratio of the net increment to gross task energy can be made arbitrarily small by stiffer regulation, and it vanishes under the hard cap.}

For the same reason, neither the task's share of the energy above committed use nor the net increment identifies the price, since budgets that differ in their margin, their regulation and the value of their competing uses can produce the same share and the same increment at different prices (Proposition 5; Supplementary Section S9.2). A task-evoked increment of a few percent above rest is thus compatible with a high price of energy at the margin, and the reading of such increments as evidence that cognition is cheap draws a conclusion about the price from statistics that do not identify it \citep{jamadar2025,kondrakiewicz2025}.

\subsection{Telling a price from a technology}

A change of physiological state can change the price of energy, the coding technology, or both, and behavior alone cannot tell them apart, since a lower capacity per signal raises imprecision just as a higher price does. The two separate once the energy of the encoding population, which for numerosity lies in parietal cortex, where number encoding adapts to the range of the prior \citep{pratcarrabin2025}, is measured alongside imprecision, because they move the two measures in different directions. A higher price lowers the number of signals that the population buys, so that imprecision rises while the energy spent falls, whereas a change in technology that makes signals costlier or less informative raises imprecision and raises the energy spent on the task, so that the two measures move together (Proposition 6). In the coordinates of Figure 4, an observed change of state can therefore be decomposed into a price component and a technology component, and the sign of the joint change reveals which of them dominates.

\textbf{Proposition 6 (sign test).} \emph{For the same task, prior, loss, decoder, and task scale across states, at the interior continuous optimum described above, let} \(x = \Delta\log p\)\emph{,} \(y = \Delta\log e\) \emph{and} \(z = \Delta\log K\)\emph{, where} \(p = \mu/\beta\) \emph{is the budget price per unit loss, and define} \(T = y - 2z\)\emph{. Then} \(\Delta\log\sigma = (x+T)/4\) \emph{and} \(\Delta\log E = (T-x)/2\)\emph{, while} \(\Delta\log(\sigma^{2}E) = T\) \emph{for any change. For a nonzero pure price change} (\(x \ne 0, T = 0\))\emph{, the log--log slope of imprecision against task energy is} \(-1/2\)\emph{; for a nonzero pure technology change} (\(x = 0, T \ne 0\))\emph{, it is} \(+1/2\)\emph{. The finite slope is defined only when} \(T \ne x\)\emph{; when} \(T=x\ne0\)\emph{, task energy is unchanged while imprecision changes, and when} \(x=T=0\)\emph{, both are unchanged.}

Two conditions qualify the test. Changes in the stakes of the task act exactly as changes in the price, since precision responds to the price per unit of loss, and a state that changes the value of rewards, as hunger does for food, must therefore be held apart by incentives whose value does not depend on the state. The energy of the encoding population is also measured together with displaced activity: a regional task-evoked change in glucose uptake records the task's energy minus the activity that it displaces in the same region, which preserves the slopes when displacement is proportional to the task's energy and can bend or reverse them when it is not (Supplementary Section S9.3). When both conditions are met, a pure price change also satisfies an overidentifying restriction that links the change in imprecision to the change in energy, and either measure then estimates the price ratio itself.

\begin{figure}[H]\centering
\includegraphics[alt={Sign-test plot of changes in log imprecision against changes in log task energy. A pure price change follows slope minus one-half, whereas a pure technology change follows slope plus one-half. Model markers include a mixed scarcity-and-technology change on the price-dominant side.},width=0.96\linewidth]{figures/fig4_sign_test.pdf}
\caption{Conditional sign restriction for the budget price and technology. Changes in log imprecision are plotted against changes in log task energy. A pure change in p at fixed e and K moves the measures in opposite directions with slope \ensuremath{-}\ensuremath{\frac{1}{2}}; a pure technology change moves them together with slope +\ensuremath{\frac{1}{2}}. The markers show model comparative statics, including a mixed case in which a doubling of p outweighs lower energy per signal and capacity. The mapping identifies the combined technology index T; additional measurements of e and K resolve its components.}\label{fig:figure4}
\end{figure}

\subsection{Simulation-based design assessment}

The behavioral half of the test can be assessed by simulation, with the behavioral layer calibrated to the published numerosity tasks and a within-subject design with two states (Methods). Rejection rates rise with the size of the price change and with the number of participants (Figure 5): in the discrimination task, a doubling of the price, which raises imprecision by about a fifth, is detected in nearly all simulated studies with 40 participants, whereas a rise of a quarter requires substantially larger samples. The simulations also expose the main threat to the inference. When the rate of lapses changes between states, a fit that assumes a common lapse rate attributes the change to imprecision and reports a price change that did not occur, whereas a fit with state-specific lapse rates removes this bias at a cost in power, so that lapses must be measured state by state. Rejection rates, Monte Carlo intervals and fit diagnostics are reported in Supplementary Section S9.4.

\begin{figure}[H]\centering
\includegraphics[alt={Simulation chart of rejection rates across effective price ratios, sample sizes, and task designs. Rejection rises with larger price changes and larger samples under the stated design assumptions. Error bars are 95 percent Wilson intervals; horizontal references mark the nominal 0.05 test size and 0.80 rejection rate.},width=0.96\linewidth]{figures/fig5_power.pdf}
\caption{Simulated rejection rates for detecting a state-dependent change in the effective per-signal price in within-subject designs. Discrimination curves report the one-sided t-test on fitted log f with 200 trials per state; effective per-signal price ratios of 1, 1.25, 1.5 and 2 correspond to the labeled common factors in imprecision. The star marks the distinct estimation design using a one-sided signed-rank test, 120 trials per state and a doubled effective per-signal price. Error bars are 95\% Wilson intervals from 200 replications for discrimination and 100 for estimation. Horizontal lines mark the nominal 0.05 test size and 0.80 rejection rate; lapse rates are held constant across these main design simulations.}\label{fig:figure5}
\end{figure}

\subsection{Matched-numerosity response variability}

The encoding layer can be confronted with the published estimation data of \citet{pratcarrabin2026}, released as 12,960 trials from 36 participants \citep{pratcarrabin_data} (Methods), whose reconstructed standard deviations reproduce the published summaries and thereby anchor the calibration of the behavioral layer to the raw trials. Holding numerosity fixed at the 21 levels that all three priors include, residual response variance rises with prior width, with median ratios of 2.86 for widths 40 and 20 and 4.25 for widths 60 and 20; the pooled ratio exceeds one at all 21 numerosities in both comparisons with width 20 and at 19 of the 21 for widths 60 and 40 (Figure 6). Participant-cluster bootstrap intervals and multiplicity-adjusted p-values are reported in the Supplementary Materials as nominal diagnostics, since a calibration pilot under a fitted null indicates that the procedure is liberal (Supplementary Section S10). The calibrated encoding layer, whose two parameters were fitted only at numerosity 60, predicts these ratios out of sample (2.73, 4.57 and 1.68), a descriptive comparison, since the calibration targets condition-level variability while the ratios use within-participant residuals. The comparison checks the encoding component on which the price operates and leaves the price, neural energy and the budget untested, and, because prior width is linked to block position in the published design, it does not identify a causal effect of width; the inferential apparatus and its sensitivity analyses are reported in Supplementary Sections S6 to S8.

\begin{figure}[H]\centering
\includegraphics[alt={Three-panel plot of pooled residual-variance ratios for prior-width contrasts 40 over 20, 60 over 20, and 60 over 40 across 21 shared numerosities. The equality line marks a ratio of one. Navy points show the observed estimates at each numerosity; rust curves show the model-implied width-scaling profile calibrated at numerosity 60 and displayed across the shared range. Panel subtitles summarize each contrast's median estimate and number of numerosity-specific estimates above one.},width=0.96\linewidth]{figures/fig6_empirical.pdf}
\caption{Matched-numerosity response variability in the published estimation cohort. Panels A--C show pooled residual-variance ratios for prior-width pairs 40/20, 60/20, and 60/40 at the 21 shared numerosities. Navy points are observed ratios after centering within participant \ensuremath{\times} width \ensuremath{\times} numerosity cells; the dashed vertical reference marks equal residual variance. Rust curves show the model-implied width-scaling profile, calibrated to published condition-level standard-deviation targets at numerosity 60 and displayed across the shared range. Each panel reports the observed median and corresponding model-profile median; a second subtitle gives the number of numerosity-specific observed point estimates above one: 21/21 for 40/20 and 60/20, and 19/21 for 60/40.}\label{fig:figure6}
\end{figure}

\section{Discussion}

The model connects a marginal physiological price to the precision of an internal representation, and it shows that this price can be high while the energy that a task draws, and the whole-brain increment that imaging records, remain small. The constancy of total use that underlies the reading of cognition as cheap \citep{kondrakiewicz2025} is, in this account, the condition that makes the price bind: a computation costs the energy that it displaces from competing uses, valued at the margin, and the small task-evoked increment \citep{jamadar2025} reflects the reallocation of energy within a budget held near its set point. The exact fourth-root restriction applies when coding technology is fixed and tasks face the same price ratio, and the equilibrium analysis states when a common factor is a useful benchmark and when finite task demand requires task-specific prices.

Several rival accounts bear on these predictions. An account in which cognition is metabolically cheap predicts no metabolic modulation of precision, whereas the model predicts a common factor on imprecision across tasks and priors whenever the energy margin changes, and an account with a fixed cost of precision fitted to each task predicts no restriction linking the state effects of different tasks. Opportunity-cost accounts of effort \citep{kurzban2013} are nested in the model, since the shadow price equals the marginal value of energy in competing uses, so that opportunity cost gives the cost its form and metabolism gives it its currency; the two readings separate under manipulations of supply that leave the value of alternative activities unchanged. Limits on multitasking that reflect a trade-off between learning efficacy and processing efficiency in neural architectures \citep{musslick2021} can be represented as constraints on the capacity K, and the sign test separates changes in capacity, which move imprecision and energy together, from changes in the price, which move them in opposite directions. Accounts based on the depletion of energetic reserves \citep{christie2015} concern dynamics within a session, which the static model leaves aside, and the adaptation of coding technology under scarcity \citep{padamsey2022} enters through the energy per signal and the capacity; because technology responds to the price over longer timescales than the price itself, prolonged scarcity combines the two changes, and the sign test still classifies a change of state as a change in the price whenever the price change dominates.

A metabolic account of the cost of cognition must also be kept apart from the proposal that self-control draws on blood glucose as a limited fuel \citep{gailliot2007}, whose support has eroded: the depletion effect on which it rested has not survived a preregistered multilaboratory replication \citep{hagger2016}, the evidence for the glucose model has limited evidential value \citep{vadillo2016}, and neurophysiological evidence makes it unlikely that self-control tasks lower glucose relative to control tasks \citep{kurzban2010}. In the present model, the cost of a computation is a price set at the margin of a regulated budget, and neither blood glucose nor the level of a reserve enters it directly. An ingested load of glucose moves the price only insofar as it widens the margin of a budget that supply limitation and regulation hold close to its set point, and the fourth-root law then attenuates whatever change in the price occurs: in the illustrative baseline, a tightening of the margin by 30\% multiplies imprecision by only 1.06. Small and inconsistent behavioral effects of glucose manipulations are therefore what the model predicts, and they bear on the difference between a fuel that is depleted and a price that is set at the margin.

For economics, the model offers a physical anchor for the cost of precision. A change of state rescales the cost of information by the ratio of prices while preserving the shape of the cost function, a restriction that rational-inattention experiments run across physiological states can test \citep{dean2023}, and which bears on the variety of resource constraints that resource-rational models assume \citep{ma2020}. Since imprecise representations have been proposed as a source of small-stakes risk aversion \citep{khaw2021}, a state-dependent price of precision implies state-dependent risk attitudes of a specific form, which connects the energetics of the brain to the economics of choice under risk.

The reanalysis of the published cohort characterizes an established pattern, since the source study already reports unequal response variance across prior conditions \citep{pratcarrabin2026}, at a finer resolution: it holds numerosity fixed, respects the repeated-measures structure of the design, and sets the observed ratios against the predictions of a calibration that used a single numerosity. The agreement, which is descriptive, is consistent with the encoding component on which the price operates, while the price itself and its link to the budget remain to be measured. The design simulations quantify the detection of a price change under specified trial counts, heterogeneity, lapse processes, fitting rules and tests, and they make state-specific lapse measurement a consequential design choice, since a change in lapses can otherwise mimic a change in the price.

The model can fail in three ways, each of which existing paradigms can test. It fails if a change of state that leaves coding technology and stakes unchanged alters imprecision by different factors in tasks that draw on the same budget, since a common price must rescale them alike. It fails if imprecision and stimulus-linked task energy move together under a manipulation of supply that leaves coding technology intact, since a pure change in the price moves them in opposite directions. And it fails if, with technology and stakes fixed, task-evoked energy follows the reallocation margin while precision remains unchanged, since in the model the margin reaches behavior only through the price that it sets.

The same restriction can locate the boundary of the shared budget, which the model leaves open when it allows prices to be set region by region. The common factor should hold for tasks that share an encoding population, such as tasks that rely on the parietal encoding of numbers \citep{pratcarrabin2025}, and it should weaken as tasks draw on more distant circuits whose energy is supplied and regulated separately, so that the pattern of state factors across tasks would trace the spatial scale at which the brain prices energy, and the point at which the restriction breaks down would mark the boundary of the market.

Some implications are suggested by the model without being shown by it. The temporal variability of cerebral glucose use, which tracks cognition more closely than its baseline rate and declines with age \citep{deery2026}, is a candidate measure of the reallocation margin; signals of metabolic state that regulate the complexity of behavioral policies \citep{tesmer2025} are candidate carriers of the price; the conservation of energy that the aging brain achieves by suppressing low-priority processes \citep{shaulson2024} implies a rising price with age, and therefore a common rise of imprecision across tasks in older adults unless coding technology changes as well; and predictable input, which lowers the energetic cost of cortical processing \citep{hechler2023}, would read in the model as a change in technology.

The model and the analyses have limits that bound these conclusions. The model is a static equilibrium that treats coding technology and task stakes as stable for its common-price comparison, and it does not represent within-session depletion or specify a neural signal that coordinates allocation. The committed budget, which the static model takes as given within each physiological state, is itself endogenous on the slow timescales of development and evolution, since mammalian connectomes accept higher wiring costs in exchange for computational capability and robustness to damage \citep{dendorfer2026}. A costlier architecture commits more of the budget, which narrows the margin and raises \ensuremath{\mu}, while it can lower the energy per signal by shortening the paths that messages travel, so that the sign of its net effect on the price of precision, \ensuremath{\lambda} = e\ensuremath{\mu}/\ensuremath{\beta}, remains an open question for later work. The human analysis is a secondary reanalysis of a published cohort, without preregistration or an independent validation sample; observed response variance does not isolate latent Fisher precision or neural energy, prior width is linked to block position in the observed sequences, and in a pilot of 100 simulated cohorts under a fitted Gaussian null the bootstrap procedure rejected at least one of the 63 comparisons in 22\% of cohorts (Wilson 95\% interval 15.0\% to 31.1\%) against a nominal 5\%, so that its intervals and adjusted p-values serve as nominal diagnostics. For imaging designs, glucose uptake can diverge from oxidative cost, because activation increases glycolysis beyond oxidation and lactate can leave the activated tissue \citep{dienel2025}, and PET integrates activity over minutes. The design simulations are conditional on the specified sample sizes, lapse processes and fitting choices, and frequent boundary estimates under state-specific lapse fits limit the generality of the displayed operating characteristics.

These limits define the next empirical step, which measures behavior and stimulus-linked neural energy in the same participants and states. Such a design would vary the available margin or the competing demand, estimate displacement with independently calibrated control conditions, fit state-specific lapse and motor-noise components, and repeat states across tasks, so as to test whether the predicted common price factor transfers across tasks, whether the sign restriction survives the observed displacement, and whether physiology constrains the price separately from the energy per signal, the task stakes and the coding capacity. Within its limits, the model provides a common variable, the shadow price of energy at the margin, through which the neuroscience of brain energy and the economics of imprecise choice can be tested against each other.

\section{Methods}

Our methods combine closed-form derivations with independent numerical verification, a participant-clustered reanalysis of the public trial record, and prospective design simulations. This separation makes each prediction traceable to its assumptions and keeps theoretical implications, human estimates, and simulated operating characteristics distinct.

\subsection{Derivations of the encoding results}

For Lemma 1, set s(x) = I\textsubscript{1}(x)\textsuperscript{\ensuremath{\frac{1}{2}}}. At fixed n, the objective becomes (1/n)\ensuremath{\int}\ensuremath{\pi}(x)\textsuperscript{a}/s(x)\textsuperscript{2}dx under \ensuremath{\int}s(x)dx \ensuremath{\leq} K. Pointwise minimization of \ensuremath{\pi}(x)\textsuperscript{a}/s(x)\textsuperscript{2} + \ensuremath{\nu}s(x), where \ensuremath{\nu} is the multiplier on the information-allocation bound, gives s(x) \ensuremath{\propto} \ensuremath{\pi}(x)\textsuperscript{a/3}; the positive second derivative confirms a minimum. Normalizing to exhaust K yields s(x) = K\ensuremath{\pi}(x)\textsuperscript{a/3}/Z\textsubscript{a} and loss Z\textsubscript{a}\textsuperscript{3}/(nK\textsuperscript{2}). Thus the optimal shape depends on the prior, while n scales total precision without changing that shape.

For Proposition 1, minimize Z\textsubscript{a}\textsuperscript{3}/(nK\textsuperscript{2}) + \ensuremath{\lambda}n over n > 0. The first-order condition gives n\textsuperscript{\ensuremath{\ast}} = Z\textsubscript{a}\textsuperscript{3/2}/(K\ensuremath{\lambda}\textsuperscript{\ensuremath{\frac{1}{2}}}), at which the allocation loss and signal cost are equal. Substitution yields I(x) = n\textsuperscript{\ensuremath{\ast}}s(x)\textsuperscript{2} = KZ\textsubscript{a}\textsuperscript{\ensuremath{-}\ensuremath{\frac{1}{2}}}\ensuremath{\lambda}\textsuperscript{\ensuremath{-}\ensuremath{\frac{1}{2}}}\ensuremath{\pi}(x)\textsuperscript{2a/3} and \ensuremath{\sigma}(x) = Z\textsubscript{a}\textsuperscript{\ensuremath{\frac{1}{4}}}\ensuremath{\lambda}\textsuperscript{\ensuremath{\frac{1}{4}}}K\textsuperscript{\ensuremath{-}\ensuremath{\frac{1}{2}}}\ensuremath{\pi}(x)\textsuperscript{\ensuremath{-}a/3}. Under a uniform prior of width w, Z\textsubscript{a} = w\textsuperscript{1\ensuremath{-}a/3}, so \ensuremath{\sigma} \ensuremath{\propto} w\textsuperscript{(1+a)/4} and n\textsuperscript{\ensuremath{\ast}} \ensuremath{\propto} w\textsuperscript{(3\ensuremath{-}a)/2}. These expressions make the task-specific width scaling and common price elasticity explicit. The derivation therefore isolates a common price response while retaining the distinct way each task uses prior information.

The closed-form optimum separates the task-specific prior shape from the common response to price:

\[\begin{aligned} n^\ast &= \frac{Z_{a}^{3/2}}{K\lambda^{1/2}},\\ \sigma(x) &= Z_{a}^{1/4}\lambda^{1/4}K^{-1/2}\pi(x)^{-a/3}. \end{aligned}\tag{1}\]

The paired expressions in (1) show why the same fourth-root price response applies across prior distributions while the prior continues to determine the shape of imprecision across stimuli.

Proposition 2 follows because log \ensuremath{\sigma} is affine in log \ensuremath{\lambda} with coefficient \ensuremath{\frac{1}{4}} at each x, and \ensuremath{\lambda} = e p, where p = \ensuremath{\mu}/\ensuremath{\beta} is the budget price per unit loss. The cross-state fourth-root comparison maps to a \ensuremath{\mu} ratio only when e, K and \ensuremath{\beta} are held fixed; with stable e, the \ensuremath{\lambda} ratio equals the p ratio, and with stable \ensuremath{\beta}, the p ratio equals the \ensuremath{\mu} ratio. Under the generalized bound \ensuremath{\int}I\textsubscript{1}\textsuperscript{\ensuremath{\gamma}/2}dx \ensuremath{\leq} K, the optimal allocation still separates from n and the optimized loss remains proportional to 1/n; hence n\textsuperscript{\ensuremath{\ast}} \ensuremath{\propto} \ensuremath{\lambda}\textsuperscript{\ensuremath{-}\ensuremath{\frac{1}{2}}} and \ensuremath{\sigma} \ensuremath{\propto} \ensuremath{\lambda}\textsuperscript{\ensuremath{\frac{1}{4}}}, although the exponents in w change. A convex signal cost \ensuremath{\lambda}n\textsuperscript{\ensuremath{\kappa}} instead gives n\textsuperscript{\ensuremath{\ast}} \ensuremath{\propto} \ensuremath{\lambda}\textsuperscript{\ensuremath{-}1/(\ensuremath{\kappa}+1)} and \ensuremath{\sigma} \ensuremath{\propto} \ensuremath{\lambda}\textsuperscript{1/(2(\ensuremath{\kappa}+1))}. The discrete-count calculation uses the baseline specification \ensuremath{\gamma} = \ensuremath{\kappa} = 1: for integer n, it minimizes G/n + \ensuremath{\lambda}n over n \ensuremath{\in} \{1,2,\ldots{}\}, with G = Z\textsubscript{a}\textsuperscript{3}/K\textsuperscript{2}. Comparing adjacent integer costs yields intervals in which the optimal count changes; the reported 3--10 range is computed by evaluating these exact discrete optima on a fine grid of continuous optima.

For Proposition 3, multiplying \(n^\ast\) by \(e\) gives \(E^\ast \propto e^{1/2}\mu^{-1/2}K^{-1}\) when \(\beta\) is fixed. In loss units, \(L^\ast=\lambda n^\ast\) with \(\lambda=e\mu/\beta\); equivalently, \(\beta L^\ast=\mu E^\ast\). Under the normalization \(\beta=1\), this reduces to \(L^\ast=\lambda n^\ast=\mu E^\ast\). Together, these expressions bridge prior-weighted precision, task energy, and valued loss, allowing the budget mechanism to determine a behavioral price rather than treating that price as an external input.

\subsection{Derivations of the budget results}

At each state, the finite-demand allocation solves the joint problem \(\max_{n>0,\,E_{o}>0,\,d>0}\{V(E_{o})-\beta L_{a}(n)-R(d):\ en+E_{o}=M+d\}\), with \(e,K,\beta,c,v,k,\rho>0\) and \(R'(d)=cd^{k}\). At an interior solution, the first-order conditions are \(\beta[-L_{a}'(n)]/e=V'(E_{o})=R'(d)=\mu\), so the per-signal price is \(\lambda=e\mu/\beta\). The equilibrium price is therefore determined jointly with task demand; the price-taking result in Lemma 2 is the limiting case as this task's demand becomes negligible relative to the rest of the budget.

For Proposition 4, write d(\ensuremath{\mu}) = (\ensuremath{\mu}/c)\textsuperscript{1/k}, E(\ensuremath{\mu}) \ensuremath{\propto} \ensuremath{\mu}\textsuperscript{\ensuremath{-}\ensuremath{\frac{1}{2}}}, and E\textsubscript{o}(\ensuremath{\mu}) = (v/\ensuremath{\mu})\textsuperscript{1/\ensuremath{\rho}}. The equilibrium condition and its derivative are:

\[\begin{aligned} F(\mu) &= d(\mu)+M-E(\mu)-E_{o}(\mu)=0,\\ F'(\mu) &= \frac{d(\mu)}{k\mu}+\frac{E(\mu)}{2\mu}+\frac{E_{o}(\mu)}{\rho\mu}>0. \end{aligned}\tag{2}\] Strict monotonicity in (2) gives at most one equilibrium price; for finite c it tends to opposite signs at the positive-price boundaries, establishing existence and uniqueness. Implicit differentiation gives \ensuremath{\partial}\ensuremath{\mu}/\ensuremath{\partial}M = \ensuremath{-}1/F\ensuremath{^\prime} < 0 and positive comparative statics with respect to v and c. Under the hard cap d = 0, the residual tends to M at large \ensuremath{\mu}, so a finite equilibrium requires M > 0. This monotonicity is the analytical basis for the budget curves in Figure 3.

For Lemma 2, scale the task demand by s so the equilibrium equation contains sE(\ensuremath{\mu}). Implicit differentiation at s = 1 gives \ensuremath{\partial}\ensuremath{\mu}/\ensuremath{\partial}s = E/F\ensuremath{^\prime}, and the corresponding elasticity is (E/\ensuremath{\mu})/F\ensuremath{^\prime}. As task demand becomes negligible, this elasticity tends to zero: the budget sets the price, while a small task takes it as given.

For Lemma 3, introduce task demand gradually as \ensuremath{\varepsilon}E\textsuperscript{u}(\ensuremath{\mu}), where E\textsuperscript{u} is task energy at unit scale, and denote the no-task equilibrium by \ensuremath{\mu}\textsubscript{0}. At that baseline, d\ensuremath{\mu}/d\ensuremath{\varepsilon} = E\textsuperscript{u}(\ensuremath{\mu}\textsubscript{0})/\(F'_0\), where \(F'_0\) = d(\ensuremath{\mu}\textsubscript{0})/(k\ensuremath{\mu}\textsubscript{0}) + E\textsubscript{o}(\ensuremath{\mu}\textsubscript{0})/(\ensuremath{\rho}\ensuremath{\mu}\textsubscript{0}). Since d\ensuremath{\Delta}B/d\ensuremath{\varepsilon} = d\ensuremath{^\prime}(\ensuremath{\mu}\textsubscript{0})d\ensuremath{\mu}/d\ensuremath{\varepsilon}, the first-order expansion and its allocation share are:

\[\begin{aligned} \Delta B &= \phi_0\varepsilon E^{u}(\mu_0)+o(\varepsilon),\\ \phi_0 &= \frac{d(\mu_0)/k}{d(\mu_0)/k+E_{o}(\mu_0)/\rho}. \end{aligned}\tag{3}\]

The coefficient in (3), \ensuremath{\phi}\textsubscript{0}, measures the local share of an infinitesimal task-energy increase that remains in total use after competing demand adjusts. For a local comparison of no-task states differing through M alone, holding c, v, k, \ensuremath{\rho}, e, K, task stakes and task scale fixed, \ensuremath{\phi}\textsubscript{0} has elasticity (1\ensuremath{-}\ensuremath{\phi}\textsubscript{0})(1/k + 1/\ensuremath{\rho}) with respect to \ensuremath{\mu}\textsubscript{0}. Because E\textsuperscript{u}(\ensuremath{\mu}\textsubscript{0}) \ensuremath{\propto} \ensuremath{\mu}\textsubscript{0}\textsuperscript{\ensuremath{-}\ensuremath{\frac{1}{2}}}, the first-order net increment has elasticity (1\ensuremath{-}\ensuremath{\phi}\textsubscript{0})(1/k + 1/\ensuremath{\rho}) \ensuremath{-} \ensuremath{\frac{1}{2}} and rises with \ensuremath{\mu}\textsubscript{0} under this comparison when (1\ensuremath{-}\ensuremath{\phi}\textsubscript{0})(1/k + 1/\ensuremath{\rho}) > \ensuremath{\frac{1}{2}}. In particular, \ensuremath{\phi}\textsubscript{0} < \ensuremath{\frac{1}{2}} and 1/k + 1/\ensuremath{\rho} \ensuremath{\geq} 1 provide a transparent sufficient parameter condition. The numerical baseline separately compares this no-task first-order expression with the exact finite-task equilibrium change.

For Proposition 5, normalize task-plus-competing use to one. For any target share \(s\in(0,1)\) and price \(\mu>0\), set \(e=1\), choose the task-scale primitive so that \(E(\mu)=s\), and set \(E_{o}=1-s\). Choose any \(d\in(0,1)\) and let \(M=1-d\); then \(d+M=E+E_{o}=1\). Choosing \(\bar B>M\) and \(C=\bar B-M\) keeps committed use positive, while \(c=\mu/d^{k}\) and \(v=\mu E_{o}^\rho\) give \(R'(d)=V'(E_{o})=\mu\). The resulting share of energy above committed use is \(E/(E+E_{o})=s\). This establishes the construction for the full interior domain; the grid of sixteen pairs checks its numerical implementation, with maximum relative error 2.22 \ensuremath{\times} 10\textsuperscript{-16}. Varying \(c\) while adjusting \(v\) to hold \(\mu\) fixed shows that the net-to-gross increment ratio is not uniquely determined by price. The construction makes precise why the task share and net increment are complementary observations rather than substitutes for the joint price-and-precision restrictions.

For the joint non-identification claim, take \(k=\rho=1\) and hold the task-demand coefficient \(A\) fixed, so \(E(\mu)=A\mu^{-1/2}\). Choose a common \(\Delta>0\) and two distinct prices \(\mu_j\) for which \(E_{j}=A\mu_j^{-1/2}>2\Delta\). Set \(r_j=E_{j}/(2E_j-\Delta)\), \(\mu_{0j}=r_j\mu_j\), \(v_j=\mu_jE_j\), \(c_j=\mu_j(1-r_j)/\Delta\), and \(M_{j}=E_{j}/r_j-r_j\Delta/(1-r_j)\). These primitives are positive. They imply \(E_{o}(\mu_j)=E_{j}\) and \(E_{o}(\mu_{0j})=E_{j}/r_j=2E_j-\Delta\), so each task share is \(1/2\); they also imply \(d(\mu_j)-d(\mu_{0j})=\Delta\). The two distinct prices therefore generate the same task share and net increment with the same task technology \(A\), while the budget primitives differ. This construction clarifies why aggregate shares and net increments are complementary observations, while the joint behavioral-energy restrictions carry cross-state information about price.

\subsection{The sign test, measurement, and numerical verification}

For each fixed task specification across states, define the budget price per unit loss \(p=\mu/\beta\), so that \(\lambda=ep\) is the effective price per signal, and let \(E=en^*\) denote gross encoding energy. Taking log changes in \(\sigma\propto(ep)^{1/4}K^{-1/2}\) and \(E\propto e^{1/2}p^{-1/2}K^{-1}\) gives the joint log-change restriction:

\[\begin{aligned} \Delta\log\sigma &= \frac{x+y}{4}-\frac{z}{2}, & \Delta\log E &= \frac{y-x}{2}-z,\\ \frac{\Delta\log\sigma}{\Delta\log E} &= \frac{x+T}{2(T-x)}, & \Delta\log(\sigma^{2}E) &= T. \end{aligned}\tag{4}\]

Here T = y\ensuremath{-}2z is the combined technology index. The joint restriction in (4) defines the finite slope only when \ensuremath{\Delta}logE \ensuremath{\neq} 0 (T \ensuremath{\neq} x); when T = x \ensuremath{\neq} 0, gross energy is unchanged while imprecision changes, and when x = T = 0 both measures are unchanged. A pure budget-price change has T = 0 and slope \ensuremath{-}\ensuremath{\frac{1}{2}}; a pure technology change has x = 0 and slope +\ensuremath{\frac{1}{2}}. The pair of observables therefore identifies the change in budget price p and the combined index T under the model, but not \ensuremath{\mu} separately when \ensuremath{\beta} changes, nor e and K separately. At fixed p, offsetting technology changes with T = \ensuremath{\Delta}log(e/K\textsuperscript{2}) = 0 leave both \ensuremath{\sigma} and E unchanged; these outcomes alone therefore cannot distinguish them from unchanged technology. Price claims about \ensuremath{\mu} consequently require stable or measured stakes.

For regional measurement, let the observed task-evoked quantity be E\ensuremath{-}D, with D representing displaced activity and \ensuremath{\delta} = dD/dE. Differentiation gives dlog(E\ensuremath{-}D) = (1\ensuremath{-}\ensuremath{\delta})dE/(E\ensuremath{-}D), which rescales the theoretical slope by (E\ensuremath{-}D)/(E(1\ensuremath{-}\ensuremath{\delta})). The numerical illustration uses a price doubling with baseline displacement D\textsubscript{0} = 0.6E\textsubscript{0}; all scenarios are checked to keep D and E\ensuremath{-}D nonnegative. With proportional displacement, the pure-price slope remains \ensuremath{-}\ensuremath{\frac{1}{2}}; fixed or state-changing displacement alters its magnitude and can reverse its sign if the observed increment changes in the opposite direction. Because both measures have error, the empirical slope should be estimated with an errors-in-variables method, with the sign as the primary test. The joint design thus uses the directional restriction as its clearest diagnostic and the additional magnitude relation to sharpen price comparisons.

The illustrative baseline, in normalized units with the set point equal to one, sets the reallocation margin at M = 0.05, the stiffness of regulation at c = 100 with k = 1, the value of competing uses at v = 0.05 with curvature \ensuremath{\rho} = 1, the energy per signal at e = 1, and the task's scale Z\textsubscript{a}\textsuperscript{3/2}/K at 0.002. Task stakes are normalized to \ensuremath{\beta} = 1 for these illustrative calculations, so the budget price per unit loss is p = \ensuremath{\mu} and the effective price per signal is \ensuremath{\lambda} = e\ensuremath{\mu}. The equilibrium price is found by bisection on the logarithm of the price, which agrees with an independent root finder within a relative error of 10\textsuperscript{-13} over 300 random parameter draws that include extreme convexity, curvature and stiffness. Elasticities are computed by central differences, and the exact net increment of Lemma 3 by solving the budget with and without the task.

An independent grid optimizer that does not use the closed forms evaluates 160 points from 10 to 90, with K = 1 and per-signal prices 0.001 and 0.002. It compares the optimum with the analytical solution for uniform, log-uniform and bimodal priors and reproduces the closed form within relative error 10\textsuperscript{-6}; the common-factor identity is reproduced within 10\textsuperscript{-6}. Symbolic checks and invariant tests cover the elasticity formulas, budget sign conditions, displacement feasibility and integer-count cases. The agreement between independent root-finding and grid-based calculations anchors the reported comparative statics in numerically distinct verification routes, strengthening the numerical basis for the displayed equilibrium and encoding predictions.

\subsection{Behavioral model, calibration and power}

The behavioral component follows the numerosity estimation and discrimination tasks reported by Prat-Carrabin and Woodford. In estimation, integer stimuli are drawn from the task's prior range, decoded as posterior means under a uniform prior, and combined with truncated motor noise before rounding. In discrimination, each of two colors contains five integers drawn from the prior range; observers compare noisy representations of the two sample averages, with lapses choosing either color at random. Component calibration uses the published summary targets. The secondary estimation reanalysis uses the public trial-level file from the same 36-person cohort (12,960 trials), available from the OSF project cited under Data and Code Availability. At numerosity 60, the pooled residual SD subtracts the condition-level mean response to reconstruct the published calibration anchor. Matched-numerosity comparisons use the 21 numerosities shared by prior widths 20, 40 and 60 (50--70 inclusive). Within each participant \ensuremath{\times} width \ensuremath{\times} numerosity cell with at least two trials, responses are centered on the cell mean. The pooled residual-variance estimator at numerosity m and prior width w is:

\[\widehat V_{wm}=\frac{\displaystyle\sum_{i:n_{iwm}\ge 2}\sum_{t=1}^{n_{iwm}}\left(Y_{iwmt}-\bar Y_{iwm}\right)^{2}}{\displaystyle\sum_{i:n_{iwm}\ge 2}(n_{iwm}-1)}\tag{5}.\]

This pooled estimator centers responses within participant before combining residual sums of squares and degrees of freedom, so each available cell contributes in proportion to its residual degrees of freedom. Because released trial counts vary across cells, paired participant support can vary by prior-width contrast and numerosity. At each numerosity, the primary descriptive profile compares all three prior-width pairs using the pooled ratios above. A participant-cluster bootstrap additionally summarizes log-ratio variation: each draw resamples participants separately within the two recorded block-order strata, 20\ensuremath{\rightarrow}40\ensuremath{\rightarrow}60 (n = 20) and 60\ensuremath{\rightarrow}40\ensuremath{\rightarrow}20 (n = 16), carrying each selected participant's cells across widths. The defined family contains 63 numerosity-by-width contrasts, with 500000 draws per contrast; the supplement reports the nominal procedure-based Bonferroni outputs and Wilson Monte Carlo intervals for its tail fractions. A complementary post hoc participant-level directional analysis pools each participant's available within-cell residual sums of squares and degrees of freedom over numerosity 50--70, then forms one wider/narrower variance ratio per participant. A two-sided exact binomial sign test evaluates equal positive and negative direction counts, with Bonferroni adjustment across the three width contrasts. The level-specific sign sensitivity instead compares eligible participants' sample variances at each numerosity and width pair, requires at least two observations in both cells, retains one-zero pairs as directional signs, and records equal variances as ties; its 63 tests are adjusted as a separate family. A participant-equal sensitivity averages participant log ratios and is reported once in Supplementary Table S3 alongside the fixed-numerosity analysis. These complementary summaries make the matched-level profile and participant-level direction directly inspectable. The participant is the resampling unit; the behavioral analyses use the same released cohort of 36 participants, with trials nested within participant. For estimation, \ensuremath{\nu}w\textsuperscript{\ensuremath{\frac{1}{2}}} with \ensuremath{\nu} = 1.06 and motor noise \ensuremath{\sigma}\textsubscript{0} = 3.02 reproduces the three reported standard deviations at numerosity 60: 4.27, 6.68 and 8.45, compared with 4.2, 6.8 and 8.4. The continuous-prior approximation is checked against the experiment's integer prior; refitting under that discrete prior changes \ensuremath{\nu} by 0.8\% (\ensuremath{\nu} = 1.07) and gives \ensuremath{\sigma}\textsubscript{0} = 2.80. In discrimination, \ensuremath{\nu} = 0.195 with width exponent \ensuremath{\frac{3}{4}} and lapse rate 0.144 jointly fit four reported psychometric points to within 0.01. The replication configuration records the seeds and simulation counts. Simulation settings, seeds and replication counts are fixed in the supplied configuration; the simulated outcomes evaluate the specified designs.

Each prospective design simulates two within-subject states. Participant heterogeneity has standard deviation 0.2 for log \ensuremath{\nu}, 0.3 for the lapse logit, and 0.03 for the true log state effect; estimation additionally varies log motor noise with standard deviation 0.2. Discrimination uses 200 trials per state over prior range 35--65, and tests participant-level log f with a one-sided t-test at \ensuremath{\alpha} = 0.05. Estimation uses 120 trials per state over 40--80, 40 participants and a one-sided Wilcoxon signed-rank test. The reported replications are 200 and 100, respectively. A separate robustness simulation multiplies second-state lapse odds by 1.5 under price ratios 1 and 2, tests designs with 40 and 80 participants, and fits shared versus state-specific lapse rates; it uses 200 replications per condition. In the discrimination power analysis, the N = 20 and N = 40 estimates use nested participant subsets from the N = 80 sample within each replication, so rejection rates across sample sizes are paired summaries rather than independent simulation batches.

All subject fits use L-BFGS-B with bounds matched to the likelihood parameterization: log \ensuremath{\nu} in [\ensuremath{-}12, 12], logit lapse in [\ensuremath{-}15, 15], log f in [\ensuremath{-}5, 5], and log \ensuremath{\sigma}\textsubscript{0} in [\ensuremath{-}12, 12]. The optimizer uses a 2,000-iteration ceiling, ftol 10\textsuperscript{-12} and gtol 10\textsuperscript{-7}. Success and boundary contact are recorded for every fit. Rejection-rate uncertainty is summarized with two-sided 95\% Wilson intervals and Monte Carlo standard errors, including estimates at zero or one. The discrimination campaign uses a seeded generator; estimation uses independently seeded chunks of ten replications, with a final partial chunk retained when the requested total is not divisible by ten. Cache names fingerprint the configuration, calibration, code and numerical-library environment; cached chunk lengths are checked before reuse.

The calibrated estimation model's ratios are computed at each shared numerosity by Gauss--Hermite quadrature over the representation, using the posterior-mean decoder under the uniform prior, truncated motor noise, and the model's rounding step (\texttt{model/predicted\_variance\_ratios.py}). Parameters are fitted at numerosity 60; the model profile is then compared with the 20 additional shared levels without refitting, giving a transparent out-of-calibration-numerosity benchmark within the released cohort.

Reproducibility is supported by the model source, configuration, symbolic derivations, tests, figure scripts and generated results. From the \texttt{model} directory, \texttt{python run\_all.py} regenerates the analytic checks, calibrated and simulated model results, the primary participant-clustered reanalysis, and the two model figures. The complete release workflow is \texttt{reproduce.ps1} on Windows or \texttt{reproduce.sh} on Unix-like systems: it additionally runs the pair-specific complete-case and participant-sign analyses, regenerates the manuscript figures and filled manuscript sources, compiles the main and supplementary TeX/PDF files, and runs the model test suite. The separation of these commands makes each analysis output identifiable while allowing the complete sequence to be regenerated in one workflow. Estimation-recovery chunks and the slower trial-level bootstrap are reused only when their fingerprints and requested replication counts match the current inputs and configuration. The model is released under the MIT license. Together, the analytical results, numerical checks, behavioral reanalysis and design projections remain separately traceable to their assumptions and inputs, while the workflow supports end-to-end reproduction.

\section*{Data and code availability}

The estimation reanalysis uses trial-level data released in the Open Science Framework project of Prat-Carrabin and Woodford (2026), cited in the references. The raw CSV is not redistributed because file-level reuse terms could not be independently confirmed; readers can obtain it from the OSF project and place it at \texttt{data/published\_estimation.csv} within the replication package, which records the SHA-256 checksum against which the file should be verified. The replication package, available at \url{https://pierluigisacco.eu/wp-content/uploads/2026/10/The_marginal_brain_replication.zip}, includes the analysis code, configuration, derived results and figure-generation scripts, and regenerates every number and figure in the manuscript. Behavioral calibration also uses the published summary values.

\section*{Competing interests}

The authors declare no competing interests.

\section*{Funding}

This work received no specific funding.

\section*{Author contributions}

Fabio Lokwani-Di Matteo: formal analysis, software, validation, writing -- original draft, writing -- review and editing. Pier Luigi Sacco: conceptualization, methodology, formal analysis, writing -- original draft, writing -- review and editing.

\begingroup
\footnotesize
\setlength{\parskip}{0pt}
\begin{thebibliography}{99}
\setlength{\itemsep}{0pt}
\setlength{\parsep}{0pt}
\bibitem[Attwell and Laughlin(2001)]{attwell2001} Attwell, D., Laughlin, S. B. (2001). \emph{An energy budget for signaling in the grey matter of the brain}. Journal of Cerebral Blood Flow \& Metabolism, 21(10), 1133--1145. DOI: \href{https://doi.org/10.1097/00004647-200110000-00001}{10.1097/00004647-200110000-00001}.
\bibitem[Balasubramanian et al.(2001)]{balasubramanian2001} Balasubramanian, V., Kimber, D., Berry, M. J., II (2001). \emph{Metabolically efficient information processing}. Neural Computation, 13(4), 799--815. DOI: \href{https://doi.org/10.1162/089976601300014358}{10.1162/089976601300014358}.
\bibitem[Bruckmaier et al.(2020)]{bruckmaier2020} Bruckmaier, M., Tachtsidis, I., Phan, P., Lavie, N. (2020). \emph{Attention and capacity limits in perception: A cellular metabolism account}. The Journal of Neuroscience, 40(35), 6801--6811. DOI: \href{https://doi.org/10.1523/jneurosci.2368-19.2020}{10.1523/jneurosci.2368-19.2020}.
\bibitem[Christie and Schrater(2015)]{christie2015} Christie, S. T., Schrater, P. (2015). \emph{Cognitive cost as dynamic allocation of energetic resources}. Frontiers in Neuroscience, 9. DOI: \href{https://doi.org/10.3389/fnins.2015.00289}{10.3389/fnins.2015.00289}.
\bibitem[Dean and Neligh(2023)]{dean2023} Dean, M., Neligh, N. (2023). \emph{Experimental tests of rational inattention}. Journal of Political Economy, 131(12), 3415--3461. DOI: \href{https://doi.org/10.1086/725174}{10.1086/725174}.
\bibitem[Deery et al.(2026)]{deery2026} Deery, H. A., Liang, E., Moran, C., Egan, G. F., Jamadar, S. D. (2026). \emph{Brain glucodynamic variability is an essential feature of the metabolism--cognition relationship}. Proceedings of the National Academy of Sciences, 123(3), e2510850123. DOI: \href{https://doi.org/10.1073/pnas.2510850123}{10.1073/pnas.2510850123}.
\bibitem[Dendorfer et al.(2026)]{dendorfer2026} Dendorfer, A., Mousley, A., Páscoa dos Santos, F., Astle, D. E., Fakhar, K. (2026). \emph{Mapping the trade-offs that shape mammalian brain network architecture}. bioRxiv. DOI: \href{https://doi.org/10.64898/2026.10.07.757032}{10.64898/2026.10.07.757032}.
\bibitem[Dienel et al.(2025)]{dienel2025} Dienel, G. A., Rothman, D. L., Mangia, S. (2025). \emph{A bird's-eye view of glycolytic upregulation in activated brain: The major fate of lactate is release from activated tissue, not shuttling to nearby neurons}. Journal of Neurochemistry, 169(6), e70111. DOI: \href{https://doi.org/10.1111/jnc.70111}{10.1111/jnc.70111}.
\bibitem[Gailliot et al.(2007)]{gailliot2007} Gailliot, M. T., Baumeister, R. F., DeWall, C. N., Maner, J. K., Plant, E. A., Tice, D. M., Brewer, L. E., Schmeichel, B. J. (2007). \emph{Self-control relies on glucose as a limited energy source: Willpower is more than a metaphor}. Journal of Personality and Social Psychology, 92(2), 325--336. DOI: \href{https://doi.org/10.1037/0022-3514.92.2.325}{10.1037/0022-3514.92.2.325}.
\bibitem[Grujic et al.(2022)]{grujic2022} Grujic, N., Brus, J., Burdakov, D., Polan\'{i}a, R. (2022). \emph{Rational inattention in mice}. Science Advances, 8(9), eabj8935. DOI: \href{https://doi.org/10.1126/sciadv.abj8935}{10.1126/sciadv.abj8935}.
\bibitem[Gusnard and Raichle(2001)]{gusnard_raichle2001} Gusnard, D. A., Raichle, M. E. (2001). \emph{Searching for a baseline: Functional imaging and the resting human brain}. Nature Reviews Neuroscience, 2, 685--694. DOI: \href{https://doi.org/10.1038/35094500}{10.1038/35094500}.
\bibitem[Hagger et al.(2016)]{hagger2016} Hagger, M. S., Chatzisarantis, N. L. D., Alberts, H., Anggono, C. O., Batailler, C., Birt, A. R., Brand, R., Brandt, M. J., Brewer, G., Bruyneel, S., Calvillo, D. P., Campbell, W. K., Cannon, P. R., Carlucci, M., Carruth, N. P., Cheung, T., Crowell, A., De Ridder, D. T. D., Dewitte, S., Elson, M., Evans, J. R., Fay, B. A., Fennis, B. M., Finley, A., Francis, Z., Heise, E., Hoemann, H., Inzlicht, M., Koole, S. L., Koppel, L., Kroese, F., Lange, F., Lau, K., Lynch, B. P., Martijn, C., Merckelbach, H., Mills, N. V., Michirev, A., Miyake, A., Mosser, A. E., Muise, M., Muller, D., Muzi, M., Nalis, D., Nurwanti, R., Otgaar, H., Philipp, M. C., Primoceri, P., Rentzsch, K., Ringos, L., Schlinkert, C., Schmeichel, B. J., Schoch, S. F., Schrama, M., Schütz, A., Stamos, A., Tinghög, G., Ullrich, J., vanDellen, M., Wimbarti, S., Wolff, W., Yusainy, C., Zerhouni, O., Zwienenberg, M. (2016). \emph{A multilab preregistered replication of the ego-depletion effect}. Perspectives on Psychological Science, 11(4), 546--573. DOI: \href{https://doi.org/10.1177/1745691616652873}{10.1177/1745691616652873}.
\bibitem[Harris et al.(2012)]{harris2012} Harris, J. J., Jolivet, R., Attwell, D. (2012). \emph{Synaptic energy use and supply}. Neuron, 75(5), 762--777. DOI: \href{https://doi.org/10.1016/j.neuron.2012.08.019}{10.1016/j.neuron.2012.08.019}.
\bibitem[Haueis and Cola\c{c}o(2026a)]{haueis2026a} Haueis, P., Cola\c{c}o, D. (2026a). \emph{Giving metabolism its due}. Behavioral and Brain Sciences, 49, e486. DOI: \href{https://doi.org/10.1017/s0140525x26107298}{10.1017/s0140525x26107298}.
\bibitem[Haueis and Cola\c{c}o(2026b)]{haueis2026b} Haueis, P., Cola\c{c}o, D. (2026b). \emph{Metabolic considerations for cognitive modeling}. Behavioral and Brain Sciences, 49, e453. DOI: \href{https://doi.org/10.1017/s0140525x25103956}{10.1017/s0140525x25103956}.
\bibitem[Hechler et al.(2023)]{hechler2023} Hechler, A., de Lange, F. P., Riedl, V. (2023). \emph{The energy metabolic footprint of predictive processing in the human brain}. bioRxiv. DOI: \href{https://doi.org/10.1101/2023.12.08.570804}{10.1101/2023.12.08.570804}.
\bibitem[Herculano-Houzel(2011)]{herculanohouzel2011} Herculano-Houzel, S. (2011). \emph{Scaling of brain metabolism with a fixed energy budget per neuron: Implications for neuronal activity, plasticity and evolution}. PLoS ONE, 6(3), e17514. DOI: \href{https://doi.org/10.1371/journal.pone.0017514}{10.1371/journal.pone.0017514}.
\bibitem[Herculano-Houzel and Rothman(2022)]{herculanohouzel2022} Herculano-Houzel, S., Rothman, D. L. (2022). \emph{From a demand-based to a supply-limited framework of brain metabolism}. Frontiers in Integrative Neuroscience, 16, 818685. DOI: \href{https://doi.org/10.3389/fnint.2022.818685}{10.3389/fnint.2022.818685}.
\bibitem[Jamadar et al.(2025)]{jamadar2025} Jamadar, S. D., Behler, A., Deery, H., Breakspear, M. (2025). \emph{The metabolic costs of cognition}. Trends in Cognitive Sciences, 29(6), 541--555. DOI: \href{https://doi.org/10.1016/j.tics.2024.11.010}{10.1016/j.tics.2024.11.010}.
\bibitem[Khaw et al.(2021)]{khaw2021} Khaw, M. W., Li, Z., Woodford, M. (2021). \emph{Cognitive imprecision and small-stakes risk aversion}. The Review of Economic Studies, 88(4), 1979--2013. DOI: \href{https://doi.org/10.1093/restud/rdaa044}{10.1093/restud/rdaa044}.
\bibitem[Kondrakiewicz and Nawrocka(2025)]{kondrakiewicz2025} Kondrakiewicz, K., Nawrocka, J. (2025). \emph{Brains are expensive, but cognition is often cheap}. Neuroscience \& Biobehavioral Reviews, 179, 106450. DOI: \href{https://doi.org/10.1016/j.neubiorev.2025.106450}{10.1016/j.neubiorev.2025.106450}.
\bibitem[Kurzban(2010)]{kurzban2010} Kurzban, R. (2010). \emph{Does the brain consume additional glucose during self-control tasks?}. Evolutionary Psychology, 8(2), 244--259. DOI: \href{https://doi.org/10.1177/147470491000800208}{10.1177/147470491000800208}.
\bibitem[Kurzban et al.(2013)]{kurzban2013} Kurzban, R., Duckworth, A., Kable, J. W., Myers, J. (2013). \emph{An opportunity cost model of subjective effort and task performance}. Behavioral and Brain Sciences, 36(6), 661--679. DOI: \href{https://doi.org/10.1017/s0140525x12003196}{10.1017/s0140525x12003196}.
\bibitem[Laughlin et al.(1998)]{laughlin1998} Laughlin, S. B., de Ruyter van Steveninck, R. R., Anderson, J. C. (1998). \emph{The metabolic cost of neural information}. Nature Neuroscience, 1(1), 36--41. DOI: \href{https://doi.org/10.1038/236}{10.1038/236}.
\bibitem[Lennie(2003)]{lennie2003} Lennie, P. (2003). \emph{The cost of cortical computation}. Current Biology, 13(6), 493--497. DOI: \href{https://doi.org/10.1016/s0960-9822(03)00135-0}{10.1016/s0960-9822(03)00135-0}.
\bibitem[Ma and Woodford(2020)]{ma2020} Ma, W. J., Woodford, M. (2020). \emph{Multiple conceptions of resource rationality}. Behavioral and Brain Sciences, 43, e15. DOI: \href{https://doi.org/10.1017/s0140525x19001754}{10.1017/s0140525x19001754}.
\bibitem[Ma\'{c}kowiak et al.(2023)]{mackowiak2023} Ma\'{c}kowiak, B., Mat\v{e}jka, F., Wiederholt, M. (2023). \emph{Rational inattention: A review}. Journal of Economic Literature, 61(1), 226--273. DOI: \href{https://doi.org/10.1257/jel.20211524}{10.1257/jel.20211524}.
\bibitem[Markman and Otto(2011)]{markman2011} Markman, A. B., Otto, A. R. (2011). \emph{Cognitive systems optimize energy rather than information}. Behavioral and Brain Sciences, 34(4), 207--207. DOI: \href{https://doi.org/10.1017/s0140525x11000355}{10.1017/s0140525x11000355}.
\bibitem[Musslick and Cohen(2021)]{musslick2021} Musslick, S., Cohen, J. D. (2021). \emph{Rationalizing constraints on the capacity for cognitive control}. Trends in Cognitive Sciences, 25(9), 757--775. DOI: \href{https://doi.org/10.1016/j.tics.2021.06.001}{10.1016/j.tics.2021.06.001}.
\bibitem[Otto et al.(2026)]{otto2026} Otto, A. R., Westbrook, A., Daunizeau, J. (2026). \emph{Why is cognitive effort experienced as costly?}. Trends in Cognitive Sciences, 30(7), 632--645. DOI: \href{https://doi.org/10.1016/j.tics.2025.10.012}{10.1016/j.tics.2025.10.012}.
\bibitem[Padamsey et al.(2022)]{padamsey2022} Padamsey, Z., Katsanevaki, D., Dupuy, N., Rochefort, N. L. (2022). \emph{Neocortex saves energy by reducing coding precision during food scarcity}. Neuron, 110(2), 280--296.e10. DOI: \href{https://doi.org/10.1016/j.neuron.2021.10.024}{10.1016/j.neuron.2021.10.024}.
\bibitem[Prat-Carrabin et al.(2025)]{pratcarrabin2025} Prat-Carrabin, A., de Hollander, G., Bedi, S., Gershman, S. J., Ruff, C. C. (2025). \emph{Distributed range adaptation in human parietal encoding of numbers}. bioRxiv. DOI: \href{https://doi.org/10.1101/2025.09.25.675916}{10.1101/2025.09.25.675916}.
\bibitem[Prat-Carrabin and Woodford(2026)]{pratcarrabin2026} Prat-Carrabin, A., Woodford, M. (2026). \emph{Endogenous precision of the number sense}. eLife, 13, RP101277. DOI: \href{https://doi.org/10.7554/eLife.101277.4}{10.7554/eLife.101277.4}.
\bibitem[Prat-Carrabin and Woodford(n.d.)]{pratcarrabin_data} Prat-Carrabin, A., Woodford, M. (n.d.). \emph{Endogenous precision of the number sense: Data \& code}. Open Science Framework [Data set]. \url{https://osf.io/d6k3m/}.
\bibitem[Raichle et al.(2001)]{raichle2001} Raichle, M. E., MacLeod, A. M., Snyder, A. Z., Powers, W. J., Gusnard, D. A., Shulman, G. L. (2001). \emph{A default mode of brain function}. Proceedings of the National Academy of Sciences, 98(2), 676--682. DOI: \href{https://doi.org/10.1073/pnas.98.2.676}{10.1073/pnas.98.2.676}.
\bibitem[Raichle and Mintun(2006)]{raichle_mintun2006} Raichle, M. E., Mintun, M. A. (2006). \emph{Brain work and brain imaging}. Annual Review of Neuroscience, 29, 449--476. DOI: \href{https://doi.org/10.1146/annurev.neuro.29.051605.112819}{10.1146/annurev.neuro.29.051605.112819}.
\bibitem[Shaulson et al.(2024)]{shaulson2024} Shaulson, E. D., Cohen, A. A., Picard, M. (2024). \emph{The brain--body energy conservation model of aging}. Nature Aging, 4(10), 1354--1371. DOI: \href{https://doi.org/10.1038/s43587-024-00716-x}{10.1038/s43587-024-00716-x}.
\bibitem[Tesmer et al.(2025)]{tesmer2025} Tesmer, A. L., Pola, C. D., Gilli, D., Grujic, N., Bracey, E. F., Patriarchi, T., Peleg-Raibstein, D., Polania, R., Burdakov, D. (2025). \emph{Neurometabolic signaling and control of policy complexity}. bioRxiv. DOI: \href{https://doi.org/10.1101/2025.02.24.639890}{10.1101/2025.02.24.639890}.
\bibitem[Vadillo et al.(2016)]{vadillo2016} Vadillo, M. A., Gold, N., Osman, M. (2016). \emph{The bitter truth about sugar and willpower: The limited evidential value of the glucose model of ego depletion}. Psychological Science, 27(9), 1207--1214. DOI: \href{https://doi.org/10.1177/0956797616654911}{10.1177/0956797616654911}.
\bibitem[Woodford(2020)]{woodford2020} Woodford, M. (2020). \emph{Modeling imprecision in perception, valuation, and choice}. Annual Review of Economics, 12(1), 579--601. DOI: \href{https://doi.org/10.1146/annurev-economics-102819-040518}{10.1146/annurev-economics-102819-040518}.
\end{thebibliography}
\endgroup
\end{document}
