## Supplementary material for "The marginal brain": Replication package: supplement.md

---
title: Supplementary derivations and simulation design
authors: Fabio Lokwani-Di Matteo^{a} and Pier Luigi Sacco^{a}
affiliations: ^{a} Department of Neurosciences, Imaging and Clinical Sciences, University of Chieti-Pescara, Italy
style: author-date
page: letter
spacing: 1.2
justify: true
---

## S1. Model primitives and equilibrium

For a task with prior density π(x), the observer chooses the precision allocation I₁(x) of one signal and continuous effective signal mass n. Fisher information adds, so total precision is I(x) = nI₁(x). The capacity constraint is ∫I₁(x)^(1/2)dx ≤ K. Estimation and discrimination are represented by losses Lₐ[I] = ∫π(x)^a/I(x)dx, with a = 1 and a = 2, respectively. In the budget model, the task loss is scaled by β and each unit of effective signal mass n incurs marginal budget cost eμ. Dividing this objective by β yields the per-signal price λ = eμ/β = ep in the normalized loss problem, where p = μ/β is the budget price per unit loss.

Total energy use is B = C + E + Eₒ, where C is committed use, E = en is gross task energy, and Eₒ is energy allocated to competing uses. In normalized energy units, V(Eₒ) = v(Eₒ^(1−ρ) − 1)/(1−ρ) for ρ ≠ 1 and V(Eₒ) = v log Eₒ for ρ = 1; both imply V′(Eₒ) = vEₒ^(−ρ). We assume v > 0 and ρ > 0, so competing-use value is increasing and concave. Above a set point B̄, regulation incurs cost R(d), with d = B − B̄ and R′(d) = cd^k for c,k > 0. At an interior optimum, μ = R′(d) = V′(Eₒ) is the shared shadow price. For general task stakes β, p = μ/β is the budget price per unit loss and λ = ep = eμ/β is the effective price per signal; with β = 1, p = μ and λ = eμ. Defining M = B̄ − C gives the equilibrium condition d(μ) + M − E(μ) − Eₒ(μ) = 0. This common-budget structure formalizes the model's physiology-motivated account of resource allocation.

Because E(μ) depends on task scale, a finite-demand task can shift its own equilibrium price. The exact fourth-root relation is conditional on the effective per-signal price ratio faced by the task; a common cross-task factor is the price-taking benchmark when task demand is small relative to competing uses, or can be assessed by solving the task-specific equilibrium equations jointly. The main-text baseline reports the local sensitivity of μ to task scale.

## S2. Optimal encoding allocation

Set s(x) = I₁(x)^(1/2). At fixed n, the loss is n^(−1)∫π(x)^a/s(x)^2dx subject to ∫s(x)dx ≤ K. Pointwise minimization of π(x)^a/s(x)^2 + νs(x), where ν is the multiplier on the capacity constraint, gives s(x) ∝ π(x)^(a/3); the second derivative is positive. Normalizing to exhaust K yields s(x) = Kπ(x)^(a/3)/Zₐ, with Zₐ = ∫π(x)^(a/3)dx. The minimized loss conditional on n is Zₐ³/(nK²).

Minimizing Zₐ³/(nK²) + λn over n > 0 gives n* = Zₐ^(3/2)/(Kλ^(1/2)); at this optimum the allocation loss and signal cost are equal. Substitution gives I(x) = KZₐ^(−1/2)λ^(−1/2)π(x)^(2a/3) and σ(x) = Zₐ^(1/4)λ^(1/4)K^(−1/2)π(x)^(−a/3). Hence the price elasticity of imprecision is 1/4 throughout the prior's support. Under a uniform prior of width w, holding capacity K and effective signal price λ fixed, Zₐ = w^(1−a/3), giving σ ∝ w^((1+a)/4): the exponents are 1/2 for estimation and 3/4 for discrimination.

Under a generalized bound ∫I₁(x)^(γ/2)dx ≤ K, the optimal allocation still separates from n and the minimized loss remains proportional to 1/n. Therefore the price elasticity remains 1/4, although the exponents in w depend on γ. If signal cost is λn^κ, the optimum instead gives n* ∝ λ^(−1/(κ+1)) and σ ∝ λ^(1/(2(κ+1))). The discrete-count calculation uses the baseline specification γ = κ = 1 and is presented as a separate count-scale sensitivity illustration. When n is restricted to positive integers, it minimizes G/n + λn over n ∈ {1,2,…}, where G = Zₐ³/K². Comparing adjacent integer costs yields intervals in which the optimal count does not change; at larger counts the discrete response approaches the continuous fourth-root result. The normalized budget baseline in Methods implies an effective signal mass of {{baseline_effective_signal_mass}} normalized units, which anchors a separate count-scale sensitivity illustration. This illustration complements the human-data calibration and is presented as an effective activity measure rather than an anatomical spike count.

## S3. Budget equilibrium and net energy increment

From the encoding solution, unit-scale task energy satisfies Eᵘ(μ) ∝ μ^(−1/2) at fixed e and K. The value-maximizing allocation to competing uses is Eₒ(μ) = (v/μ)^(1/ρ), and regulation gives d(μ) = (μ/c)^(1/k). For task scale ε ≥ 0, write task energy as εEᵘ(μ). The equilibrium residual is

\[
F(\mu;\varepsilon)=\left(\frac{\mu}{c}\right)^{1/k}+M-\varepsilon E^{u}(\mu)-\left(\frac{v}{\mu}\right)^{1/\rho}.\tag{S1}
\]

Equation (S1) summarizes the budget balance as a single strictly increasing function of the shadow price: F_μ(μ;ε) = d/(kμ) + εEᵘ/(2μ) + Eₒ/(ρμ) > 0 for positive primitives and μ > 0. For finite c, its opposite boundary signs establish a unique equilibrium. Under a hard cap d = 0, a finite equilibrium requires M > 0. Implicit differentiation gives ∂μ/∂M = −1/F_μ < 0; increasing v or c raises μ.

For the no-task equilibrium, write \(\mu_0\) and evaluate \(d_0=d(\mu_0)\), \(E_{o,0}=E_o(\mu_0)\), and \(E^u_0=E^u(\mu_0)\) at that baseline. At \(\varepsilon=0\), \(F_\mu(\mu_0;0)=d_0/(k\mu_0)+E_{o,0}/(\rho\mu_0)\), so \(d\mu/d\varepsilon=E^u_0/F_\mu(\mu_0;0)\). Because total-use change equals the change in regulatory excess, \(d\Delta B/d\varepsilon=d'(\mu_0)d\mu/d\varepsilon\). Using \(d'(\mu_0)=d_0/(k\mu_0)\) gives \(\Delta B=\phi_0\varepsilon E^u_0+o(\varepsilon)\), with

\[
\phi_0=\frac{d(\mu_0)/k}{d(\mu_0)/k+E_o(\mu_0)/\rho}.\tag{S2}
\]

Equation (S2) makes the local allocation explicit: \(\phi_0\) is the fraction of an infinitesimal task-energy increment that appears as a net increase in total use, while \(1-\phi_0\) is initially reallocated from competing uses. For a local comparison of no-task states differing through \(M\) alone, holding \(c,v,k,\rho,e,K\), task stakes and task scale fixed, \(\phi_0\) has elasticity \((1-\phi_0)(1/k+1/\rho)\) with respect to \(\mu_0\). Because \(E^u(\mu_0)\propto\mu_0^{-1/2}\), the first-order net increment has elasticity \((1-\phi_0)(1/k+1/\rho)-1/2\) and rises with \(\mu_0\) under this comparison when \((1-\phi_0)(1/k+1/\rho)>1/2\). A transparent sufficient region is \(\phi_0<1/2\) and \(1/k+1/\rho\geq1\); the computational analysis also compares the no-task first-order approximation with the exact finite-task equilibrium change.

## S4. Joint identification of price and technology

Let p = μ/β denote the budget price per unit loss, so the effective price per signal is λ = ep; write x = Δlog p, y = Δlog e and z = Δlog K. The encoding solution implies Δlog σ = (x+y)/4−z/2 and Δlog E = (y−x)/2−z. Define the combined technology change T = y−2z. Eliminating y and z yields

\[
\begin{aligned}
\frac{\Delta\log\sigma}{\Delta\log E}&=\frac{x+T}{2(T-x)}, &
\Delta\log(\sigma^2 E)&=T,\\
x&=2\Delta\log\sigma-\Delta\log E, &
T&=2\Delta\log\sigma+\Delta\log E.
\end{aligned}\tag{S3}
\]

A pure change in p at fixed e and K has T = 0 and slope −1/2; a pure technology change has x = 0 and slope +1/2. The finite slope is defined only when ΔlogE ≠ 0 (T ≠ x). If T = x ≠ 0, task energy does not change while imprecision does; if x = T = 0, neither observable changes and the slope is undefined. The last two identities in (S3) give the direct inverse map from the observed log changes to the budget-price change x and combined technology index T, including when the slope is undefined. Measuring e and K separately decomposes T, while measuring task stakes β maps p to μ. A pure change in p at fixed e and K also satisfies (σ₁/σ₀)^4 = (E₀/E₁)^2. These restrictions require stable coding technology outside the modeled price channel; claims about μ also require stable or measured stakes.

If measured regional energy is E−D, where D is displaced activity, define δ = dD/dE. Then \(d\log(E-D)=(1-\delta)dE/(E-D)\), so the measured slope is the theoretical slope multiplied by (E−D)/(E(1−δ)). Proportional displacement preserves pure-change slopes, and measuring displacement places regional energy observations on the model's gross-energy scale across states. The numerical scenarios use a baseline displacement D₀ = {{headline.displacement_baseline_D:.2f}} and keep both D and measured net energy within their feasible range.

## S5. Behavioral calibration and simulation design

The behavioral component implements the numerosity estimation and discrimination tasks summarized by Prat-Carrabin and Woodford (2026; the full bibliographic entry appears in the main manuscript references). Estimation draws integer stimuli from the task's prior range, decodes noisy representations with the posterior mean under a uniform prior, adds truncated motor noise and rounds responses. Discrimination compares noisy representations of two averages, each formed from five sampled integers; lapses choose either option at random. Calibration uses published summary statistics, with a trial-level reconstruction of the estimation anchor reported in S6.

The simulations vary two within-subject states and assume participant heterogeneity in log representation noise, lapse rate, state effect and, for estimation, motor noise. Discrimination uses {{config.power.trials_per_state}} trials per state, prior range {{config.power.lo}}–{{config.power.hi}}, sample sizes {{config.power.N_grid}} and one-sided t and Wilcoxon tests on fitted log f. Estimation uses {{config.estimation_recovery.trials_per_state}} trials per state, prior range {{config.estimation_recovery.lo}}–{{config.estimation_recovery.hi}}, {{config.estimation_recovery.N}} participants and one-sided signed-rank and t tests. Each condition uses {{config.power.reps}} discrimination and {{config.estimation_recovery.reps}} estimation replications. The N = 20 and N = 40 discrimination estimates are computed from nested subsets of each N = 80 simulated cohort, so power estimates across sample sizes are paired rather than independent. A lapse sensitivity analysis multiplies second-state lapse odds by {{config.lapse_robustness.lapse_odds_mult}} under price ratios 1 and 2, fitting shared or state-specific lapse rates, with {{config.lapse_robustness.reps}} replications per condition.

Subject fits use bounded L-BFGS-B optimization as specified in the main Methods. Convergence and boundary contact are recorded for each fit. Rejection rates are accompanied by Wilson intervals, including estimates at 0 or 1. Estimation runs in independently seeded chunks; cache fingerprints include the active source, configuration, calibration and numerical environment. Complete condition-level results are stored in `model/results.json`, which also supplies manuscript values and figures.

## S6. Trial-level reconstruction and matched-numerosity analysis

We reanalyzed the public estimation-trial file from the 36-person sample reported by Prat-Carrabin and Woodford. For each prior width and presented numerosity, the pooled anchor analysis subtracts the condition-level mean response. At the common numerosity 60, the reconstructed residual standard deviations can be compared directly with the published rounded targets and the number of available trials (Table S1), reproducing the calibration anchor from the released records.

| Prior width | Trials at numerosity 60 | Reconstructed residual SD | Published SD |
|:--:|--:|--:|--:|
| 20 | {{headline.empirical_n_width_20}} | {{headline.empirical_sd_width_20:.2f}} | 4.2 |
| 40 | {{headline.empirical_n_width_40}} | {{headline.empirical_sd_width_40:.2f}} | 6.8 |
| 60 | {{headline.empirical_n_width_60}} | {{headline.empirical_sd_width_60:.2f}} | 8.4 |

**Table S1.** Reconstructed pooled residual variability at numerosity 60 by prior width. The estimates closely reproduce the published summaries; the largest absolute difference is {{headline.empirical_max_abs_sd_difference:.2f}} response units.

The close reconstruction shows that the released trial file recovers the source study's three calibration anchors at the common numerosity before the matched-level comparisons are formed. It also gives the reader the trial counts underlying each anchor, making the empirical starting point transparent and independently checkable.

For the matched-numerosity comparisons, we use the 21 numerosities shared by all three prior ranges, 50–70 inclusive. Let \(n_{iwx}\) be the number of responses for participant i, width w and numerosity x, and let \(r_{iwxt}\) be response t. In cells with at least two trials, define \(SSE_{iwx}=\sum_t(r_{iwxt}-\bar r_{iwx})^2\) and \(df_{iwx}=n_{iwx}-1\); cells with fewer than two trials contribute zero SSE and zero residual degrees of freedom. For each fixed x and w, the primary pooled residual-variance estimator is

\[
\widehat V(x,w)=\frac{\sum_i SSE_{iwx}}{\sum_i df_{iwx}}.\tag{S4}
\]

For each x, the variance ratio compares these pooled estimates. In each bootstrap replicate, we resample participants within the recorded Order groups—20 from Widening and 16 from Narrowing—and carry each selected participant's cells across widths, preserving both the paired design and observed sequence composition. The two-sided test uses the centered bootstrap distribution of the log variance ratio. Let $\hat\theta=\log\{\widehat V(x,w_2)/\widehat V(x,w_1)\}$ denote the observed log variance ratio and let $\hat\theta_b^*$ be its estimate in bootstrap replicate $b$, for $b=1,\ldots,B$. The Monte Carlo tail probability is $\hat p=\{1+\sum_{b=1}^{B}\mathbf{1}(|\hat\theta_b^*-\hat\theta|\geq|\hat\theta|)\}/(B+1)$, which applies the plus-one correction to the centered, two-sided bootstrap test. P-values are Bonferroni-adjusted over 21 numerosity levels × 3 width pairs = 63 contrasts. For each contrast, write $q=\alpha/63$ and $k_{\max}=\lceil q(B+1)-1\rceil-1$, where B is the number of bootstrap draws. Let c be the $(B-k_{\max})$-th ordered value of $|\hat\theta_b^*-\hat\theta|$; then $[\hat\theta-c,\hat\theta+c]$ is the interval on the log-ratio scale, transformed back to a variance ratio. The nominal per-contrast confidence level is {{headline.empirical_familywise_interval_level_percent:.4f}}%; the Bonferroni adjustment targets a nominal 5% familywise level. These nominal summaries are reported as outputs of the designated procedure; their inferential scope is described once under Limitations and scope conditions in the main text. The interval is decision-dual to the reported nominal Bonferroni-adjusted test, including the finite-replication rule. Every contrast uses {{config.empirical_reanalysis.bootstrap_reps}} draws. The results file also reports Wilson 95% Monte Carlo intervals for the raw bootstrap tail fractions, including when the simulated tail count is zero; these intervals quantify simulation error in tail estimation.

The fixed-numerosity contrasts are assembled in Table S2, so the complete set of ratios and nominal multiplicity-adjusted outputs can be inspected directly. The support column separates participants contributing residual degrees of freedom at each width from those with paired support for each width contrast; each contrast cell places the variance ratio before its adjusted p-value.

| x; n by width; paired n | 40 / 20 ratio (nominal adj. p) | 60 / 20 ratio (nominal adj. p) | 60 / 40 ratio (nominal adj. p) |
|:--|:--|:--|:--|
{{empirical_per_number_rows}}

**Table S2.** Pooled residual-variance ratios, nominal Bonferroni-adjusted bootstrap p-values and participant-support counts for each of the 21 shared numerosities. In the support column, *x* is numerosity; the first support triplet gives the number of participants with residual degrees of freedom at widths 20, 40 and 60, in that order; the second gives the number with residual degrees of freedom at both widths for contrasts 20/40, 20/60 and 40/60, in that order. Each contrast entry reports the pooled variance ratio followed by its nominal adjusted centered-bootstrap p-value in parentheses.

Across the matched stimulus range, point estimates are above one at every numerosity for the 20-to-40 and 20-to-60 contrasts. Table S2 reports the complete ratio profile and outputs from the designated interval procedure, while the 40-to-60 panel presents the distinct pattern for the wider-width comparison.

The pooled estimator weights participants by their residual degrees of freedom. To assess sensitivity to that weighting, we pool each participant's within-cell residual sums of squares and degrees of freedom over that participant's available cells in the common 50–70 numerosity range, compute a participant-level log variance ratio, and average those log ratios with equal participant weights. This gives each participant the same weight while retaining the observed cell and residual-degree-of-freedom contributions within that participant, with numerosity-level support determined by the released observations. The bootstrap resamples paired participant-level ratios within Widening and Narrowing sequences, retaining the observed group sizes; intervals and p-values are nominal Bonferroni-adjusted outputs across the three width pairs, using nominal per-contrast tests. This distinct equal-participant-weighted geometric-mean estimand adds a participant-balanced view alongside the numerosity-specific primary analysis. The three estimates and their nominal familywise intervals are presented together in Table S3, making the alternative participant-weighted summary directly comparable across width pairs.

| Prior-width contrast | Paired participants | Equal-weight geometric mean ratio | Nominal Bonferroni-adjusted 98.33% interval | Nominal adjusted p |
|:--|--:|--:|--:|--:|
{{empirical_equal_weight_rows}}

**Table S3.** Equal-participant-weighted residual-variance ratios over the common numerosity range, with nominal Bonferroni-adjusted bootstrap intervals and p-values based on nominal per-contrast tests.

Under this participant-level estimand, the 40/20 geometric-mean variance ratio is {{headline.empirical_equal_weight_ratio_20_40:.2f}} (nominal 98.33% per-contrast interval {{headline.empirical_equal_weight_ratio_20_40_lo:.2f}}–{{headline.empirical_equal_weight_ratio_20_40_hi:.2f}}; nominal Bonferroni-adjusted p = {{headline.empirical_equal_weight_ratio_20_40_p:.3g}}). The 60/20 estimate is {{headline.empirical_equal_weight_ratio_20_60:.2f}} ({{headline.empirical_equal_weight_ratio_20_60_lo:.2f}}–{{headline.empirical_equal_weight_ratio_20_60_hi:.2f}}; nominal adjusted p = {{headline.empirical_equal_weight_ratio_20_60_p:.3g}}), and the 60/40 estimate is {{headline.empirical_equal_weight_ratio_40_60:.2f}} ({{headline.empirical_equal_weight_ratio_40_60_lo:.2f}}–{{headline.empirical_equal_weight_ratio_40_60_hi:.2f}}; nominal adjusted p = {{headline.empirical_equal_weight_ratio_40_60_p:.3g}}). These participant-equal estimates are a distinct summary of the available-cell data; the fixed-numerosity results in Table S2 remain the primary analysis.

The sequence-specific check assesses whether the participant-equal variance pattern is similar in the two observed block orders. For each participant, residual sums of squares and degrees of freedom are pooled over the shared 50–70 numerosity support, a within-participant variance ratio is calculated, and log ratios are averaged within sequence. Participants are resampled within sequence, retaining the observed group sizes. Nominal Bonferroni adjustments span the six sequence-by-contrast estimates and, separately, the three between-sequence differences. The resulting summaries show how the observed variance pattern is expressed across the two recorded block orders.

Sequence-specific ratios and their between-sequence differences appear together in Table S4. This paired presentation makes the common direction of the main contrasts easy to compare while keeping sequence heterogeneity and its uncertainty visible.

| Prior-width contrast | Widening sequence, ratio (nominal adjusted interval) | Narrowing sequence, ratio (nominal adjusted interval) | Difference in log ratios (nominal adjusted interval) |
|:--|--:|--:|--:|
{{empirical_order_rows}}

**Table S4.** Sequence-stratified participant-equal variance ratios and nominal Bonferroni-adjusted bootstrap intervals over the common numerosity range.

In both recorded sequences, the point estimates for 40/20 and 60/20 exceed one, consistent with larger observed within-cell response dispersion under the wider prior conditions. The 40/60 estimates provide the more modest contrast and are shown separately rather than being used to imply a uniform monotonic pattern. The sequence comparison is informative about descriptive robustness, while its intervals and the blocked design preclude a claim that the width association is free of every order-related influence.

The within-block position check splits each participant × width × numerosity cell at trial 60. Within each half, cells with at least two observations are centered on their half-specific mean; their residual sums of squares and degrees of freedom are pooled across participants and the 21 common numerosities. A paired participant bootstrap resamples within the recorded Widening and Narrowing groups. Each contrast uses {{config.empirical_reanalysis.position_sensitivity_bootstrap_reps}} draws, and nominal Bonferroni adjustment spans three width pairs × two block halves. The paired half-block estimates describe how the variance pattern appears across trial position.

Table S5 assembles the six within-block estimates and interval and p-value outputs from the designated bootstrap procedure. A ratio above one indicates greater residual response variance in the wider-width condition for that block half.

| Block half | Prior-width contrast | Variance ratio (nominal adjusted interval) | Nominal adjusted p (raw-tail MC 95% interval) | Participants with residual df, narrower / wider |
|:--|--:|--:|--:|--:|
{{empirical_position_rows}}

**Table S5.** Within-block half-specific variance ratios, nominal Bonferroni-adjusted bootstrap intervals and p-values, and effective participant counts.

The table makes the scale and uncertainty of the half-specific comparisons visible. It shows the 40/20 and 60/20 contrasts in both halves and places the 40/60 estimates alongside them, giving a within-block view of how the full-sample pattern is expressed across trial position. These estimates provide a complementary sensitivity view of the released trials.

Condition-level results for the specified discrimination and estimation tests are brought together in Table S6, with lapse specifications, Wilson intervals and replication counts shown alongside each rejection rate. The display lets readers inspect the full design-specific operating characteristics alongside the comparisons highlighted in the main Results.

| Simulation condition | Price ratio | N | Test / lapse model | Rejection rate | Wilson 95% interval | Replications |
|:--|--:|--:|:--|--:|:--|--:|
{{simulation_summary_rows}}

**Table S6.** Simulated rejection rates with Wilson intervals, lapse specifications and replication counts.

Across these simulations, larger price ratios and samples raise detection, while lapse specification affects both null rejection and sensitivity to a price change. The nested samples make comparisons across N paired within each simulated cohort; Wilson intervals convey Monte Carlo precision. Fit diagnostics are summarized separately: {{lapse_fit_diagnostics_supplement}}

To assess sensitivity to cell availability, we retained participants with at least two responses in both conditions for each width-pair-by-numerosity and pooled their within-cell residual sums of squares and degrees of freedom. Matched sets vary by row. Table S7 summarizes the resulting pair-specific median ratios, ranges and participant support; the complete set of centered-bootstrap outputs remains in the accompanying machine-readable sensitivity file.

| Prior-width contrast | Matched participants per numerosity, median (range) | Median pooled variance ratio (range) | Numerosity levels with point estimate above 1 |
|:--|--:|--:|--:|
{{complete_case_sensitivity_rows}}

**Table S7.** Pair-specific complete-case variance ratios and matched-participant support. Each point estimate is based on participants with usable cells in both conditions at that numerosity; consequently, the participant set can differ across rows and contrasts. The final column counts numerosity-specific point estimates above one.

The two contrasts anchored at width 20 retain an above-one point estimate at every shared level, while the 60/40 profile remains above one at most levels. This matched-set view complements the fixed-support profile and shows the stability of the principal directional pattern across available-cell definitions.

## S7. Participant-level directional sensitivity

This participant-level sensitivity compares each participant's sample variance at the wider and narrower prior for each shared numerosity. A participant is eligible when both cells contain at least two trials. Positive and negative entries count participants whose observed wider-condition sample variance is larger or smaller; exact ties are omitted from the binomial denominator, while a one-zero pair remains directional. The two-sided exact sign test evaluates balance between positive and negative signs among eligible participants, with a nominal Bonferroni adjustment across the 63 numerosity-by-width contrasts in this sensitivity family. The full set of level-specific counts, ties and exact and adjusted p-values appears in Table S8.

| Numerosity | Width pair | Eligible N | Sign counts (+ / - / ties) | Exact p | Adjusted p |
|--:|:--:|--:|:--:|--:|--:|
{{participant_sign_sensitivity_rows}}

**Table S8.** Exact participant-level sign-balance results for observed sample-variance differences. Each row reports the wider/narrower width pair, eligible participant count, positive/negative/tied signs, exact p-value, and Bonferroni-adjusted p-value for the complete 63-test family.

The exact sign counts show directionally consistent 40/20 contrasts at selected numerosities, with the complete 60/20 and 60/40 results reported in Table S8. A separate participant-level aggregation pools available within-cell residual sums of squares and degrees of freedom over numerosity 50–70 before forming one ratio per participant. The resulting direction test is {{aggregate_participant_sign_summary}}; its exact p-values use Bonferroni adjustment across the three width contrasts. This aggregate directional summary complements the fixed-numerosity pooled estimates with one equally interpretable participant-level comparison.

## S8. Matched-numerosity reanalysis in full

This section reports the matched-numerosity reanalysis of the published estimation trials in full; Figure 6 of the main text displays the per-numerosity ratios together with the ratios that the calibrated model predicts.

The released estimation trials provide a detailed component-level check of how prior width relates to response dispersion. At numerosity 60, pooled residual standard deviations reconstructed from 12,960 trials in the published 36-person sample closely reproduce the published summaries (Table S1). The matched-numerosity reanalysis holds numerosity fixed at all 21 levels from 50 through 70 observed in every prior-width condition, comparing prior widths at matched stimulus values throughout the shared range.

For each numerosity x and prior width w, we subtract each participant's response mean within that participant × width × numerosity cell. We apply the pooled residual-variance estimator in Supplementary equation (S4), weighting each participant by the observed residual degrees of freedom. We compare all three width pairs separately at each x. The released file records Widening and Narrowing sequences; the first observed trial time in each width block confirms the sequences 20→40→60 (20 participants) and 60→40→20 (16 participants). Each paired participant bootstrap draw resamples separately within those two groups, maintaining their observed composition while carrying all of a selected participant's cells together across widths. The procedure applies a nominal Bonferroni adjustment over the complete family of 63 numerosity-by-width contrasts. The resampling unit is the participant, not the trial. A within-block split-half sensitivity divides each block into trials 1–60 and 61–120, re-centers within participant × width × numerosity × half, and applies a nominal Bonferroni adjustment across the six half-by-width contrasts; it asks whether the descriptive pattern appears throughout a block without treating the trials as independent participants.

Table S9 brings together pooled residual-variance ratios and interval outputs from the designated centered-bootstrap procedure. The point estimates show higher residual variance under wider priors in the 20-to-40 and 20-to-60 contrasts, with the 40-to-60 profile providing the complementary wider-width comparison. Reporting all three profiles makes the magnitude and shape of the association visible across the shared numerosity range.

**Table S9.** Matched-numerosity residual-variance contrasts in the released estimation sample. Each entry compares pooled within-participant residual variance at the wider and narrower prior; the range spans the 21 shared numerosities. “Nominal procedure-based interval exclusions” counts two-sided participant-cluster bootstrap intervals that exclude a ratio of one after the designated nominal Bonferroni adjustment over all 63 numerosity-by-width contrasts.

| Prior-width contrast | Median variance ratio (range across numerosity) | Nominal interval exclusions | Largest nominal adjusted p (raw-p MCSE) |
|:--|--:|--:|--:|
| 20 → 40 | {{headline.empirical_variance_ratio_20_40:.2f}} ({{headline.empirical_variance_ratio_20_40_min:.2f}}–{{headline.empirical_variance_ratio_20_40_max:.2f}}) | {{headline.empirical_variance_ratio_20_40_significant_numerosities}} / 21 | {{headline.empirical_variance_ratio_20_40_max_adjusted_pvalue:.3g}} ({{headline.empirical_variance_ratio_20_40_max_adjusted_pvalue_mcse:.1g}}) |
| 20 → 60 | {{headline.empirical_variance_ratio_20_60:.2f}} ({{headline.empirical_variance_ratio_20_60_min:.2f}}–{{headline.empirical_variance_ratio_20_60_max:.2f}}) | {{headline.empirical_variance_ratio_20_60_significant_numerosities}} / 21 | {{headline.empirical_variance_ratio_20_60_max_adjusted_pvalue:.3g}} ({{headline.empirical_variance_ratio_20_60_max_adjusted_pvalue_mcse:.1g}}) |
| 40 → 60 | {{headline.empirical_variance_ratio_40_60:.2f}} ({{headline.empirical_variance_ratio_40_60_min:.2f}}–{{headline.empirical_variance_ratio_40_60_max:.2f}}) | {{headline.empirical_variance_ratio_40_60_significant_numerosities}} / 21 | {{headline.empirical_variance_ratio_40_60_max_adjusted_pvalue:.3g}} ({{headline.empirical_variance_ratio_40_60_max_adjusted_pvalue_mcse:.1g}}) |

The pooled ratios show a substantial and consistent narrow-to-intermediate descriptive difference, while the 40-to-60 estimates map how that association varies across numerosity. The participant-level exact sign analysis supplies a complementary directional cross-check alongside the full pooled pattern. Paired participant support ranges from {{empirical_support_summary}}; Supplementary Table S2 reports the exact level-specific counts and bootstrap results.

A complementary participant-level sign-balance analysis asks whether observed sample variances tend to be larger in the wider condition, giving each eligible participant one directional contribution. Within its 63-test family, corrected directional results occur for {{participant_sign_sensitivity_summary}}. Because this summary weights participants equally rather than pooling residual degrees of freedom, it localizes directional evidence to selected numerosity levels and provides a distinct view of the primary pattern. All 63 tests, ties and eligible participant counts are listed in Supplementary Table S8.

The sequence-stratified, participant-equal sensitivity provides a complementary view of the two largest contrasts. For width 40 relative to width 20, the geometric mean variance ratios are {{headline.empirical_order_widening_ratio_20_40:.2f}} in the Widening sequence ({{headline.empirical_order_widening_ci_20_40}}) and {{headline.empirical_order_narrowing_ratio_20_40:.2f}} in the Narrowing sequence ({{headline.empirical_order_narrowing_ci_20_40}}). For width 60 relative to width 20, the corresponding estimates are {{headline.empirical_order_widening_ratio_20_60:.2f}} ({{headline.empirical_order_widening_ci_20_60}}) and {{headline.empirical_order_narrowing_ratio_20_60:.2f}} ({{headline.empirical_order_narrowing_ci_20_60}}). The smaller 60/40 contrast is also above one in both sequences: {{headline.empirical_order_widening_ratio_40_60:.2f}} ({{headline.empirical_order_widening_ci_40_60}}) for Widening and {{headline.empirical_order_narrowing_ratio_40_60:.2f}} ({{headline.empirical_order_narrowing_ci_40_60}}) for Narrowing, with nominal intervals adjusted across the six sequence-by-contrast estimates. The bootstrap keeps the participant count fixed within each sequence and shows whether the main pattern is visible in both recorded orders; the complete contrasts and between-sequence uncertainty appear in Supplementary Table S4. Similar estimates across sequences strengthen the descriptive robustness of the matched-level pattern.

Heterogeneity across stimulus magnitudes is visible in the 21 fixed-numerosity variance ratios in Figure 6. Supplementary Table S2 reports the corresponding point estimates and nominal adjusted p-values; Table S9 summarizes the number of nominal procedure-based intervals excluding one in each contrast.


Across the 21 shared numerosities, the point estimates exceed one for all 21 width-40-versus-20 contrasts and all 21 width-60-versus-20 contrasts, and for 19 width-60-versus-40 contrasts (Figure 6). The participant-equal aggregate estimates appear once in Table S3 and complement this level-specific profile.

The pair-specific complete-case sensitivity evaluates each contrast using participants with usable response cells in both conditions at that numerosity. {{complete_case_sensitivity_summary}} The contributing participants vary across levels, and Supplementary Table S7 reports the corresponding estimates and point-estimate direction, providing a direct view of cell-availability weighting.

The split-half sensitivity provides a second descriptive view of how the matched-width pattern behaves across trial position. For 40 relative to 20, the variance ratios are {{headline.empirical_position_ratio_first_half_20_40:.2f}} in trials 1–60 and {{headline.empirical_position_ratio_second_half_20_40:.2f}} in trials 61–120; for 60 relative to 20 they are {{headline.empirical_position_ratio_first_half_20_60:.2f}} and {{headline.empirical_position_ratio_second_half_20_60:.2f}}. The corresponding 60-to-40 ratios are {{headline.empirical_position_ratio_first_half_40_60:.2f}} and {{headline.empirical_position_ratio_second_half_40_60:.2f}}. The split-half point estimates retain the same broad direction as the full-block results, including for the 60-to-40 contrast. The table presents the procedure-based interval and p-value outputs for both halves, while the estimates quantify pooled half-block patterns alongside the full-block results.

The model derivation establishes a formal link between brain energetics and decision precision and identifies a concrete empirical design target. Jointly measuring precision and stimulus-linked task energy can test the model's price-versus-technology restriction, and adding physiological measures can test the shared-budget mechanism directly.

## S9. Numerical companion to the Results

This section reports the quantitative results that the main-text Results summarize, with the same headings; the figures are those of the main text.

### S9.1. Precision and its price

The first set of results concerns the encoding layer, with the per-signal price λ taken as given; the budget enters these results only through λ = e μ/β, and every result is derived and verified in Methods.

The optimal encoding allocation separates cleanly from the number of signals. Because n enters the loss only through the factor 1/n, the optimal allocation for a single signal is independent of n: precision is allocated in proportion to π(x)^(2a/3), so that I₁(x)^½ ∝ π(x)^(a/3). The minimized loss after this allocation is Z_a³/(nK²), where Z_a = ∫ π(x)^(a/3) dx summarizes the prior's shape. This separation makes the model's task-specific prior structure transparent while yielding explicit comparative statics for the common resource response.

With linear signal cost λn, the optimal signal mass is n* = Z_a^(3/2)/(K λ^½), and imprecision is σ(x) = Z_a^(¼) λ^(¼) K^(−½) π(x)^(−a/3). Under a uniform prior of width w, imprecision grows as w^½ in estimation and w^¾ in discrimination, reproducing the prior-width scalings reported by Prat-Carrabin and Woodford; optimal signal mass grows as w in estimation and w^½ in discrimination. The result preserves the established task-specific width patterns while identifying the shared resource price that shifts precision across states.

The central comparative static is an exact fourth-root response: the elasticity of imprecision to the effective per-signal price is ¼ at every magnitude, for every prior, task, and capacity. Since λ = eμ/β, when coding technology and stakes are unchanged, a common price ratio multiplies imprecision by its fourth root at each magnitude while preserving each task's prior-width scaling. Across estimation and discrimination, this conditional invariance appears as a common upward shift while the task-specific prior-width slopes remain intact (Figure 2). Doubling the effective price raises imprecision by {{headline.common_factor_price_x2:.3f}}; the same precise restriction across priors therefore creates a jointly testable behavioral signature. Finite task demand feeds back on μ, so an exact cross-task comparison requires the tasks to face the same effective-price ratio; when task demand is small, the price-taking approximation yields a direct benchmark. Lemma 2 quantifies this feedback: in the illustrative baseline, the task uses {{headline.baseline_task_share:.1%}} of energy above committed use and its local task-scale elasticity is {{headline.el_mu_taskscale:.3f}}. Joint estimation can therefore distinguish the common-factor benchmark from task-specific equilibrium prices. The fourth-root exponent follows from additive Fisher information and linear cost per signal. It persists under the generalized bound ∫ I₁(x)^(γ/2) dx ≤ K, although the prior-width exponents change (at γ = ½ they are {{robustness.gammas.gamma0.5.a1.el_sigma_w:.2f}} for estimation and {{robustness.gammas.gamma0.5.a2.el_sigma_w:.2f}} for discrimination). A convex signal cost λn^κ changes the price elasticity to 1/(2(κ + 1)) while preserving its positive direction. In a separate count-scale sensitivity analysis, exact integer optima evaluated over hypothetical continuous optima from 3 to 10 yield an imprecision factor between {{headline.integer_range_3_10_lo:.2f}} and {{headline.integer_range_3_10_hi:.2f}} after a price doubling; the discrete staircase approaches the continuous prediction at larger counts. This illustration is distinct from the normalized budget baseline, which implies an effective signal mass of {{baseline_effective_signal_mass}} normalized units and does not represent a literal neural count or a calibration target for the human-data analysis.

The same optimum links precision to expenditure. Holding energy per signal, capacity, task scale and stakes fixed, task energy is E* = e n* ∝ μ^(−½): a higher shadow price reduces the encoding population's expenditure, while task loss valued at its stakes equals that expenditure valued at the margin, βL* = μE*. Under β = 1, this becomes L* = μE*. This relationship shows why a modest energetic draw can coexist with a meaningful decision cost; expenditure alone does not measure the behavioral importance of a computation. The aggregate increment depends in turn on how the surrounding budget reallocates energy, which is the next result.

### S9.2. The budget and the price

The budget equilibrium translates the available reallocation margin into a unique shadow price. The condition F(μ) = d(μ) + M − E(μ) − Eₒ(μ) = 0, where d(μ) = (μ/c)^(1/k) is use above the set point, has a unique solution for every convexity k > 0 because F increases strictly in μ; under a hard cap, existence requires a positive margin. The comparative statics are intuitive and testable: μ falls as M widens and rises with the value of competing uses or regulatory stiffness. A tighter margin thus raises the price faced by funded computations and, through the fourth-root restriction, raises imprecision. In the illustrative baseline (Methods), the price elasticity with respect to M is {{headline.el_mu_M_baseline:.2f}} and the corresponding imprecision elasticity is {{headline.el_sigma_M_baseline:.2f}}. Tightening M by 30% multiplies the price by {{headline.tighten30_mu_ratio:.2f}}, imprecision by {{headline.tighten30_sigma_ratio:.3f}}, and task energy by {{headline.tighten30_energy_ratio:.3f}} (Figure 3A). Under a hard cap, the price elasticity approaches the negative curvature of competing uses ({{headline.el_mu_M_hard_rho1:.2f}} at curvature 1 and {{headline.el_mu_M_hard_rho2:.2f}} at curvature 2); a graded precision response therefore arises when competing uses display diminishing returns.

The equilibrium also clarifies when a task can be treated as taking the price as given. The elasticity of μ with respect to task-demand scale is (E/μ)/F′(μ), and it vanishes as the task's share of energy above committed use approaches zero. The illustrative baseline uses {{headline.baseline_task_share:.1%}} of that energy and has a local elasticity of {{headline.el_mu_taskscale:.3f}}. Thus, the price is determined by the wider budget and transmitted to the task, even when the task itself has a small energy share.

The distinction between gross task energy and net total-use change follows from equilibrium reallocation. Switching on a task changes the price from μ₀ to μ and total use by ΔB = d(μ) − d(μ₀). If demand is scaled by ε and Eᵘ is gross energy at unit scale, the local expansion is ΔB = φ₀εEᵘ(μ₀) + o(ε), with φ₀ = [d(μ₀)/k]/[d(μ₀)/k + Eₒ(μ₀)/ρ], evaluated at the no-task equilibrium. The coefficient is the share of an infinitesimal task-energy increment that initially appears in total use; the balance is reallocated from competing uses. Under quadratic regulation (k = 1), φ₀ = 1/[1 + cEₒ(μ₀)/(ρμ₀)], and it approaches zero under a hard cap. For a local comparison of no-task states differing only in margin M, with c, v, k, ρ, e, K, stakes and task scale held fixed, φ₀ has elasticity (1 − φ₀)(1/k + 1/ρ) with respect to μ₀. Since Eᵘ ∝ μ₀^(−½), the first-order net increment rises with μ₀ when (1 − φ₀)(1/k + 1/ρ) > ½; a sufficient condition is φ₀ < ½ and 1/k + 1/ρ ≥ 1. This local condition complements the finite-task equilibrium in Figure 3B, where the net increment is {{headline.net_over_gross_baseline:.2f}} of gross task energy ({{headline.reallocated_share_baseline:.0%}} reallocated), and the no-task approximation differs from the finite-task result by {{headline.lemma3_approx_rel_err:.1%}}. The result makes explicit how a small aggregate increment can coexist with substantial gross task demand. The local comparative static is conditional: sufficiently convex regulation combined with highly curved competing uses can make the net increment decline with price.

Neither the task's share of use above committed expenditure nor its net increment identifies μ on its own. For any chosen pair consisting of a share s ∈ (0, 1) and a positive μ, budget primitives can reproduce that pair; at a fixed price, regulatory stiffness also changes the fraction of gross task energy recorded in total use. In the illustrative baseline that fraction falls from {{headline.same_price_net_over_gross_c1:.2f}} at c = 1 to {{headline.same_price_net_over_gross_c1e5:.1e}} at c = 10⁵ and reaches zero under a hard cap. Measuring task-evoked changes alongside displaced use and regulatory primitives therefore provides the information needed to interpret a net increment as a budget response. The joint non-identification of share and net increment follows constructively in the main-text proof: with $k=\rho=1$, fixed $E(\mu)=A\mu^{-1/2}$, share $1/2$, and common net increment $\Delta$, any two distinct prices satisfying $E(\mu)>2\Delta$ admit positive budget primitives that reproduce both quantities.

### S9.3. Telling a price from a technology

Behavior alone does not separate a change in the budget price per unit loss from a change in coding technology, since both rescale imprecision: lower capacity K raises imprecision with elasticity −½ in K, while a higher p = μ/β raises it with elasticity ¼ when e and K are fixed. Joint measurement of task energy and imprecision distinguishes these channels.

The joint sign restriction distinguishes a price change from a technology change. Let p = μ/β be the budget price per unit loss, λ = ep the effective price per signal, and define x = Δlog p, y = Δlog e and z = Δlog K. The encoding solution gives Δlog σ = (x + y)/4 − z/2 and Δlog E = (y − x)/2 − z. A pure price change at fixed e and K moves imprecision and energy in opposite directions, with slope Δlog σ/Δlog E = −½; a pure technology change moves them together with slope +½ (Figure 4). More generally, the slope is (x + T)/(2(T − x)), where T = y − 2z is the combined technology index. It is negative when the price change dominates (|T| < |x|) and positive when technology dominates. Equivalently, Δlog(σ²E) = T: under a pure price change at fixed e and K, σ²E is invariant. These two measurements identify changes in p and the combined index T; measuring task stakes maps p to μ, while an additional channel can distinguish e from K. Holding reward value comparable across states makes this restriction empirically operational. The mixed-change illustration doubles p, lowers the scenario input e by 29%, and lowers the scenario input K by 20%, yielding a model slope of {{headline.padamsey_like_slope:.2f}}. The 29% scenario is numerically inspired by the reported reduction in ATP expenditure associated with excitatory postsynaptic currents in food-restricted mouse visual cortex [[padamsey2022]], but the cited study does not estimate the model's energy-per-signal parameter e; the 20% capacity change is likewise a scenario input rather than a measured change in K.

Jointly measuring stimulus-linked energy in the encoding population provides the empirical counterpart; for numerosity, parietal cortex is a candidate because number encoding adapts to the prior range [[pratcarrabin2025]]. A regional task-evoked glucose-uptake change measures E − D, where D is activity displaced in the same region. If δ = dD/dE, the observed slope is the theoretical slope multiplied by (E − D)/(E(1 − δ)). Proportional displacement preserves the pure-price and pure-technology slopes of −½ and +½; when displacement changes with task energy, it changes slope magnitude and can reverse its sign if δ exceeds one.

The numerical displacement scenarios use a common baseline D₀ = 0.60E₀ and a doubled price. They yield measured slopes of {{headline.displacement_fixed_slope:.2f}} when displacement stays fixed, {{headline.displacement_half_slope:.2f}} when it falls by half the gross-energy decline, and {{headline.displacement_more_slope:.2f}} when it falls by 1.5 times that decline. All scenarios keep D and measured energy E − D nonnegative. Together, these scenarios map how displacement calibration shapes the measured slope and highlight the value of paired control conditions for future joint-measurement tests.

A practical design can compare matched blocks of trials across states and map the encoding population together with its surround. With stakes held fixed, a pure change in p at fixed e and K additionally satisfies (σ₁/σ₀)⁴ = (E₀/E₁)², an overidentifying restriction violated by technology changes that alter e/K². Under these conditions either side estimates the budget-price ratio p₁/p₀; this equals μ₁/μ₀ when β is stable.

### S9.4. Simulation-based design assessment

The behavioral layer is calibrated to published summary statistics from numerosity estimation and discrimination tasks [[pratcarrabin2026]]. The simulations assess a prospective within-subject design with two states. In these simulations, β = 1 and e and K are fixed across states, so the per-signal price ratio λ₁/λ₀ equals the budget-price ratio p₁/p₀ and the shadow-price ratio μ₁/μ₀. Representation noise therefore changes by f = (λ₁/λ₀)^(¼) = (p₁/p₀)^(¼) = (μ₁/μ₀)^(¼). Rejection rates are compared across sample sizes, effective per-signal price ratios and task formats, with uncertainty bars conveying Monte Carlo precision.

Across the simulated designs, rejection rates rise with larger state effects and samples, while Wilson intervals show Monte Carlo precision. The estimation marker identifies a distinct task and trial design, and the no-change series supplies a design-specific reference for the changed-price curves.

For discrimination, each state supplies 200 trials. Under the illustrative heterogeneity and lapse assumptions in Methods, a price doubling is detected in {{headline.power_disc_ratio2.0_N20:.2f}} (95% Wilson interval {{headline.power_disc_ratio2.0_N20_mc95_lo:.2f}}–{{headline.power_disc_ratio2.0_N20_mc95_hi:.2f}}), {{headline.power_disc_ratio2.0_N40:.2f}} ({{headline.power_disc_ratio2.0_N40_mc95_lo:.2f}}–{{headline.power_disc_ratio2.0_N40_mc95_hi:.2f}}) and {{headline.power_disc_ratio2.0_N80:.2f}} ({{headline.power_disc_ratio2.0_N80_mc95_lo:.2f}}–{{headline.power_disc_ratio2.0_N80_mc95_hi:.2f}}) of simulated studies with 20, 40 and 80 participants, respectively. At 80 participants, a 50% price increase gives a rejection rate of {{headline.power_disc_ratio1.5_N80:.2f}} ({{headline.power_disc_ratio1.5_N80_mc95_lo:.2f}}–{{headline.power_disc_ratio1.5_N80_mc95_hi:.2f}}), compared with {{headline.power_disc_ratio1.25_N80:.2f}} ({{headline.power_disc_ratio1.25_N80_mc95_lo:.2f}}–{{headline.power_disc_ratio1.25_N80_mc95_hi:.2f}}) for a 25% increase. Under no price change, estimated rejection rates at N = 20, 40 and 80 are {{headline.power_disc_ratio1.0_N20:.2f}} ({{headline.power_disc_ratio1.0_N20_mc95_lo:.2f}}–{{headline.power_disc_ratio1.0_N20_mc95_hi:.2f}}), {{headline.power_disc_ratio1.0_N40:.2f}} ({{headline.power_disc_ratio1.0_N40_mc95_lo:.2f}}–{{headline.power_disc_ratio1.0_N40_mc95_hi:.2f}}) and {{headline.power_disc_ratio1.0_N80:.3f}} ({{headline.power_disc_ratio1.0_N80_mc95_lo:.2f}}–{{headline.power_disc_ratio1.0_N80_mc95_hi:.2f}}); the intervals quantify Monte Carlo precision around these scenario-specific rejection estimates. The fitted state factor's largest absolute mean bias across price conditions is {{headline.disc_max_abs_bias_log_f:.2f}} log units. In estimation, 120 trials per state and 40 participants yield a rejection rate of {{headline.power_est_ratio2.0_N40:.2f}} (95% Wilson interval {{headline.power_est_ratio2.0_N40_mc95_lo:.2f}}–{{headline.power_est_ratio2.0_N40_mc95_hi:.2f}}) for a doubled price; the median log-factor bias is {{headline.bias_est_median_ratio2.0:.2f}}. Across these simulations, optimizer convergence and boundary contact are recorded for every participant rather than inferred from an arbitrary parameter threshold.

The lapse analysis identifies a consequential design feature. When lapse odds rise by 50% in the second state while the price is unchanged, fitting a shared lapse rate yields {{lapse_false_positive_intervals}}. With a doubled price and state-specific lapse rates, simulated rejection is {{lapse_power_intervals}}. Each lapse condition uses {{config.lapse_robustness.reps}} replications; Wilson intervals in Table S6 show the Monte Carlo precision. The 100-replication estimation recovery gives {{estimation_null_calibration}}. The estimation fits succeed in {{headline.estimation_fit_success_ratio1:.1%}} of cases at a unit price ratio and {{headline.estimation_fit_success_ratio2:.1%}} at a doubled ratio; full convergence and boundary diagnostics appear in Table S6. {{lapse_fit_diagnostics_main}} Together, the design projections show how lapse specification changes detectability and identify joint lapse and precision measurement as a useful feature of future tests.

## S10. Calibration pilot of the participant-cluster bootstrap

To assess the finite-sample calibration of the centered participant-cluster bootstrap used in Sections S6 to S8, we simulated 100 cohorts under a conditional Gaussian null (model/conditional_null_calibration.py, seed 20261019). Each simulated cohort resamples observed participant profiles within the recorded sequence strata, retains each participant's design and support across the 21 numerosities and three prior widths, and draws within-cell residual sums of squares from scaled chi-square distributions whose width-specific scales make the expected pooled residual variance equal across widths, so that every one of the 63 contrasts satisfies the null. Each cohort was analyzed with the procedure of the reanalysis, with 10,000 bootstrap draws per test, plus-one p-values and Bonferroni correction over the 63 tests.

The procedure rejected at least one contrast in 22 of the 100 cohorts (familywise rate 0.22; Wilson 95% interval 0.150 to 0.311), against a nominal 0.05, and the mean unadjusted rejection rate across tests was 0.091 (cohort-level 95% interval 0.082 to 0.100), against a nominal 0.05. A balanced two-width toy validation with 300 cohorts and 2,000 draws per test gave an unadjusted rejection rate of 0.063 (Wilson 95% interval 0.041 to 0.097), close to the nominal level. The pilot therefore indicates that the procedure is liberal in the unbalanced design of the published cohort, although its size and its single null do not establish general finite-sample properties. For this reason, the bootstrap intervals and adjusted p-values in Sections S6 to S8 are treated as nominal diagnostics, and the main text relies on the point estimates and on the comparison with the calibrated model. The pilot summary, the test-level results and the toy validation are provided in model/outputs/conditional_null_calibration.
