## Supplementary material for "The marginal brain": Replication package: typeset.py

```
"""Typographic pass on the filled manuscript: true minus signs before numbers and powers of ten for e-notation."""
import re, sys
SUP = str.maketrans('0123456789-', '⁰¹²³⁴⁵⁶⁷⁸⁹⁻')
t = open(sys.argv[1], encoding='utf-8').read()
def sci(m):
    mant, exp = m.group(1), str(int(m.group(2)))
    return (f'{mant} × 10' if mant != '1' else '10') + ('-' + exp).translate(SUP)
t = re.sub(r'\b(\d(?:\.\d+)?)e-(\d+)\b', sci, t)
t = re.sub(r'(?<=[\s(])-(?=\d)', '−', t)
open(sys.argv[2], 'w', encoding='utf-8').write(t)
```
