## Supplementary material for "The marginal brain": Replication package: README.md

# The marginal brain: reproduction package

This reproduction package accompanies the theoretical analysis by Fabio Lokwani-Di Matteo and Pier Luigi Sacco. It contains the model, analytic derivations, configuration, behavioral calibration, simulation code, figure scripts, tests, and the manuscript source.

## Requirements

- Python 3.10 or newer with the packages listed in `model/requirements.txt`.
- LuaLaTeX or XeLaTeX (MiKTeX or TeX Live) to rebuild the manuscript PDFs from the supplied TeX sources; the build uses the standard article class and falls back to XeLaTeX when LuaLaTeX cannot load the OpenType fonts.

The primary numerical campaign's runtime configuration is recorded in `model/results.json`. For a close numerical reproduction, use that recorded Python, NumPy, and SciPy environment; `requirements.txt` lists the minimum libraries required by the scripts.

Install the Python requirements with:

```powershell
py -3 -m pip install -r model/requirements.txt
```

On systems without the Windows `py` launcher, use the active Python executable instead. If it is not on `PATH`, set `REPRO_PYTHON` to its full path before running the build script.

## Reproduce the analysis

From this directory, run:

```powershell
Push-Location model
py -3 run_all.py
py -3 tests/test_model.py
Pop-Location
```

To rebuild the numerical results, the matched-numerosity participant-clustered reanalysis, the pair-specific matched-support sensitivity, the participant-level sign-balance sensitivity, all six figures, both TeX sources and both PDFs, run `reproduce.ps1` in PowerShell or `sh reproduce.sh` in a POSIX shell. The supplied `paper/Manuscript.tex` and `paper/Supplementary_Materials.tex` compile from the `paper/` directory; `paper/build/compile_tex.ps1` and `paper/build/compile_tex.sh` provide platform-specific commands. The PDFs use standard LaTeX output with embedded title/author metadata; this MiKTeX build does not currently support the tagging option required for tagged-PDF output. Primary numerical values are filled from `model/results.json`; matched-support and sign-balance sensitivity values in the main text and supplementary tables are filled from their separately recorded JSON outputs. The participant-level sign analysis uses the same cohort and tests a distinct directional estimand from the primary pooled residual-variance ratio. Seeds and design settings are recorded with their corresponding outputs.

The POSIX script supports staged runs: `sh reproduce.sh paper` rebuilds figures, filled tables, TeX sources and PDFs from the bundled JSON result files without rerunning the numerical campaign; `sh reproduce.sh model` reruns the numerical analyses and their tests; and `sh reproduce.sh all` (the default) runs both stages. The Windows `reproduce.ps1` runs the complete workflow. The recorded primary campaign took about 5,473 seconds in its recorded environment; runtime will vary by machine. The `paper` stage is the appropriate route for rebuilding the presentation from the frozen outputs, while a full numerical rerun should be scheduled separately.

The estimation-recovery simulation is cached in independently seeded chunks. A chunk is reused only when its fingerprint matches the active source, configuration, calibration and numerical environment; a final partial chunk is included when the requested replication count is not divisible by the chunk size. The slower trial-level bootstrap is cached against a fingerprint of its analysis source, input data, settings and numerical runtime, so unchanged reruns do not repeat those draws.

`model/refresh_derived.py` is a compatibility entry point for the same complete workflow; `model/run_all.py` is the canonical command.

## Package map

- `model/`: closed-form model, numerical validation, behavioral calibration, matched-numerosity participant-clustered reanalysis of the published trial data, simulations, tests and generated results.
- data/: source information and setup instructions for the estimation-task trial records; the raw CSV is obtained from OSF and is not redistributed.

A separate conditional-null calibration pilot is included under `model/outputs/conditional_null_calibration/`, with its script and command in `model/conditional_null_calibration.py`; it is an explicit model-conditional diagnostic, not part of the default run. The manuscript reports its calibration warning once in the Limitations subsection. The campaign and its secondary sensitivity are reported as separate analyses. The trial-level reanalysis uses the same published cohort and does not identify neural energy use, the shadow price, or an integrated physiological controller. Simulations are design projections under explicit assumptions, not empirical treatment effects.

The software is provided under the MIT License. No new human data were collected. The model calibration uses published summary statistics; reproducing the trial-level reanalysis requires obtaining the released source CSV from the cited OSF project and placing it in data/.


## Package integrity

The results file stores the source fingerprint captured when its numerical campaign ran. The separate package manifest is regenerated at the end of each release workflow and records SHA-256 hashes for every file shipped in the replication archive; the manifest excludes itself from its hash list. The delivery includes the manuscript PDFs and LaTeX sources alongside the code and configuration. The source trial-level CSV must be obtained from the cited OSF project before reproducing the human-data analysis.

## Model-predicted variance ratios

`model/predicted_variance_ratios.py` computes, by deterministic quadrature, the response-variance ratios that the calibrated estimation model predicts at each of the 21 shared numerosities, using only the parameters fitted at numerosity 60. It writes `model/model_predicted_ratios.json`, which the build merges into the manuscript numbers (`fill_numbers.py --extra=...`) and Figure 6 overlays on the observed ratios.
