## Supplementary material for "The marginal brain": Replication package: reproduce.ps1

```
$ErrorActionPreference = 'Stop'
$root = Split-Path -Parent $MyInvocation.MyCommand.Path
$trialData = Join-Path $root 'data\published_estimation.csv'
if (-not (Test-Path -LiteralPath $trialData)) { throw "Trial-level data are not bundled. Obtain published_estimation.csv from https://osf.io/d6k3m/ and place it at $trialData (see data\README.md)." }
$script:PythonExecutable = $env:REPRO_PYTHON
$script:PythonPrefix = @()
if (-not $script:PythonExecutable -and $env:VIRTUAL_ENV) {
    $candidate = Join-Path $env:VIRTUAL_ENV 'Scripts\python.exe'
    if (Test-Path $candidate) { $script:PythonExecutable = $candidate }
}
if (-not $script:PythonExecutable) {
    $launcher = Get-Command py -ErrorAction SilentlyContinue
    if ($launcher) {
        & $launcher.Source -3 -c 'import sys' *> $null
        if ($LASTEXITCODE -eq 0) {
            $script:PythonExecutable = $launcher.Source
            $script:PythonPrefix = @('-3')
        }
    }
}
if (-not $script:PythonExecutable) {
    $candidate = Get-Command python -ErrorAction SilentlyContinue
    if ($candidate -and $candidate.Source -notmatch '\\WindowsApps\\') {
        $script:PythonExecutable = $candidate.Source
    }
}
if (-not $script:PythonExecutable) {
    throw 'Python 3 was not found. Install Python and add it to PATH, activate a virtual environment, or set REPRO_PYTHON to its executable.'
}
function Invoke-Python {
    param([Parameter(ValueFromRemainingArguments = $true)][string[]]$Arguments)
    & $script:PythonExecutable @script:PythonPrefix @Arguments
    if ($LASTEXITCODE -ne 0) { throw "Python failed with exit code ${LASTEXITCODE}: $($Arguments -join ' ')" }
}
Push-Location (Join-Path $root 'model')
try {
    Invoke-Python run_all.py
    Invoke-Python complete_case_sensitivity.py
    Invoke-Python participant_sign_sensitivity.py
    Invoke-Python predicted_variance_ratios.py
} finally { Pop-Location }
Push-Location (Join-Path $root 'paper')
try {
    New-Item -ItemType Directory -Force figures | Out-Null
    Invoke-Python fig_schematic.py
    Invoke-Python fig_common_factor.py
    Invoke-Python fig_rest.py
    Invoke-Python fig6_empirical.py
    Invoke-Python ..\model\fill_numbers.py body.md ..\model\results.json ..\model\complete_case_sensitivity.json ..\model\participant_sign_sensitivity.json --extra=..\model\model_predicted_ratios.json _main_filled_raw.md
    Invoke-Python typeset.py _main_filled_raw.md _main_filled.md
    Invoke-Python build\mathfix.py _main_filled.md _main_manuscript.md
    Invoke-Python ..\model\fill_numbers.py supplement.md ..\model\results.json ..\model\complete_case_sensitivity.json ..\model\participant_sign_sensitivity.json --extra=..\model\model_predicted_ratios.json _supplement_filled_raw.md
    Invoke-Python typeset.py _supplement_filled_raw.md _supplement_filled.md
    Invoke-Python build\mathfix.py _supplement_filled.md _supplement_manuscript.md
} finally { Pop-Location }
& (Join-Path $root 'paper\build\compile_tex.ps1')
if ($LASTEXITCODE -ne 0) { throw "TeX/PDF build failed with exit code $LASTEXITCODE" }
Push-Location (Join-Path $root 'model')
try { Invoke-Python tests\test_model.py; Invoke-Python write_package_manifest.py } finally { Pop-Location }
Write-Output 'Results, figures, TeX sources, PDFs and package hashes regenerated.'
```
