## Supplementary material for "The marginal brain": Replication package: reproduce.sh

```
#!/bin/sh
# Regenerate numerical results, figures, TeX sources, and PDFs.
# Usage: sh reproduce.sh [all|model|paper] (default: all)
set -eu
STAGE=${1:-all}
ROOT=$(CDPATH= cd -- "$(dirname -- "$0")" && pwd)
if [ "$STAGE" = all ] || [ "$STAGE" = model ]; then
  if [ ! -f "$ROOT/data/published_estimation.csv" ]; then
    echo "Trial-level data are not bundled. Obtain published_estimation.csv from https://osf.io/d6k3m/ and place it in $ROOT/data (see data/README.md)." >&2
    exit 2
  fi
fi
PYTHON=${REPRO_PYTHON:-${PYTHON:-python3}}

if [ "$STAGE" = all ] || [ "$STAGE" = model ]; then
  cd "$ROOT/model"
  "$PYTHON" run_all.py
  "$PYTHON" complete_case_sensitivity.py
  "$PYTHON" participant_sign_sensitivity.py
  "$PYTHON" predicted_variance_ratios.py
fi

if [ "$STAGE" = all ] || [ "$STAGE" = paper ]; then
  cd "$ROOT/paper"
  mkdir -p figures
  "$PYTHON" fig_schematic.py
  "$PYTHON" fig_common_factor.py
  "$PYTHON" fig_rest.py
  "$PYTHON" fig6_empirical.py
  "$PYTHON" ../model/fill_numbers.py body.md ../model/results.json ../model/complete_case_sensitivity.json ../model/participant_sign_sensitivity.json --extra=../model/model_predicted_ratios.json _main_filled_raw.md
  "$PYTHON" typeset.py _main_filled_raw.md _main_filled.md
  "$PYTHON" build/mathfix.py _main_filled.md _main_manuscript.md
  "$PYTHON" ../model/fill_numbers.py supplement.md ../model/results.json ../model/complete_case_sensitivity.json ../model/participant_sign_sensitivity.json --extra=../model/model_predicted_ratios.json _supplement_filled_raw.md
  "$PYTHON" typeset.py _supplement_filled_raw.md _supplement_filled.md
  "$PYTHON" build/mathfix.py _supplement_filled.md _supplement_manuscript.md
  sh build/compile_tex.sh
fi

# The full invariant suite includes trial-level analyses and therefore requires
# the external CSV; the paper-only build uses the bundled frozen result files.
if [ "$STAGE" = all ] || [ "$STAGE" = model ]; then
  cd "$ROOT/model"
  "$PYTHON" tests/test_model.py
fi

# Refresh hashes after any model-only, paper-only, or full staged build.
cd "$ROOT"
"$PYTHON" model/write_package_manifest.py
```
