## Supplementary material for "The marginal brain": Replication package: verify.sh

```
#!/bin/sh
# Run the package's model invariants and numerical checks.
set -e
ROOT=$(cd "$(dirname "$0")" && pwd)
if [ ! -f "$ROOT/data/published_estimation.csv" ]; then
  echo "Trial-level source data are not bundled. Download published_estimation.csv from https://osf.io/d6k3m/ and place it in $ROOT/data before running the verification suite (see data/README.md)." >&2
  exit 2
fi
cd "$ROOT/model"
python3 tests/test_model.py
```
