## Supplementary figures and images for "The marginal brain"

### fig1_schematic.png

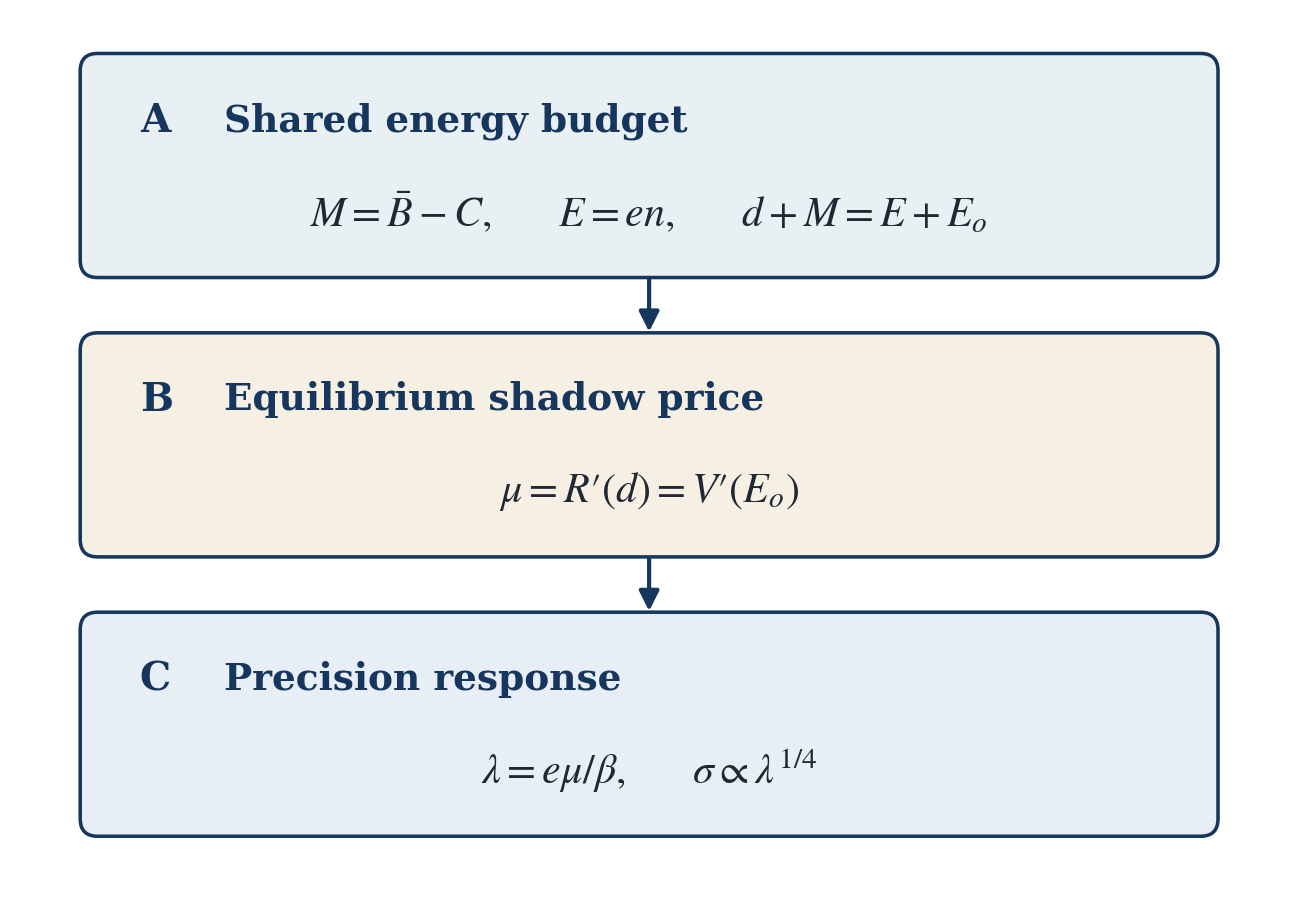

### fig1_signatures.pdf

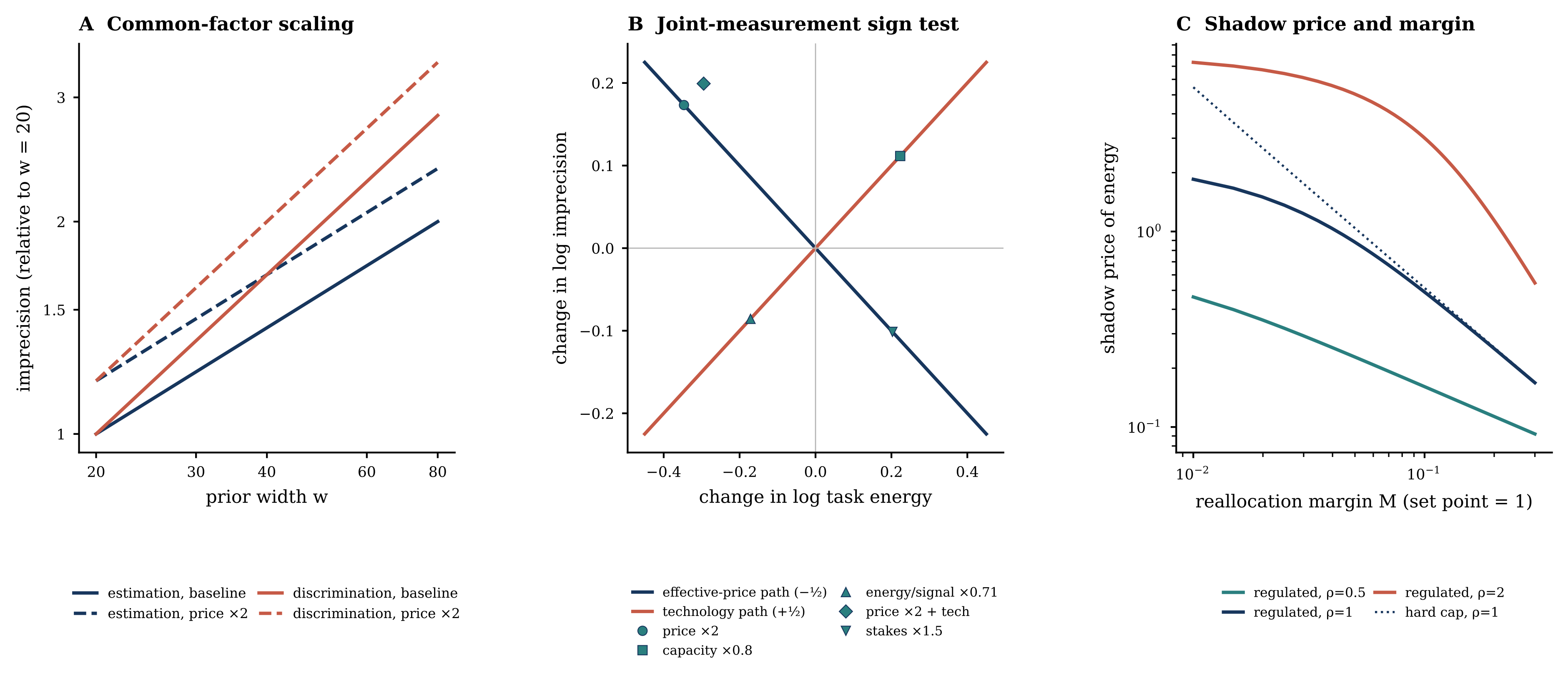

### fig1_signatures.png

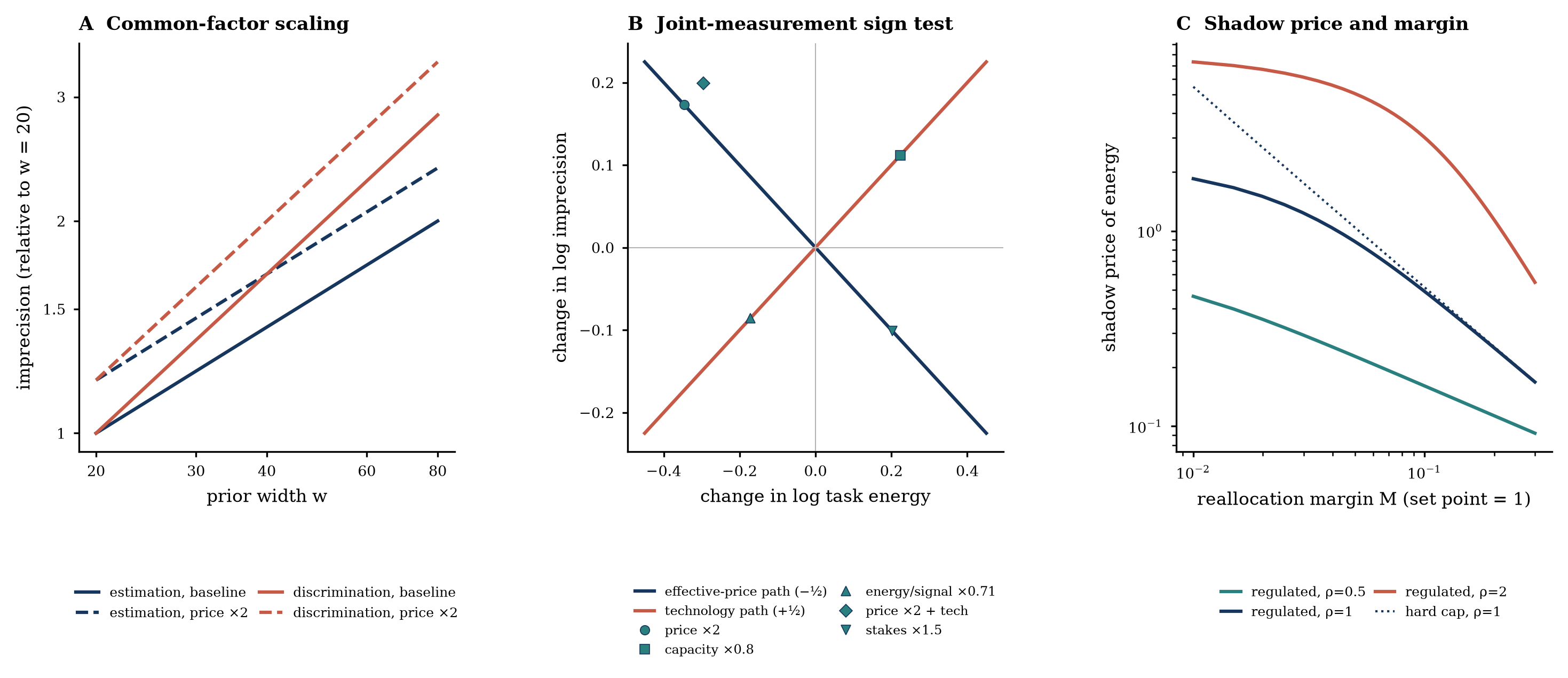

### fig2_common_factor.pdf

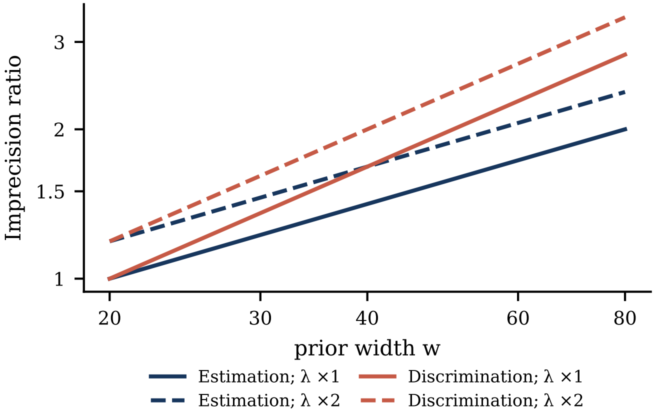

### fig2_common_factor.png

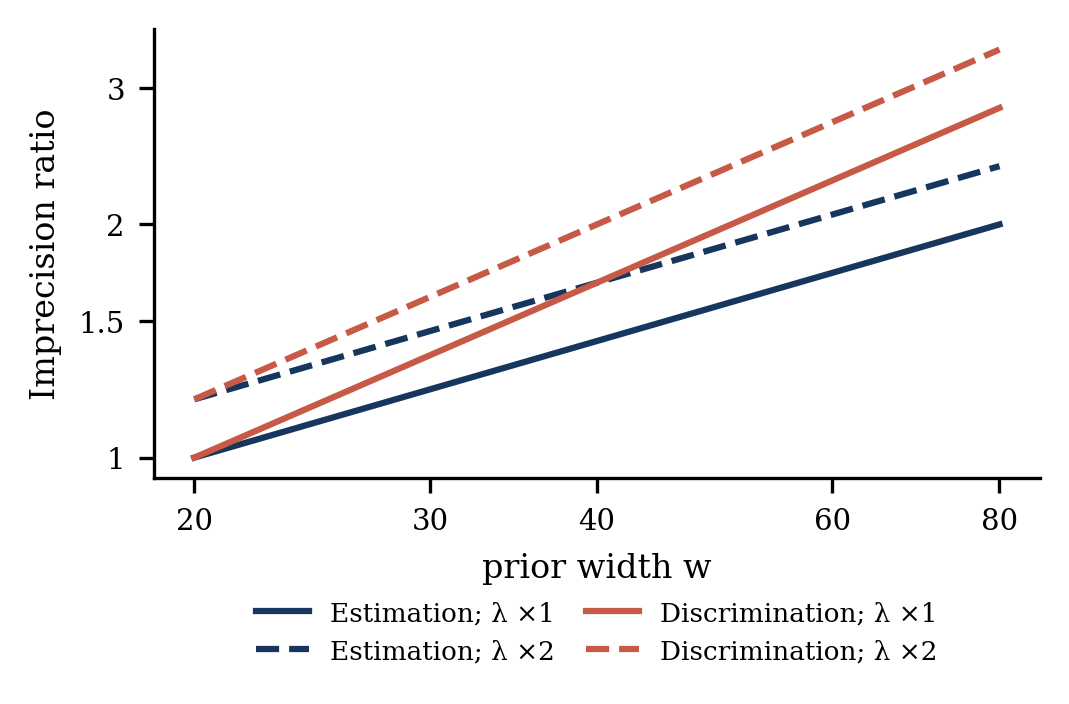

### fig2_power.pdf

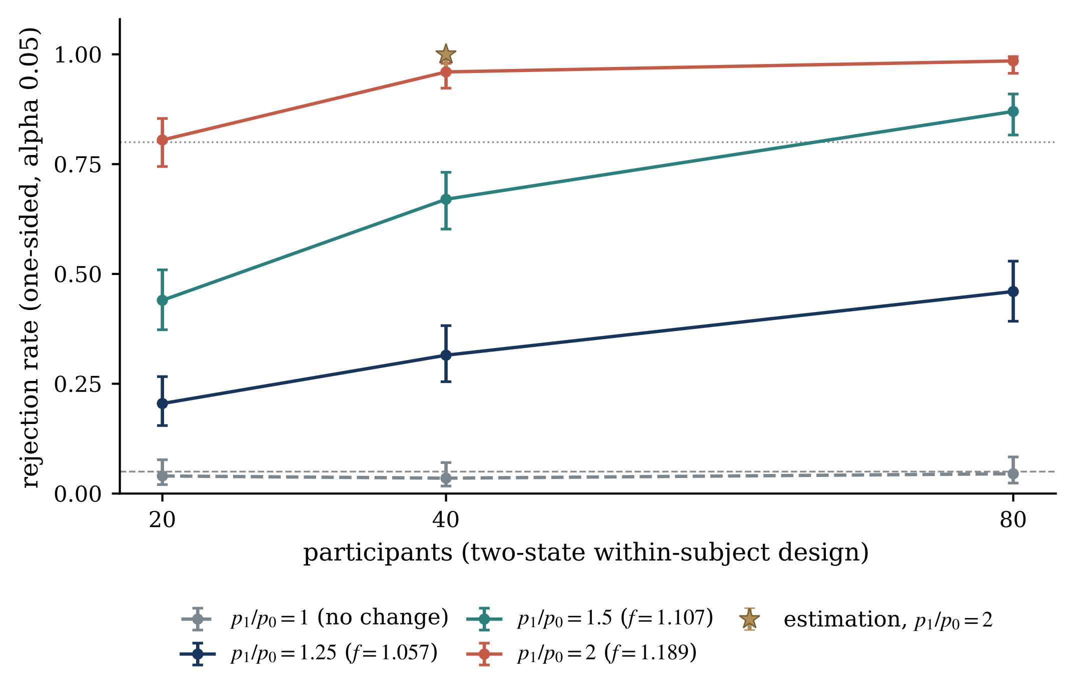

### fig2_power.png

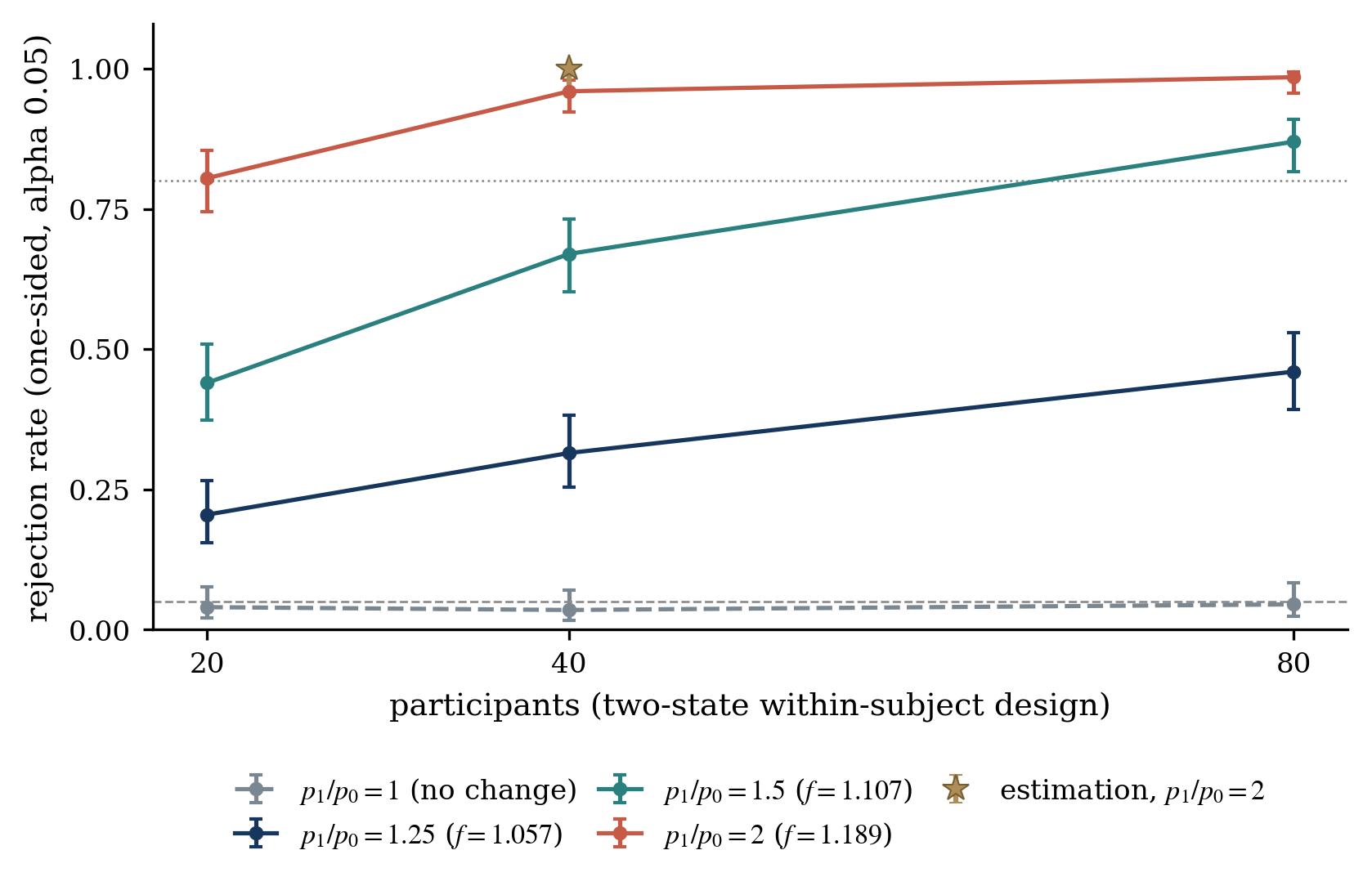

### fig3_budget.pdf

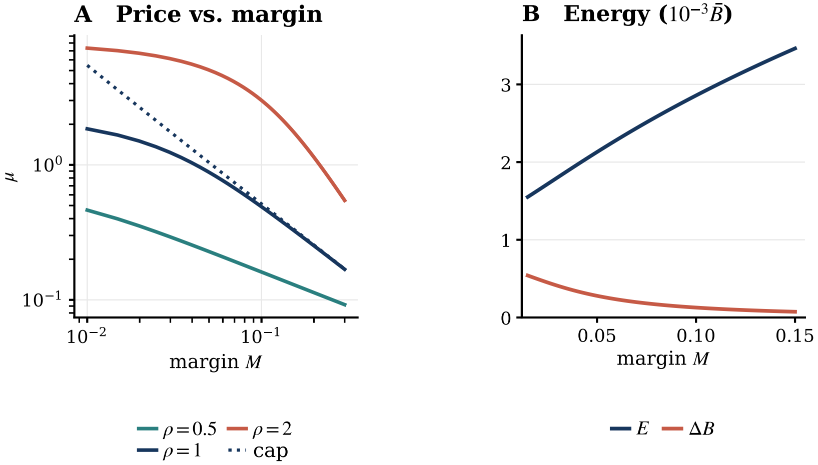

### fig3_budget.png

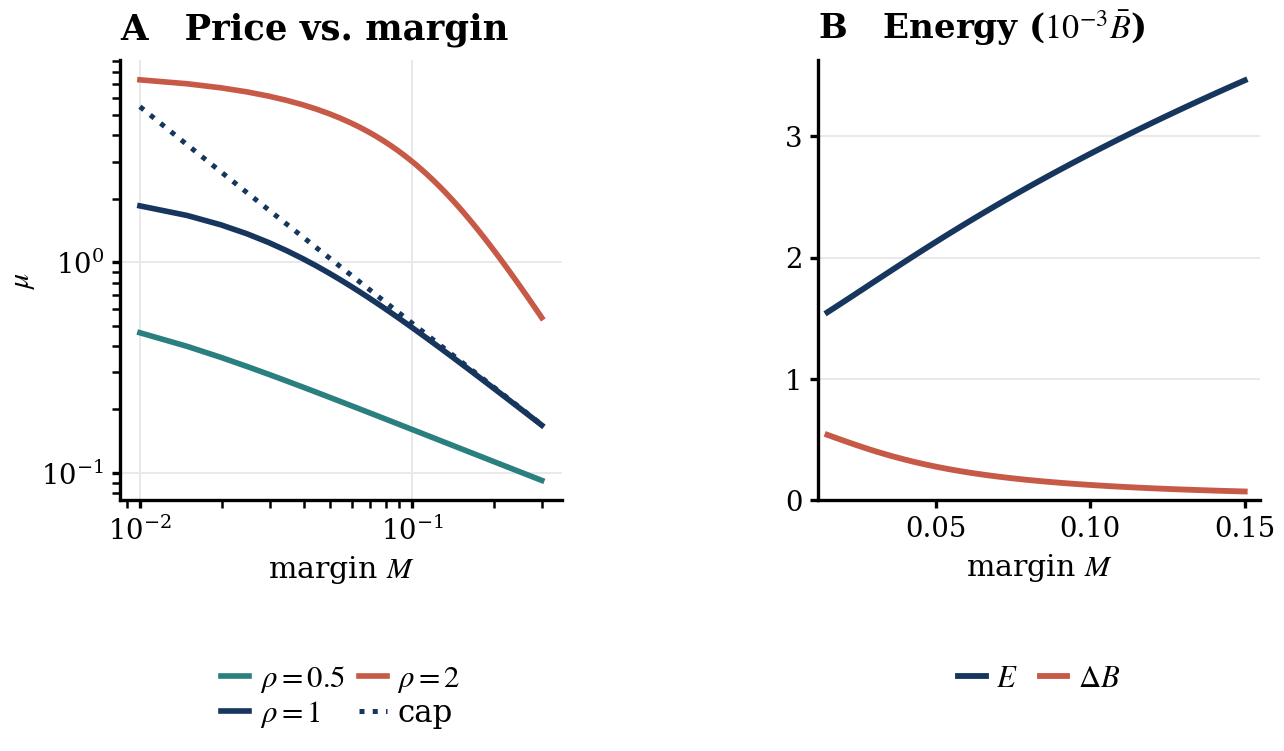

### fig4_sign_test.pdf

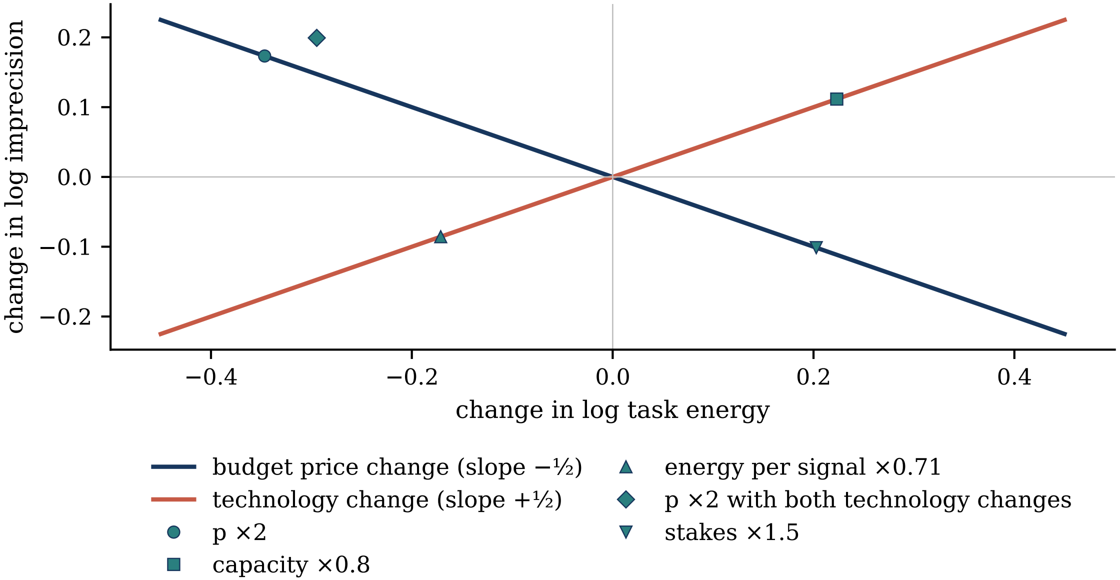

### fig4_sign_test.png

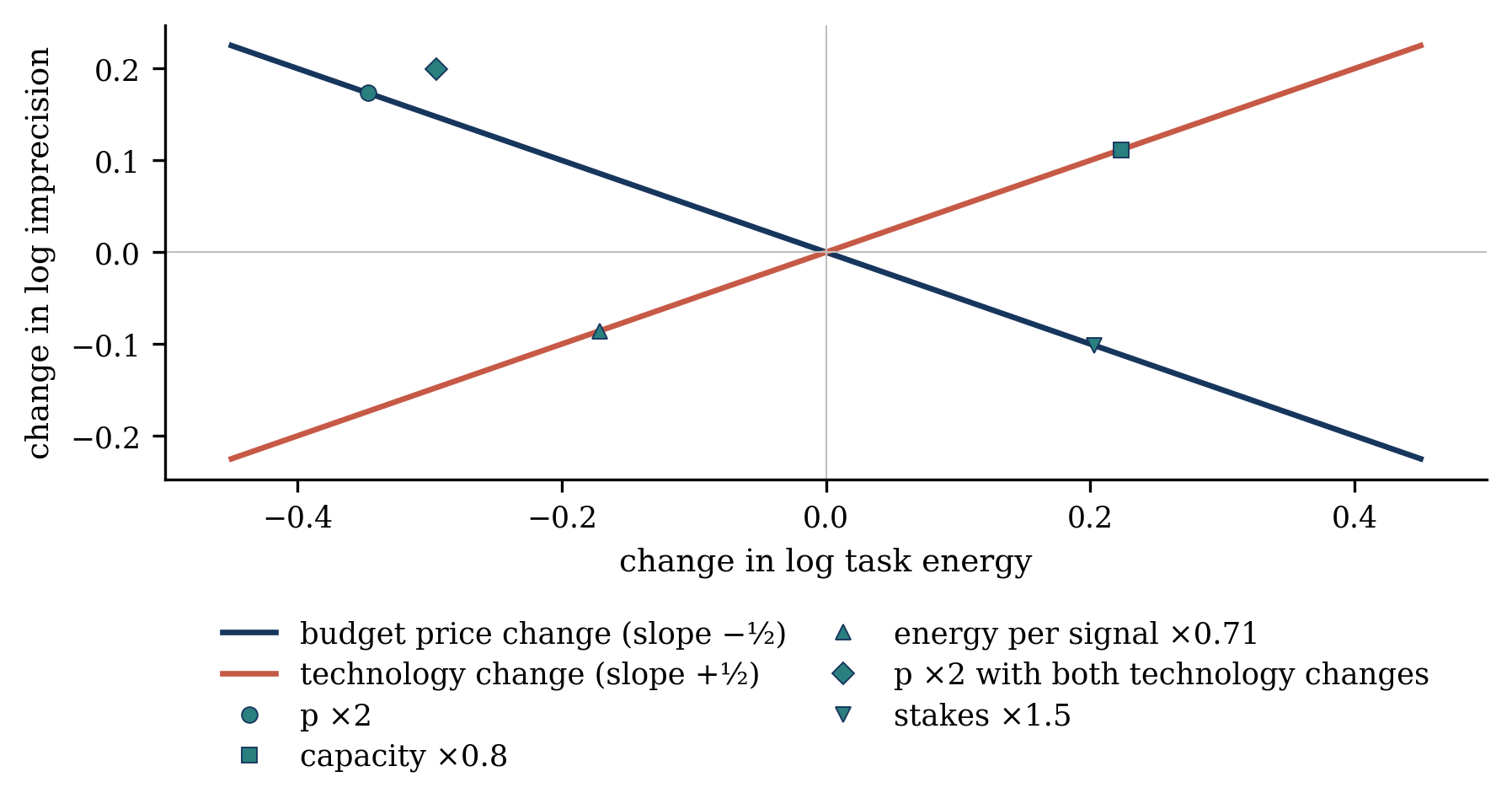

### fig6_empirical.pdf

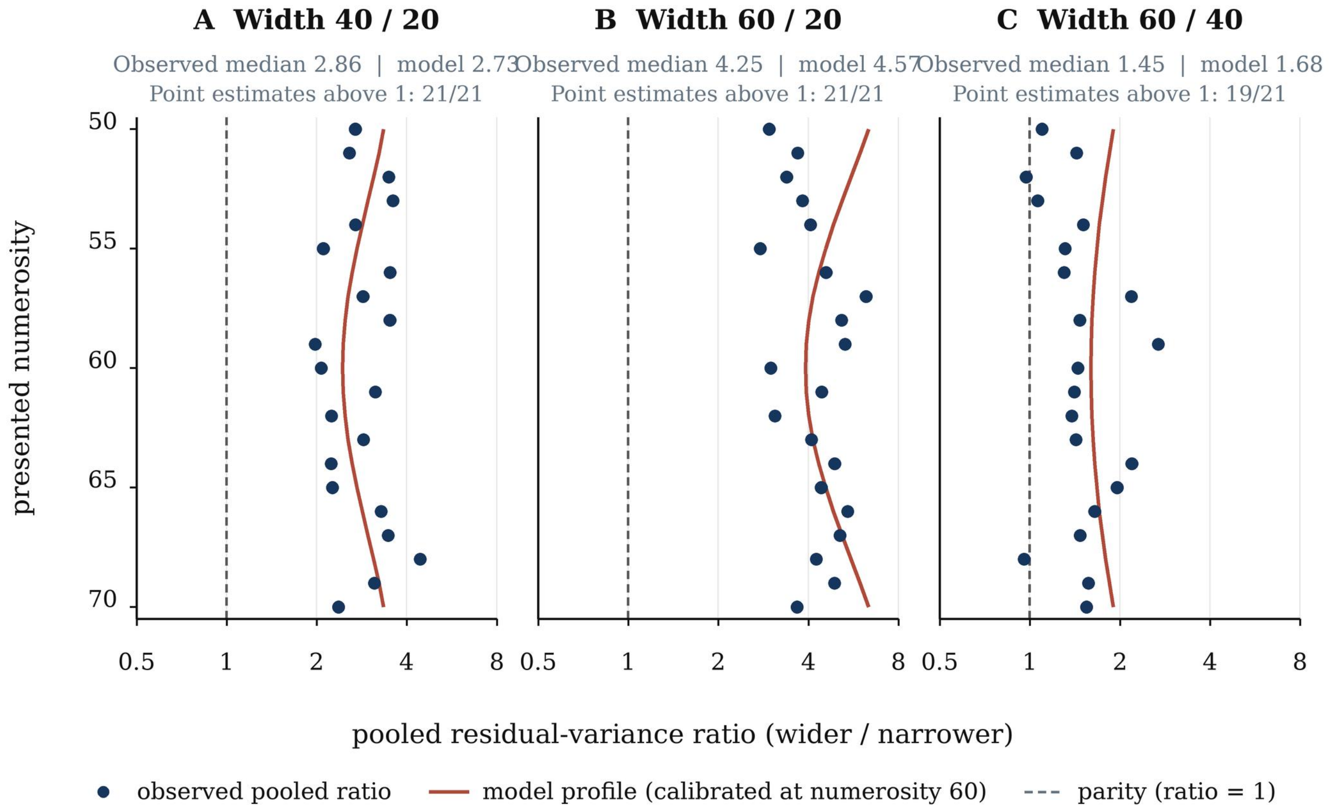

### fig6_empirical.png

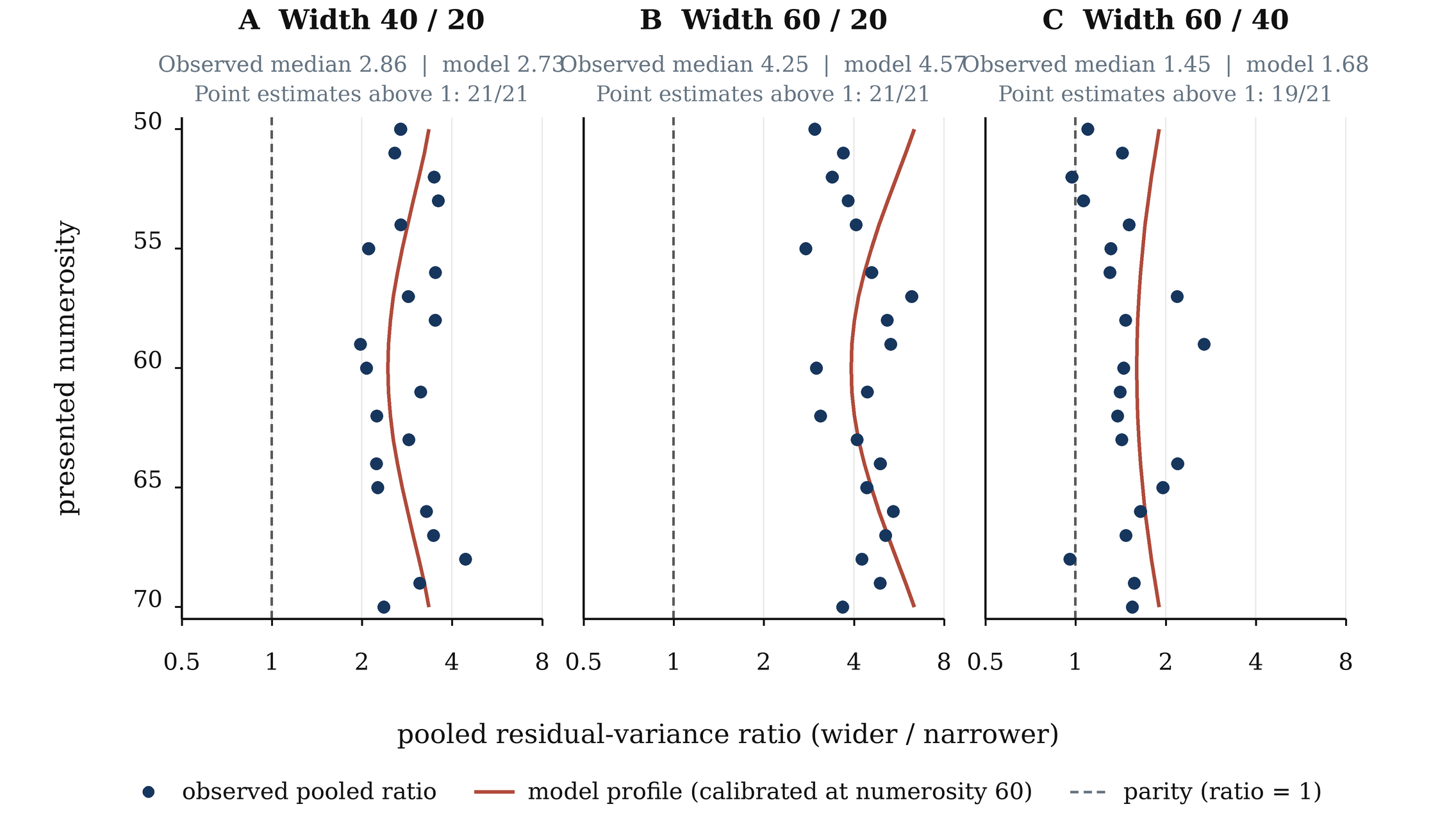
